# COMPASS: In silico reconstruction of single-cell-resolved spatial atlases via cross-omics distillation and coordinate refinement

**DOI:** 10.64898/2026.11.20.768074

**Authors:** Xuanwu Wang, Wenjun Fang, Yuqi Wang, Xingbo Guan, Nuo Li, Yihao Bai, Lixin Liang, Heng Peng, Hua He, Wei Liu, Qishi Dong

## Abstract

The spatial organization of single cells underlies tissue function, yet most spatial omics protocols capture multiple cells per spot. Recovering single-cell coordinates from spot-level data is underdetermined, and existing methods give only cell-type proportions or heuristic placements while rarely using domain structure, geometric priors, and complementary modalities. We introduce COMPASS, a constrained reconstruction framework that resolves this ambiguity by imposing three mutually reinforcing priors on cell-resolved atlas assembly. COMPASS distills auxiliary spatial modalities into an RNA-centered representation deployable to RNA-only dissociated cells, restricts cell-to-spot matching through domain-constrained optimal transport, and refines discrete assignments into dense, non-overlapping coordinates under capacity, locality, and minimum-separation constraints. Across simulations and real datasets, COMPASS outperformed twelve baselines in localization accuracy. In clinical breast cancer, the reconstructed atlas enabled spatial analyses inaccessible to spot-level methods, revealing that compact ER+ tumor nests physically exclude immune cells and suggesting a spatial barrier to immunotherapy.

## Introduction

The spatial organization of cells within a tissue shapes its function, development, and disease progression^1^. Understanding tissue function thus requires resolving both the molecular identity of each cell and its position relative to its neighbors ^2^. Single-cell RNA sequencing (scRNA-seq) profiles cellular heterogeneity at high molecular depth but discards spatial context ^3,4^. Spatial transcriptomics (ST) preserves tissue architecture, yet no single platform delivers whole-transcriptome profiling at single-cell resolution: image-based methods^5–7^ achieve single-cell resolution but are restricted to targeted gene panels, whereas sequencing-based methods ^8–11^ achieve transcriptome-wide profiling but aggregate multiple cells within each spot. Although recent high-resolution sequencing platforms ^12,13^ approach sub-cellular resolution, they remain confined to a few tissues and cohorts, leaving the large existing corpus of spatial data at spot-level resolution. Bridging this gap requires computational frameworks that integrate scRNA-seq and ST data to reconstruct a spatially resolved single-cell atlas from spot-level measurements ^14^.

Resolving precise single-cell coordinates from spot-level measurements is an underdetermined problem: the many-to-one aggregation of cells within each capture spot irreversibly conflates cellular identity with spatial position ^15,16^. Reconstruction becomes well-posed only by imposing tissue-domain structure, geometric priors, and complementary modalities ^17,18^. To assign spatial context to dissociated cells, most existing methods perform spatial deconvolution ^19–26^, decomposing each spot into estimated cell-type proportions. While such methods reveal local composition, they do not yield individual cell coordinates and therefore cannot support analyses that depend on precise cellular positioning. A second class of mapping methods aligns dissociated cells to spots through shared latent spaces ^27,28^, but their outputs remain discrete spot memberships rather than continuous coordinates, leaving sub-spot organization unresolved. These assignments are also driven mainly by transcriptional similarity, with little use of tissue-domain structure, allowing cells with similar expression to be placed in spatially distinct regions. More recently, methods have begun to concentrate on assigning explicit coordinates to individual cells ^29–31^. CellTrek ^29^ and CytoSPACE ^28^ distribute cells within each assigned spot through random or repulsive placement, whereas SIMO^30^ and CMAP^31^ displace cells across neighboring spots using local expression similarity or physics-inspired relaxation. Yet their sub-spot placement remains largely heuristic, without explicitly enforcing geometric priors such as per-spot capacity consistency, minimum inter-cell separation, and confinement to the spot-supported tissue region. Consequently, reconstructed maps may contain spatially implausible arrangements, including overlapping or overly regular cell placements and fragmented local neighborhoods.

Beyond tissue-domain structure and geometric priors, resolving the underdetermined nature of cell mapping also requires complementary information to reduce mapping ambiguity. Recent spatial multi-omics platforms ^32,33^ can jointly profile RNA with auxiliary modalities, such as surface proteins or chromatin accessibility on the same tissue section, providing signals that can sharpen cell-state distinctions otherwise masked by RNA alone. However, leveraging these signals for cell-to-spot mapping faces an inherent asymmetry: spatial measurements may contain multi-omics profiles, whereas the dissociated scRNA-seq reference contains only transcriptomes. RNA thus serves as the anchor modality shared by both sides, and auxiliary information can improve cell mapping only if it is transferred into an RNA-centered representation that is deployable to RNA-only cells. Existing multi-omics approaches are not designed to resolve this asymmetry. General joint models and weighted-neighbor frameworks, including totalVI^34^, MultiVI^35^, Seurat WNN^36^, StabMap ^37^, MOFA+^38^, and scMM ^39^ integrate multi-omics profiles but do not explicitly distill auxiliary-modality information into a shared RNA representation that remains usable on dissociated cells. Spatial multi-omics integration tools, such as SpatialGlue^40^, SpaMI ^41^, and SpaMosaic ^42^, incorporate spatial context to integrate co-measured modalities, but mainly produce fused spot-level embeddings rather than RNA-deployable representations for matching RNA-only dissociated cells to spatial spots. SIMO ^30^ combines multi-omics and spatial mapping, but auxiliary modalities are transferred after RNA-based cell placement, rather than being used to improve the placement itself. Thus, the key challenge is not simply to integrate modalities, but to selectively distill auxiliary-modality-derived information into a shared RNA backbone that can support robust cell-to-spot mapping and spatial atlas reconstruction.

To address these limitations, we developed Cross-Omics distillation, Mapping, and Poisson-disk refinement for Assembly of Single-cell Spatial atlases (COMPASS), a framework for reconstructing single-cell-resolved spatial atlases from spatial omics and dissociated scRNA-seq references. COMPASS learns a shared, domain-aware representation through a multi-objective graph autoencoder and performs domain-constrained optimal transport (OT) followed by capacity-limited assignment, restricting cell-to-spot correspondences to spatially coherent regions. To exploit complementary spatial modalities, COMPASS introduces Residual-Gated Cross-Omics Distillation (RG-COD), which selectively injects auxiliary signals through a learnable residual gate and distills auxiliary-derived domain structure into an RNA-centered backbone for dissociated-cell mapping. To resolve sub-spot positions, COMPASS introduces Poisson-disk Geometric Refinement (PDGR), which converts discrete cell-to-spot assignments into continuous single-cell coordinates while enforcing per-spot capacity consistency, adaptive minimum-separation, and confinement to the spot-supported tissue region. We systematically validated COMPASS through extensive simulation benchmarks and across diverse real spatial omics datasets, including a Xenium-validated breast cancer tissue with single-cell ground-truth coordinates, the laminar mouse olfactory bulb, sagittal mouse brain sections requiring joint cross-section reconstruction, a human lymph node spatial multi-omics dataset, and a multi-sample clinical breast cancer cohort spanning ER+ and TNBC subtypes. Across these settings, COMPASS improved spatial localization accuracy, recovered complex local cellular neighborhoods, preserved fine-scale tissue boundaries, and enabled clinically relevant tumor microenvironment analyses. By enabling the reconstruction of spatially resolved single-cell atlases from widely available spot-level and multi-omics datasets, COMPASS expands the utility of existing spatial resources and provides a general framework for studying tissue organization at cellular resolution.

## Results

### Overview of COMPASS

COMPASS reconstructs a cell-resolved spatial atlas by placing each dissociated cell at a continuous sub-spot coordinate (Fig. 1). The framework proceeds through three coordinated steps. First, a domain-aware representation module uses RG-COD to distill auxiliary spatial omics into an RNA-centered backbone, aligning spatial spots and cells in a shared latent space. Second, a mapping module assigns cells to candidate spots by domain-constrained optimal transport followed by capacity-constrained discretization. Third, a geometric refinement module uses PDGR to convert discrete spot assignments into dense, non-overlapping single-cell coordinates.

**Figure 1:**
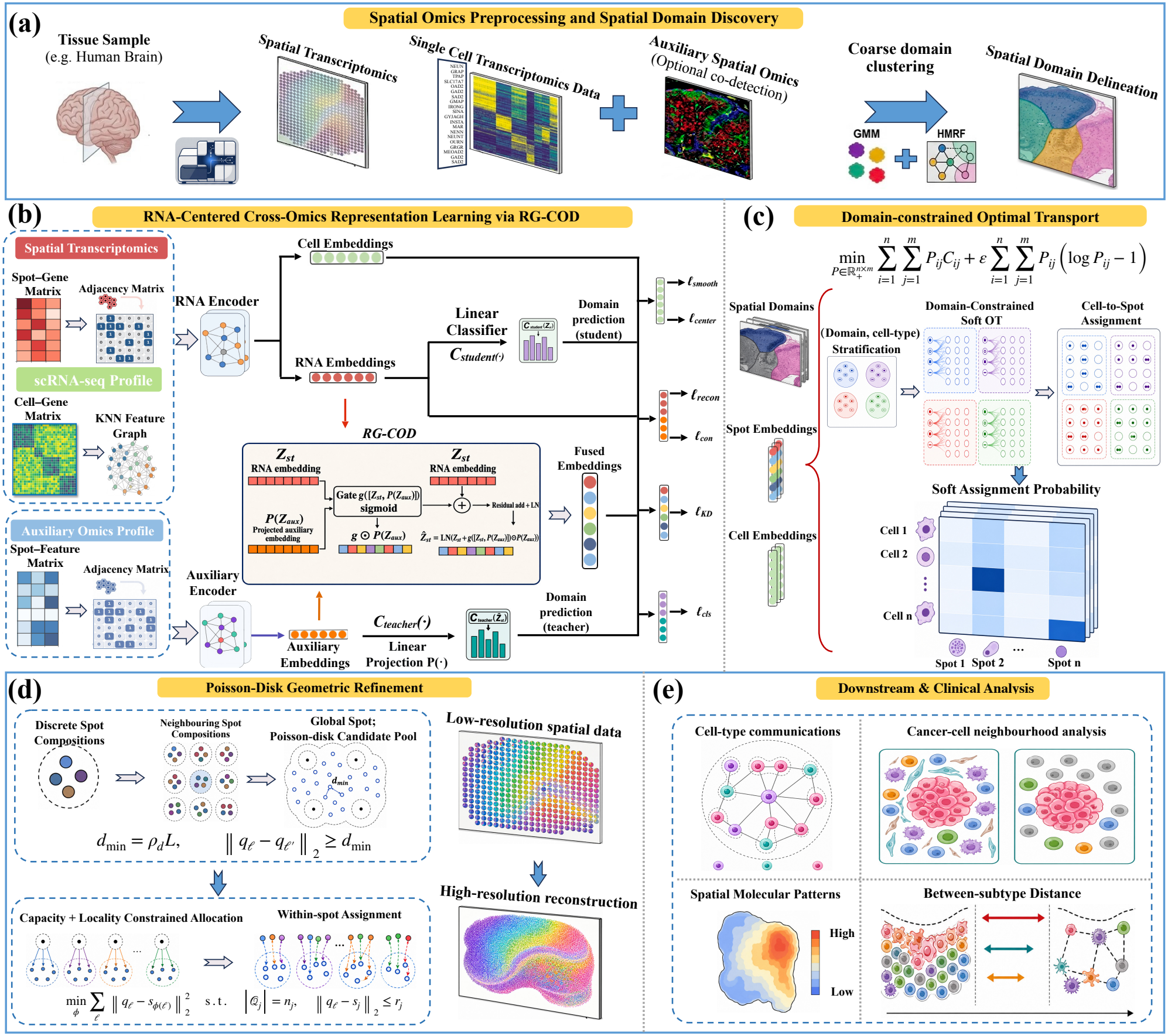
Overview of the COMPASS framework for reconstructing cell-resolved spatial atlases from spatial omics data. (a) COMPASS takes spot-level spatial transcriptomics, a dissociated scRNA-seq reference, and an optional auxiliary spatial omics modality as input. Spatial RNA and auxiliary features are jointly used to infer coarse tissue domains under a Gaussian mixture model with a hidden Markov random field prior. These domains provide structural supervision for representation learning and constrain subsequent cell-to-spot mapping. (b) Spatial spots and dissociated cells are represented on spatial-adjacency and k-NN graphs and encoded by a shared RNA encoder, whereas the auxiliary spatial modality is processed by a separate auxiliary encoder. Residual-Gated Cross-Omics Distillation (RG-COD) projects the auxiliary embedding into the RNA latent space and selectively injects it into the spatial RNA embedding through a learnable residual gate, forming a fused multi-omics teacher representation. Domain classification and knowledge distillation transfer auxiliary-derived domain structure from the teacher to an RNA-only student that remains deployable to dissociated cells. Representation learning is further guided by reconstruction, contrastive, spatial-smoothness, and domain-center objectives, yielding directly comparable spot and cell embeddings. (c) Dissociated cells are stratified according to their predicted spatial domain and annotated cell type, and entropy-regularized optimal transport is solved only between cells and candidate spots within the corresponding domain. The resulting soft cell-to-spot assignment probabilities are discretized under spot-capacity constraints to produce one-cell-to-one-spot assignments while preventing transcriptionally similar cells from being mapped across spatially incompatible tissue regions. (d) Poisson-Disk Geometric Refinement (PDGR) converts the discrete cell-to-spot assignments into continuous sub-spot coordinates. A global Poisson-disk candidate pool is generated under a minimum-separation constraint, followed by capacity- and locality-constrained allocation of candidate coordinates to individual spots. Within each spot, cells are assigned to candidate positions according to local cell-type composition, yielding a dense, non-overlapping single-cell reconstruction while preserving neighborhood continuity across spot boundaries. (e) The reconstructed single-cell spatial atlas enables downstream and clinically relevant analyses that require explicit cell coordinates, including cell-type communication analysis, cancer-cell neighborhood analysis, spatial molecular pattern characterization, and comparison of spatial distributions between clinical subtypes.

To ground learning in tissue structure, COMPASS first infers coarse spatial domains from joint RNA and auxiliary embeddings under a hidden Markov random field (HMRF) prior (Fig. 1a), then encodes each modality with a graph autoencoder over the spatial neighborhood graph. Because dissociated cells contain only RNA, a fully fused multi-omics representation would not be deployable to them at inference. RG-COD resolves the asymmetry by gating projected auxiliary signals into the RNA backbone to form a multi-omics teacher that supervises an RNA-only student (Fig. 1b), so that auxiliary-derived domain structure is selectively distilled into the shared backbone rather than uniformly fused. In the shared latent space, the mapping module stratifies dissociated cells into (domain, cell-type) buckets and restricts candidate spots to the matching domains, then solves OT within each bucket to obtain soft cell-to-spot correspondences (Fig. 1c). The confinement prevents transcriptionally similar cells from being matched across spatially distinct regions, and the soft plan is discretized into a one-cell-to-one-spot assignment.

The assigned spot still leaves the within-spot position undetermined. PDGR therefore refines each cell to a continuous coordinate by Poisson-disk sampling under three geometric priors: per-spot capacity consistency, locality around the spot center, and a minimum inter-cell separation (Fig. 1d). A within-spot assignment then places each cell where its local neighborhood is enriched for its own type, preserving neighborhood continuity across spot boundaries and yielding a dense, non-overlapping single-cell atlas that supports downstream and clinically relevant analyses (Fig. 1e).

### COMPASS stays accurate as perturbations accumulate in simulation benchmarks

To evaluate the spatial reconstruction performance of COMPASS, we designed four progressively challenging simulation scenarios spanning measurement noise in spatial gene expression, spatial RNA signal loss, and missing cell states in the dissociated scRNA-seq reference. Synthetic ST data were generated by aggregating single-cell expression profiles into pseudo-spots according to predefined spatial layouts and cell-type compositions, with auxiliary ATAC profiles co-generated on the same spots to provide a paired multi-omics input. We benchmarked COMPASS against 12 representative baselines for both spot-level composition recovery and cell-level spatial mapping. To study the contribution of multi-omics integration, we additionally compared COMPASS with its single-omics version that only used RNA (COMPASS-single) and the variant with noisy auxiliary omics (COMPASS-noise), which are both detailed in the Methods section.

We first evaluated spot-level reconstruction across the four scenarios (Fig. 2a–d, upper panels). In the unperturbed Scenario 1, COMPASS achieved dominant performance across every reconstruction metric (RMSE = 0.0931, ERGAS = 93.10, UQI = 0.8211, PCC = 0.8542; Fig. 2a; Supplementary Figs S1-3), reducing RMSE by about 5% over the next-best mapping method (CytoSPACE) and improving PCC by over 8% over the next-best deconvolution method (Tangram). Adding measurement noise to the spatial RNA modality while keeping ATAC unperturbed (Scenario 2), COMPASS again led overall (Fig. 2b; Supplementary Figs S4-6), reducing RMSE by 33.1% over COMPASS-single (0.1202 vs. 0.1798), indicating that cross-omics distillation buffers composition recovery against spatial-RNA noise. Scenario 3 further masks 50% of spot-level RNA profiles to zero while retaining the paired ATAC modality. All three COMPASS variants outperformed competing methods by a substantial margin comprehensively (Fig. 2c; Supplementary Figs S7-9), with COMPASS-single (RMSE 0.1687, PCC 0.5791) still surpassing the next-best baseline CytoSPACE (RMSE 0.1741, PCC 0.3987). Notably, the gap between COMPASS and COMPASS-single further widened relative to Scenario 2, suggesting that the auxiliary ATAC modality compensates for missing RNA signal when spatial RNA alone becomes uninformative. In Scenario 4, we introduced additional cell dropout in the scRNA-seq reference to evaluate robustness to reference incompleteness on top of spatial signal corruption. COMPASS regained the top rank (RMSE 0.1439, PCC 0.6828), with COMPASS-noise close behind (Fig. 2d; Supplementary Fig. S10). The gap between COMPASS and COMPASS-single (RMSE 0.1439 vs 0.1950, PCC 0.6828 vs 0.4403) remained substantial, and all three COMPASS variants continued to outperform each baseline across all reconstruction metrics.

**Figure 2:**
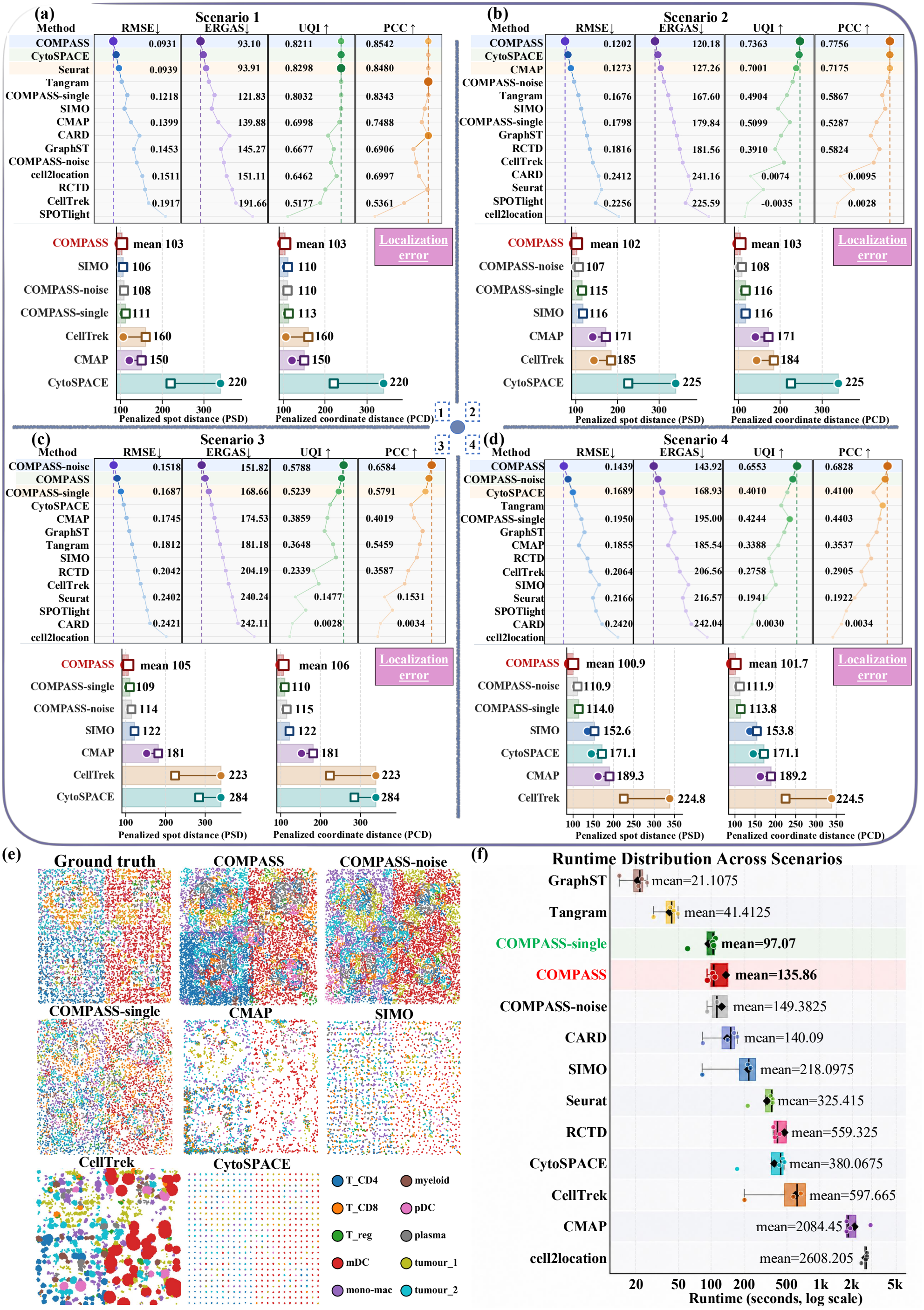
COMPASS maintains accurate single-cell spatial reconstruction under cumulative perturbations. (a)-(d) Benchmarking of COMPASS, COMPASS-single, COMPASS-noise, and competing mapping and deconvolution methods across four simulated spatial multi-omics scenarios generated from paired human melanoma RNA–ATAC data. Scenario 1 uses unperturbed inputs, Scenario 2 introduces spatial RNA noise, Scenario 3 further masks 50% of spot-level RNA profiles, and Scenario 4 additionally removes cell states from the scRNA-seq reference. For each scenario, the upper panels summarize spot-level composition recovery using RMSE, ERGAS, UQI, and PCC, with values averaged over repeated simulation runs. The lower panels summarize cell-level localization errors for methods with coordinate outputs using penalized spot distance (PSD) and penalized coordinate distance (PCD); squares and circles denote the mean and median errors, respectively, with the mean value annotated. (e) Representative ground-truth and reconstructed single-cell maps under Scenario 1, with cells colored by cell type. COMPASS best preserves the simulated tissue organization and local cell-type neighborhoods. (f) Runtime distribution of all benchmarked methods across the four simulation scenarios, with mean runtime annotated on a logarithmic scale.

Since accurate spot-level composition does not by itself guarantee that individual cells are placed at correct positions, we further evaluated cell-level localization error using penalized spot distance (PSD) and penalized coordinate distance (PCD) against ground-truth coordinates (Fig. 2a–d, lower panels). In Scenario 1, COMPASS achieved the highest localization accuracy with the smallest PCD (103), and the reconstructed map faithfully reproduced the spatial organization of the ground truth, including the tumour 1/tumour 2 boundary and T cell–enriched regions (Fig. 2e). By contrast, CMAP produced visually fragmented assignments that disrupted local neighborhood structure (PCD 111). CytoSPACE generated an overly regular lattice pattern (PCD 160), and SIMO yielded sparse, discontinuous reconstructions (PCD 220), which deviate substantially from the ground-truth layout. As perturbations accumulated across Scenarios 2 to 4, COMPASS still maintained consistently low distances (PCD 103–114) and visibly preserved the nested tumour 1/tumour 2 architecture in the reconstructed maps, whereas baselines exhibited distinct degradation modes: CMAP and CellTrek progressively lost cells and fragmented region boundaries, CytoSPACE retained a rigid lattice pattern with weakened cell-type contrast, and SIMO drifted toward noise-dominated layouts (Supplementary Figs S11-14). Deconvolution methods collapsed more sharply under RNA corruption. CARD was reduced to only a handful of predicted spots from Scenario 2 onward, cell2location and Seurat homogenized toward a single dominant cell type, and Tangram/SPOTlight blurred into uniform regional fills (Supplementary Figs S15-18). Across both spot- and cell-level evaluations, COMPASS retained a consistent margin over COMPASS-single, which per-spot diagnostics on Scenario 1 attribute to RG-COD: the residual gate admits ATAC selectively (median contribution ratio 2.43), while the multi-omics teacher and the RNA-only student stay closely aligned, confirming that the auxiliary domain structure is faithfully distilled into the deployable RNA encoder (Supplementary Figs S19-23). However, these gains depend on every module rather than on RG-COD alone, as confirmed by targeted ablations of the five components of COMPASS under Scenario 4, the most challenging setting. Their contributions were nonetheless unequal: removing the domain prior (Supplementary Figs S24-26) or the reconstruction and knowledge-distillation losses (Supplementary Figs S27-29) degraded composition recovery most severely, and weakening RG-COD’s distillation (Supplementary Figs. S30-32) or PDGR’s cell-type-coherent placement (Supplementary Figs. S33-35) caused consistent losses in structural fidelity and localization accuracy, whereas the OT-assignment variants showed the narrowest spread (Supplementary Figs S36-38). No ablated variant matched the full model across all metrics, confirming that the domain-aware representation, cross-omics distillation, and geometric refinement jointly drive the observed gains. Importantly, this multi-module design did not incur a prohibitive computational cost, as COMPASS completed the four simulation settings with a median run time of approximately 106 s (Fig. 2f), despite integrating representation learning, domain-constrained assignment, and geometric coordinate refinement.

### COMPASS enables accurate cell-level localization using paired Visium–Xenium data

To directly assess whether COMPASS assigns each dissociated cell to a spatially accurate coordinate, we leveraged paired 10x Xenium ^7^ and 10x Visium ^8^ data from adjacent serial sections of breast cancer tissue, where Xenium provides independent, cell-resolved spatial coordinates as ground truth. Since the two platforms profile neighboring rather than identical sections, we performed spatial registration following the companion workflow released by 10x Genomics to extract the overlap region, which covered 78% of the full Visium area, and restricted our analysis to this area.

We first evaluated whether individual mapped cells land at biologically correct positions. At the tissue scale, the spatial map reconstructed by COMPASS closely matched the Xenium reference with the lowest minimum distance error (MDE) of 0.16 *µ*m (Fig. 3a; Supplementary Figs S39-S40), preserving clear spatial boundaries and gradual transitions between DCIS 1/DCIS 2, *invasive tumor*, and stromal regions, whereas baseline methods exhibited excessive spatial dispersion (CellTrek and SIMO), over-aggregation of tumor populations (CMAP), and sparse cellular recovery (CytoSPACE). Zoomed-in comparisons further showed that COMPASS preserved local cell-type organization within representative regions, with predicted cells occupying spatially corresponding neighborhoods relative to the Xenium reference (Fig. 3b). This cell-level positional accuracy was quantitatively confirmed by measuring the distance between each mapped cell and its nearest same-type Xenium neighbor (Fig. 3c). Compared with the best-performing baseline, COMPASS reduced the mean and median error distances by 13.20% and 14.88%, respectively.

**Figure 3:**
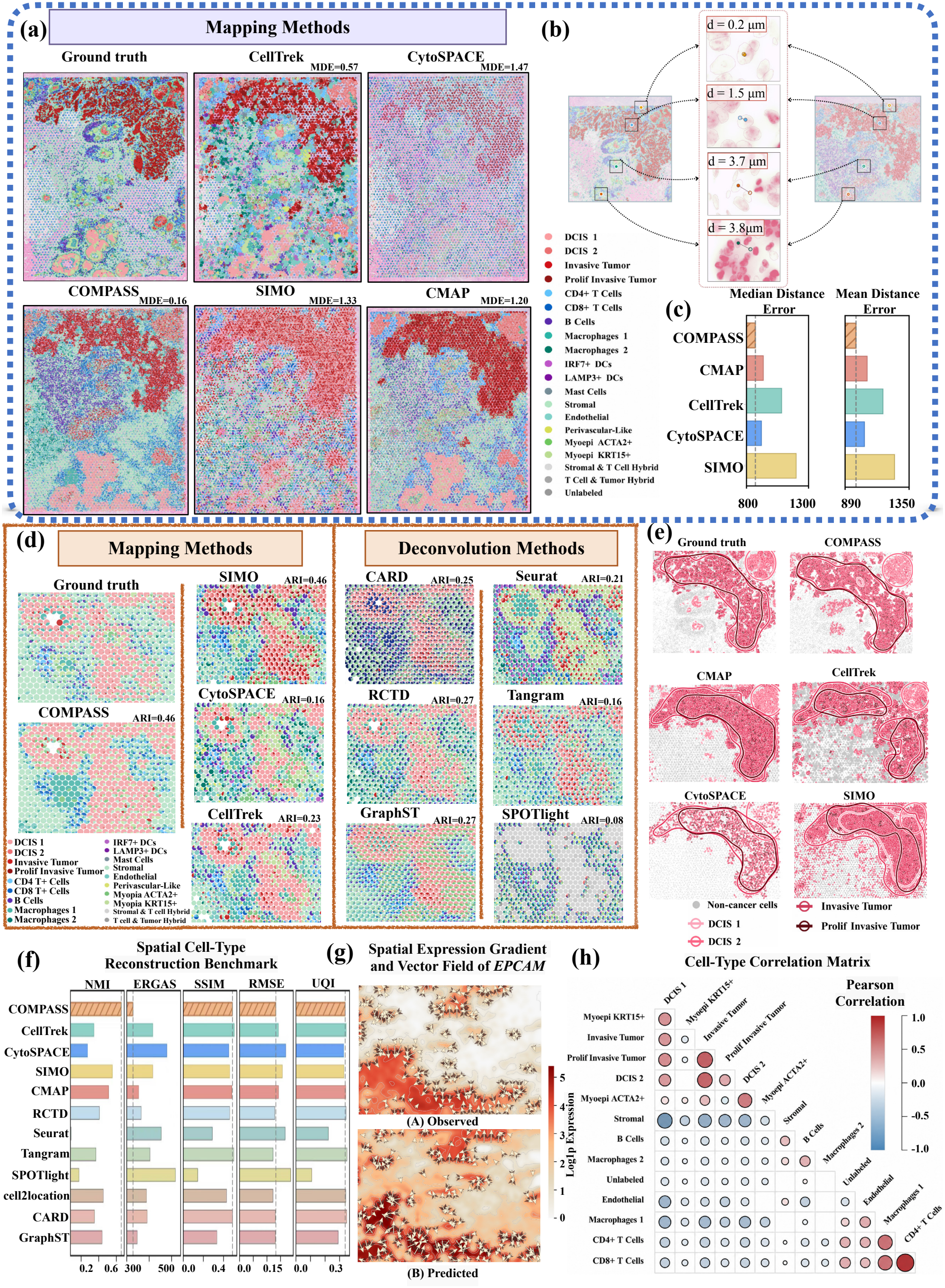
COMPASS assigns spatially precise single-cell coordinates on paired Xenium-Visium breast cancer data. (a) Tissue-scale single-cell maps reconstructed from paired 10x Xenium and 10x Visium breast cancer sections within the registered overlap region, using Xenium cell coordinates as the cell-resolved reference. COMPASS and competing mapping methods are compared against the Xenium ground truth. For each mapping method, the annotated minimum distance error (MDE) denotes the smallest nearest same-type Xenium-neighbor distance among mapped cells, measured in *µ*m. (b) Zoomed-in views of representative regions in (a), showing local cell-position recovery at the neighborhood scale. (c) Cell-level localization accuracy quantified by median and mean distance errors, where the distance error of each mapped cell is defined as the distance to its nearest same-type Xenium neighbor. Lower values indicate more accurate single-cell localization. Spot-level cell-type composition maps obtained by aggregating reconstructed cells onto the Visium grid for mapping methods, or directly using predicted spot-level proportions for deconvolution methods. The ARI value shown above each method quantifies agreement with the Xenium-derived spot-level reference, with higher values indicating better compositional consistency. (e) Spatial distributions of tumor-associated populations, including DCIS 1, DCIS 2, *invasive tumor*, and *proliferative invasive tumor*. (f) Quantitative evaluation of spot-level reconstruction using NMI, ERGAS, SSIM, RMSE, and UQI. (g) Observed and reconstructed spatial expression gradient of the epithelial marker *EPCAM*, with arrows indicating local gradient direction and magnitude. (h) Pairwise Pearson correlation matrices of spot-level cell-type compositions, summarizing recovered spatial relationships among tumor, immune, and stromal populations.

Accurate cell-level localization should yield faithful spot-level cell-type compositions. To test this, we projected the mapped cell atlas onto the Visium spot grid and compared the resulting proportion maps with the Xenium-derived reference, where COMPASS attained the highest ARI of 0.46 (Fig. 3d; Supplementary Fig. S41). COMPASS best recapitulated the spatial distributions and compositional gradients of major populations across tumor, immune, and stromal regions, whereas competing mapping methods introduced scattered mixing (SIMO, CellTrek) and deconvolution methods suffered from oversmoothing (Tangram, SPOTlight). We further examined tumor-specific spatial organization, including DCIS 1, DCIS 2, *invasive tumor*, and *prolif invasive tumor* (Fig. 3e). COMPASS retained the connectivity and boundary morphology of each tumor subclone, with fewer spillovers and boundary shifts relative to the Xenium reference. Across a comprehensive panel of metrics (Fig. 3f; Supplementary Fig. S42), COMPASS showed consistently favorable performance, with high NMI, SSIM, and UQI and low ERGAS and RMSE., confirming that the accuracy of individual cell placements translates into superior recovery of both local composition and tissue-scale spatial structure.

Beyond compositional accuracy, precise cell coordinates should also preserve the spatial relation-ships that underlie biological function. COMPASS reproduced the spatial expression gradient of the epithelial cell adhesion molecule gene (*EPCAM*), a canonical epithelial marker (Fig. 3g), capturing rapid local expression changes at the tumor-stroma interface and further supporting spatial consistency. The reconstructed cell-type correlation matrix also recovered key organizational features of the tumor microenvironment (Fig. 3h; Supplementary Figs S43-44), including the strong spatial co-variation of CD4^+^ and CD8^+^ T cells (*ρ* = 0.912), internal coupling of tumor populations, and the classic spatial segregation between tumor epithelial and stromal components ^43^. Together, the multi-scale evaluations demonstrate that the single-cell atlas reconstructed by COMPASS assigns biologically meaningful and spatially precise coordinates to each mapped cell.

### COMPASS recovers laminar and cross-section architecture in mouse brain tissues

To test whether COMPASS recovers anatomically faithful tissue organization, we evaluated it on two mouse brain datasets that probe two complementary scales of spatial reconstruction, namely intra-section laminar fidelity and cross-section harmonization across slices. We first applied it to the mouse olfactory bulb (MOB) dataset generated by ST v1.0^8^, with a dissociated scRNA-seq reference ^44^ spanning five major cell types. The MOB has a well-defined concentric laminar organization that provides a known anatomical reference, offering a stringent test of intra-section reconstruction. COMPASS produced a dense, spatially complete single-cell map that closely recapitulated the laminar architecture of the MOB^45^ (Fig. 4a). The granule cell layer (GC) formed a continuous central compartment, whereas periglomerular cells (PGC) were enriched along the peripheral glomerular region, consistent with the known concentric organization of the olfactory bulb. By contrast, CMAP produced fragmented tissue boundaries, CellTrek left substantial unmapped regions that disrupted laminar continuity, SIMO introduced pronounced grid-like artifacts, and CytoSPACE yielded an overly sparse reconstruction with incomplete tissue coverage. Quantitatively, COMPASS outperformed all benchmarked methods across assignment quality, spatial coherence, and structural recovery metrics. It achieved the most accurate cell-to-spot assignment, with the highest mean objective and JSD scores alongside the lowest coefficient of variation and entropy (Fig. 4b), while achieving the highest Moran’s I and lowest Geary’s C (Fig. 4c), indicating robust spatial coherence in the reconstructed map.

**Figure 4:**
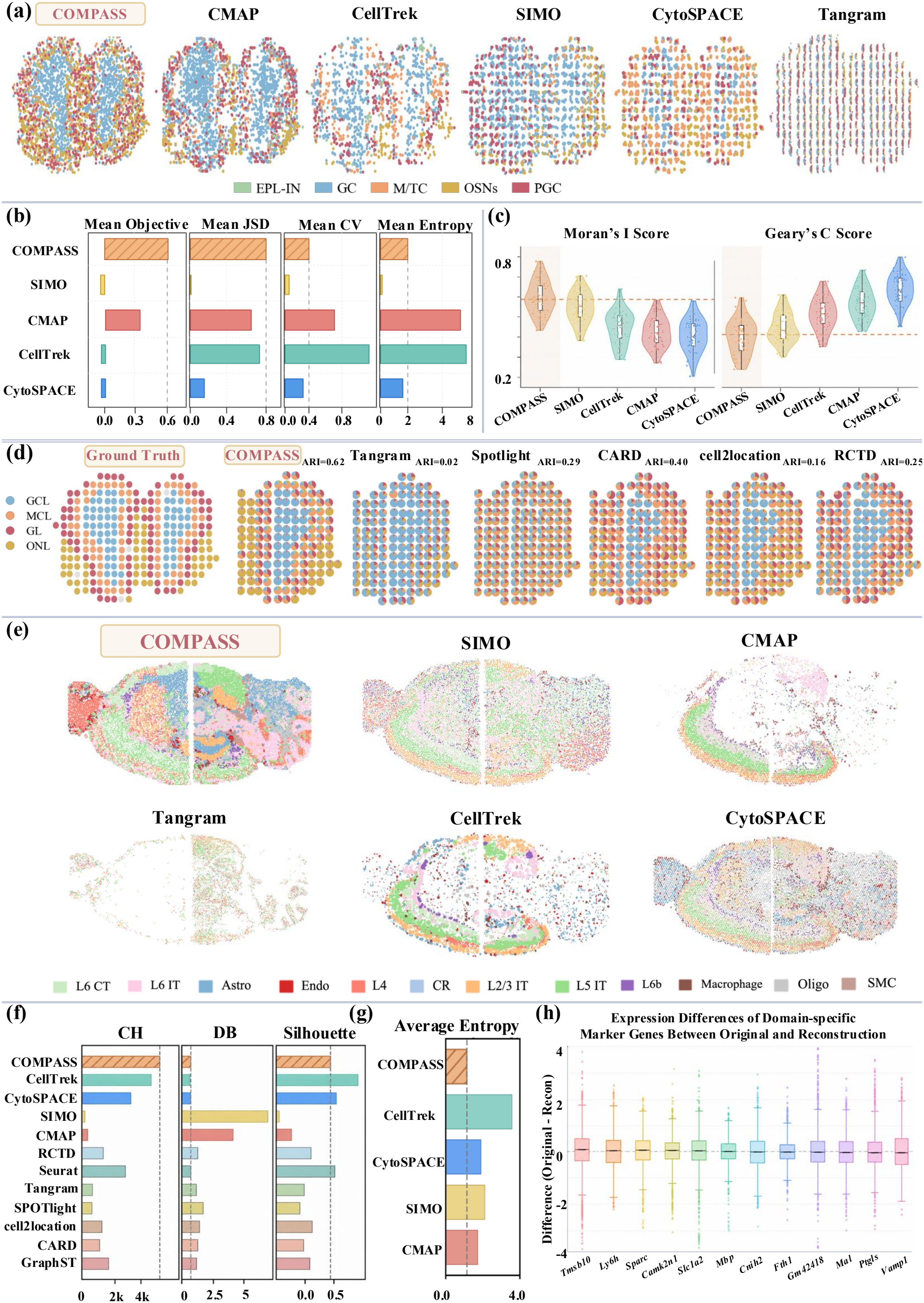
COMPASS recovers laminar and cross-section architecture in mouse brain tissues. (a) Reconstructed single-cell maps of the mouse olfactory bulb (MOB) generated by COMPASS and competing mapping methods, with cells colored by five major cell types. The MOB provides a laminar tissue structure for evaluating the intra-section spatial reconstruction. (b) Spot-level evaluation on the MOB by mean objective score, mean JSD, CV, and entropy. The mean objective score combines two complementary terms, the Jensen–Shannon divergence (JSD) capturing inter-domain separation minus the coefficient of variation (CV) penalizing intra-domain heterogeneity, so that higher values reflect both well-separated and internally homogeneous domains. (c) Spatial autocorrelation of reconstructed MOB maps measured by Moran’s I and Geary’s C. (d) Inferred cell-type proportion maps for the MOB compared with the ground truth and deconvolution methods, colored by laminar layers (GCL, MCL, GL, and ONL); for clarity, only one of the two symmetric bulbs is shown, with the complete section provided in Supplementary Fig. S47. Per-method ARI values against the laminar reference are annotated above each map. COMPASS yields a GCL-dominant center and ONL-enriched periphery with sharper laminar separation than competing methods. (e) Cell-type maps of two jointly reconstructed sagittal mouse brain sections generated by COMPASS and competing methods. (f) Clustering quality of reconstructed sagittal brain sections measured by Silhouette, Calinski–Harabasz (CH), and Davies–Bouldin (DB) scores. (g) Average spot-level entropy across mapping methods. (h) Differences in domain-specific marker-gene expression between original and reconstructed data, summarizing recovery of spatial transcriptional programs.

We further assessed whether COMPASS preserves tissue structure at the spot level by examining inferred cell-type proportions in spatial context (Fig. 4d; Supplementary Fig. S47). This translated into the best agreement with known laminar architecture, with COMPASS ranking first across all clustering metrics (ARI=0.62; Supplementary Figs S45-46). COMPASS yielded more anatomically faithful proportion maps ^45^, with a GCL-dominant center and ONL-enriched periphery that remained more sharply separated than in competing deconvolution methods. In contrast, benchmark baselines showed stronger cross-layer admixture and blurred the boundaries between neighboring laminae. This improved compositional accuracy was further supported by the domain detection results, where both the cluster view and the mapped-cell view produced by COMPASS remained spatially continuous and highly concordant (Supplementary Fig. S48), indicating that anatomically meaningful structure is preserved from cell-level reconstruction to spot-level composition. Consistent with the structural fidelity, the shared RNA latent space underlying these results showed clearly separated spot and cell clusters aligned with the predicted domains (Supplementary Fig. S49).

We next turned to the second scale, jointly reconstructing two sagittal mouse brain sections^46^ profiled by the 10x Genomics Visium platform ^8^. Without a laminar reference for direct comparison, this setting evaluates whether COMPASS can align shared anatomical structures across slices while remaining robust to section-specific variation. COMPASS jointly harmonized the two sections, resolved the major cortical laminae, including L2/3 IT, L5 IT, and L6 populations, and preserved the localized distributions of glial populations such as astrocytes and oligodendrocytes (Fig. 4e; Supplementary Fig. S50). This joint reconstruction was robust to processing choices. The shared UMAP embeddings stayed consistent across domain numbers and under both joint and separate clustering of the two sections (Supplementary Figs S51-52), and the recovered spatial domains remained stable across domain numbers and per-type capacity caps (Supplementary Fig. S53), confirming that the harmonization reflects shared anatomy rather than a specific parameter setting. By contrast, deconvolution-based approaches tended to over-smooth across neighboring structures and blur laminar boundaries, while mapping-based methods exhibited complementary failures, including massive cell dropout (CMAP, CellTrek, Tangram), fragmented layouts (CytoSPACE), and pronounced cross-region mixing (SIMO) (Supplementary Figs S54-55). These qualitative observations were also consistent with the quantitative evaluation. COMPASS achieved competitive Silhouette, Calinski–Harabasz (CH), and Davies–Bouldin (DB) scores, together with the lowest average entropy across spots (Fig. 4f-g), indicating superior spatial coherence, cluster separation, and compositional accuracy. Beyond these structural metrics, expression differences in domain-specific marker genes between the original and reconstructed data remained centered near zero (Fig. 4h), confirming faithful recovery of the underlying spatial transcriptional programs.

### COMPASS integrates spatial multi-omics data in the human lymph node

To study how COMPASS leverages complementary multi-omic information for spatial reconstruction, we applied it to a human lymph node dataset ^24^ profiled by 10x Genomics Visium CytAssist ^8^, which simultaneously captures spatial RNA and antibody-derived tag (ADT) protein measurements on the same tissue section. For the ground-truth reference, we relied on H&E-based tissue annotations that capture major anatomical structures, including the outer capsule, pericapsular adipose tissue, internal cortex, follicle, hilum, and medullary regions. To facilitate cross-method comparison, we grouped mapped cells into five major populations on the basis of their biological identity: B cells, T cells, NK/ILC cells, myeloid cells, and endothelial–stromal cells. Compared with the candidate mapping methods, the annotated tissue architecture was faithfully recovered by COMPASS, which integrates spatial RNA and protein information (Fig. 5a). The reconstructed cell atlas preserved the layered organization of the lymph node, with B cells concentrated in the cortical and follicular interior, T cells enriched in the paracortical zone, and endothelial–stromal cells delineating the peripheral capsule and hilar regions, consistent with the H&E-annotated reference ^47^. The competing methods are restricted to transcriptomic-only input and exhibited two complementary failures. CytoSPACE introduced substantial local mixing between lymphoid and stromal populations, fragmenting the boundary between cortex and medulla, while CellTrek generated overly coarse regional patterns that merged compositionally distinct subregions into uniform blocks. On the other hand, CMAP left noticeable unmapped or weakly assigned areas, disrupting tissue continuity, whereas SIMO achieved low within-region variation but at the cost of assigning highly similar compositions across the entire section, which is inconsistent with the known structural heterogeneity of lymph node tissue.

**Figure 5:**
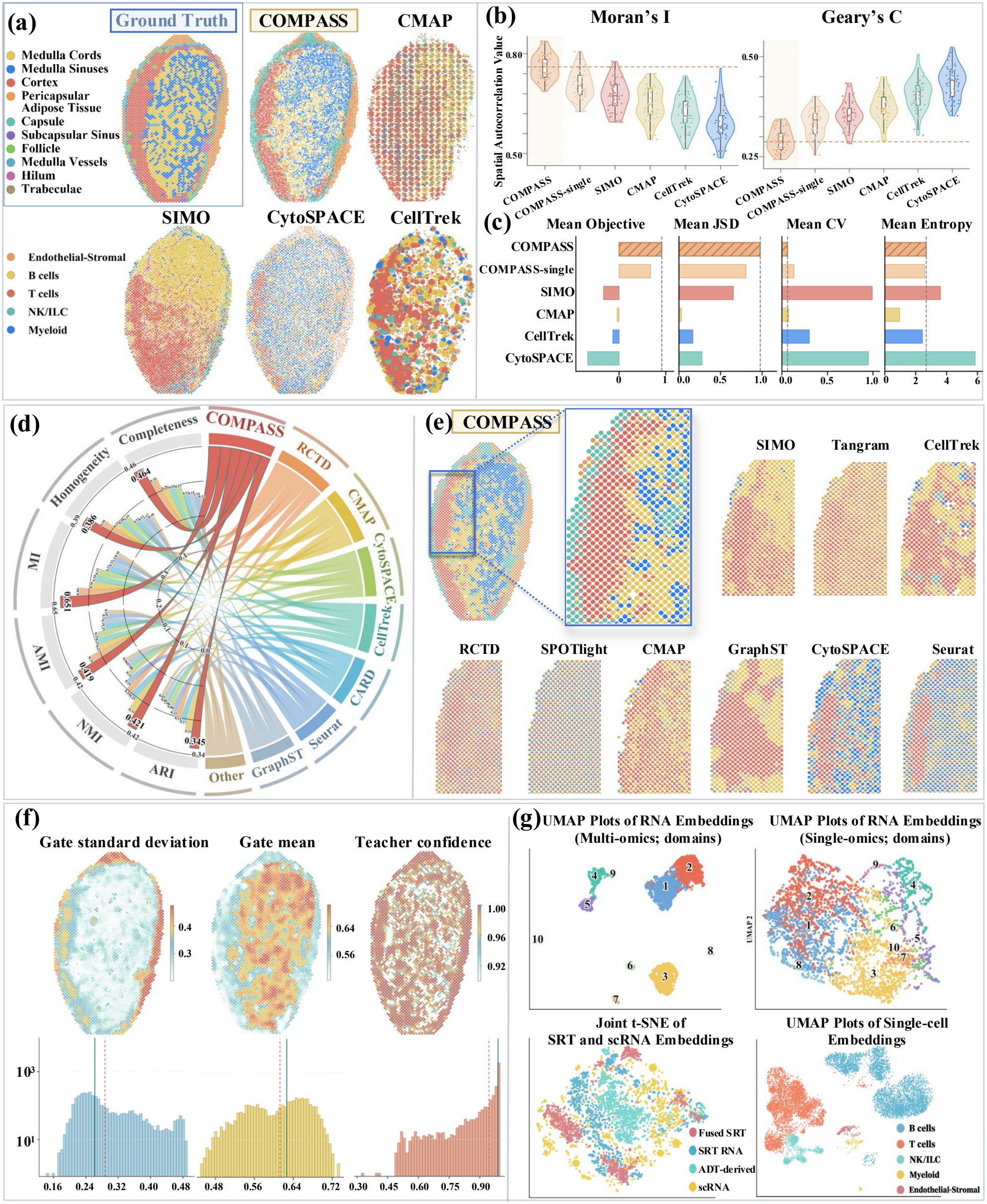
COMPASS integrates spatial RNA and protein measurements to reconstruct the human lymph node. (a) Spatial maps of dominant cell populations reconstructed from a 10x Visium CytAssist human lymph node dataset with paired spatial RNA and ADT protein measurements. Reconstructed maps from COMPASS and competing mapping methods are compared with H&E-based anatomical annotations. Mapped cells are grouped into five major populations: B cells, T cells, NK/ILC cells, myeloid cells, and endothelial–stromal cells. (b) Spatial autocorrelation of reconstructed maps measured by Moran’s I and Geary’s C across COMPASS, COMPASS-single, and competing methods. (c) Spot-level evaluation using mean objective value, mean JSD, mean CV, and mean entropy. (d) Concordance between inferred spatial structure and H&E-annotated regions measured by homogeneity, completeness, MI, AMI, NMI, and ARI. (e) Cell-type composition maps with a magnified region, comparing the spatial organization of major lymph node populations across methods. (f) Per-spot RG-COD diagnostics, including gate standard deviation, gate mean, and teacher confidence, summarize how ADT information is incorporated during multi-omics training. (g) Embedding diagnostics of RG-COD in the human lymph node dataset. The joint t-SNE visualization shows the learned RNA-centered latent space, including the fused SRT teacher embedding, SRT RNA student embedding, projected ADT-derived component, and scRNA embedding. The overlap between SRT and scRNA embeddings indicates that dissociated cells and spatial spots are aligned in a shared manifold, while the separated domain structure supports the selective transfer of ADT-derived information into the deployable RNA encoder.

Quantitatively, COMPASS outperformed all baselines. It achieved the highest Moran’s I and lowest Geary’s C among mapping methods (Fig. 5b), indicating the strongest spatial coherence in the reconstructed layout. At the spot level, COMPASS also attained the best objective score and the strongest inter-domain separation (highest mean JSD), while maintaining the most homogeneous intra-domain cell allocation (lowest mean CV and entropy) (Fig. 5c). Beyond spatial coherence, we further evaluated whether the mapped cells preserved their molecular identity. Canonical lineage markers showed strong and specific enrichment in the expected populations (Supplementary Figs S56-57, and S58(a)), including *CD79A* in B cells ^48^, *CD3D* in T cells ^49,50^, *KLRD1* and *NKG7* in NK/ILC cells ^51,52^, *LILRB2* in myeloid cells, and *IGFBP7* and *CRIP2* in endothelial–stromal cells. The cell-level and spot-level advantages translated into the highest concordance with H&E-annotated anatomical regions across all clustering metrics (Fig. 5d; Supplementary Figs S58(b), and S59-60), and the most spatially coherent composition map (Fig. 5e; Supplementary Fig. S61), with a clear endothelial–stromal peripheral rim and a B-cell-dominant interior compartment.

However, the gains do not reveal whether RG-COD uses the ADT signal as the intended selective residual correction that is faithfully distilled into the deployable RNA-only encoder. Ablation confirms that the protein modality drives part of the improvement (removing ADT reduced the Mean Objective Score from 2.4852 to 1.6854), but performance gains alone cannot establish how that signal is used. We therefore examined the per-spot gate diagnostics (Fig. 5f). The gate admitted ADT at appreciable strength, with a median residual contribution ratio of 5.2, yet modulated it in a domain- and dimension-specific rather than uniform manner, while the fused teacher stayed highly confident and its KL and JS divergences from the RNA-only student remained near zero, with medians of 0.008 and 0.002 (Supplementary Figs S62-64). This agreement persisted even at the spots with the strongest ADT injection, where the teacher relied most on ADT signal unavailable to the student, with matched teacher-student confidence and margin and a near-diagonal confusion matrix (Supplementary Figs S65-66), confirming that the ADT-derived domain structure is faithfully carried into the deployable encoder. The combined effect is evident in the embedding geometry (Fig. 5g), where the inferred domains separate cleanly under multi-omics training and spatial spots and dissociated scRNA share a common manifold.

### COMPASS characterizes the tumor microenvironment in breast cancer

To assess whether the single-cell spatial atlas reconstructed by COMPASS can yield clinically relevant insights into the tumor microenvironment (TME), we applied it to spatial transcriptomics data from four primary breast cancer samples encompassing two major clinical subtypes: ER+ (CID4290, CID4465) and TNBC (CID44971, CID4535) ^53^. These subtypes differ markedly in immune infiltration and immunotherapy response, yet the spatial organization underlying these clinical differences remains poorly characterized at single-cell resolution ^54,55^. COMPASS produced spatially complete and biologically coherent cell maps across all four samples (Fig. 6a; Supplementary Fig. S67). The reconstructed atlases captured the expected compositional asymmetry between subtypes: ER+ samples were overwhelmingly dominated by cancer epithelial cells forming dense, contiguous tumor nests, whereas TNBC samples showed a more balanced representation of immune, stromal, and epithelial populations. Co-localization network analysis further revealed that TNBC harbours more complex inter-cell-type spatial associations than ER+, with extensive connections involving T cells, CAFs, and myeloid cells. By contrast, competing mapping methods failed to recover these subtype-specific spatial patterns (Fig. 6b). CellTrek and CMAP yielded fragmented or incomplete coverage across multiple samples, CytoSPACE introduced local mixing that obscured population boundaries, and SIMO over-homogenized regional compositions.

**Figure 6:**
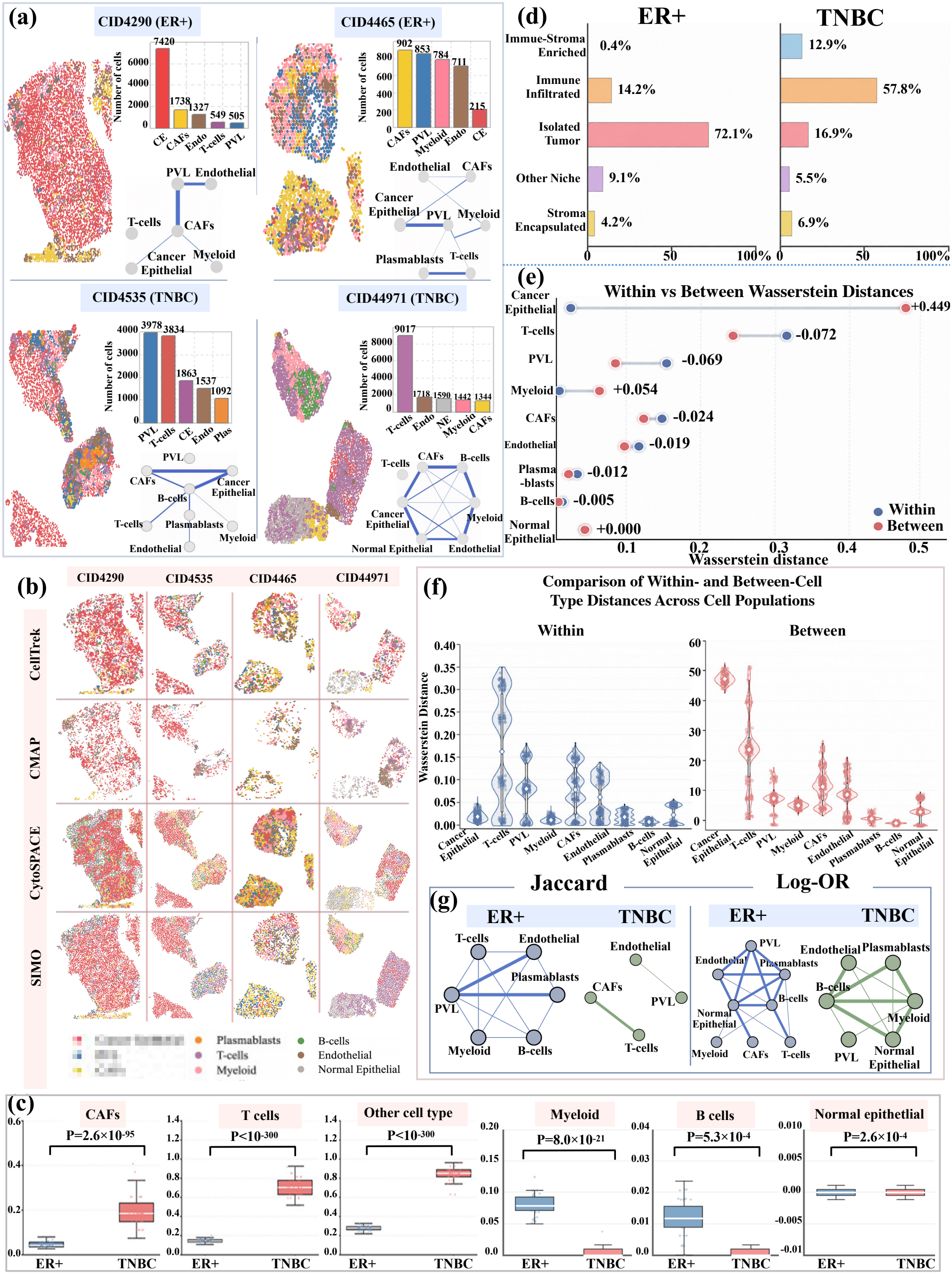
COMPASS reconstructs single-cell spatial atlases that distinguish the ER+ and TNBC breast cancer microenvironment. (a) COMPASS was applied to spatial transcriptomics data from four primary breast cancer samples spanning two clinical subtypes: ER+ (CID4290 and CID4465) and TNBC (CID44971 and CID4535). Reconstructed single-cell maps are shown together with cell-type composition bar plots and co-localization networks, summarizing subtype-specific cellular organization across samples. (b) Spatial maps reconstructed by competing mapping methods across the same four samples, with cells colored by cell type. (c) Local neighborhood composition of cancer cells in ER+ and TNBC samples, quantifying the proximity of cancer cells to CAFs, T cells, myeloid cells, B cells, normal epithelial cells, and other cell populations. Two-sided P values are annotated for subtype comparisons. (d) Tissue-area fractions assigned to spatial niche categories, including immune-stroma enriched, immune infiltrated, isolated tumor, other niche, and stroma encapsulated regions. (e) Within-subtype and between-subtype Wasserstein distances of cell-type spatial distributions across the four breast cancer samples. For each cell population, the annotated value denotes the spatial divergence gap, defined as the between-subtype distance minus the within-subtype distance (Between − Within). Positive values indicate subtype-specific spatial organization, whereas negative values indicate that patient-level spatial variation exceeds subtype-level divergence. (f) Distributions of within- and between-subtype Wasserstein distances across cell populations, summarizing subtype-level and patient-level spatial variation. (g) Boundary-zone interaction networks at tumor margins in ER+ and TNBC samples. Physical co-occurrence is quantified by the Jaccard coefficient, whereas abundance-corrected interaction preference is quantified by the log odds ratio (Log-OR), distinguishing crowding-driven contacts from selective spatial associations

To dissect how these subtype-level compositional differences translate into local cellular contacts, we quantified the local neighborhood composition of cancer cells in each subtype (Fig. 6c). TNBC cancer cells showed significantly greater physical proximity to T cells and CAFs compared with ER+ (*p*-value=2.6 × 10^−55^). Conversely, ER+ cancer cells showed only marginal enrichment for myeloid neighbors (*p*-value=8×10^−11^) and were consistently depleted for T cell and CAF proximity. TNBC cancer cells are extensively exposed to immune and stromal contact, whereas ER+ cancer cells are physically shielded within dense tumor nests, which is consistent with the characterization of ER+tumors as immunologically cold^56,57^. Extending this contact-level view to the tissue scale, spatial niche analysis quantified these architectural differences (Fig. 6d). In TNBC, 57.8% of the tissue area was classified as immune-infiltrated niches, with an additional 12.9% forming immuno-stromal dual-enrichment zones. By contrast, ER+ samples allocated 72.1% of tissue area to isolated tumor niches with minimal immune content, and only 14.2% to immune-infiltrated regions. Such enrichment disparity of immune-infiltrated niches in TNBC relative to ER+ provides a spatial explanation for the well-documented difference in immunotherapy responsiveness between these subtypes ^55^.

Given that both contact- and niche-level analyses point to coordinated spatial divergence, we next asked which specific cell populations drive this divergence by computing pairwise Wasserstein distances of cell-type spatial distributions across all four samples and decomposed them into within-subtype Wasserstein distances (ER+ vs. ER+, TNBC vs. TNBC), and between-subtype Wasserstein distances (ER+ vs. TNBC) (Fig. 6e-f). A positive gap (between minus within) indicates that a cell type is spatially distributed in a subtype-specific manner, whereas a negative gap indicates that patient-to-patient variation dominates over subtype differences. Cancer epithelial cells showed a markedly positive gap (+0.449), indicating that the spatial organization of tumor cells is the primary feature distinguishing ER+ from TNBC tissue architecture. In contrast, most non-cancer populations, including T cells (−0.072), PVL (−0.069), CAFs (−0.024), and endothelial cells (−0.019) showed negative gaps, indicating that their spatial distributions vary more across individual patients. These findings could argue for patient-level spatial profiling rather than subtype-based stratification when assessing immune and stromal architecture for therapeutic decision-making.

The macro-scale analyses above locate each cell type across the tissue, but functional cross-talk is exerted through direct contact, which concentrates at the cancer/non-cancer boundary^58^. There, two measures diverged (Fig. 6g): Physical co-occurrence (Jaccard) inverted the macro-scale co-localization ordering, with ER+ now forming the denser network, whereas abundance-corrected preference (log odds ratio, Log-OR) preserved it, leaving TNBC the stronger. ER+’s denser co-occurrence reflects crowding rather than coordination, because ER+ cancer cells fill the tissue ^56,57^ and compress the scarce immune and stromal cells into narrow gaps, where forced proximity yields indiscriminate contact (Supplementary Fig. S68). Correcting for abundance (Log-OR)^59^ collapsed the ER+ network toward chance, whereas TNBC retained stronger enrichment of interactions among B cells, myeloid cells, and PVL after correcting for abundance. Therefore, the subtypes differ not in how crowded their margins are but in whether the contacts are organized: the TNBC margin is a coordinated immune-stromal interface^53^, whereas the ER+ margin reflects immune exclusion rather than interaction. This organized TNBC boundary is consistent with its greater immune infiltration and immunotherapy responsiveness ^54,55^.

Collectively, these analyses enabled by COMPASS identify the spatial organization of cancer epithelial cells as the primary driving factor distinguishing ER+ and TNBC microenvironments, and suggest that the spatial compactness of ER+ tumor nests represents a potential barrier to effective immunotherapy. Critically, these analyses depend on accurate single-cell spatial coordinates that preserve local neighborhood structure. Spot-level deconvolution estimates cell-type proportions without resolving individual cell positions, and therefore cannot support the adjacency quantification, niche classification, and boundary-zone analyses presented here. The ability of COMPASS to reconstruct spatially faithful single-cell atlases thus enables a class of clinically relevant spatial analyses that may inform therapeutic stratification for breast cancer.

## Discussion

Reconstructing a cell-resolved spatial atlas from spot-level measurements is inherently underdetermined, since each capture spot aggregates multiple cells, irreversibly entangling cellular identity with spatial position. COMPASS addresses this challenge through three coupled modules. RG-COD uses a learnable residual gate and teacher-student distillation to adaptively transfer auxiliary spatial omics into the RNA backbone, providing the complementary information needed to enrich the shared representation. Domain-constrained optimal transport limits matching to spatially coherent domain and cell-type groups, enforcing tissue-domain structure to prevent transcriptionally similar cells from being assigned across spatially distinct regions. PDGR converts discrete spot memberships into continuous coordinates under three biologically grounded geometric priors, capacity consistency, locality, and minimum separation, and further promotes cell-type-coherent placement to preserve neighborhood continuity across spot boundaries. Rather than treating sub-spot reconstruction as a heuristic post-processing step, COMPASS transforms cell-to-coordinate recovery into a principled pipeline jointly enforcing tissue-domain structure, geometric priors, and complementary modalities, providing the structural foundation for the empirical gains observed across our benchmarks.

Across diverse evaluation settings, COMPASS consistently outperformed 12 existing baselines, but the more informative pattern lies in how these gains arise. First, the framework remains robust under compound measurement degradation. In simulations with various noise levels, COMPASS maintained both spot-level composition recovery and cell-level localization accuracy, and the progressively widening gap between COMPASS and its RNA-only ablation indicates that RG-COD acts as a compensatory signal that absorbs auxiliary information precisely when RNA is degraded. On the paired Visium–Xenium breast cancer benchmark, this robustness translated into faithful single-cell placement against cell-resolved ground truth and accurate recovery of tumor boundary morphology, confirming that the reconstructed coordinates correspond to biologically meaningful positions rather than merely improved global metrics. Second, COMPASS recovers tissue architecture that is anatomically interpretable. In the laminar mouse olfactory bulb and the multi-section sagittal mouse brain, COMPASS reproduced known concentric and cortical organization and maintained coherent cellular neighborhoods across spot boundaries and tissue sections, demonstrating that the geometric priors imposed by PDGR generate spatially continuous maps that respect the underlying tissue plan. Third, the reconstructed atlas enables downstream analyses that spot-level methods cannot support. In the human lymph node multi-omics dataset, integrating ADT through RG-COD’s gated transfer improved both reconstruction quality and domain separation, demonstrating selective rather than indiscriminate auxiliary incorporation. In the ER+ and TNBC breast cancer cohort, the cell-resolved atlas supported neighborhood enrichment, niche classification, spatial distribution comparison across tumor subtypes, and tumor–stromal boundary interaction analysis, which require precise single-cell coordinates and cannot be performed on spot-level proportions alone. Together, these results indicate that COMPASS contributes not only quantitative gains in localization accuracy but also a qualitative expansion of the biological questions accessible from spot-level spatial omics data.

Beyond the spatial atlas reconstruction, COMPASS illustrates cross-modal distillation as a deployable-modality-centered paradigm for asymmetric multi-modal integration. When auxiliary modalities are available only on one side of a comparison, transferring their structural information into the shared modality through gated distillation offers a general strategy for asymmetric multi-modal integration, rather than building a fused representation. In addition, COMPASS first learns tissue-domain structure from spatial data, then uses this structure to constrain cell-level matching, and finally imposes geometric priors at the sub-spot scale. Such a design transforms a globally underdetermined reconstruction into a sequence of locally regularized subproblems. This cascading-prior strategy is naturally suited to other spatial inference problems where uncertainty arises simultaneously at tissue, cellular, and subcellular scales.

Currently, COMPASS rests on several modeling assumptions that could be relaxed in future work. First, PDGR derives its geometric priors from spot spacing, which stabilizes sub-spot reconstruction under approximately uniform tissue density but may be less flexible in tissues with strongly hetero-geneous local density, such as densely packed lymphoid follicles or bone marrow. Extending PDGR with density-adaptive priors learned from local tissue context is a natural next step. Second, as a reference-based framework, COMPASS depends on the cell-state coverage and annotation quality of the dissociated scRNA-seq reference. Missing rare populations, transitional states, or mislabeled cells may propagate into the reconstructed atlas, motivating future work on uncertainty-aware mapping and self-correcting reference refinement, particularly for cell states absent from the reference. Third, our current evaluation of RG-COD spans paired RNA–ATAC in simulation and RNA–ADT in the human lymph node, but its behavior across broader multi-omics combinations remains to be systematically examined. As single-cell and spatial omics technologies continue to diversify across modalities, scales, and time dimensions, frameworks that combine principled multi-level priors with adaptive cross-modal representation learning will be essential for building the spatially resolved, multi-modal cell atlases that the next generation of tissue biology requires.

## Method

### Model Specification

COMPASS aims to reconstruct a cell-resolved spatial atlas by assigning and refining each dissociated cell to a continuous sub-spot coordinate (Fig.1). The framework comprises three coupled components: (i) a domain-aware representation module that distills auxiliary spatial omics information into the deployable RNA backbone via RG-COD to align spatial spots and dissociated cells in a shared RNA latent space and; (ii) a domain-constrained mapping module that assigns cells to candidate spots through domain-aware optimal transport followed by capacity-limited assignment; and (iii) a geometric refinement module that converts discrete cell-to-spot assignments into dense, non-overlapping single-cell coordinates via PDGR.

### Representation Learning via Residual-Gated Cross-Omics Distillation

Let *X*_*st*_ *∈* ℝ^*n*×*g*^ denote the spatial transcriptomic matrix for *n* spots and *g* genes, with spot coordinates 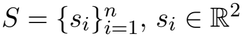. Let *X*_*aux*_ *∈* ℝ^*n*×*p*^ denote an auxiliary spatial omics measured on the same spots. COMPASS aims to learn a deployable RNA representation in which spatial spots and dissociated cells are directly comparable, leveraging *X*_*aux*_ during training while remaining applicable to RNA-only dissociated cells at inference. In our experiments, when only the primary modality is available (COMPASS-single) and no auxiliary spatial omics is measured, we set *X*_*aux*_ = *X*_*st*_, so that the primary modality itself serves as the auxiliary input and the same RG-COD formulation applies without modification. To anchor downstream representation learning in tissue-level structure, COMPASS first infers coarse spatial domains as a structural prior, where a PCA-based clustering embedding *u*_*i*_ is constructed from concatenated RNA and auxiliary embeddings. A Gaussian mixture model (GMM) is fit to 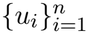 with a hidden Markov random field ^60^ and Potts penalty to encourage locally coherent domain assignments:

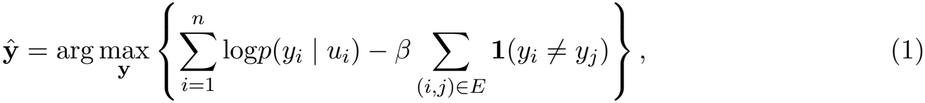

where *E* is the set of edges in the spatial neighborhood graph, and *β >* 0 controls the smoothness strength. The inferred domain labels *y*_*i*_ *∈* {1, …, *K*} obtained from Eqn (1) serve two distinct downstream roles: as supervisory signals in representation learning, and as a structural prior partitioning the cell-to-spot mapping problem.

COMPASS then encodes each spatial omics into a modality-specific embedding space using a two-layer graph encoder. Let 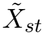 and 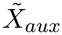 denote the low-dimensional PCA features of spatial RNA and auxiliary omics, respectively. The RNA spot embedding *Z*_*st*_ and the auxiliary spot embedding *Z*_*aux*_ are obtained as

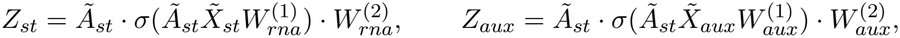

where 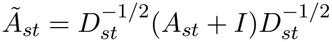 is the shared normalized adjacency matrix, and *σ*(·) denotes the ReLU activation.

To align spatial spots and dissociated cells in a shared RNA latent space, COMPASS needs to address modality asymmetry. Directly learning a fully fused multimodal representation is undesirable because the fused space would depend on modalities that are unavailable for dissociated cells at inference. Instead, we propose a *Residual-Gated Cross-Omics Distillation* (RG-COD) to adaptively distill auxiliary spatial omics into the deployable RNA encoder backbone to enhance spatial representation learning. The resulting distilled RNA backbone is then shared with dissociated cells, enabling both spots and cells to be encoded into a common RNA latent space for downstream comparison and mapping. Specifically, we define the fused embedding 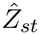 as a multi-omics teacher representation:

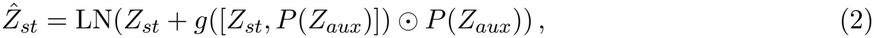

where the auxiliary embedding is projected into the RNA space through the learnable projection *P* (·), *g*(·) is a sigmoid gate, and LN(·) denotes layer normalization. The gate in Eqn (2) is introduced to modulate the contribution of auxiliary omics in a spot-specific manner, so that cross-omics information acts as a selective residual correction to the RNA backbone rather than uniformly overwriting the RNA representation. In parallel, the RNA-only representation *Z*_*st*_ serves as the student representation.

Both teacher and student are supervised by the inferred domain labels **ŷ**, with logits

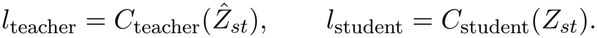

The student is additionally trained to match the softened predictions of the teacher through knowledge distillation, yielding the RG-COD objective

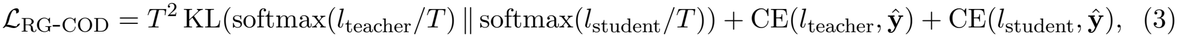

where CE(·, ·) denotes the cross-entropy loss, *T* is the distillation temperature. The full training objective combines *ℒ*_RG-COD_ with omics-specific reconstruction and regularization terms, including spatial smoothness, supervised contrastive, center, and decorrelation regularization (Supplementary Notes, S1). After training, both spatial spots and dissociated scRNA-seq cells are encoded by the same distilled RNA backbone into a shared RNA latent space, yielding spot embeddings *Z*_*st*_ and dissociated-cell embeddings *Z*_*sc*_, with dissociated cells organized on a *k*-nearest-neighbor graph in this aligned space. The shared representation renders *Z*_*st*_ and *Z*_*sc*_ directly comparable, providing the input to the domain-constrained cell-to-spot mapping module.

### Domain-Constrained Cell-to-Spot Mapping

To reconstruct cell-resolved spatial coordinates from the learned embeddings, COMPASS first performs cell-to-spot assignment by solving domain-aware optimal transport problems within each predicted domain *k* and cell-type *t* bucket, thereby preventing transcriptionally similar cells from being matched across spatially incompatible regions. The inferred domain labels **ŷ** from representation learning are propagated to dissociated cells through domain prediction in the shared RNA space, and combined with cell-type annotations from the scRNA-seq reference to define a partition over the joint (domain, cell-type) space. For each (*k, t*) bucket containing *n*_*k,t*_ cells and *m*_*k*_ candidate spots, the imbalance across buckets can collapse minority domains. COMPASS therefore performs quota-based sampling that approximately balances the cell-type and domain marginals while retaining high-confidence cells, with each cell type contributing at most *M*_*max*_ cells (default *M*_*max*_ = 1000), and re-encodes this balanced subset on a *k*-NN graph to produce the refined *Z*_*sc*_ used for transport (Supplementary Notes, S1). The cell-to-spot assignment is obtained by solving an entropy-regularized balanced optimal transport problem ^31,61^:

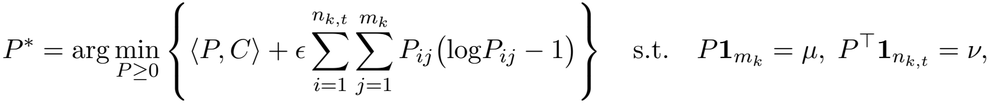

where we define the transport cost by 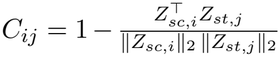 in the shared RNA space, 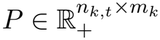 is the transport plan, *µ* and *ν* denote the prescribed cell and spot marginals, and *ϵ >* 0 controls the strength of entropic regularization. Unless an external prior is provided, COMPASS sets *µ*_*i*_ = 1 for every cell and *ν*_*j*_ = *n*_*k,t*_*/m*_*k*_ for every spot within each (domain, cell-type) bucket, so that per-spot cell load arises from the transport plan rather than a user-imposed cap. The resulting soft plan *P*\* is then converted into a discrete one-cell-one-spot assignment by integerizing the spot loads 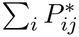 into spot capacities via Hamilton rounding ^62^ and solving a capacity-constrained Hungarian matching, yielding a spot-assignment map *π*(*c*) *∈* {1, …, *n*} for each cell *c*.

### Poisson-Disk Geometric Coordinate Refinement

The discrete spot membership of each cell does not uniquely determine its continuous subspot location, making coordinate reconstruction inherently underdetermined. We assume that cells remain spatially localized around their assigned spot, respect the inferred spot capacity, and maintain a non-overlapping local arrangement. Under the priors, we introduce *Poisson-Disk Geometric Refinement* (PDGR) to convert discrete spot memberships into dense, non-overlapping single-cell coordinates under three biologically plausible geometric priors: *capacity consistency, locality, minimum separation*. Let 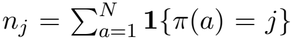denote the number of cells assigned to spot *j*, and J = {*j*: *n*_*j*_ *>* 0} the set of occupied spots. We define the local spot spacing 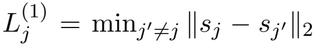 and the global characteristic spacing 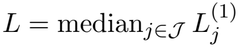. We then derive three scale-dependent parameters:

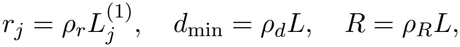

where *r*_*j*_ enforces *locality* by bounding each cell’s deviation from its spot center, *d*_min_ enforces *minimum separation* by setting the exclusion distance between any two reconstructed cells, and the per-spot count *n*_*j*_ enforces *capacity consistency* by fixing the number of coordinates allocated to each spot. Denoting *D*(*c, R*) = {*p ∈* ℝ^2^ : ∥*p −c∥ ≤R}* as the closed disk of radius *R* centered at *c*, the global support region is 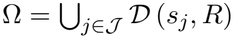.

A pool of candidate coordinates 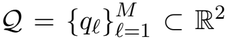 is generated using Bridson’s Poisson-disk sampling ^63^ with exclusion radius *d*_min_. Starting from a random seed *q*_1_, each new candidate is proposed uniformly from the annulus around an active point *q*:

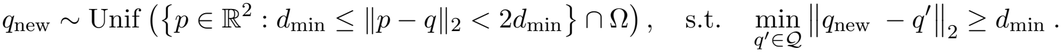

The procedure terminates when no further valid candidates can be proposed, yielding a blue-noise-distributed set satisfying 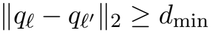 for all *ℓ* ≠ *ℓ*^′^ by construction. By design, *M ≫ N* so that candidates are available in excess of the total cell count. Each candidate coordinate is allocated to an occupied spot via a mapping *ϕ* : {1, …, *M*} → *J* that minimizes total squared displacement subject to per-spot capacity and locality constraints:

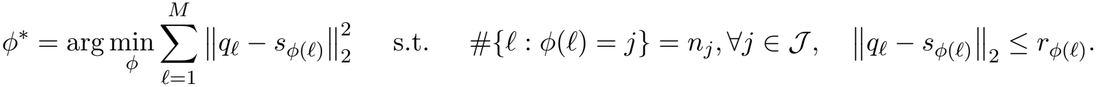

For each occupied spot *j*, this yields an allocated point set Q_*j*_ = {*q*_*ℓ*_ : *ϕ*\*(*ℓ*) = *j*} with |Q_*j*_| = *n*_*j*_. Let *A*_*j*_ = {*a* : *π*(*a*) = *j*} denote the cell set with cell-type labels 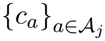. Since |Q_*j*_| = |*A*_*j*_| = *n*_*j*_, a bijection *σ*_*j*_ : *A*_*j*_ → Q_*j*_ is needed to assign each cell a unique coordinate. PDGR solves a local assignment that places each cell at the candidate position whose spatial neighborhood is most enriched for that cell’s type. For each candidate *q*_*ℓ*_ *∈* Q_*j*_, a cell-type affinity vector **f** (*q*_*ℓ*_) *∈* ℝ^|C|^ summarizes the expected composition in the vicinity of *q*_*ℓ*_ using the known compositions of neighboring spots:

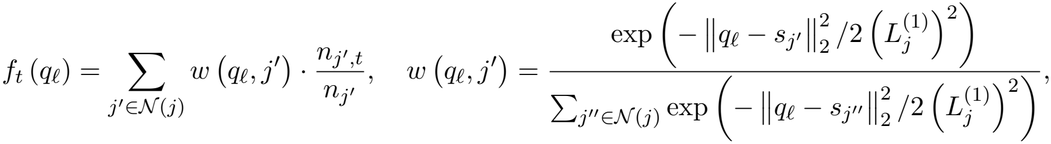

where *N*(*j*) denotes the spatial neighbors of spot *j*, and 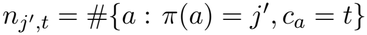 is the number of type-*t* cells assigned to spot *j*^′^. The within-spot assignment is obtained by solving a balanced linear assignment:

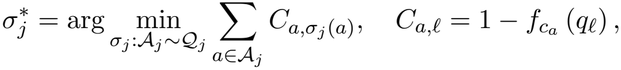

where the cost *C*_*a,ℓ*_ is low when the neighborhood around *q*_*ℓ*_ is enriched for the cell type *c*_*a*_ of cell *a*. The final reconstructed coordinate for each cell *a* is 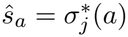, *∀a ∈ A*_*j*_. By construction, {*ŝ*_*a*_} satisfy locality *ŝ*_*a*_ *∈ D s*_*π*(*a*)_, *r*_*π*(*a*)_, capacity consistency (|Q_*j*_| = *n*_*j*_), and minimum separation (*∥ ŝ*_*a*_ − *ŝ*_*b*_*∥*_2_ ≥ *d*_min_). The type-coherent assignment further promotes cross-boundary continuity by placing cells near like-typed neighbors in adjacent spots. Finally, COMPASS produces a unified single-cell spatial atlas 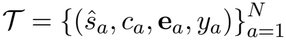, where *ŝ*_*a*_ *∈* ℝ^2^ is the reconstructed spatial coordinate, *c*_*a*_ is the cell-type annotation transferred from the scRNA-seq reference, **e**_*a*_ *∈* ℝ^*g*^ is the original single-cell gene expression profile, and *y*_*a*_ *∈* {1, …, *K*} is the spatial domain label inherited from spot *π*(*a*).

### Model training

COMPASS is trained end-to-end by minimizing a single composite objective. This objective comprises an omics-specific reconstruction term, the domain classification and knowledge distillation terms that jointly form the RG-COD supervision in Eqn (3), and a set of regularizers that promote spatial coherence, identity discrimination, and cross-modal alignment of the learned embeddings. We define each term below and give its weighted combination at the end of this section.

#### Reconstruction loss

Two modality-specific linear decoders are appended to 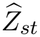 to reconstruct the standardized inputs as 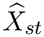 and 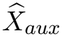, supervised by mean squared error:

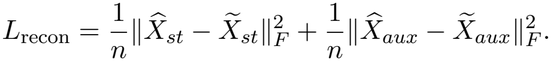

#### Domain classification loss

Both the RNA student and the multi-omics teacher are supervised by the inferred domain labels *y*. The domain classification loss is defined as

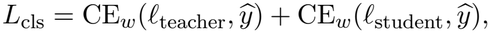

where CE_*w*_ denotes class-balanced cross-entropy.

#### Knowledge distillation loss

To transfer auxiliary-derived domain structure into the deployable RNA backbone, the RNA student is trained to match the temperature-softened predictions of the multi-omics teacher:

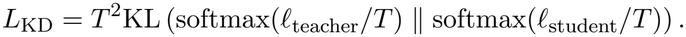

#### Contrastive loss

A projection head maps 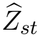 into a contrastive space. Let *ℐ* denote the set of spatial spots used for domain supervision, and let *q*_*i*_ be the normalized projected embedding of spot *i*. For each anchor *i*, define its positive set as

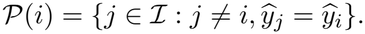

The supervised contrastive loss is

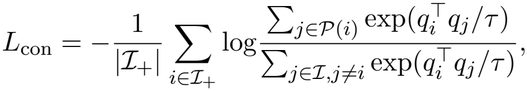

where *ℐ*_+_ = {*i ∈ ℐ* : |*P*(*i*)| *>* 0}. This loss pulls spots sharing *y* together and pushes apart spots from different domains.

#### Distribution control

This group of regularizers shapes the spatial, domain-level, feature-level, and cross-modal structure of the learned embeddings. At the spatial level, a smoothness term drives each spot embedding toward agreement with its spatial neighbors:

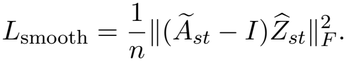

At the domain level, a center loss pulls each spot toward a learnable centroid of its assigned domain, tightening within-domain clusters:

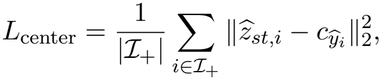

where 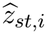 is the *i*-th row of 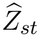 and 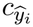 is the learnable centroid of domain *ŷ*_*i*_. At the feature level, a decorrelation loss penalizes the off-diagonal entries of the embedding covariance, thereby suppressing redundancy across embedding dimensions:

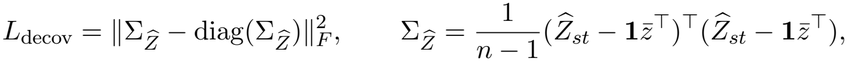

where 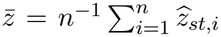. At the cross-modal level, the spot and dissociated-cell embeddings are aligned in the shared RNA latent space by jointly minimizing their maximum mean discrepancy and CORAL distances:

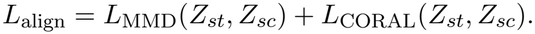

#### Composite objective

The complete objective is a weighted sum of all preceding terms:

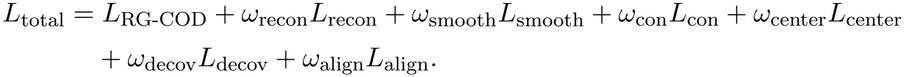

Here *L*_RG-COD_ = *L*_cls_ + *L*_KD_ groups the domain classification and knowledge distillation terms defined above, recovering the RG-COD supervision of Eqn (3).

#### Optimization

All parameters were optimized with Adam at learning rate 10^−3^ and weight decay 10^−4^, while the centroids in *L*_center_ used a smaller learning rate of 5 × 10^−4^ without weight decay. Dissociated cells were processed in mini-batches of 512, each batch using a precomputed nearest-neighbor subgraph for scalability, while spatial spots were forwarded in a single batch. Mixed-precision training was optionally enabled on GPU. Class-balanced cross-entropy weights, set to the inverse domain frequency, were applied to all classification components. Unless otherwise specified, *T* = 2, *τ* = 0.2, and

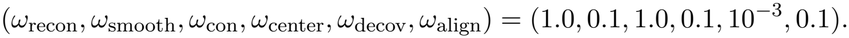

For single-omics use, additional optional regularizers on spot-level domain frequency and joint cell-type-domain distribution can be enabled. Complete loss forms and the full hyperparameter list are provided in Supplementary Notes, S1.

### Implementation details

#### Data processing

Both single-cell and spatial transcriptomics data are required as input to COMPASS, and additional spatial omics measurements, such as ADT or ATAC, can be incorporated when available. When no auxiliary modality is available, we reuse the primary modality as the auxiliary input so that the same architecture applies unchanged. The spatial transcriptomics matrix *X*_st_ was preprocessed by optional removal of mitochondrial genes, library-size normalization to 10^4^ counts per spot, and log-normalization. Highly variable genes were then identified from the normalized matrix using the variance-stabilizing procedure of Seurat v3, and the top 1000 genes were retained as the shared feature space for spatial spots and dissociated cells. Single-cell profiles were aligned to this gene template by matching shared genes and imputing missing entries with their per-gene mean expression across spots in *X*_st_. Each gene was subsequently centered and scaled using the mean and standard deviation computed from the spatial template, so that spot-level and single-cell features lie on a comparable numerical scale before downstream learning. For auxiliary modalities, protein measurements were normalized by centered log-ratio transformation, whereas chromatin accessibility peak matrices were normalized by term frequency–inverse document frequency followed by log-normalization.

#### Dimensionality reduction

Principal component analysis was applied to the standardized spatial transcriptomics matrix to obtain the low-dimensional representation 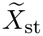 with 50 components, and the same projection was applied to single-cell features to yield a matched representation in the shared gene space. For high-dimensional or sparse inputs, truncated singular value decomposition was used as a numerically stable alternative. For the auxiliary modality, the resulting feature matrix 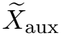 was obtained as follows: protein measurements were further reduced by PCA to 30 components after centered log-ratio transformation and feature-level standardization, whereas chromatin accessibility matrices were projected to 30 components by latent semantic indexing on TF–IDF features and subsequently standardized. When required for coarse spatial domain inference, the RNA and auxiliary representations were concatenated with modality-specific weights and further compressed by a joint PCA to 20 components, which stabilized covariance estimation in the downstream Gaussian mixture model.

### Graph construction

The spatial adjacency underlying the normalized matrix *Ã*_st_ was constructed from spot coordinates 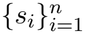 by a nearest-neighbor strategy with six neighbors per spot under Euclidean distance. The graph was then symmetrized and augmented with self-loops before applying symmetric degree normalization, as specified in Model Specification. Edge weights were binary by default, and Gaussian distance weighting could be optionally enabled to attenuate the contribution of distant neighbors. For dissociated cells, a nearest-neighbor graph with 15 neighbors per cell was built in the shared RNA latent space to capture transcriptional neighborhood structure. This graph was precomputed once on the full single-cell reference and reused across mini-batches through lightweight subgraph extraction to support scalable training. All adjacency matrices were stored as sparse tensors for efficient propagation in the graph encoder.

### Downstream analysis

All downstream analyses were performed on the reconstructed single-cell spatial atlas 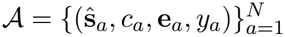, as defined in Model Specification. Here, ŝ_*a*_ denotes the reconstructed coordinate of cell *a, c*_*a*_ its cell-type label, **e**_*a*_ its single-cell expression profile inherited from the scRNA-seq reference, and *y*_*a*_ its spatial domain label. For any spatial unit *R*, including a capture spot, anatomical region, inferred domain, tumour region, or regular grid bin, we denote by *A*_*R*_ = {*a* : ŝ_*a*_ *∈ R*} the mapped cells contained in that unit.

#### Spot-level and region-level composition reconstruction

For each spatial unit *R*, the proportion of cell type *c* was computed as

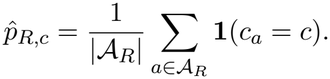

When spot-level assignment rather than continuous coordinates was required, cells were grouped by their final assigned capture spot. The resulting compositions were used to generate proportion maps across spots, regions, and inferred domains, compare with ground-truth, reference-derived, or annotation-based proportions, and assess concordance with anatomical annotations. Pairwise Pearson correlations between cell-type proportion profiles across spatial units were further used to summarize spatial co-variation among cell populations.

#### Spatial expression and marker-program recovery

Expression profiles inherited from the scRNA-seq reference were projected back through the mapped cells and aggregated within the same spatial units. For gene *g*, the reconstructed expression in unit *R* was defined as

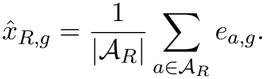

Marker-program scores were obtained analogously by averaging across predefined marker sets or differentially expressed genes associated with a given cell type, tissue domain, or pathological compartment. These maps were used to assess the recovery of lineage markers, domain-specific transcriptional programs, and pathological expression gradients.

#### Cell-cell proximity and co-localization analysis

A spatial graph *G* = (*A, ℰ*) was constructed from reconstructed coordinates using either a *k*-nearest-neighbor or fixed-radius rule, with the neighborhood size or radius chosen according to the spatial scale of each dataset. For each pair of cell types (*c, c*^′^), the observed adjacency count 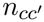 was obtained from graph edges connecting cells of the two types, and the expected count 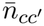 was estimated by permuting cell-type labels while keeping the graph fixed. Contact enrichment was defined as

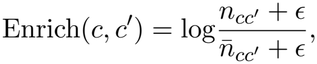

where *ϵ* is a small pseudocount. The resulting enrichment matrix and co-localization network summarized cell-type proximity patterns, while cell-centered neighborhood compositions supported downstream niche analysis.

#### Spatial niche classification

For each spatial unit *R*, the composition vector 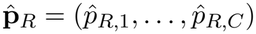 over *C* cell types was obtained as above. Niche labels were assigned either by predefined composition thresholds, yielding interpretable categories such as tumour-dominant, immune-infiltrated, stromalenriched, or immuno-stromal niches, or by unsupervised clustering of 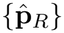 pooled within and across samples. Niche maps, area fractions, and mapped-cell fractions were then compared across samples and clinical subtypes.

#### Cross-sample spatial distribution and boundary-zone analysis

Reconstructed coordinates were first normalized to a common tissue coordinate system. For each cell type *c*, empirical spatial distributions were constructed from cells of that type in each sample, and pairwise Wasserstein distances between samples were summarized into within-subtype and between-subtype means. The spatial divergence gap was defined as

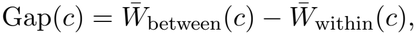

where positive values indicate subtype-specific organization and negative values indicate stronger patient-level variability. To further dissect tumour-margin organization, boundary zones were defined as cells or spatial units whose local neighborhood on *G* contained both cancer and non-cancer populations. Within these zones, physical co-occurrence between non-cancer cell types was quantified by the Jaccard coefficient of neighborhood memberships, and interaction preference by the log odds ratio of observed against expected adjacencies under cell-type label permutation, yielding sample- and subtype-level summaries of boundary organization.

### Evaluation

#### Simulation framework

To systematically evaluate cell-resolved spatial atlas reconstruction under controlled conditions, we generated simulated spatial transcriptomic and chromatin-accessibility datasets by aggregating cells from a matched single-cell multi-omics reference (Supplementary Notes, S2). Spatial spots were placed on a structured lattice, assigned to predefined tissue regions, and populated by sampling cells from region-specific cell-type compositions before aggregating their RNA and ATAC profiles. This design mimics spot-level spatial multi-omics measurements while retaining ground-truth cell-type proportions and cell-level coordinates, enabling joint assessment of composition recovery, cell-to-spot assignment, and continuous coordinate reconstruction.

#### Common spatial multi-omics generative framework

The robustness of COMPASS was evaluated using simulated spatial transcriptomic and chromatin-accessibility data generated from a human melanoma single-cell multi-omics reference with jointly measured RNA and ATAC profiles. The reference comprised immune and malignant cell populations spanning T-cell, myeloid, dendritic, and tumour compartments. To preserve realistic tissue organization, spatial spots were arranged on a structured lattice and assigned to predefined tissue sub-regions with region-specific cell-type compositions. Spot-level RNA and ATAC measurements were then generated by aggregating single-cell profiles sampled from the corresponding regional compositions. This design retained ground-truth cell-type proportions and cell-level coordinates, allowing direct assessment of both spatial mapping accuracy and proportion reconstruction.

#### Matched multimodal baseline simulation

The matched multimodal simulation served as a baseline setting in which RNA and ATAC measurements were available throughout the tissue, and the single-cell reference was compositionally matched to the simulated spatial data. Cell types were assigned to spatial regions with moderate regional specificity, while localized extreme-composition areas were introduced to mimic heterogeneous tumour microenvironments. In the primary baseline setting, RNA profiles were perturbed by spatially varying technical noise and partial dilution of highly variable genes associated with locally dominant cell types, whereas the ATAC modality was preserved without analogous additive noise. This scenario was designed to evaluate performance when both modalities provide informative signals and only moderate spatial and transcriptomic perturbations are present.

#### Complementary multi-omics simulation

A second, more challenging simulation was designed to create a complementary multi-omics setting in which chromatin accessibility could compensate for reduced transcriptomic separability. Compared with the matched baseline simulation, this complementary simulation deliberately increased spatial co-localization and within-spot mixing of biologically similar cell types, while stronger RNA-specific perturbations were applied to reduce transcriptomic separability and preserve ATAC-derived discriminative information. In this setting, biologically related and transcriptionally confusable cell-type pairs, including CD4^+^ T cells versus regulatory T cells, tumour subtype 1 versus tumour subtype 2, and mDC versus pDC, were assigned to overlapping spatial regions. RNA profiles were further degraded through increased technical noise, more extensive dilution of discriminative genes, ambient RNA contamination, and pair-dependent crosstalk between confusable cell types. This design allowed us to isolate the contribution of chromatin accessibility when RNA alone provides ambiguous evidence for cell-type identity.

#### Controlled perturbation experiments

Robustness to incomplete transcriptomic coverage was assessed by extending the complementary simulation to include spatially restricted RNA availability. Rather than removing spots or altering the underlying tissue structure, RNA measurements were retained in only one half of the tissue and masked in the other half, while ATAC measurements, spatial coordinates, metadata, and ground-truth proportions were retained. This setting tested whether COMPASS could recover spatial organization and cell-type composition when transcriptomic information was unavailable across a contiguous tissue region. In separate sensitivity analyses, ATAC-specific noise was introduced across all simulation scenarios to evaluate robustness to perturbations in chromatin-accessibility measurements.

Sensitivity to reference incompleteness was evaluated by reducing the size of the single-cell RNA reference used for mapping. The reference was subsampled in a stratified manner within each cell type, preserving broad compositional diversity while reducing the number of available reference profiles. This simulation tested whether COMPASS remains accurate when the reference atlas is limited, a common constraint in applications involving rare cell states, small patient cohorts, or costly multi-omics profiling. Across all settings, the simulated ground truth enabled quantitative comparison of inferred spatial assignments and cell-type proportions. We finally performed targeted ablations of domain-prior construction, RG-COD training, multi-objective loss design, OT-based assignment, and PDGR-based coordinate refinement, using the same composition, localization, distributional, and runtime metrics as the robustness benchmark under Scenario 4. Detailed values for lattice construction, region assignment, sampling distributions, perturbation strengths, count aggregation, ablation variants, and random seeds are provided in the Supplementary Methods.

#### Benchmarked methods

To systematically evaluate the performance of COMPASS for single-cell spatial localization and cell-type proportion reconstruction, we conducted a comprehensive benchmark against two categories of state-of-the-art methods (Supplementary Notes, S3). All comparative methods were run following their official recommended workflows with default parameters, unless otherwise specified. To dissect the contribution of multi-omics integration, we additionally evaluated two internal variants of our model: COMPASS-single, the single-omics version that uses only the spatial RNA modality without any auxiliary omics, and COMPASS-noise, which retains the auxiliary modality but corrupts it with additional measurement noise. The full COMPASS model uses both the spatial RNA modality and the paired auxiliary omics, distilling the auxiliary-derived domain structure into a deployable RNA-only encoder.

For single-cell spatial mapping tasks, we compared COMPASS (version 1.0.0) with five widely used mapping methods: CMAP (version 1.0.0), CellTrek (version 0.0.94), CytoSPACE (version 1.1.0), SIMO (version 1.0.0), and Tangram (version 1.0.4). CMAP, a divide-and-conquer spatial assignment method, was run with default parameters, except cluster k (the number of clusters) was matched to COMPASS, and the cell-filtering threshold was relaxed to 0.5 to retain more cells. CellTrek, which maps single cells via co-embedding and multivariate random forest modeling, used default parameters except repel r = NULL for automatic repelling radius calculation (one-fourth of the spot distance). CytoSPACE, an optimization-based method that achieves globally optimal cell-to-spot assignment via the shortest augmenting path algorithm, was run with default parameters. SIMO, a multimodal spatial integration method that allocates cells to spatial spots via a sequential mapping strategy, was run with default parameters, except the Leiden clustering resolution was adjusted to align the cluster count with COMPASS for each dataset. Tangram, a deep learning-based framework that aligns single-cell data to spatial data, was executed using <monospace>mode=‘constrained’</monospace> to map cells while respecting the provided cell density priors.

We further compared the cell-type proportion reconstruction performance of COMPASS with six mainstream spatial transcriptomics deconvolution methods: 1) CARD (version 1.1) utilizing the <monospace>CARD_deconvolution()</monospace> function with default settings; 2) RCTD (version 2.2.1) utilizing the <monospace>run.RCTD()</monospace> function with mode = ‘full’ on default settings; 3) <monospace>GraphST</monospace> (version 1.0.0) utilizing the GraphST() function with default settings; 4) Seurat (version 4.3.0) utilizing the <monospace>FindTransferAnchors()</monospace> and <monospace>TransferData()</monospace> functions with default settings; 5) cell2location (version 0.1.4) utilizing the <monospace>cell2location()</monospace> function with default settings; 6) SPOTlight (version 1.2.0) utilizing the <monospace>runDeconvolution()</monospace> function with default settings. For consistency, we followed the officially recommended guidelines for each method in our comparative analysis.

### Benchmarked metrics

To provide a unified assessment of spatial reconstruction performance, we evaluated COMPASS and competing methods at four complementary levels: spatial-domain recovery, spot-level composition reconstruction, single-cell mapping coherence, and cell-level localization accuracy (Supplementary Notes, S4).

To assess spatial-domain recovery, we evaluated the agreement between inferred spatial domains and ground-truth annotations using standard external clustering metrics, including adjusted Rand index (ARI), mutual information (MI), normalized mutual information (NMI), adjusted mutual information (AMI), homogeneity, and completeness. For settings without manual annotations, we used internal validation metrics, including the Silhouette coefficient, Calinski–Harabasz (CH) index, and Davies–Bouldin (DB) index, computed from per-spot cell-type proportions with inferred domains as cluster labels.

The accuracy of reconstructed cell-type proportions was assessed per spot using Root Mean Square Error (RMSE), Structural Similarity Index Measure (SSIM), Pearson correlation coefficient (PCC), Universal Quality Index (UQI), and Erreur Relative Globale Adimensionnelle de Synthèse (ERGAS), then summarized across spots. Together, the metrics capture absolute reconstruction error (RMSE, ERGAS), linear agreement (PCC), and structural or profile-level similarity (SSIM, UQI) between predicted and reference cell-type proportion profiles.

To assess single-cell spatial mapping coherence, we examined how mapped cell counts were distributed across spots and domains using Jensen–Shannon divergence (JSD) and coefficient of variation (CV), and summarized the two metrics with a Mean Objective Score. JSD quantifies between-domain discrimination of cell-count distributions, whereas CV captures within-domain variability across spots. The Mean Objective Score was defined as the difference between the mean JSD and the mean CV, with higher values indicating stronger inter-domain separation and more stable intra-domain cell allocation. We further characterized the spatial smoothness of learned embeddings using Moran’s *I* and Geary’s *C* on a binary spatial adjacency graph.

To evaluate single-cell mapping accuracy, we measured the Euclidean distance between each cell’s predicted and ground-truth location in the same full-resolution tissue coordinate system. All metrics were computed on a fixed evaluation set *E* of original cells, excluding augmentation-derived ones. For each output type *k ∈* {spot, coord}, let *M*_*k*_ *⊆ E* denote the cells with a valid predicted location. 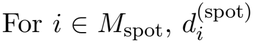 is the Euclidean distance between the ground-truth cell coordinate and the center of the assigned spatial spot, thereby evaluating the discrete cell assignment. For 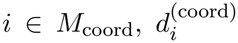 is the Euclidean distance between the ground-truth cell coordinate and the continuous sub-spot coordinate produced by Poisson-disk geometric refinement, thereby evaluating coordinate-level localization accuracy. We report the conditional mean distance over successfully mapped cells 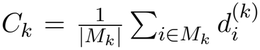, together with its median. Because *C*_*k*_ alone can be lowered by selectively mapping only easy cells, we additionally define a coverage-aware penalized cell-wise distance,

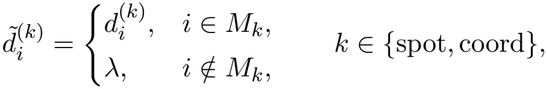

and report its mean over all evaluation cells,

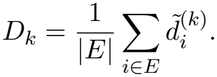

Here, *λ* is set to the diagonal length of the bounding box of the ground-truth coordinates, providing a fixed tissue-scale penalty for unmapped cells. We refer to *D*_spot_ as the penalized spot distance (PSD) and *D*_coord_ as the penalized coordinate distance (PCD).

### Computational Time of COMPASS

COMPASS achieved competitive computational efficiency while performing joint spatial reconstruction and cell-type composition analysis in both simulation studies and real-data applications. In Scenario 1, which contained 576 spots and 10 cell types, COMPASS completed the full analysis in 108.85 s in single-omics mode and 240.32 s in multi-omics mode (Supplementary Fig. S69(a)). Although several lightweight single-task methods required less time, including GraphST (14.32 s), Tangram (41.69 s), and CARD (161.37 s), COMPASS was faster than several widely used deconvolution methods, such as Seurat (350.06 s), RCTD (734.27 s), cell2location (2700.93 s), and SPOTlight (3408.00 s). COMPASS also showed a favorable runtime compared with coordinate-level spatial mapping methods, including SIMO (215.57 s), CytoSPACE (454.39 s), CellTrek (945.17 s), and CMAP (1755.30 s). Similar computational gains were observed across the remaining three simulated scenarios and the real MOB dataset, indicating that COMPASS maintains consistent efficiency across different data settings (Supplementary Figs S69(b)-(d), and S70). The ablation variants of COMPASS, covering the loss, domain, training, optimal transport, and mapping modules, were also completed within comparable runtimes (Supplementary Fig. S71), confirming that no individual component disproportionately dominates the computational cost. All runtime experiments were conducted on a Linux server with 32 Intel(R) Xeon(R) Gold 6354 CPUs and an NVIDIA GeForce RTX 3090 GPU. These results suggest that COMPASS provides a favorable balance between computational cost and analytical scope, achieving efficient runtime despite its multi-module design.

## Data availability

All datasets used in this study are publicly available, and are described in detail in Supplementary Notes, S5. These include the high-resolution Xenium data (https://www.10xgenomics.com/products/xenium-in-situ/preview-dataset-human-breast; In Situ Sample 1, Replicate 1, FRP and Visium Spatial Files), the mouse olfactory bulb ST data from Spatial Transcriptomics v1.0 (https://www.spatialresearch.org), the mouse anterior brain 10x Visium data (https://support.10xgenomics.com/spatial-gene-expression/datasets/1.0.0/V1_Mouse_Brain_Sagittal_Anterior), the mouse posterior brain 10x Visium data (https://support.10xgenomics.com/spatial-gene-expression/datasets/1.0.0/V1_Mouse_Brain_Sagittal_Posterior), the 10x Visium human lymph node data from the GEO database (accession no. GSE263617), and the clinical breast cancer data from the NCBI GEO database (accession no. GSE176078).

## Code availability

COMPASS is implemented in Python and all source code used in this study can be accessed at https://github.com/guanxingbo/COMPASS/.

## CRediT author statement

H.H, W.L., and Q.D. conceptualized and supervised the project. X.W., X.G., and Q.D. designed and developed the method. X.W., Y.W., X.G., and N.L. performed the method analyses across simulated and real datasets, while X.W. and W.F. implemented and ran benchmark analyses. X.W., W.F., Y.W, and Q.D. contributed to figure design, visualization, and presentation refinement. X.W., W.F., Y.W., X.G., and Q.D. drafted the initial manuscript, while Y.B., L.L., H.P., and W.L. collectively revised and edited the manuscript.

## Acknowledgment

This study was supported by the Shenzhen Science and Technology Program (QNXMC20250701091418025), the National Natural Science Foundation of China (12401361), the Fundamental Research Funds for the Central Universities (1082204112J06), the Natural Science Foundation of Sichuan Province (2025ZNSFSC0809), the Shanghai Hospital Development Center Foundation (SHDC12026113), the Shanghai Oriental Talents Program (BJWS2024013), the Research Grants Council of Hong Kong (HKBU 12302022), and the Initiation Grant for Faculty Niche Research Areas of Hong Kong Baptist University (RC-FNRA-IG/24-25/SCI/01).

## References

[1] Xiaowei A. Method of the year 2020: spatially resolved transcriptomics. Nat Methods. 2021;18(1):10–1038.

[2] Zhang Q, Jiang S, Schroeder A, Hu J, Li K, Zhang B, et al. Leveraging spatial transcriptomics data to recover cell locations in single-cell RNA-seq with CeLEry. Nature communications. 2023;14(1):4050.

[3] Kumar MP, D. J, Lagoudas G, Jiao Y, Sawyer A, Drummond DC, et al. Analysis of single-cell RNA-seq identifies cell-cell communication associated with tumor characteristics. Cell reports. 2018;25(6):1458–68.

[4] Smit MM, Feller KJ, You L, Storteboom J, Begce Y, Beerens C, et al. Spatially annotated single cell sequencing for unraveling intratumor heterogeneity. Frontiers in Bioengineering and Biotechnology. 2022;10:829509.

[5] Eng CHL, Lawson M, Zhu Q, Dries R, Koulena N, Takei Y, et al. Transcriptome-scale super-resolved imaging in tissues by RNA seqFISH+. Nature. 2019;568(7751):235–9.

[6] Xia C, Fan J, Emanuel G, Hao J, Zhuang X. Spatial transcriptome profiling by MERFISH reveals subcellular RNA compartmentalization and cell cycle-dependent gene expression. Proceedings of the National Academy of Sciences. 2019;116(39):19490–9.

[7] Janesick A, Shelansky R, Gottscho AD, Wagner F, Williams SR, Rouault M, et al. High resolution mapping of the tumor microenvironment using integrated single-cell, spatial and in situ analysis. Nature communications. 2023;14(1):8353.

[8] Strahl PL, Salmén F, Vickovic S, Lundmark A, Navarro JF, Magnusson J, et al. Visualization and analysis of gene expression in tissue sections by spatial transcriptomics. Science. 2016;353(6294):78–82.

[9] Rodriques SG, Stickels RR, Goeva A, Martin CA, Murray E, Vanderburg CR, et al. Slide-seq: A scalable technology for measuring genome-wide expression at high spatial resolution. Science. 2019;363(6434):1463–7.

[10] Stickels RR, Murray E, Kumar P, Li J, Marshall JL, Di Bella DJ, et al. Highly sensitive spatial transcriptomics at near-cellular resolution with Slide-seqV2. Nature biotechnology. 2021;39(3):313–9.

[11] Chen A, Liao S, Cheng M, Ma K, Wu L, Lai Y, et al. Spatiotemporal transcriptomic atlas of mouse organogenesis using DNA nanoball-patterned arrays. Cell. 2022;185(10):1777–92.

[12] Russell AJ, Weir JA, Nadaf NM, Shabet M, Kumar V, Kambhampati S, et al. Slide-tags enables single-nucleus barcoding for multimodal spatial genomics. Nature. 2024;625(7993):101–9.

[13] Oliveira MFd, Romero JP, Chung M, Williams SR, Gottscho AD, Gupta A, et al. High-definition spatial transcriptomic profiling of immune cell populations in colorectal cancer. Nature Genetics. 2025;57(6):1512–23.

[14] Bourgain-Chang E, Kuang X, Cang Z, Nie Q. Reconstructing single-cell resolution from spatial transcriptomics with CellRefiner. Nature Communications. 2026.

[15] Kabanikhin SI, et al. Definitions and examples of inverse and ill-posed problems. J Inverse Ill-Posed Probl. 2008;16(4):317–57.

[16] Chen D, Li S, Wang X. Geometric structure guided model and algorithms for complete deconvolution of gene expression data. Foundations of data science (Springfield, Mo). 2022;4(3):441.

[17] Zeng Z, Li Y, Li Y, Luo Y. Statistical and machine learning methods for spatially resolved transcriptomics data analysis. Genome biology. 2022;23(1):83.

[18] Li H, Zhou J, Li Z, Chen S, Liao X, Zhang B, et al. A comprehensive benchmarking with practical guidelines for cellular deconvolution of spatial transcriptomics. Nature communications. 2023;14(1):1548.

[19] Stuart T, Butler A, Hoffman P, Hafemeister C, Papalexi E, Mauck WM, et al. Comprehensive integration of single-cell data. cell. 2019;177(7):1888–902.

[20] Elosua-Bayes M, Nieto P, Mereu E, Gut I, Heyn H. SPOTlight: seeded NMF regression to deconvolute spatial transcriptomics spots with single-cell transcriptomes. Nucleic acids research. 2021;49(9):e50–0.

[21] Cable DM, Murray E, Zou LS, Goeva A, Macosko EZ, Chen F, et al. Robust decomposition of cell type mixtures in spatial transcriptomics. Nature biotechnology. 2022;40(4):517–26.

[22] Ma Y, Zhou X. Spatially informed cell-type deconvolution for spatial transcriptomics. Nature biotechnology. 2022;40(9):1349–59.

[23] Kleshchevnikov V, Shmatko A, Dann E, Aivazidis A, King HW, Li T, et al. Cell2location maps fine-grained cell types in spatial transcriptomics. Nature biotechnology. 2022;40(5):661–71.

[24] Long Y, Ang KS, Li M, Chong KLK, Sethi R, Zhong C, et al. Spatially informed clustering, integration, and deconvolution of spatial transcriptomics with GraphST. Nature communications. 2023;14(1):1155.

[25] Liu W, Zhang X, Chai X, Fan Z, Lin H, Chen J, et al. FAST: Scalable Factor Analysis for Spatial Dimension Reduction of Multi-section Spatial Transcriptomics. Genomics, Proteomics & Bioinformatics. 2026:qzag006.

[26] Dong Q, Huang Z, Wang X, Feng Z, Liu W, Huang B. Composition-Aware Cross-Sectional Integration for Spatial Transcriptomics. Advanced Intelligent Discovery. 2026:e202500226.

[27] Biancalani T, Scalia G, Buffoni L, Avasthi R, Lu Z, Sanger A, et al. Deep learning and alignment of spatially resolved single-cell transcriptomes with Tangram. Nature methods. 2021;18(11):1352–62.

[28] Wei R, He S, Bai S, Sei E, Hu M, Thompson A, et al. Spatial charting of single-cell transcriptomes in tissues. Nature biotechnology. 2022;40(8):1190–9.

[29] Vahid MR, Brown EL, Steen CB, Zhang W, Jeon HS, Kang M, et al. High-resolution alignment of single-cell and spatial transcriptomes with CytoSPACE. Nature biotechnology. 2023;41(11):1543–8.

[30] Yang P, Jin K, Yao Y, Jin L, Shao X, Li C, et al. Spatial integration of multiomics single-cell data with SIMO. Nature communications. 2025;16(1):1265.

[31] Ke J, Xu J, Liu J, Yang Y, Guo C, Xie B, et al. High-resolution mapping of single cells in spatial context. Nature Communications. 2025;16(1):6533.

[32] Liu Y, Yang M, Deng Y, Su G, Enninful A, Guo CC, et al. High-spatial-resolution multiomics sequencing via deterministic barcoding in tissue. Cell. 2020;183(6):1665–81.

[33] Zhang D, Deng Y, Kukanja P, Agirre E, Bartosovic M, Dong M, et al. Spatial epigenome– transcriptome co-profiling of mammalian tissues. Nature. 2023;616(7955):113–22.

[34] Gayoso A, Steier Z, Lopez R, Regier J, Nazor KL, Streets A, et al. Joint probabilistic modeling of single-cell multiomic data with totalVI. Nature methods. 2021;18(3):272–82.

[35] Ashuach T, Gabitto MI, Koodli RV, Saldi GA, Jordan MI, Yosef N. MultiVI: deep generative model for the integration of multimodal data. Nature methods. 2023;20(8):1222–31.

[36] Hao Y, Hao S, Andersen-Nissen E, Mauck WM, Zheng S, Butler A, et al. Integrated analysis of multimodal single-cell data. Cell. 2021;184(13):3573–87.

[37] Ghazanfar S, Guibentif C, Marioni JC. Stabilized mosaic single-cell data integration using unshared features. Nature biotechnology. 2024;42(2):284–92.

[38] Argelaguet R, Arnol D, Bredikhin D, Deloro Y, Velten B, Marioni JC, et al. MOFA+: a statistical framework for comprehensive integration of multi-modal single-cell data. Genome biology. 2020;21(1):111.

[39] Minoura K, Abe K, Nam H, Nishikawa H, Shimamura T. A mixture-of-experts deep generative model for integrated analysis of single-cell multiomics data. Cell reports methods. 2021;1(5).

[40] Long Y, Ang KS, Sethi R, Liao S, Heng Y, van Olst L, et al. Deciphering spatial domains from spatial multiomics with SpatialGlue. Nature Methods. 2024;21(9):1658–67.

[41] Gao C, Yang C, Zhang L. A graph neural network-based spatial multiomics data integration method for deciphering spatial domains. PLOS Computational Biology. 2025;21(9):e1013546.

[42] Yan X, Fang Z, Ang KS, Olst Lv, Edwards A, Watson T, et al. Mosaic integration of spatial multiomics with SpaMosaic. Nature Genetics. 2026:1–12.

[43] Quail DF, Joyce JA. Microenvironmental regulation of tumor progression and metastasis. Nature Medicine. 2013;19(11):1423–37.

[44] Tepe B, Hill MC, Pekarek BT, Hunt PJ, Martin TJ, Martin JF, et al. Single-cell RNA-seq of mouse olfactory bulb reveals cellular heterogeneity and activity-dependent molecular census of adult-born neurons. Cell reports. 2018;25(10):2689–703.

[45] Nagayama S, Homma R, Imamura F. Neuronal organization of olfactory bulb circuits. Frontiers in neural circuits. 2014;8:98.

[46] Tasic B, Yao Z, Graybuck LT, Smith KA, Nguyen TN, Bertagnolli D, et al. Shared and distinct transcriptomic cell types across neocortical areas. Nature. 2018;563(7729):72–8.

[47] longyahui. JinmiaoChenLab/SpatialGlue: SpatialGlue. Zenodo; 2023. Available from: 10.5281/zenodo.7879713.

[48] Tkachenko A, Kupcova K, Havranek O. B-cell receptor signaling and beyond: the role of Igα (CD79a)/Igβ (CD79b) in normal and malignant B cells. International Journal of Molecular Sciences. 2023;25(1):10.

[49] Andreatta M, Corria-Osorio J, Müller S, Cubas R, Coukos G, Carmona SJ. Interpretation of T cell states from single-cell transcriptomics data using reference atlases. Nature communications. 2021;12(1):2965.

[50] Shen WK, Zhang CY, Gu YM, Luo T, Chen SY, Yue T, et al. An automatic annotation tool and reference database for T cell subtypes and states at single-cell resolution. Science Bulletin. 2025;70(10):1659–72.

[51] Turiello R, Ng SS, Tan E, van der Voort G, Salim N, Yong MC, et al. NKG7 is a stable marker of cytotoxicity across immune contexts and within the tumor microenvironment. European Journal of Immunology. 2025;55(6):e51885.

[52] Krämer B, Nalin AP, Ma F, Eickhoff S, Lutz P, Leonardelli S, et al. Single-cell RNA sequencing identifies a population of human liver-type ILC1s. Cell reports. 2023;42(1).

[53] Wu SZ, Al-Eryani G, Roden DL, Junankar S, Harvey K, Andersson A, et al. A single-cell and spatially resolved atlas of human breast cancers. Nature genetics. 2021;53(9):1334–47.

[54] Denkert C, Von Minckwitz G, Darb-Esfahani S, Lederer B, Heppner BI, Weber KE, et al. Tumourinfiltrating lymphocytes and prognosis in different subtypes of breast cancer: a pooled analysis of 3771 patients treated with neoadjuvant therapy. The lancet oncology. 2018;19(1):40–50.

[55] Schmid P, Adams S, Rugo HS, Schneeweiss A, Barrios CH, Iwata H, et al. Atezolizumab and nab-paclitaxel in advanced triple-negative breast cancer. New England journal of medicine. 2018;379(22):2108–21.

[56] O’Leary KA, Bates AM, Jin WJ, Burkel BM, Sriramaneni RN, Emma SE, et al. Estrogen receptor blockade and radiation therapy cooperate to enhance the response of immunologically cold ER+ breast cancer to immunotherapy. Breast Cancer Research. 2023;25(1):68.

[57] Stanton SE, Disis ML. Clinical significance of tumor-infiltrating lymphocytes in breast cancer. Journal for immunotherapy of cancer. 2016;4(1):59.

[58] Fu T, Dai LJ, Wu SY, Xiao Y, Ma D, Jiang YZ, et al. Spatial architecture of the immune microenvironment orchestrates tumor immunity and therapeutic response. Journal of hematology & oncology. 2021;14(1):98.

[59] Wortman JC, He TF, Solomon S, Zhang RZ, Rosario A, Wang R, et al. Spatial distribution of B cells and lymphocyte clusters as a predictor of triple-negative breast cancer outcome. NPJ Breast Cancer. 2021;7(1):84.

[60] Zhu Q, Shah S, Dries R, Cai L, Yuan GC. Identification of spatially associated subpopulations by combining scRNAseq and sequential fluorescence in situ hybridization data. Nature biotechnology. 2018;36(12):1183–90.

[61] Dai B, Yi L, Wang P, Li H, Hu P, Song Y, et al. 3d-OT: a deep geometry-aware framework for heterogeneous slices alignment of spatial multiomics. Nature Methods. 2026:1–12.

[62] Pukelsheim F. Proportional representation. Springer; 2017.

[63] Bridson R. Fast Poisson disk sampling in arbitrary dimensions. SIGGRAPH sketches. 2007;10(1):1.

