## Supplementary Information for "COMPASS: In silico reconstruction of single-cell-resolved spatial atlases via cross-omics distillation and coordinate refinement"

**5 Data description 110**

### Supplementary Figures

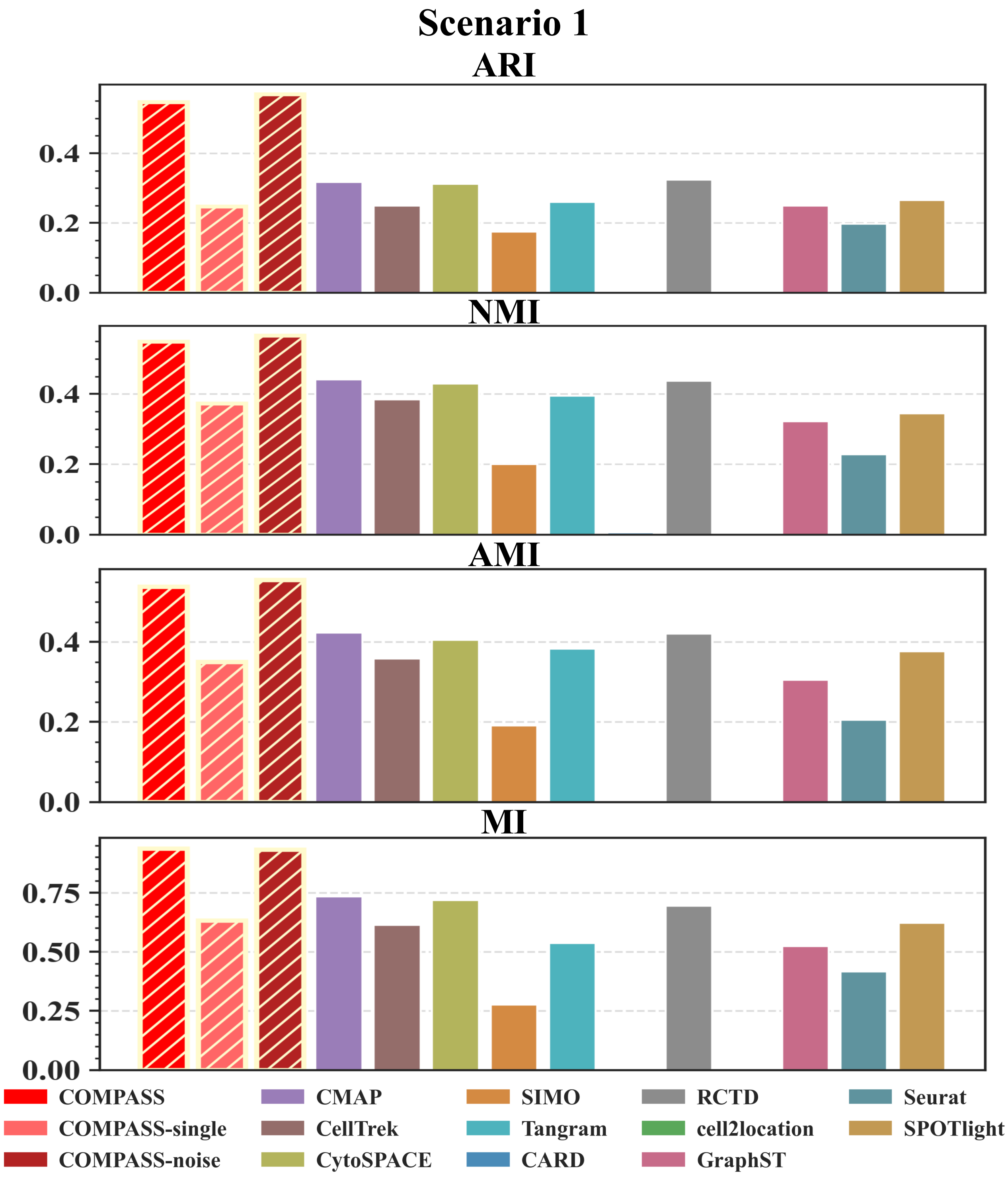

Figure S1: **External clustering agreement in Scenario 1 simulations.** Bar plots compare COMPASS and benchmark methods in the unperturbed simulation setting using four external clustering metrics: adjusted Rand index (ARI), normalized mutual information (NMI), adjusted mutual information (AMI), and mutual information (MI). The inferred spatial domains or dominant reconstructed partitions were compared with the ground-truth spatial annotations generated by the simulation framework. Higher values indicate stronger agreement with the ground-truth tissue structure.

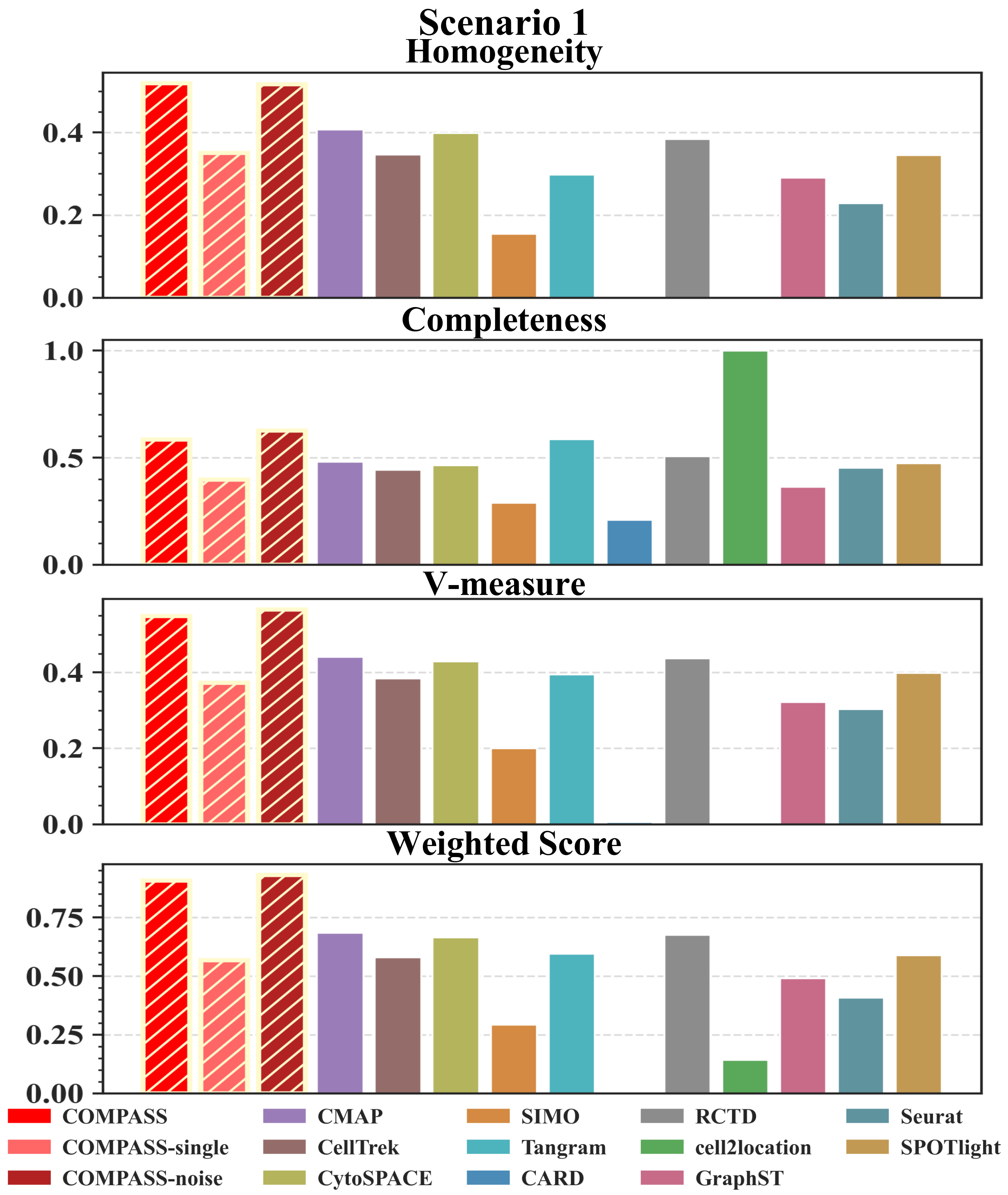

Figure S2: **Cluster purity and completeness in Scenario 1 simulations.** Homogeneity and completeness assess cluster purity and domain recovery, respectively. V-measure summarizes their balance, while the weighted score provides an overall measure of clustering concordance. Higher values indicate better recovery of the simulated spatial domain structure.

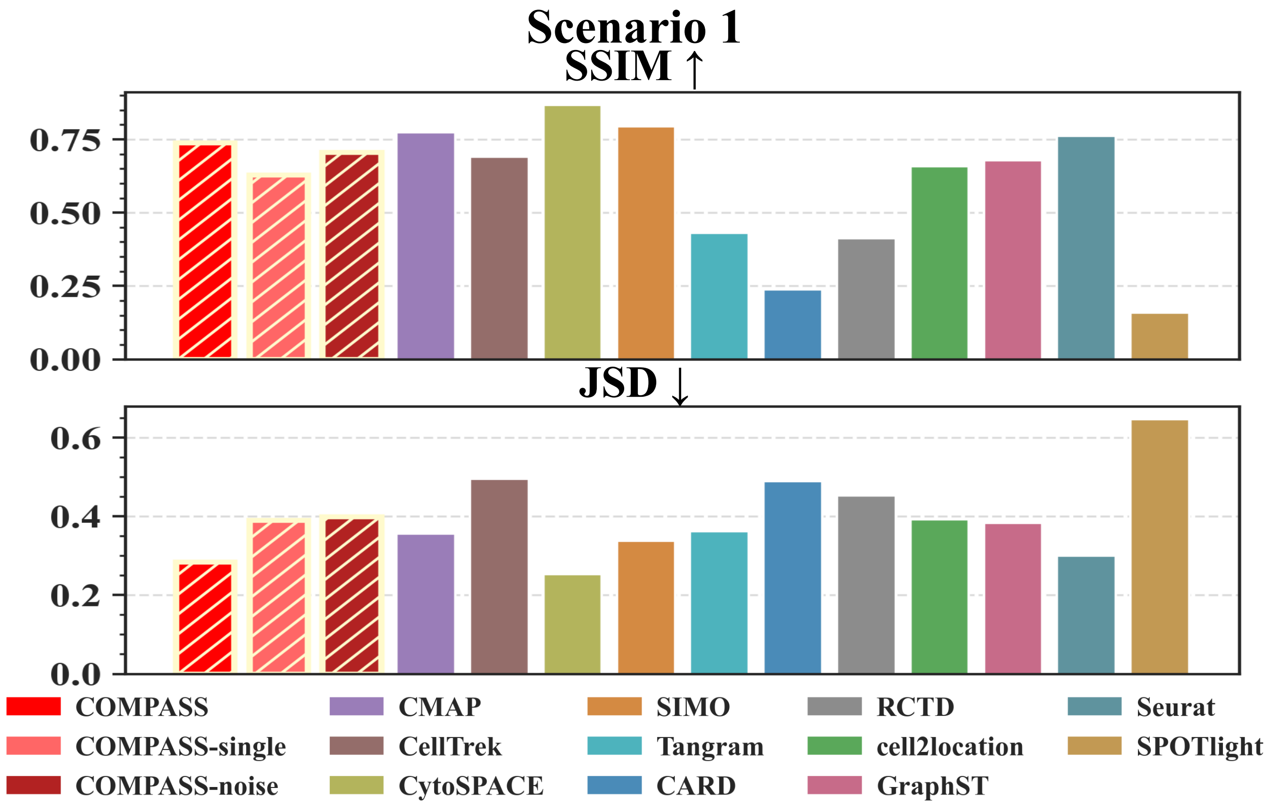

Figure S3: **Spot-level deconvolution performance in Scenario 1 simulations.** Bar plots compare COMPASS and benchmark methods in the unperturbed simulation setting using structural similarity index (SSIM) and Jensen–Shannon divergence (JSD). SSIM evaluates the preservation of spatial structure in reconstructed cell-type proportion maps, whereas JSD measures distributional discrepancy between predicted and ground-truth spot-level compositions. Higher SSIM and lower JSD indicate more accurate recovery of the simulated cell-type composition.

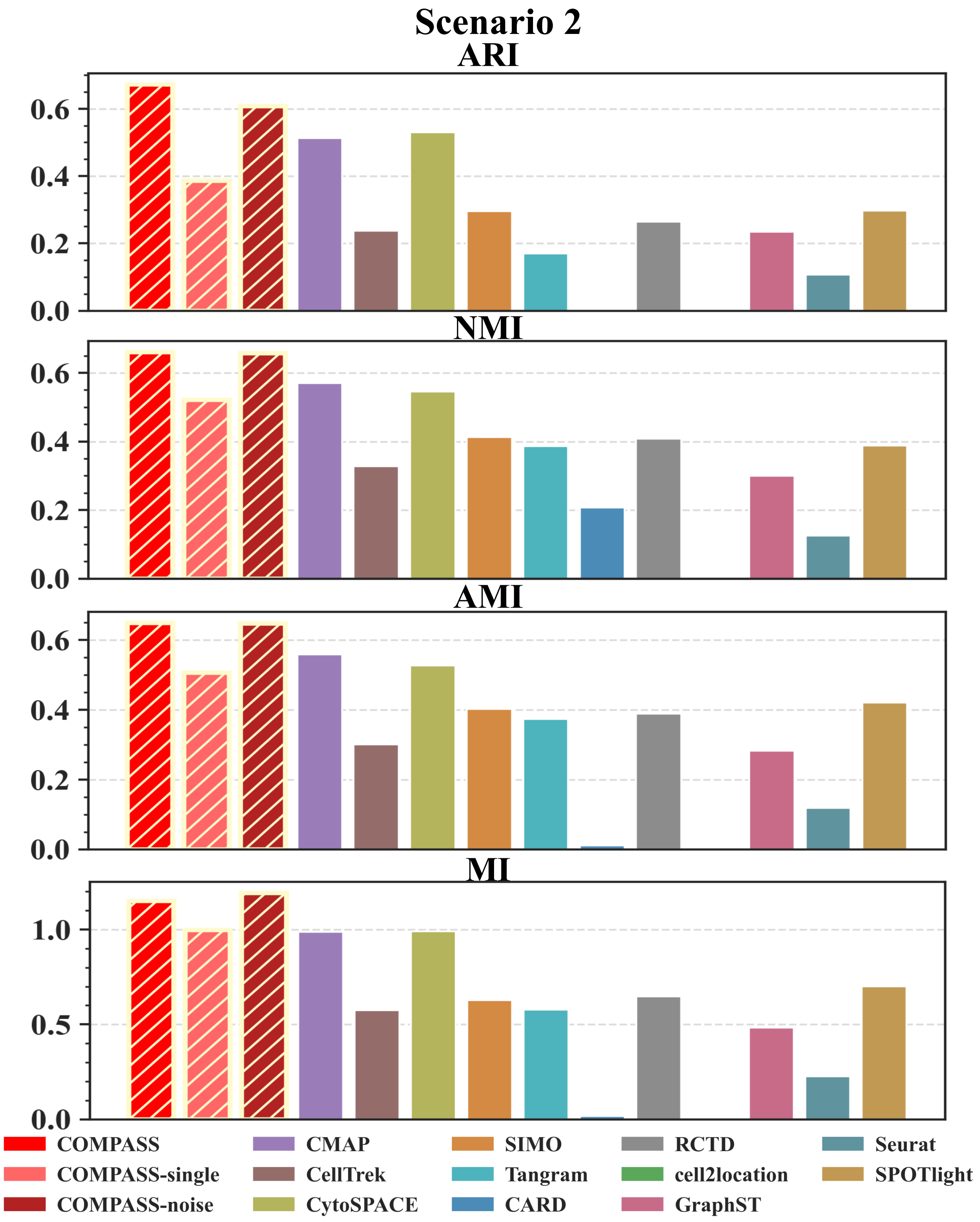

Figure S4: **External clustering agreement in Scenario 2 simulations with spatial RNA noise.** Bar plots compare COMPASS and benchmark methods after introducing measurement noise into the spatial RNA modality while retaining the auxiliary ATAC modality. Clustering concordance was evaluated using ARI, NMI, AMI, and MI between the inferred spatial partitions and the ground-truth spatial annotations. Higher values indicate better preservation of tissue-domain structure under noisy spatial transcriptomic measurements.

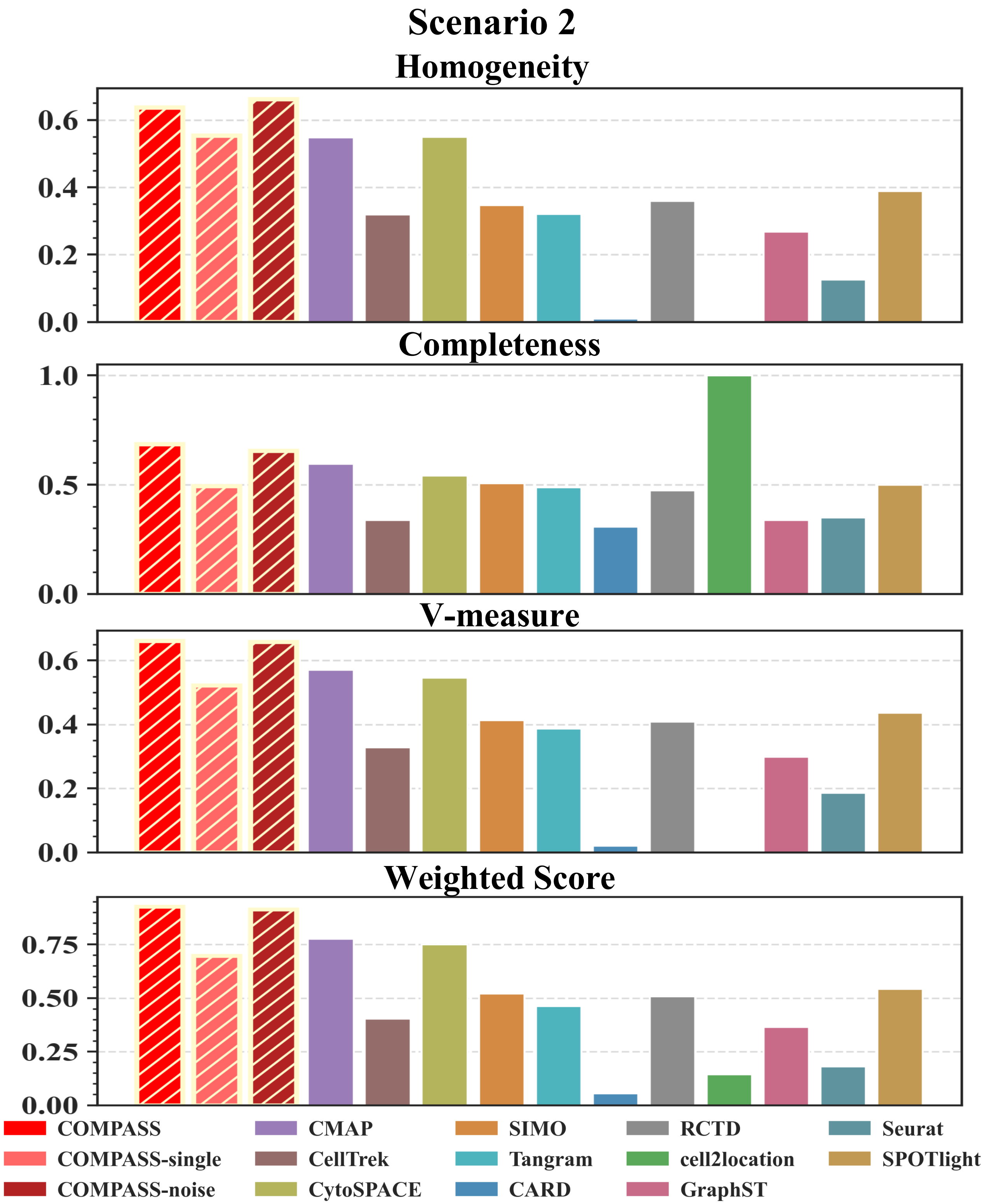

Scenario 2
Homogeneity
Completeness
V-measure
Weighted Score
0.0
0.2
0.4
0.6
0.5
1.0
0.00
0.25
0.50
0.75
COMPASS
COMPASS-single
COMPASS-noise
CMAP
CellTrek
CytoSPACE
SIMO
Tangram
CARD
RCTD
cell2location
GraphST
Seurat
SPOTlight

Figure S5: **Cluster purity and completeness in Scenario 2 simulations with spatial RNA noise.** Bar plots compare COMPASS and benchmark methods under spatial RNA measurement noise using homogeneity, completeness, V-measure, and the weighted summary score. These metrics assess whether the reconstructed partitions remain internally pure and globally consistent with the ground-truth spatial domains despite corrupted RNA measurements. Higher values indicate stronger robustness of spatial domain recovery under transcriptomic noise.

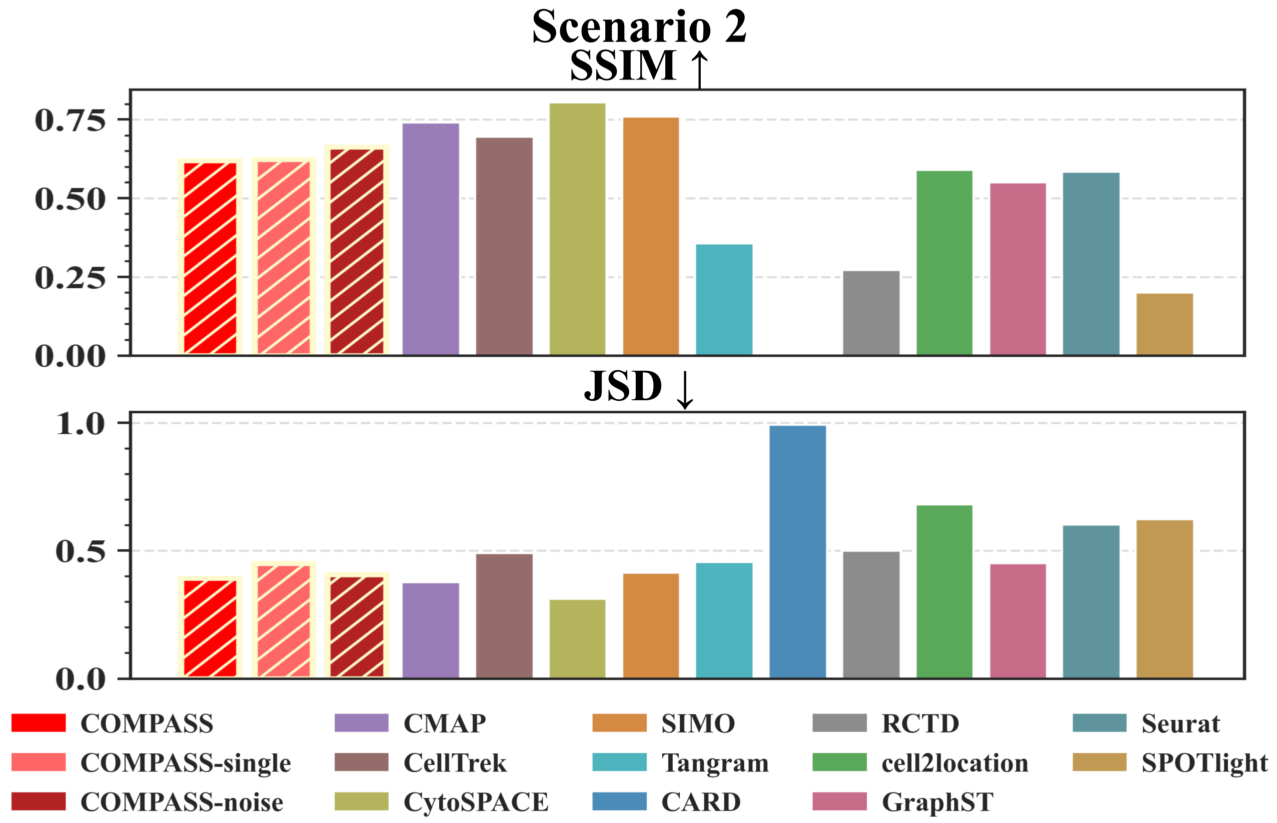

Figure S6: **Spot-level deconvolution performance in Scenario 2 simulations.** Bar plots compare COMPASS and benchmark methods after introducing measurement noise into the spatial RNA modality while retaining the auxiliary assay for transposase-accessible chromatin (ATAC) modality. Performance was evaluated using SSIM and JSD to assess both spatial structural fidelity and distributional agreement of inferred spot-level cell-type proportions. Higher SSIM and lower JSD indicate stronger robustness to noisy spatial transcriptomic measurements.

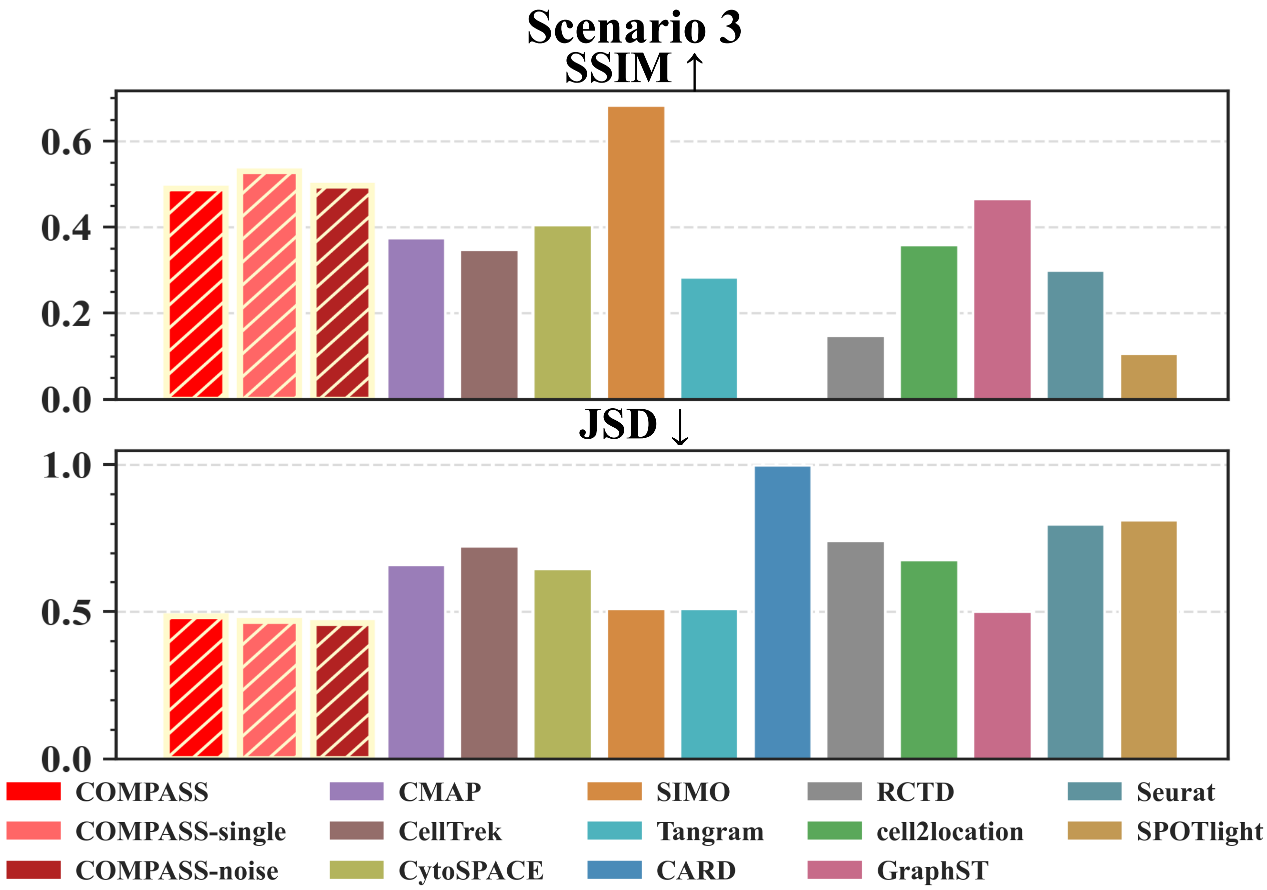

Figure S7: **Spot-level deconvolution performance in Scenario 3 simulations.** Bar plots compare COMPASS and benchmark methods under combined spatial RNA noise and partial loss of spot-level RNA signal. SSIM quantifies the structural consistency of reconstructed cell-type proportion maps, and JSD quantifies the divergence between inferred and ground-truth cell-type composition distributions. Higher SSIM and lower JSD indicate better preservation of spatial composition when RNA measurements become incomplete.

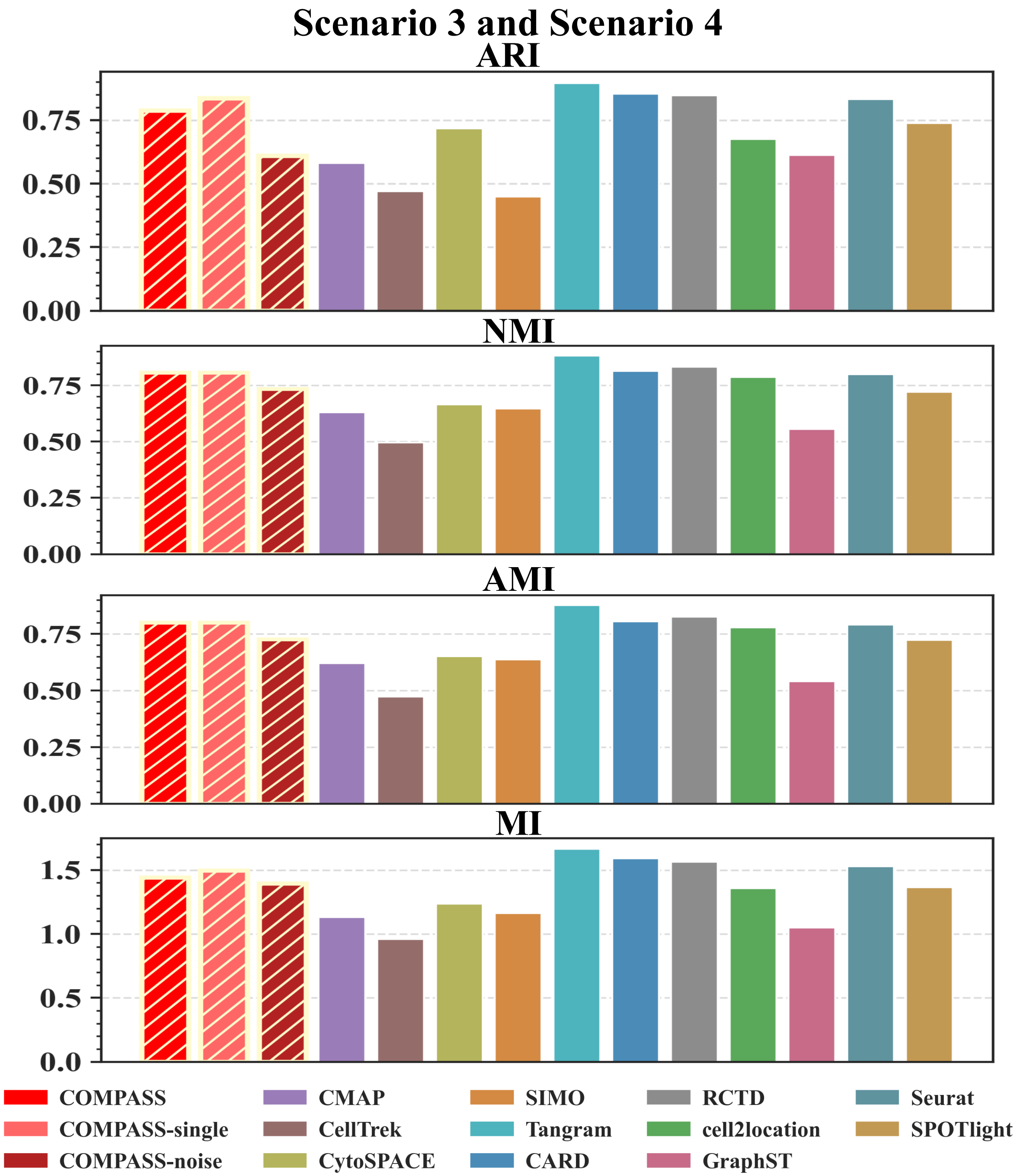
Scenario 3 and Scenario 4
ARI
NMI
AMI
MI
0.00
0.25
0.50
0.75
0.0
0.5
1.0
1.5
COMPASS
COMPASS-single
COMPASS-noise
CMAP
CellTrek
CytoSPACE
SIMO
Tangram
CARD
RCTD
cell2location
GraphST
Seurat
SPOTlight

Figure S8: **External clustering agreement in Scenario 3 and Scenario 4 simulations.** Bar plots compare COMPASS and benchmark methods using ARI, NMI, AMI, and MI in the more challenging perturbation settings. Scenario 3 further masks half of the spot-level RNA signal after spatial RNA corruption, whereas Scenario 4 additionally reduces the dissociated scRNA-seq reference used for mapping. The clustering metrics are identical for Scenario 3 and Scenario 4 because this evaluation is performed on spot-level spatial domain partitions, which are unaffected by the subsequent subsampling of the dissociated single-cell reference. Higher values indicate better agreement with the ground-truth spatial annotations.

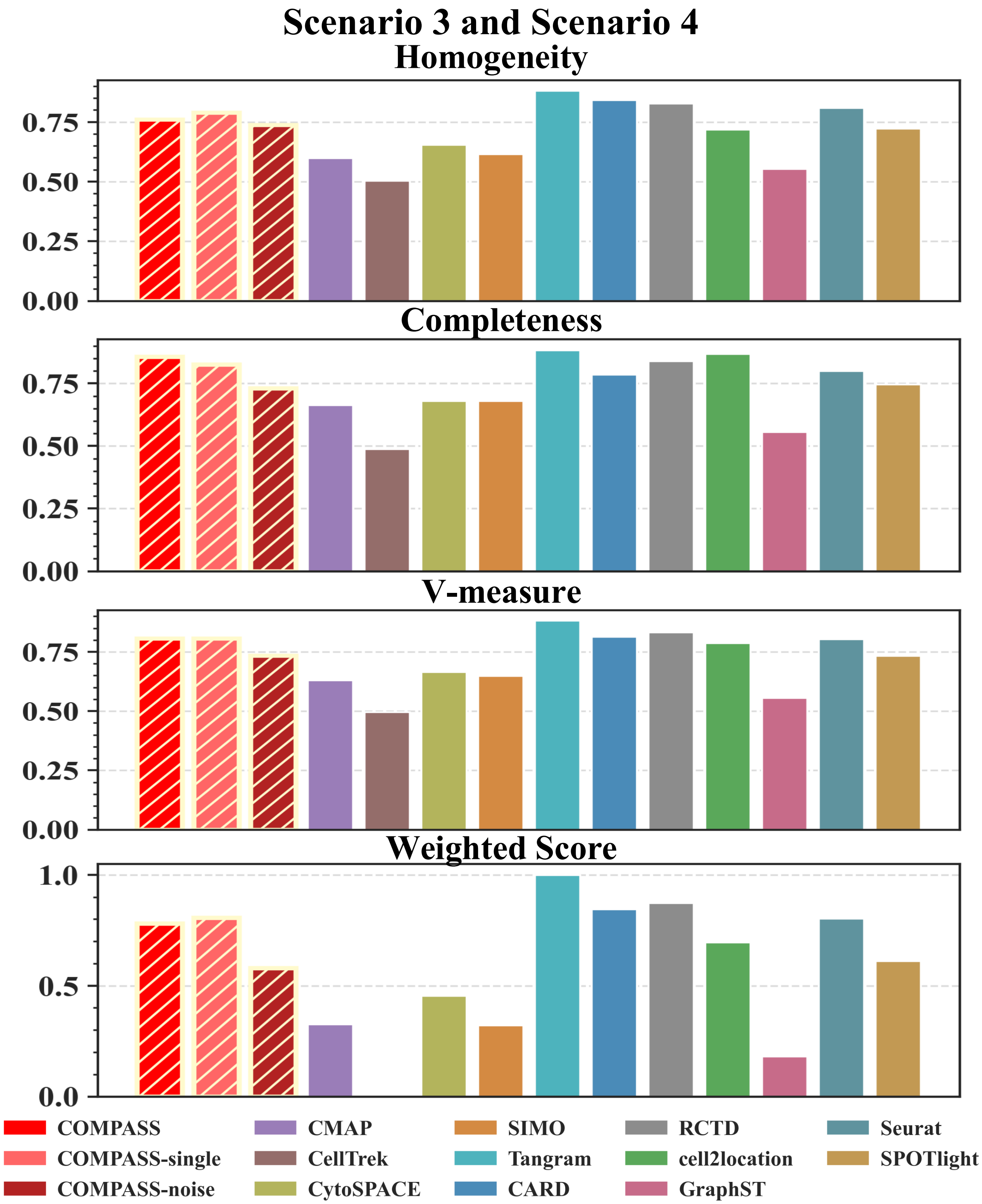
Scenario 3 and Scenario 4
Homogeneity
Completeness
V-measure
Weighted Score
0.00
0.25
0.50
0.75
0.0
0.5
1.0
COMPASS
COMPASS-single
COMPASS-noise
CMAP
CellTrek
CytoSPACE
SIMO
Tangram
CARD
RCTD
cell2location
GraphST
Seurat
SPOTlight

Figure S9: **Cluster purity and completeness in Scenario 3 and Scenario 4 simulations.** Bar plots compare COMPASS and benchmark methods using homogeneity, completeness, V-measure, and the weighted summary score under the most strongly perturbed simulation settings. Scenario 3 evaluates spatial domain recovery after combined RNA noise and partial spot-level RNA signal loss, while Scenario 4 further tests mapping with an incomplete scRNA-seq reference. Because reference subsampling in Scenario 4 does not alter the spot-level domain labels used for clustering evaluation, the Scenario 3 and Scenario 4 clustering scores are shared. Higher values indicate stronger recovery of the simulated tissue-domain organization.

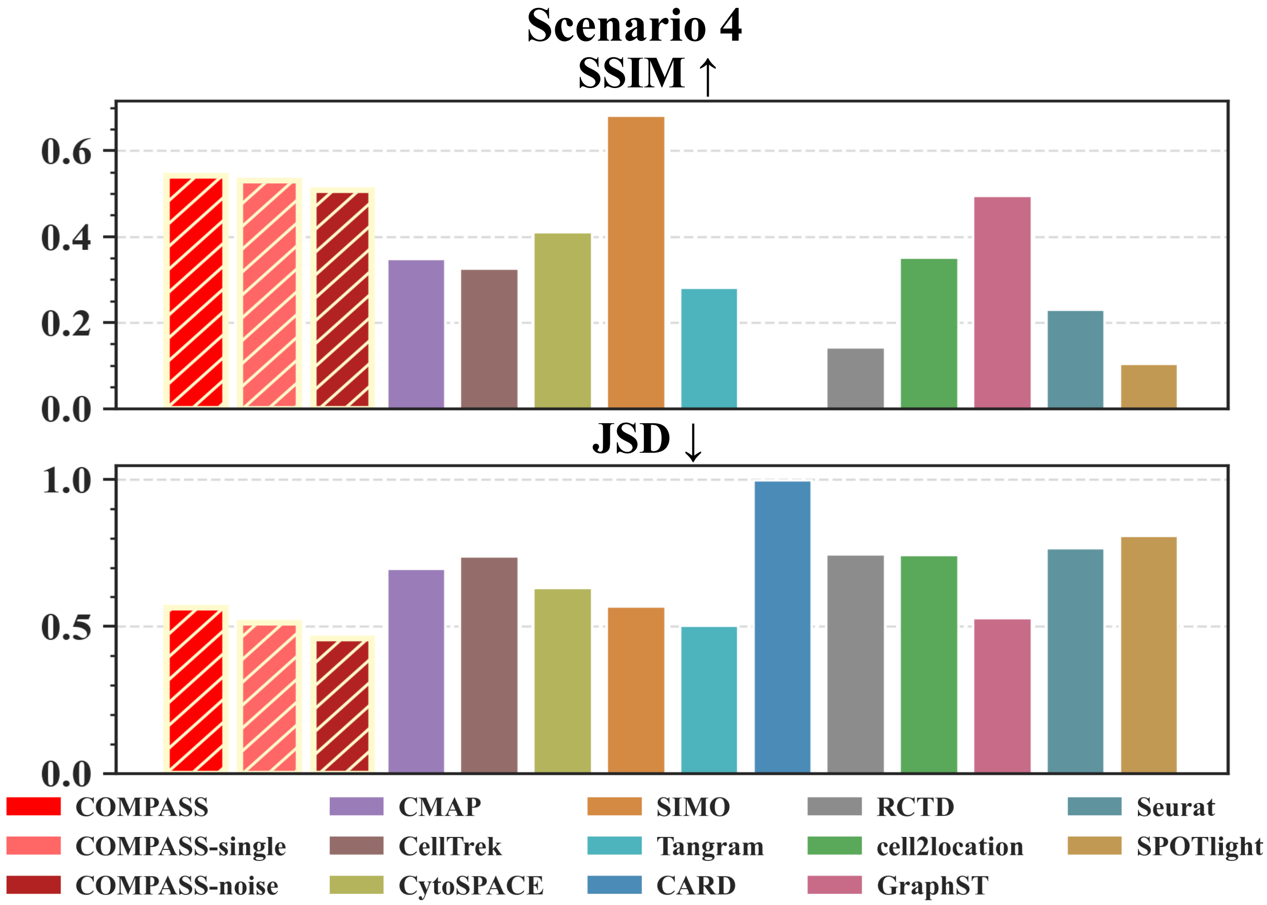

Figure S10: **Spot-level deconvolution performance in Scenario 4 simulations.** Bar plots compare COMPASS and benchmark methods in the most challenging simulation setting, where spatial RNA corruption and partial spot-level RNA signal loss are combined with reduced scRNA-seq reference coverage. SSIM and JSD jointly evaluate whether reconstructed spot-level cell-type proportions preserve both spatial structure and distributional fidelity relative to the simulated ground truth. Higher SSIM and lower JSD indicate more robust composition recovery under compound perturbations.

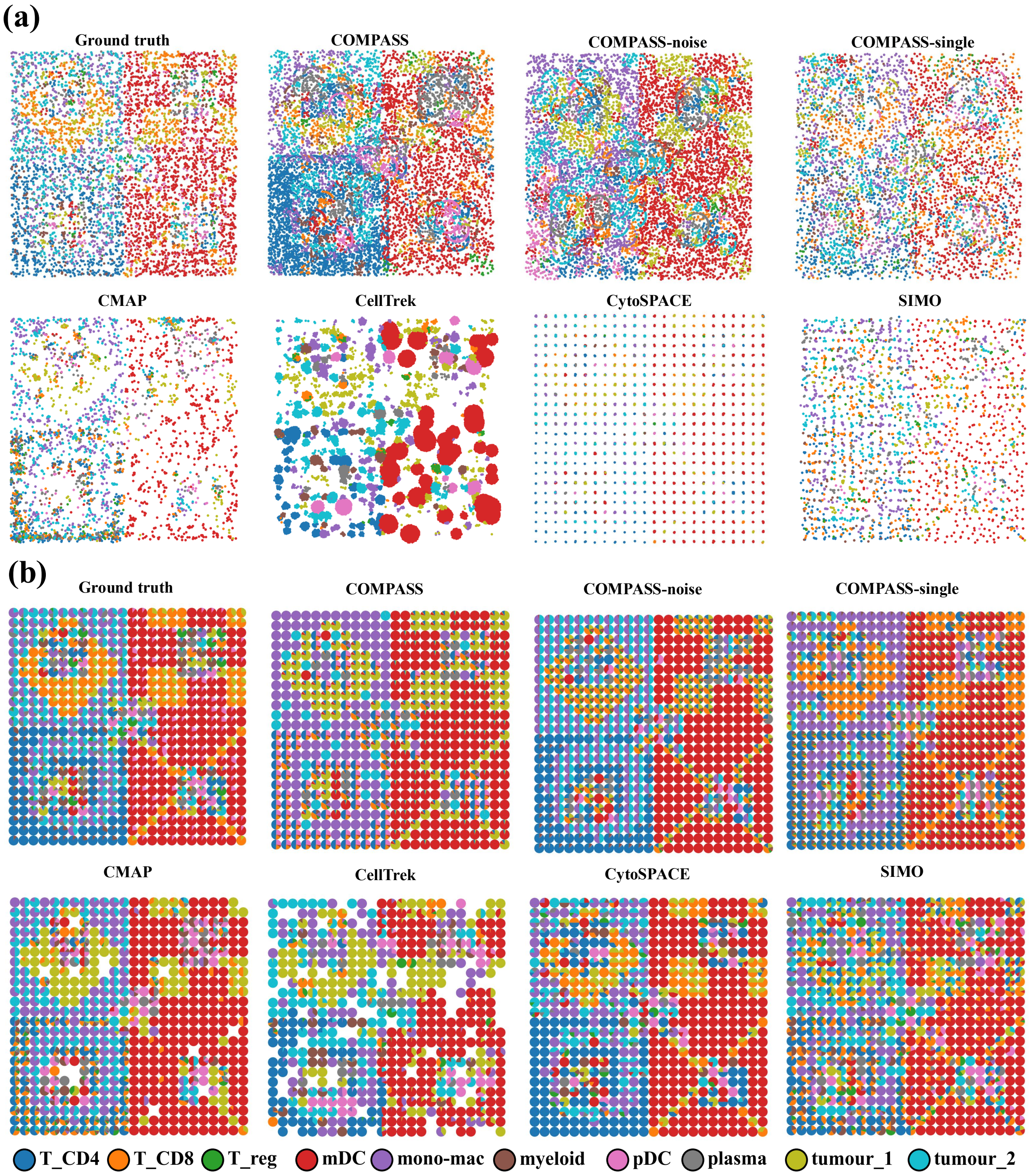

Figure S11: **Mapping-based reconstruction in Scenario 1 simulations.** Spatial maps compare COMPASS variants and representative cell-mapping baselines with the ground truth under the unperturbed setting. (a) Cell-level maps show reconstructed single-cell coordinates for each method. (b) Spot-level maps summarize the mapped cells within each pseudo-spot, providing a deconvolution-like view derived from cell assignments. Colors denote the ten simulated cell types and highlight recovery of tumour boundaries and immune-cell spatial patterns.

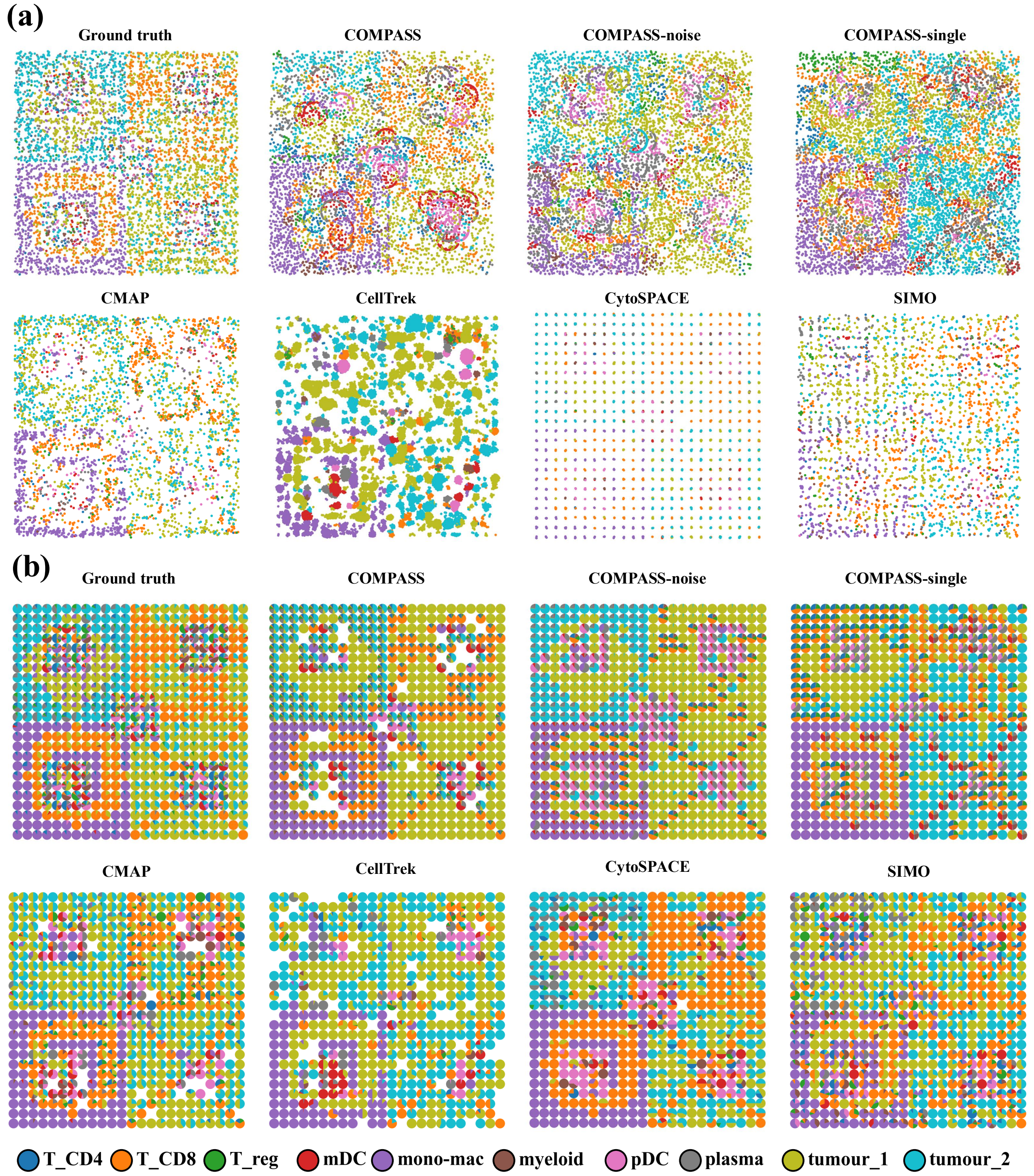

Figure S12: **Mapping-based reconstruction in Scenario 2 simulations.** Spatial maps compare COMPASS variants and representative cell-mapping baselines after adding measurement noise to the spatial RNA modality while retaining auxiliary ATAC information. (a) Cell-level maps show reconstructed single-cell coordinates. (b) Spot-level maps aggregate mapped cells within each pseudo-spot to provide a deconvolution-like summary of local composition. The comparison assesses whether mapping methods preserve tumour structure and cell-type organization when RNA similarity becomes noisy.

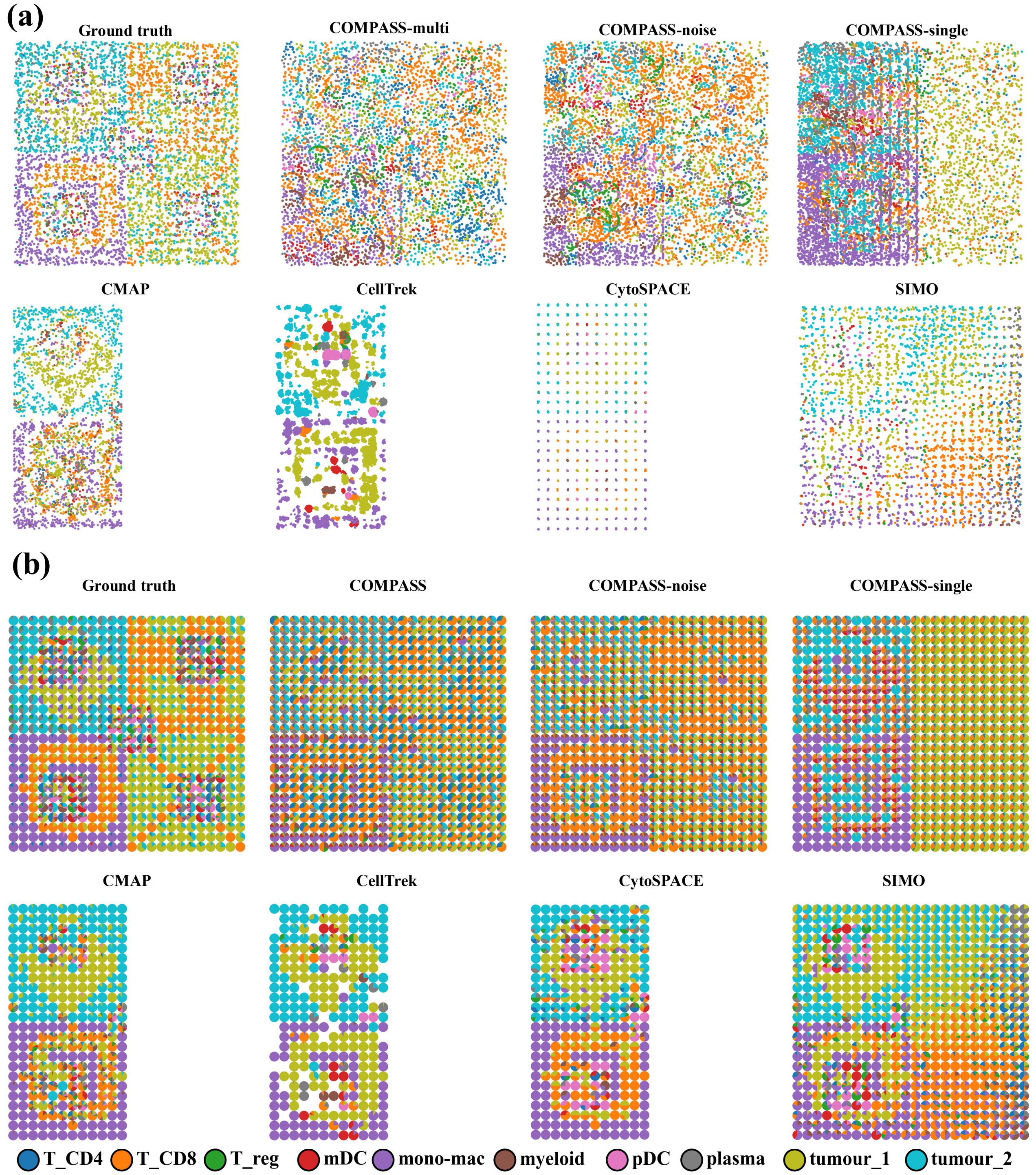

Figure S13: **Mapping-based reconstruction in Scenario 3 simulations.** Spatial maps compare COMPASS variants and representative cell-mapping baselines under combined spatial RNA noise and partial spot-level RNA signal loss. The half-width CMAP, CellTrek, and CytoSPACE panels indicate that the RNA-driven baselines produced valid mappings only for the RNA-retaining half of the lattice; the RNA-masked, ATAC-only half lacked meaningful RNA-based assignments and is therefore not shown. (a) Cell-level maps show reconstructed single-cell coordinates. (b) Spot-level maps summarize mapped cells within pseudo-spots, giving a deconvolution-like view derived from the mapping results. The figure assesses whether mapping methods maintain coherent tumour-domain organization and immune-cell patterns when RNA evidence is strongly weakened.

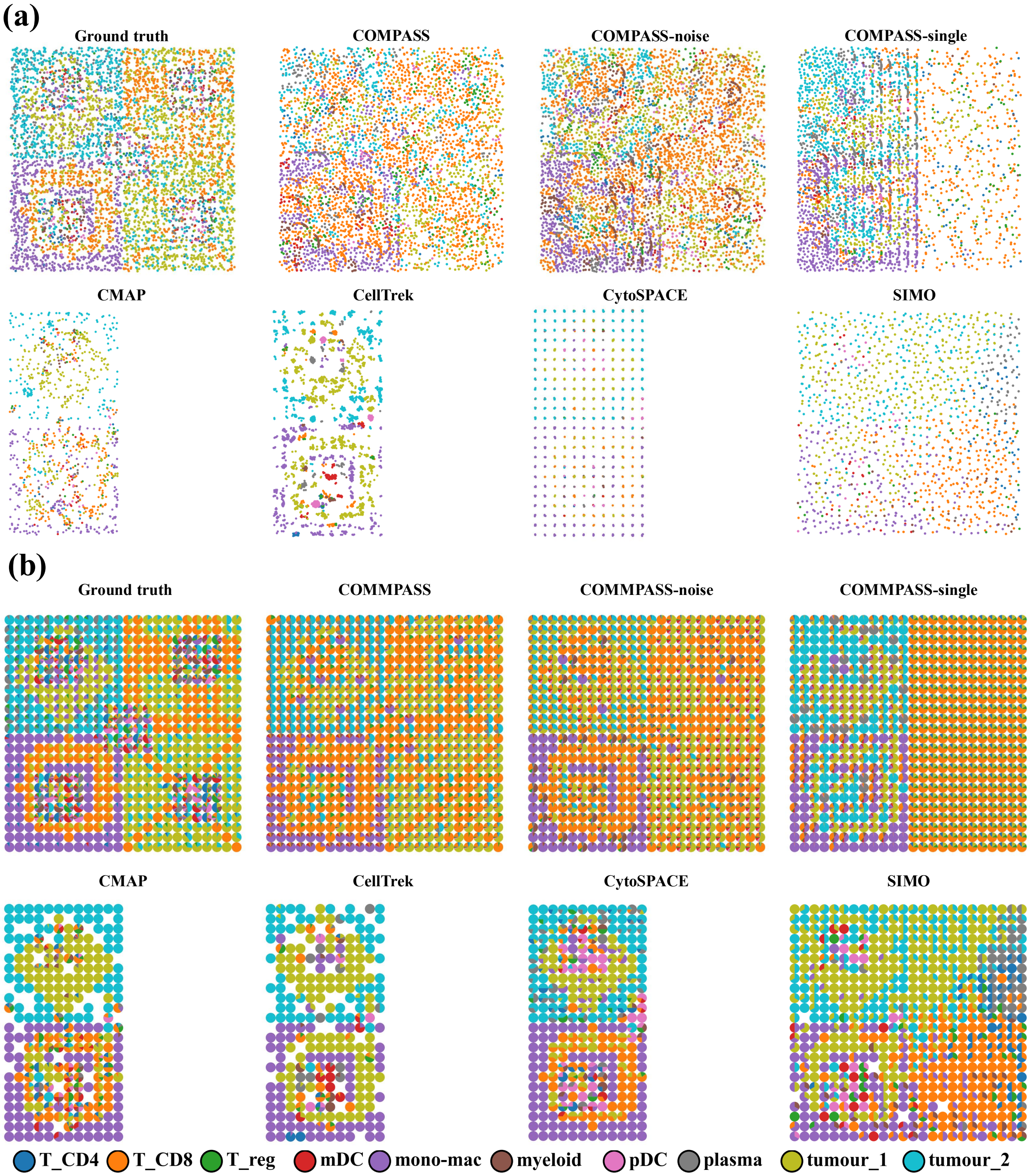

Figure S14: **Mapping-based reconstruction in Scenario 4 simulations.** Mapping-based reconstruction in Scenario 4 simulations. Spatial maps compare COMPASS variants and representative cell-mapping baselines in the most challenging setting, combining spatial RNA corruption, partial RNA signal loss, and reduced scRNA-seq reference coverage. (a) Cell-level maps show continuous coordinate reconstruction. (b) Spot-level maps aggregate mapped cells within pseudo-spots to summarize local composition. Half-width panels for some baselines indicate that valid mappings were recovered only in the RNA-retaining portion of the lattice, whereas RNA-masked regions lacked informative RNA-based assignments.

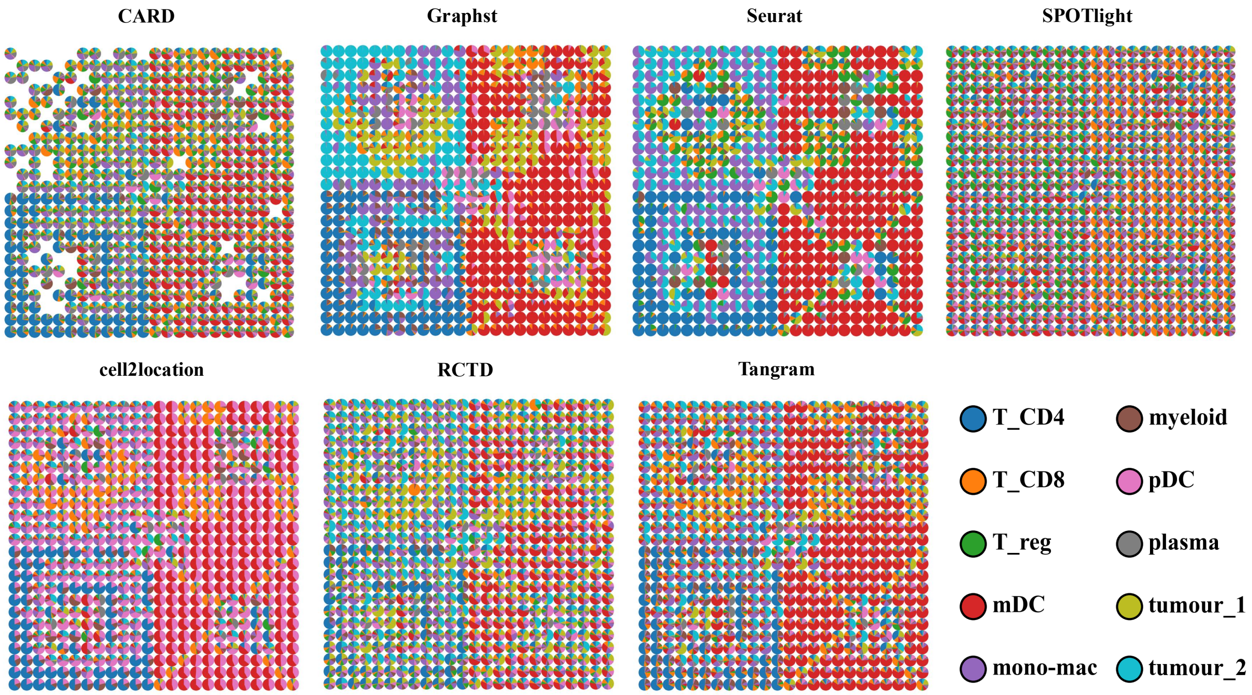

Figure S15: **Deconvolution-based composition recovery in Scenario 1 simulations.** Pie-chart maps show spot-level cell-type proportions inferred by deconvolution methods under the unperturbed setting. Each pie represents one pseudo-spot, and slice colors indicate the relative abundance of the ten simulated cell types. Blank locations in the CARD panel denote pseudo-spots for which CARD did not return a valid positive cell-type composition under the default deconvolution workflow; these positions are kept blank rather than imputed, so that incomplete spot-level recovery remains visible. The figure evaluates the recovery of regional composition patterns under ideal input conditions.

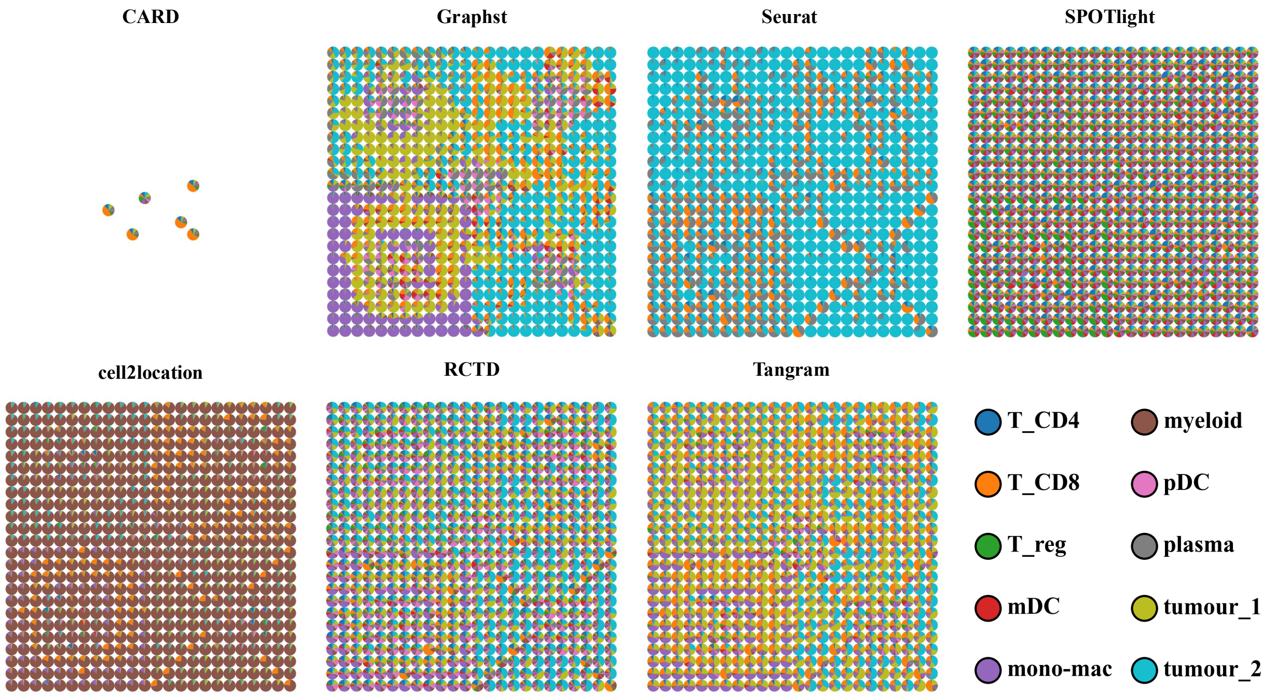

Figure S16: **Deconvolution-based composition recovery in Scenario 2 simulations.** Pie-chart maps show spot-level cell-type proportions inferred by deconvolution methods under spatial RNA noise. Each pseudo-spot is represented by a pie chart, with slices denoting the estimated abundance of the ten simulated cell types. Under RNA corruption, CARD produced valid non-zero composition estimates for only a small number of pseudo-spots, leaving the remaining lattice positions blank; this sparse output reflects the collapse of CARD-based composition recovery under noisy spatial RNA rather than missing simulated spots. The figure evaluates the robustness of composition recovery when spatial transcriptomic measurements are corrupted.

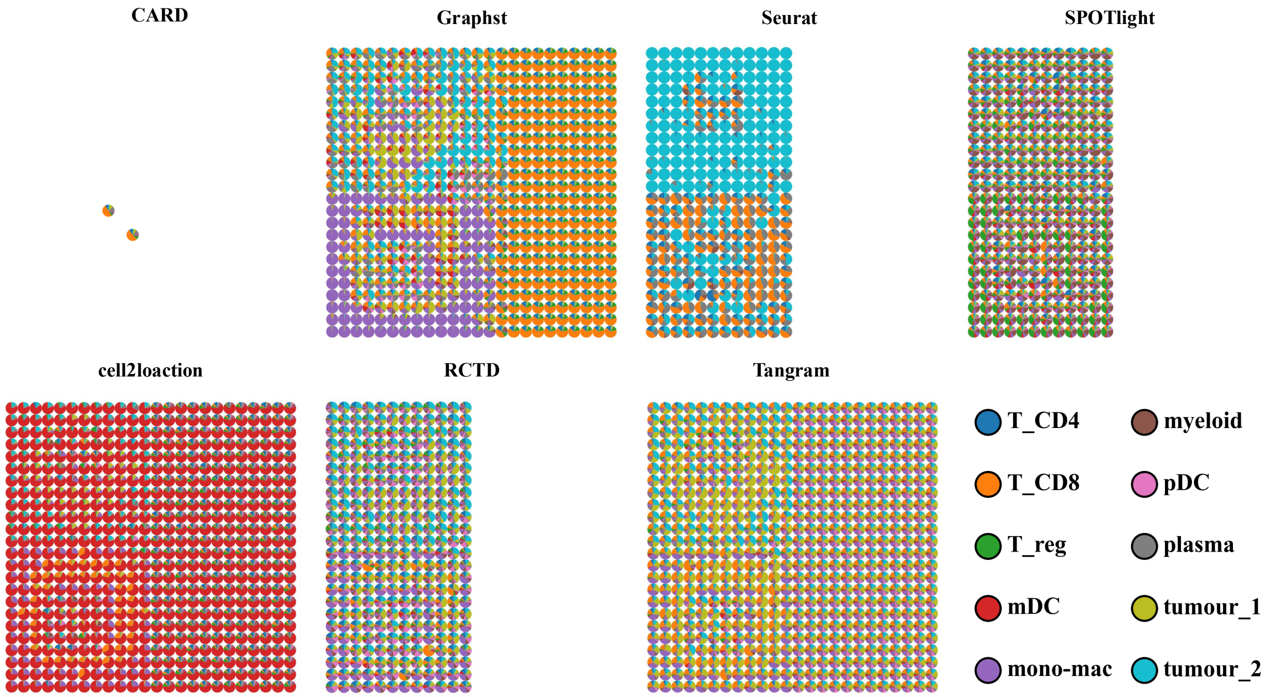

Figure S17: **Deconvolution-based composition recovery in Scenario 3 simulations.** Pie-chart maps show spot-level cell-type proportions inferred by deconvolution methods after spatial RNA noise and partial spot-level RNA masking. Each pie represents one pseudo-spot, and slice colors indicate the estimated mixture of the ten simulated cell types. The figure tests whether deconvolution methods preserve regional composition structure under severe RNA signal loss.

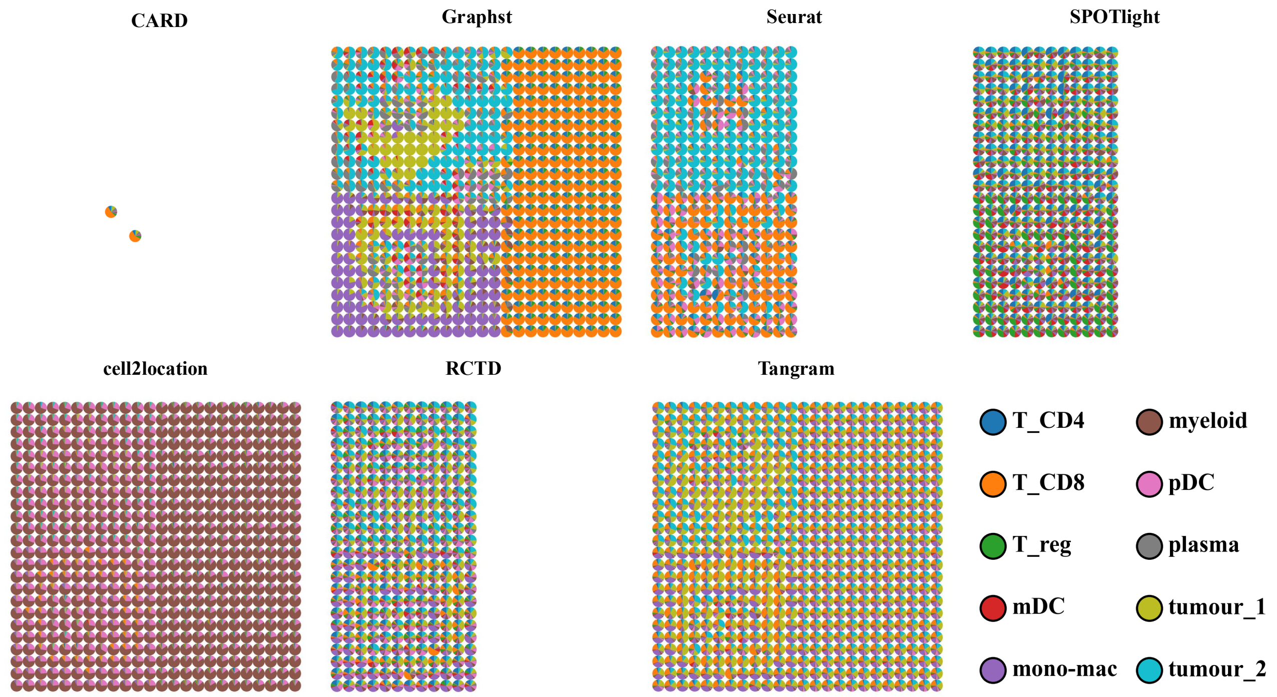

Figure S18: **Deconvolution-based composition recovery in Scenario 4 simulations.** Pie-chart maps show spot-level cell-type proportions inferred by deconvolution methods under combined RNA corruption, RNA signal loss, and incomplete scRNA-seq reference coverage. Each pseudo-spot is represented by a pie chart colored by the ten simulated cell types. The figure evaluates whether deconvolution methods retain spatially coherent composition recovery under simultaneous measurement degradation and reference incompleteness.

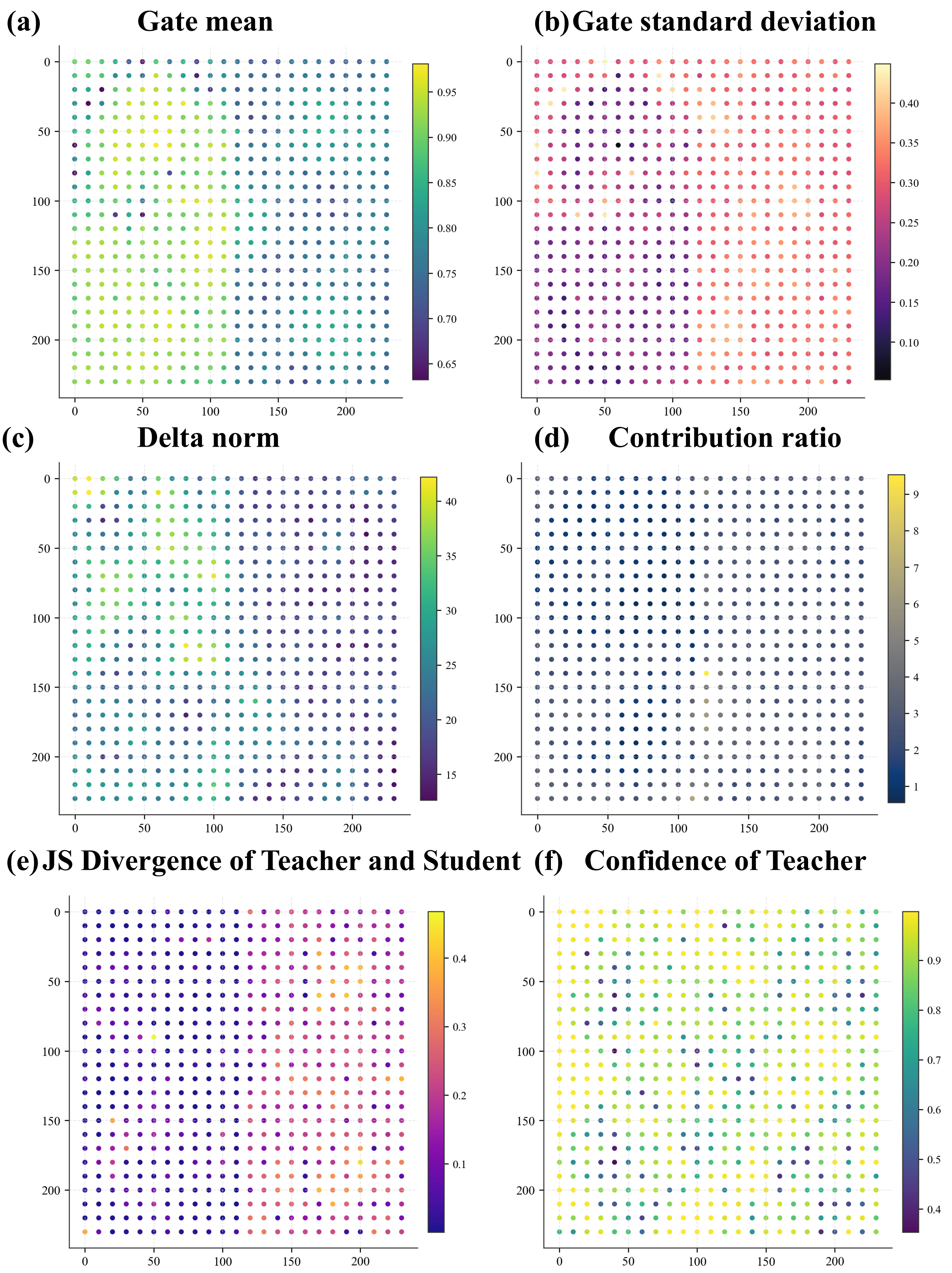
(a) Gate mean
(b) Gate standard deviation
(c) Delta norm
(d) Contribution ratio
(e) JS Divergence of Teacher and Student
(f) Confidence of Teacher

Figure S19: **Spatial distribution of RG-COD diagnostic scores in Scenario 1 simulations.** Spatial maps show spot-wise diagnostic quantities used to characterize residual-gated cross-omics distillation in the unperturbed simulation setting. (a) Mean gate value across latent dimensions, measuring the overall strength of auxiliary ATAC injection into the RNA backbone. (b) Standard deviation of gate values across latent dimensions, reflecting dimension-specific variability in auxiliary contribution. (c) Norm of the auxiliary residual after projection and gating, quantifying the magnitude of cross-omics correction. (d) Residual contribution ratio, measuring the relative contribution of the auxiliary residual to the RNA representation. (e) Jensen–Shannon divergence between teacher and student predictions, indicating local disagreement between the fused multi-omics teacher and the RNA-only student. (f) Teacher confidence, measuring the maximum predicted probability of the fused teacher branch across inferred domains.

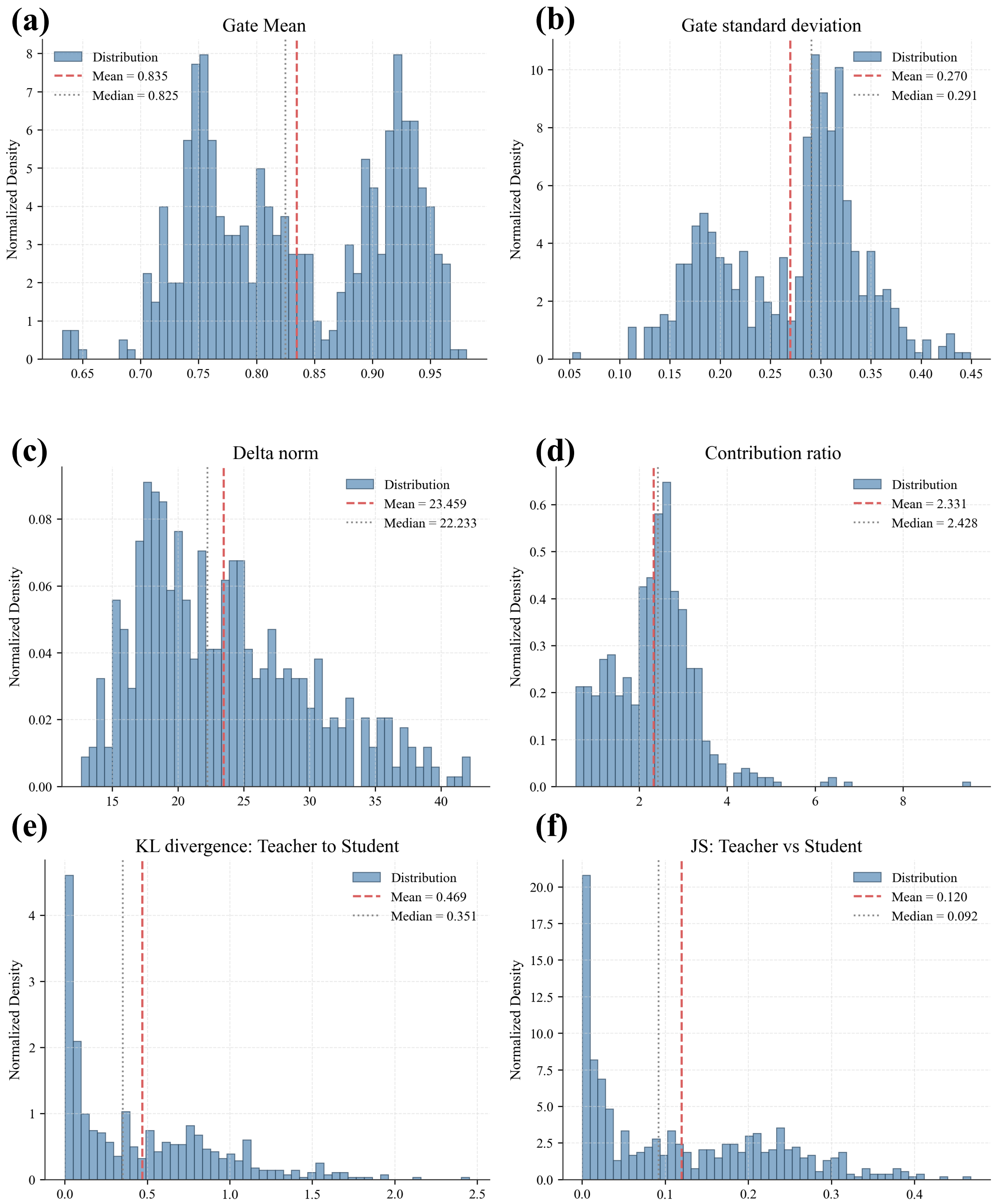

Figure S20: **Global distributions of RG-COD diagnostic scores in Scenario 1 simulations.** Histograms summarize the spot-wise distributions of the RG-COD diagnostic quantities shown in Supplementary Fig. SX. (a) Distribution of mean gate values across spots. (b) Distribution of gate standard deviations across latent dimensions. (c) Distribution of auxiliary residual norms. (d) Distribution of residual contribution ratios. (e) Distribution of teacher-to-student Kullback–Leibler divergence. (f) Distribution of teacher-to-student Jensen–Shannon divergence. Red dashed lines indicate the mean, and black dotted lines indicate the median.

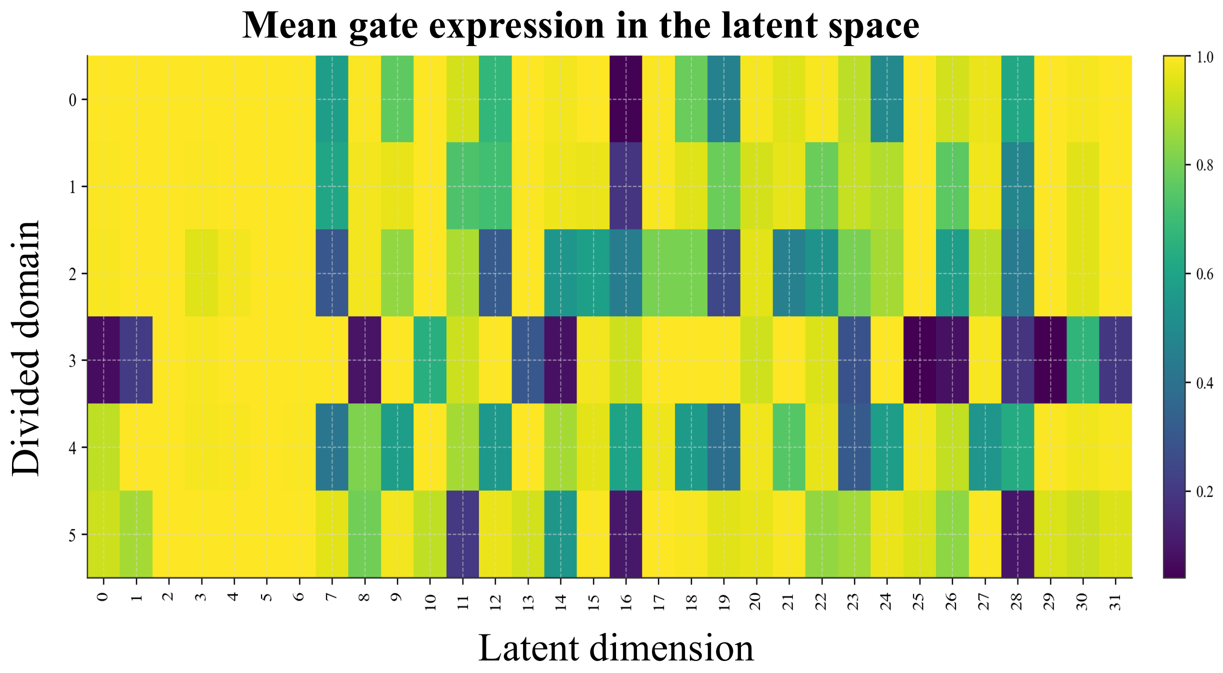

Figure S21: **Domain- and dimension-specific gate activity in Scenario 1 simulations.** Heatmap showing the mean gate value of RG-COD across inferred spatial domains and latent dimensions. Rows correspond to divided domains, and columns correspond to latent dimensions. The heterogeneous gate pattern indicates that auxiliary ATAC information is not uniformly fused into the RNA representation, but is selectively modulated across both tissue domains and representation dimensions.

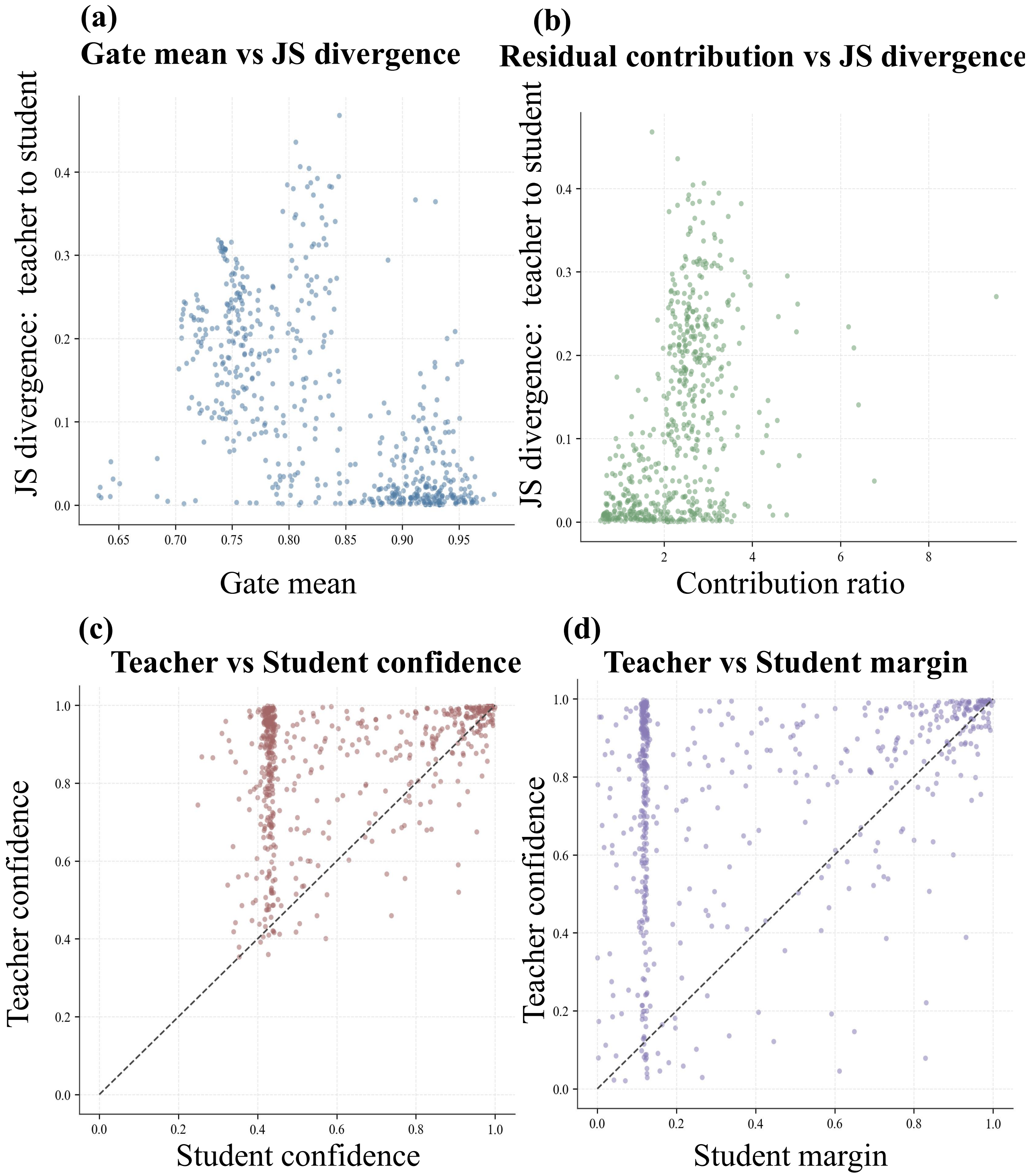

Figure S22: **Relationship between gated auxiliary transfer and teacher–student agreement in Scenario 1 simulations.** Scatter plots examine how RG-COD gating and residual contribution relate to teacher–student consistency. (a) Relationship between mean gate value and teacher–student Jensen–Shannon divergence. (b) Relationship between residual contribution ratio and teacher–student Jensen–Shannon divergence. (c) Comparison of teacher and student confidence scores, where the diagonal line indicates equal confidence. (d) Comparison of teacher and student prediction margins, where the diagonal line indicates equal margin. These analyses assess whether auxiliary-derived corrections are transferred to the RNA-only student without inducing systematic disagreement between the teacher and student branches.

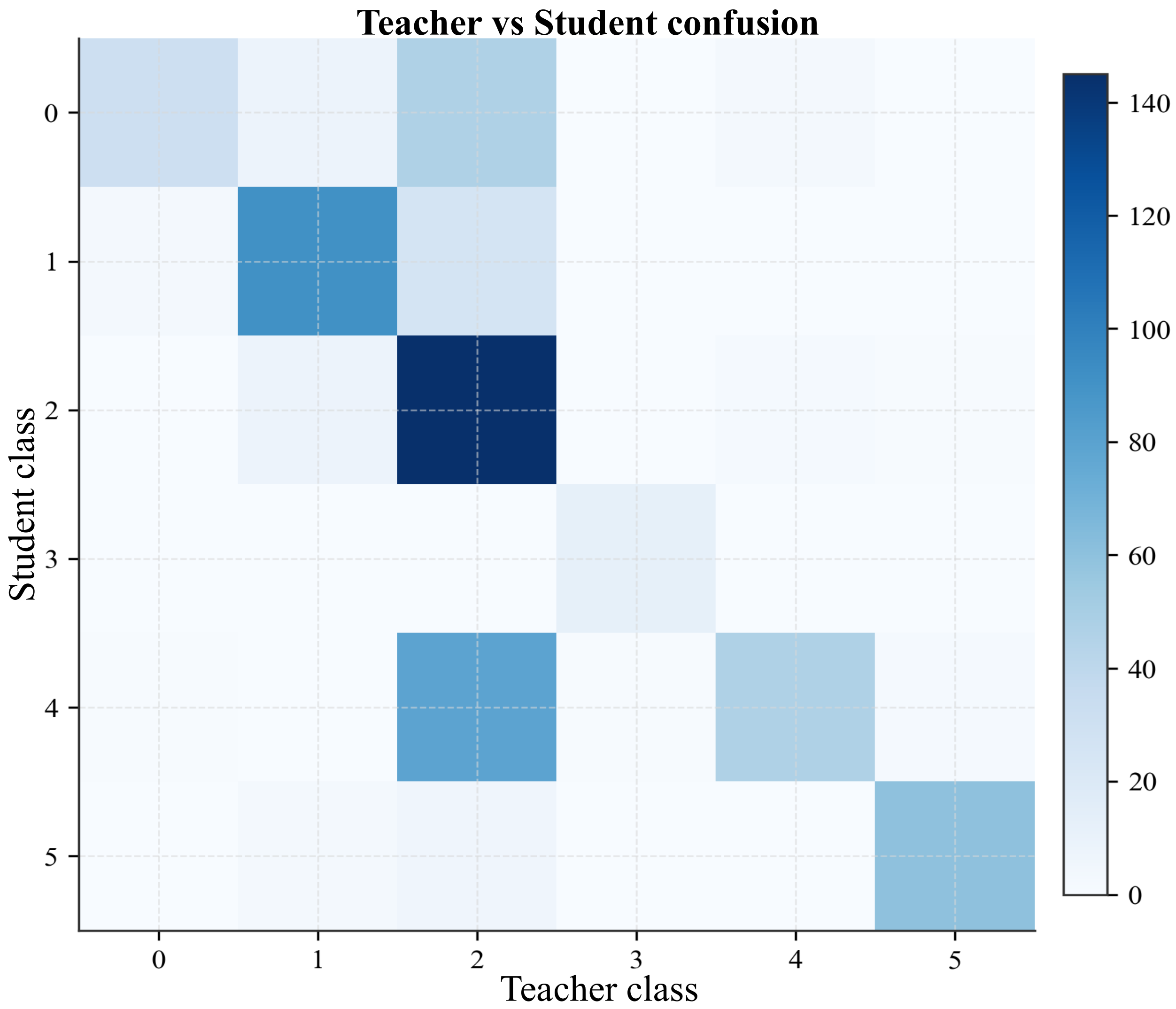

Figure S23: **Teacher–student prediction consistency in Scenario 1 simulations.** Confusion matrix comparing domain classes predicted by the fused multi-omics teacher and the RNA-only student after RG-COD distillation. Columns denote teacher-predicted classes and rows denote student-predicted classes, with color intensity representing the number of spots assigned to each teacher–student class pair. The matrix summarizes the extent to which auxiliary-derived domain structure learned by the teacher is transferred to the deployable RNA-only student while highlighting residual class-level discrepancies.

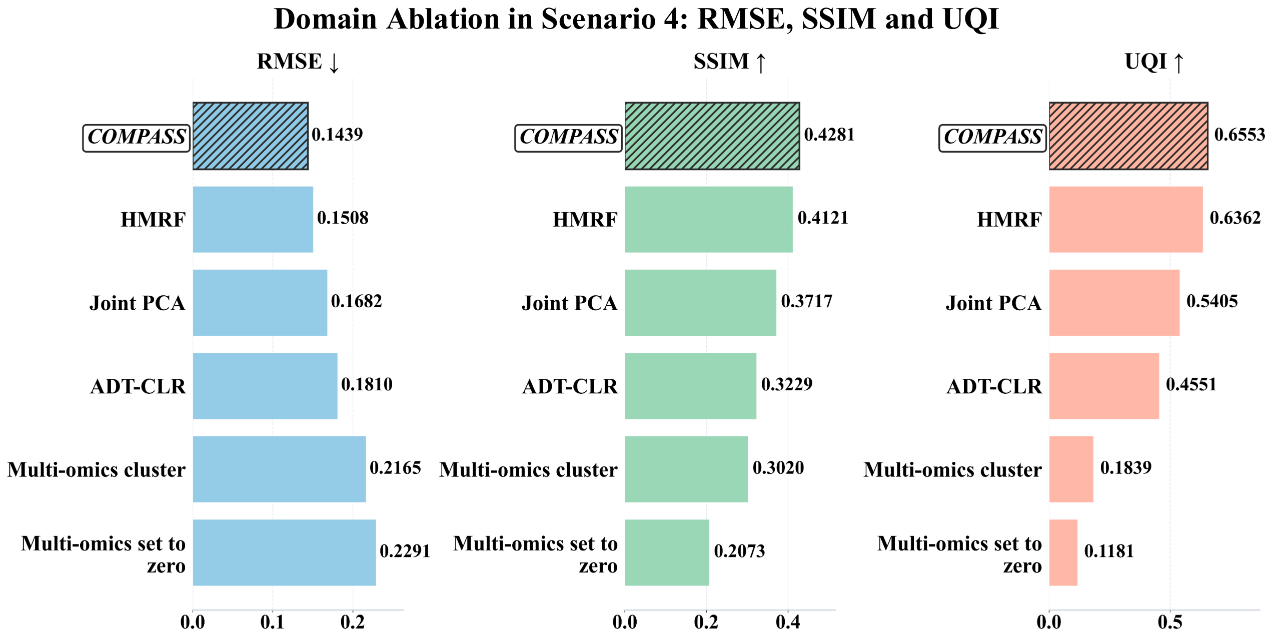

Figure S24: **Ablation of domain-prior construction in Scenario 4 evaluated by root mean square error (RMSE), structural similarity index (SSIM), and universal quality index (UQI).** Bar plots compare the full COMPASS model with variants perturbing hidden Markov random field (HMRF) smoothing, joint principal component analysis (PCA) of RNA and auxiliary omics, auxiliary centered log-ratio (CLR) normalization, multi-omics domain clustering, and complete removal of auxiliary multi-omics input. Lower RMSE and higher SSIM and UQI indicate better spot-level cell-type composition recovery.

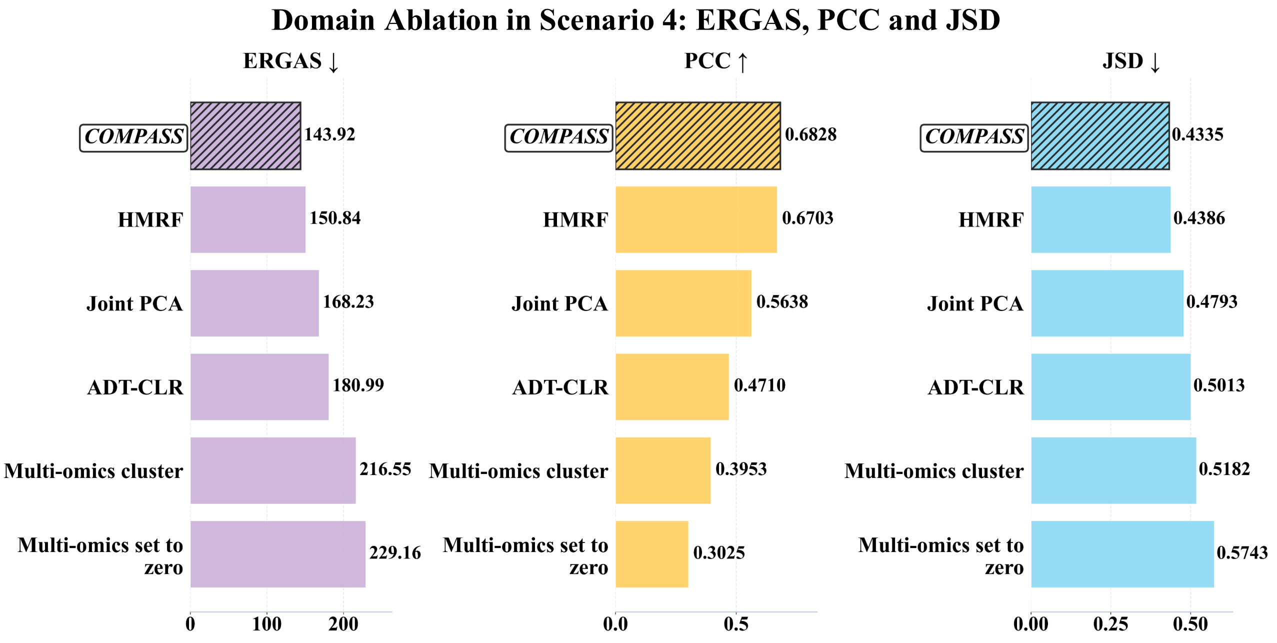

Figure S25: **Ablation of domain-prior construction in Scenario 4 evaluated by relative global dimensional synthesis error (ERGAS), Pearson correlation coefficient (PCC), and Jensen-Shannon divergence (JSD).** Bar plots assess how perturbing HMRF smoothing, joint RNA-auxiliary PCA, auxiliary CLR normalization, multi-omics clustering, and auxiliary-input removal affects global composition fidelity. Lower ERGAS and JSD and higher PCC indicate better agreement with ground-truth spot-level cell-type proportions.

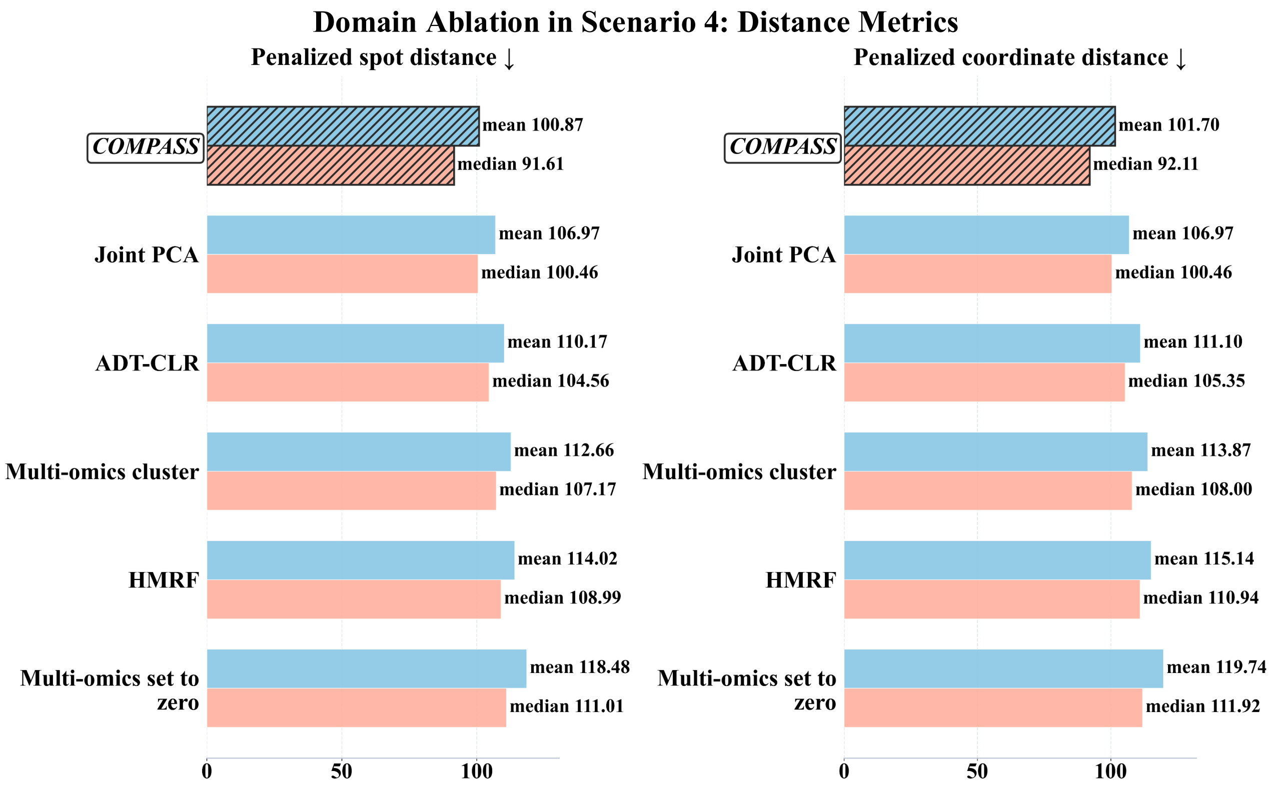

Figure S26: **Ablation of domain-prior construction in Scenario 4 evaluated by penalized spot distance (PSD) and penalized coordinate distance (PCD).** Bar plots show cell-level mapping errors after perturbing domain construction components, including spatial smoothing, joint multi-omics dimensional reduction, auxiliary-modality preprocessing, and auxiliary-aware domain initialization. Lower mean and median PSD or PCD indicate more accurate cell-to-spot assignment and continuous coordinate reconstruction.

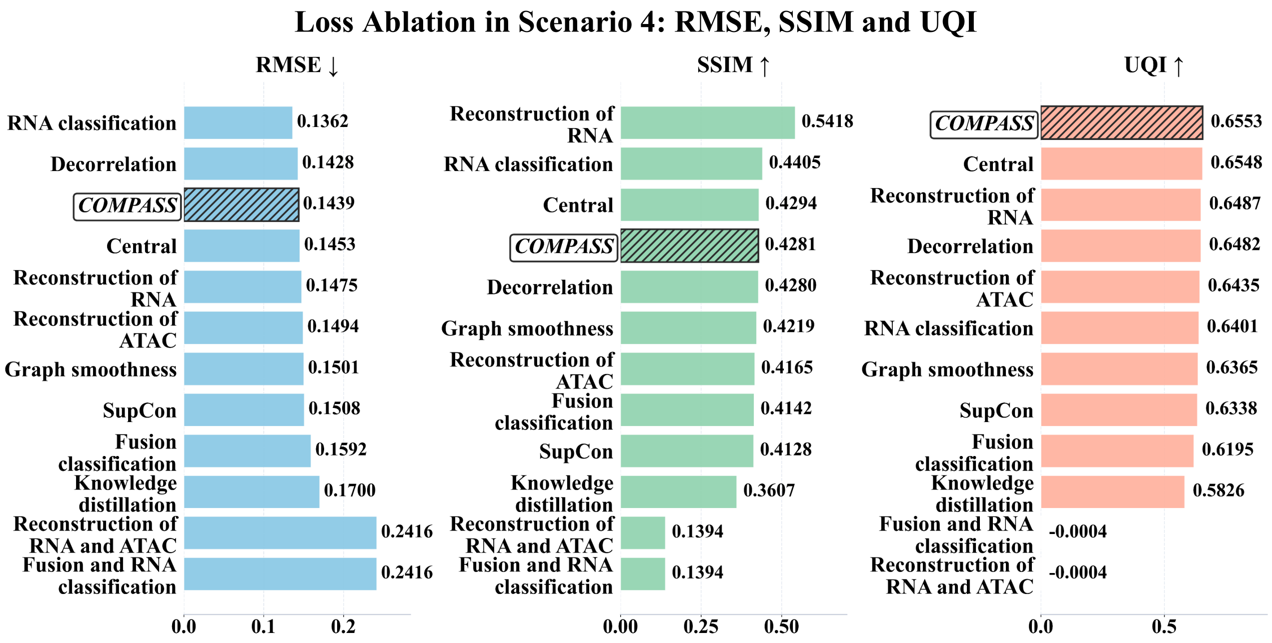

Figure S27: **Ablation of the multi-objective training loss in Scenario 4 evaluated by RMSE, SSIM, and UQI.** Bar plots compare COMPASS with variants removing RNA-domain classification, fused-teacher classification, knowledge distillation, RNA reconstruction, ATAC reconstruction, graph smoothness, supervised contrastive learning, center loss, decorrelation regularization, or paired reconstruction and classification terms. Lower RMSE and higher SSIM and UQI indicate better recovery of ground-truth cell-type proportions.

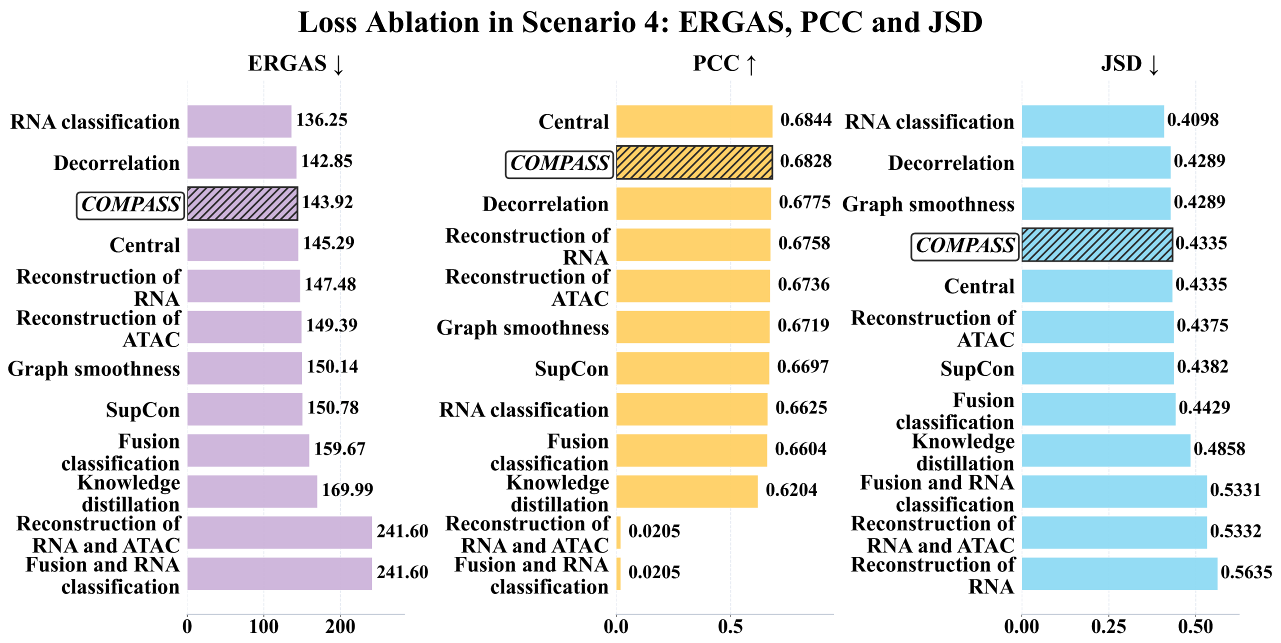

Figure S28: **Ablation of the multi-objective training loss in Scenario 4 evaluated by ERGAS, PCC, and JSD.** Bar plots quantify how removing reconstruction, classification, distillation, smoothness, contrastive, center, and decorrelation losses affects distributional and correlation-based composition accuracy. Lower ERGAS and JSD and higher PCC indicate stronger agreement with ground-truth spot-level proportions.

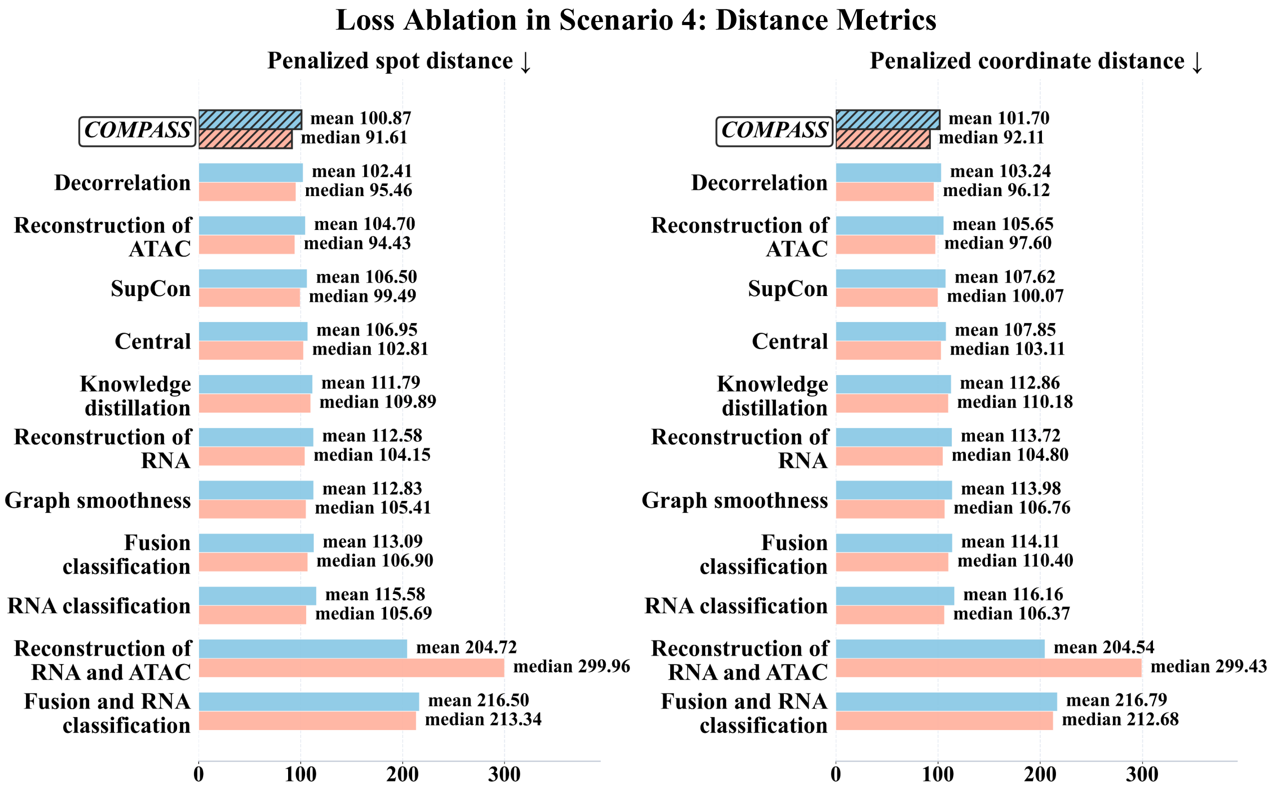

Figure S29: **Ablation of the multi-objective training loss in Scenario 4 evaluated by PSD and PCD.** Bar plots show cell-level mapping errors after removing individual or coupled loss terms, including reconstruction, domain classification, knowledge distillation, graph smoothness, supervised contrastive learning, center loss, and decorrelation regularization. Lower mean and median PSD or PCD indicate better preservation of cell-level spatial localization.

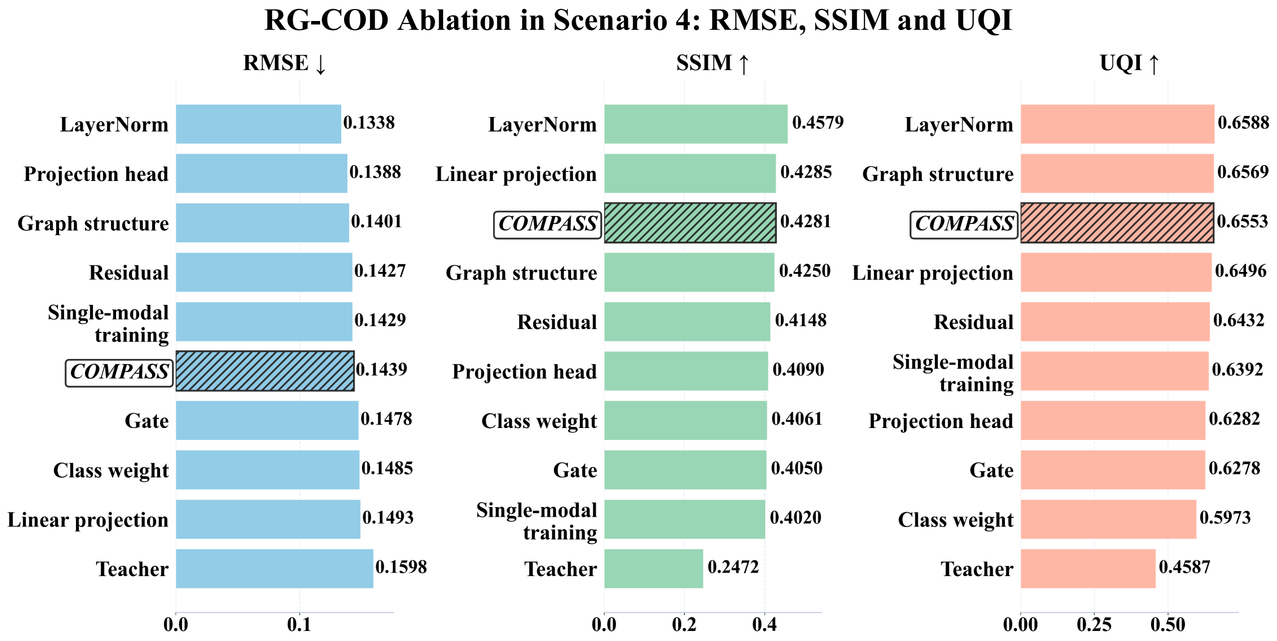

Figure S30: **Ablation of residual-gated cross-omics distillation (RG-COD) training in Scenario 4 evaluated by RMSE, SSIM, and UQI.** Bar plots compare COMPASS with variants perturbing the fused teacher branch, residual auxiliary injection, learnable gate, auxiliary-to-RNA linear projection, projection head, graph propagation, class-balanced weighting, layer normalization, and single-modality training. Lower RMSE and higher SSIM and UQI indicate better spot-level composition recovery.

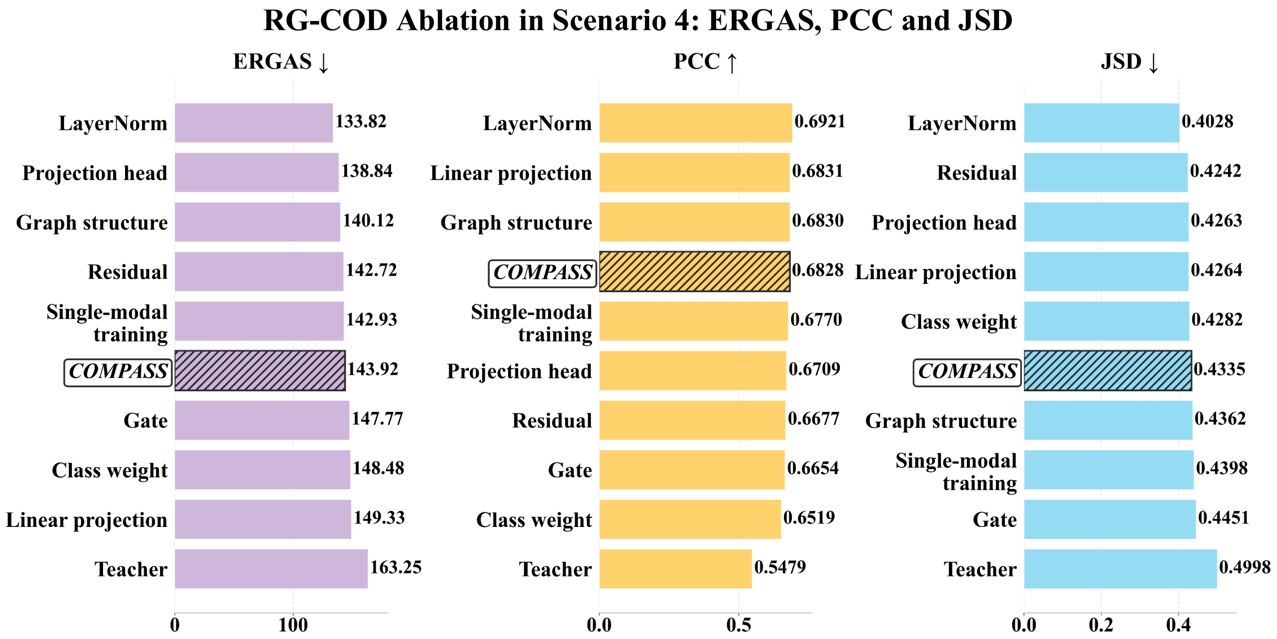

Figure S31: **Ablation of RG-COD training in Scenario 4 evaluated by ERGAS, PCC, and JSD.** Bar plots assess how perturbing the fused teacher branch, residual connection, learnable gate, auxiliary-to-RNA projection, graph structure, class-balanced weighting, layer normalization, and single-modality training affects global composition fidelity. Lower ERGAS and JSD and higher PCC indicate more accurate transfer of auxiliary-derived structure into the RNA-deployable representation.

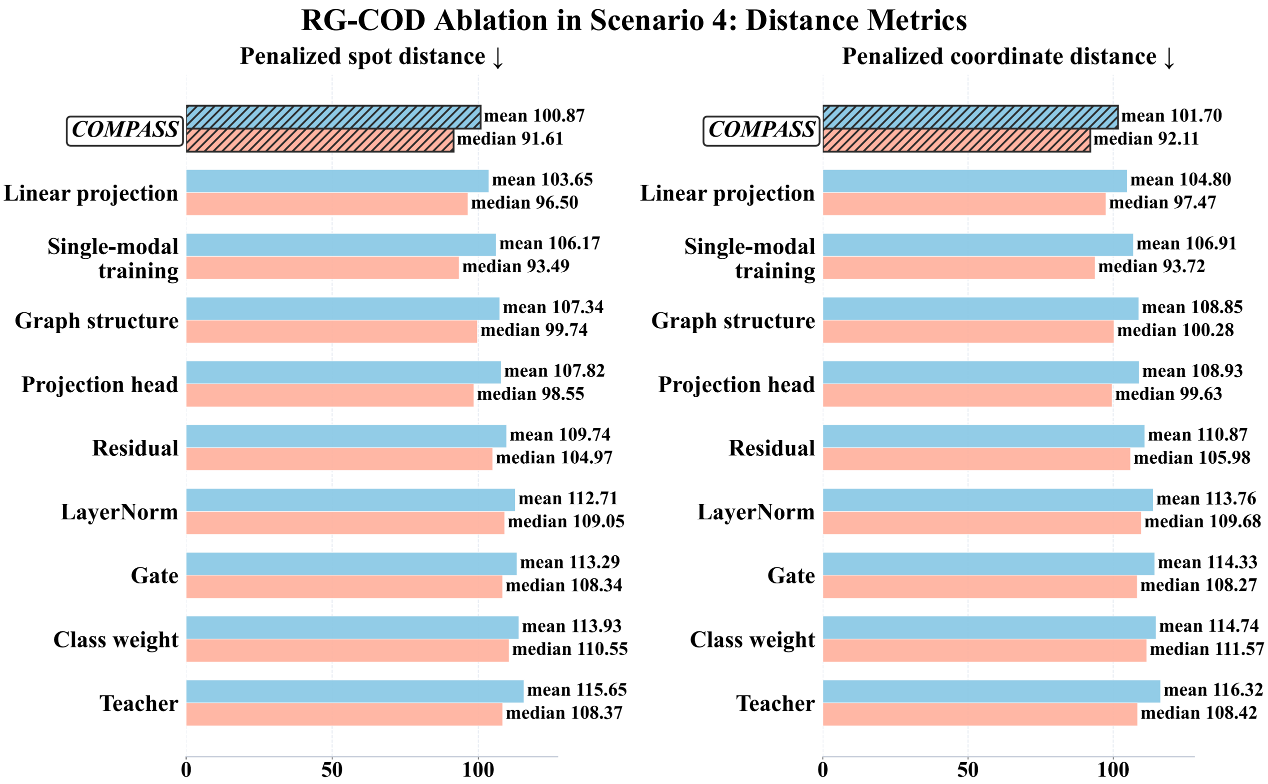

Figure S32: **Ablation of RG-COD training in Scenario 4 evaluated by PSD and PCD.** Bar plots compare cell-level mapping errors across RG-COD variants, including removal of the teacher branch, gate, residual correction, projection components, graph propagation, class weighting, layer normalization, and multi-omics training. Lower mean and median PSD or PCD indicate more accurate spatial assignment and sub-spot coordinate recovery.

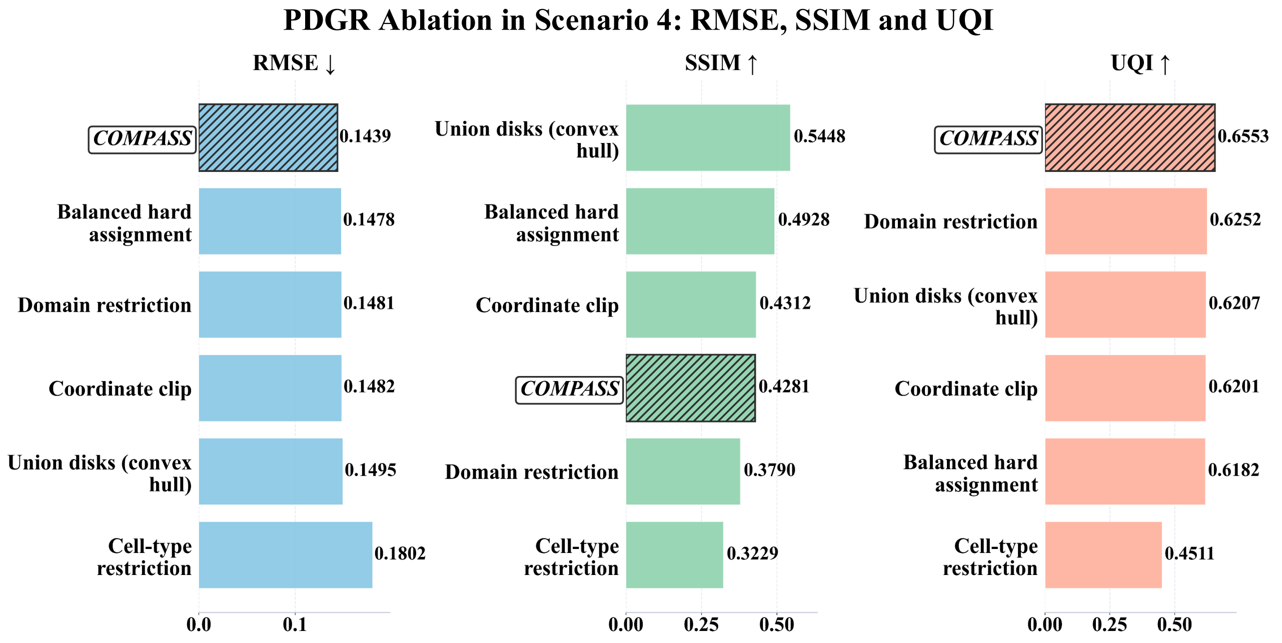

Figure S33: **Ablation of Poisson-disk geometric refinement (PDGR) in Scenario 4 evaluated by RMSE, SSIM, and UQI.** Bar plots compare COMPASS with geometric-refinement variants perturbing capacity-balanced hard assignment, domain-restricted placement, coordinate clipping, union-of-disks tissue support, and cell-type-aware coordinate allocation. Lower RMSE and higher SSIM and UQI indicate better spot-level composition recovery after continuous coordinate refinement.

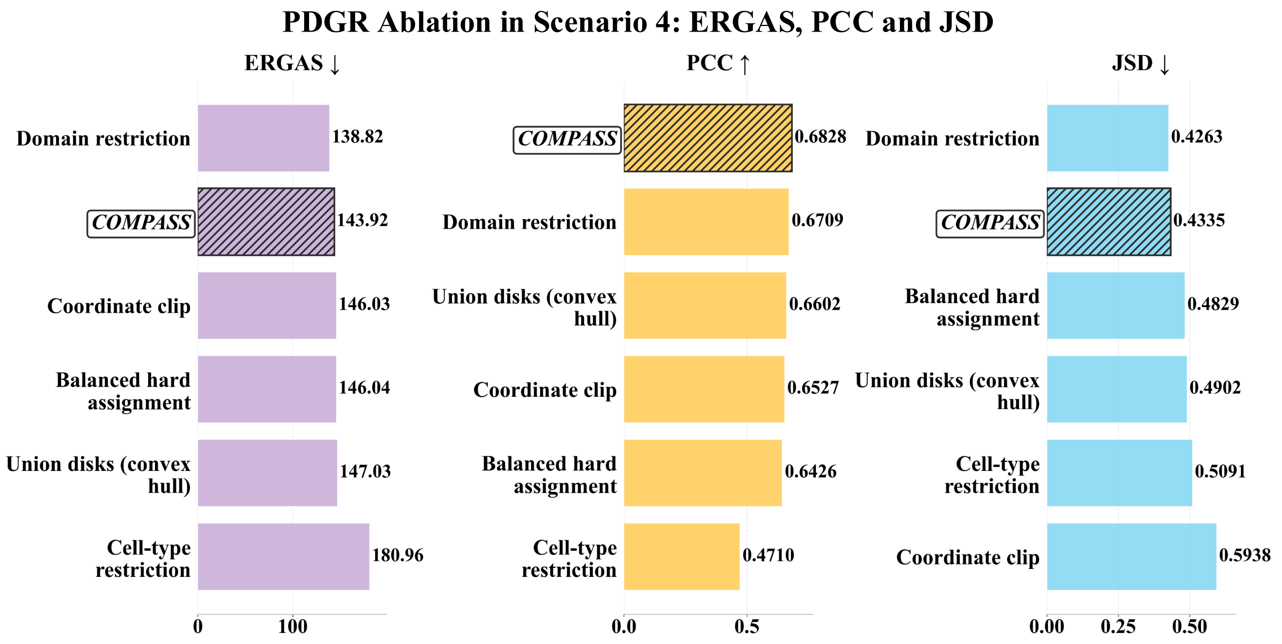

Figure S34: **Ablation of PDGR in Scenario 4 evaluated by ERGAS, PCC, and JSD.** Bar plots evaluate how capacity-balanced hard assignment, domain restriction, coordinate clipping, tissue-support definition, and cell-type-aware placement affect global composition fidelity after sub-spot coordinate reconstruction. Lower ERGAS and JSD and higher PCC indicate better preservation of ground-truth spot-level cell-type proportions.

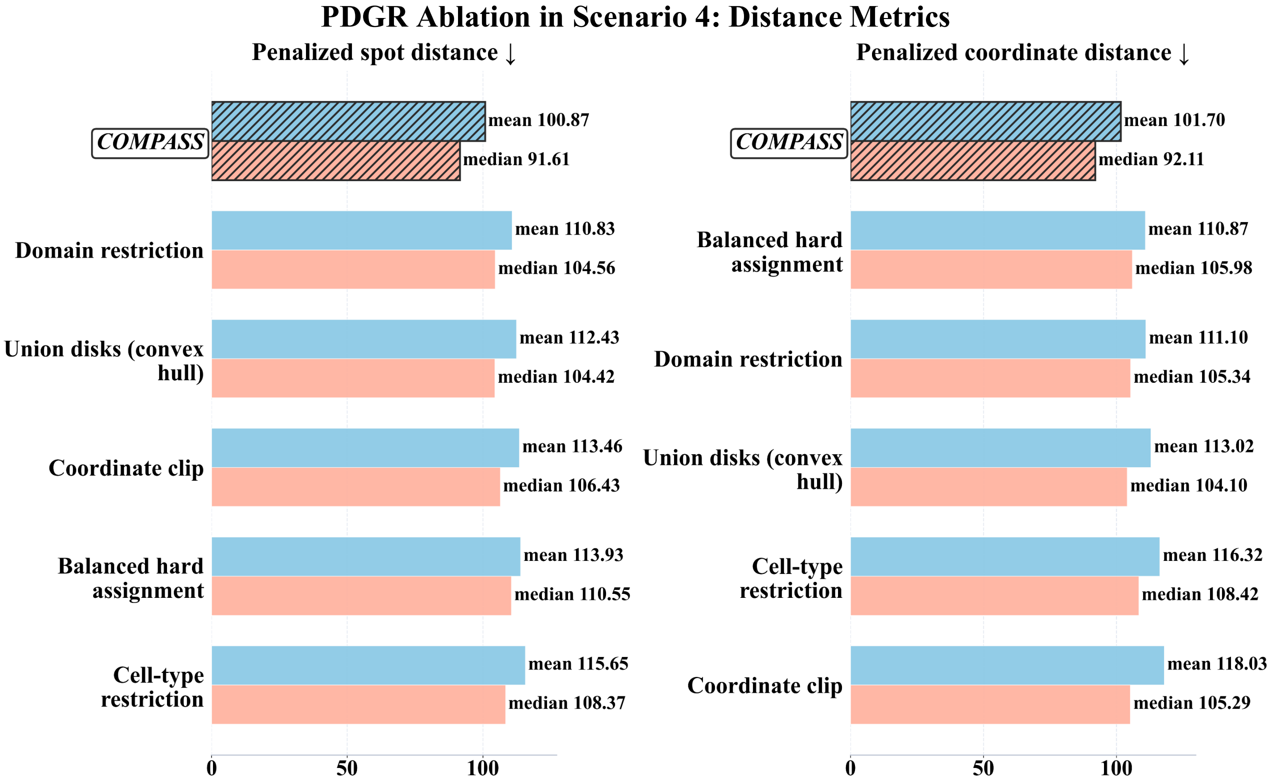

Figure S35: **Ablation of PDGR in Scenario 4 evaluated by PSD and PCD.** Bar plots show cell-level mapping errors for geometric-refinement variants, including balanced hard assignment, domain-restricted placement, coordinate clipping, replacement of union-of-disks support by convex-hull support, and removal of cell-type-aware placement. Lower mean and median PSD or PCD indicate more accurate and spatially constrained single-cell coordinate reconstruction.

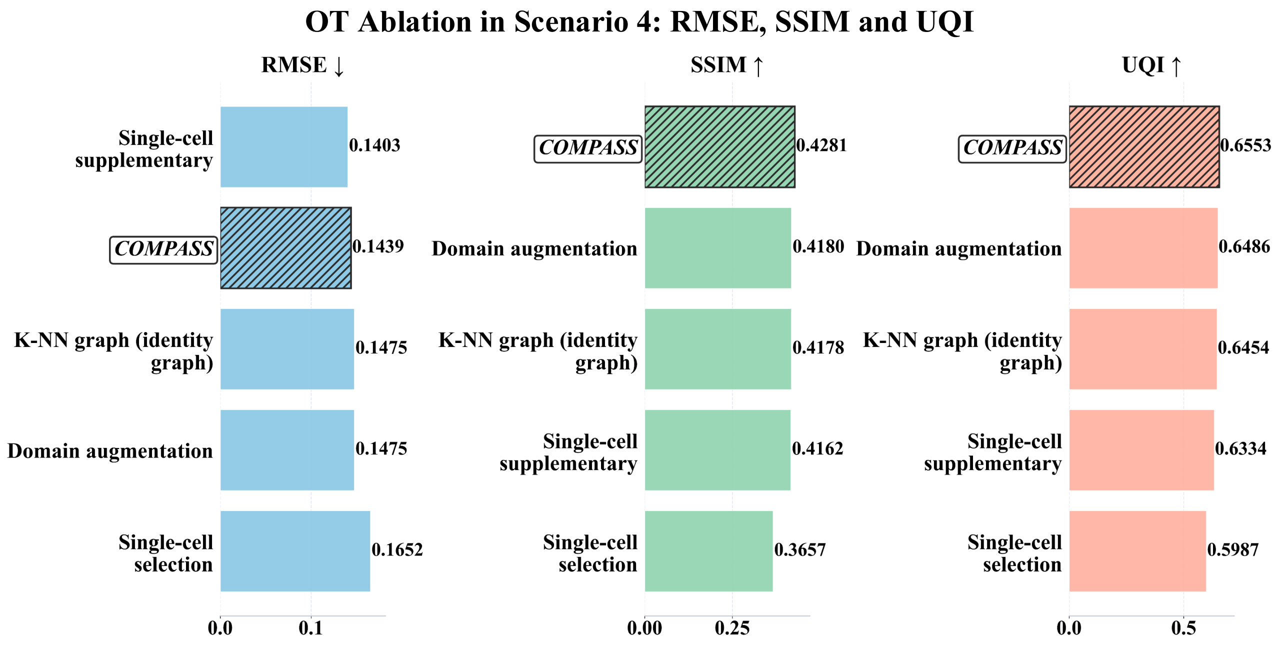

Figure S36: **Ablation of optimal-transport-based (OT-based) cell-to-spot assignment in Scenario 4 evaluated by RMSE, SSIM, and UQI.** Bar plots compare COMPASS with variants perturbing balanced single-cell selection, single-cell supplementation for sparse strata, domain augmentation, and the second-pass single-cell k-nearest-neighbor (kNN) graph. Lower RMSE and higher SSIM and UQI indicate better spot-level composition recovery after assignment.

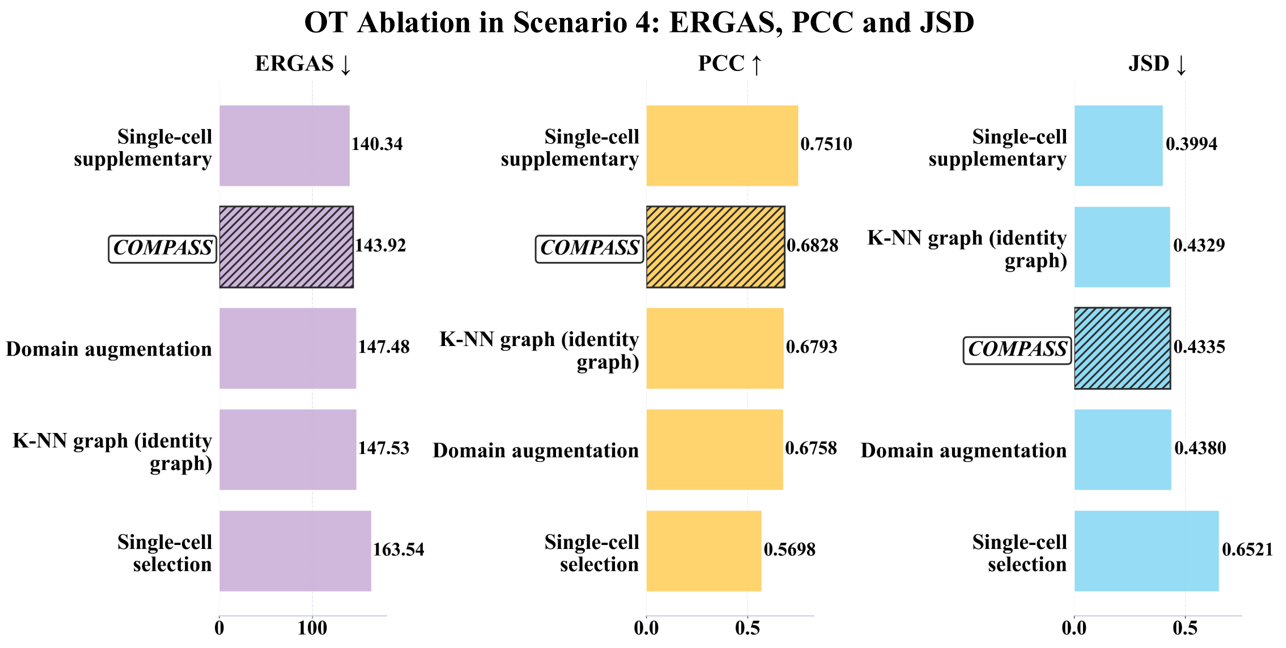

Figure S37: **Ablation of OT-based cell-to-spot assignment in Scenario 4 evaluated by ERGAS, PCC, and JSD.** Bar plots assess how balanced CellType-by-domain sampling, single-cell supplementation, domain augmentation, and second-pass kNN graph refinement affect distributional and correlation-based composition accuracy. Lower ERGAS and JSD and higher PCC indicate stronger agreement with ground-truth spot-level proportions.

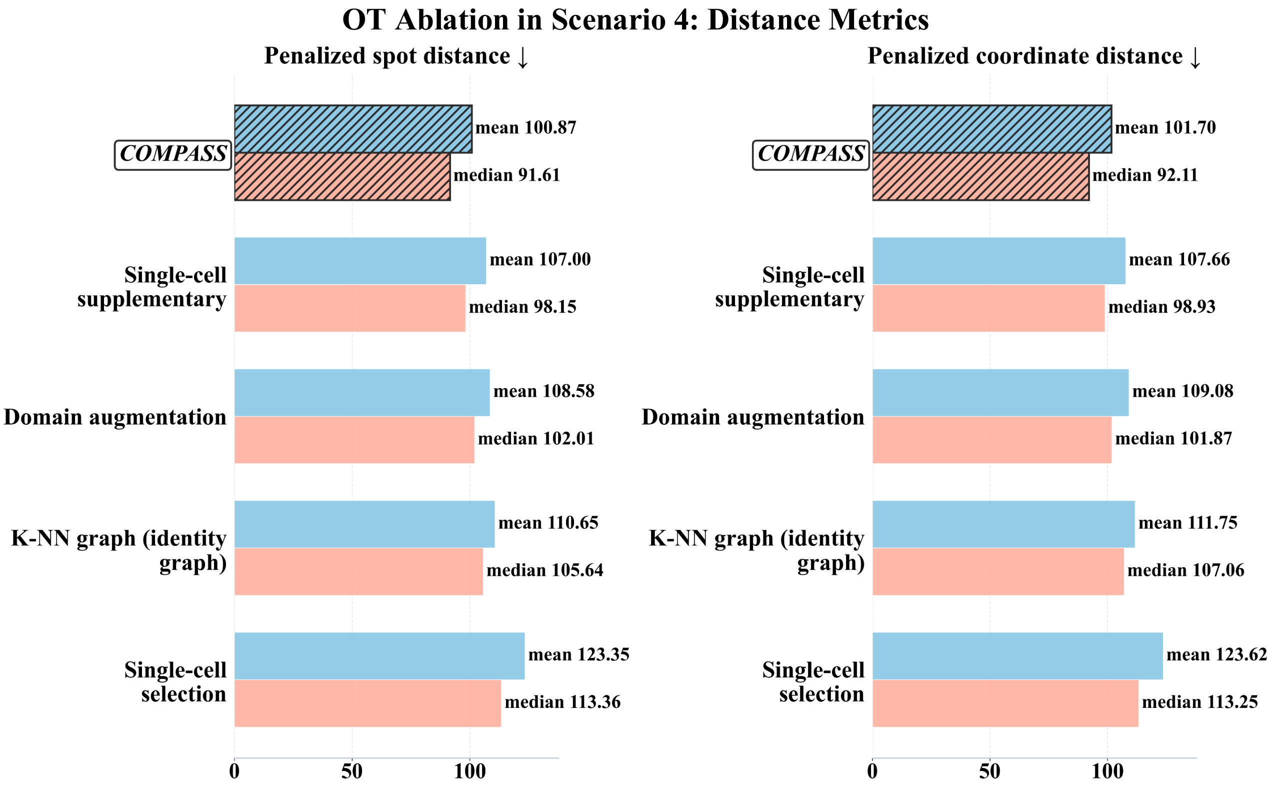

Figure S38: **Ablation of OT-based cell-to-spot assignment in Scenario 4 evaluated by PSD and PCD.** Bar plots show cell-level mapping errors for assignment-stage variants, including balanced single-cell selection, single-cell supplementation, domain augmentation, and replacement of the second-pass kNN graph by an identity graph. Lower mean and median PSD or PCD indicate more accurate conversion of learned embeddings into spatially valid cell-to-spot correspondences.

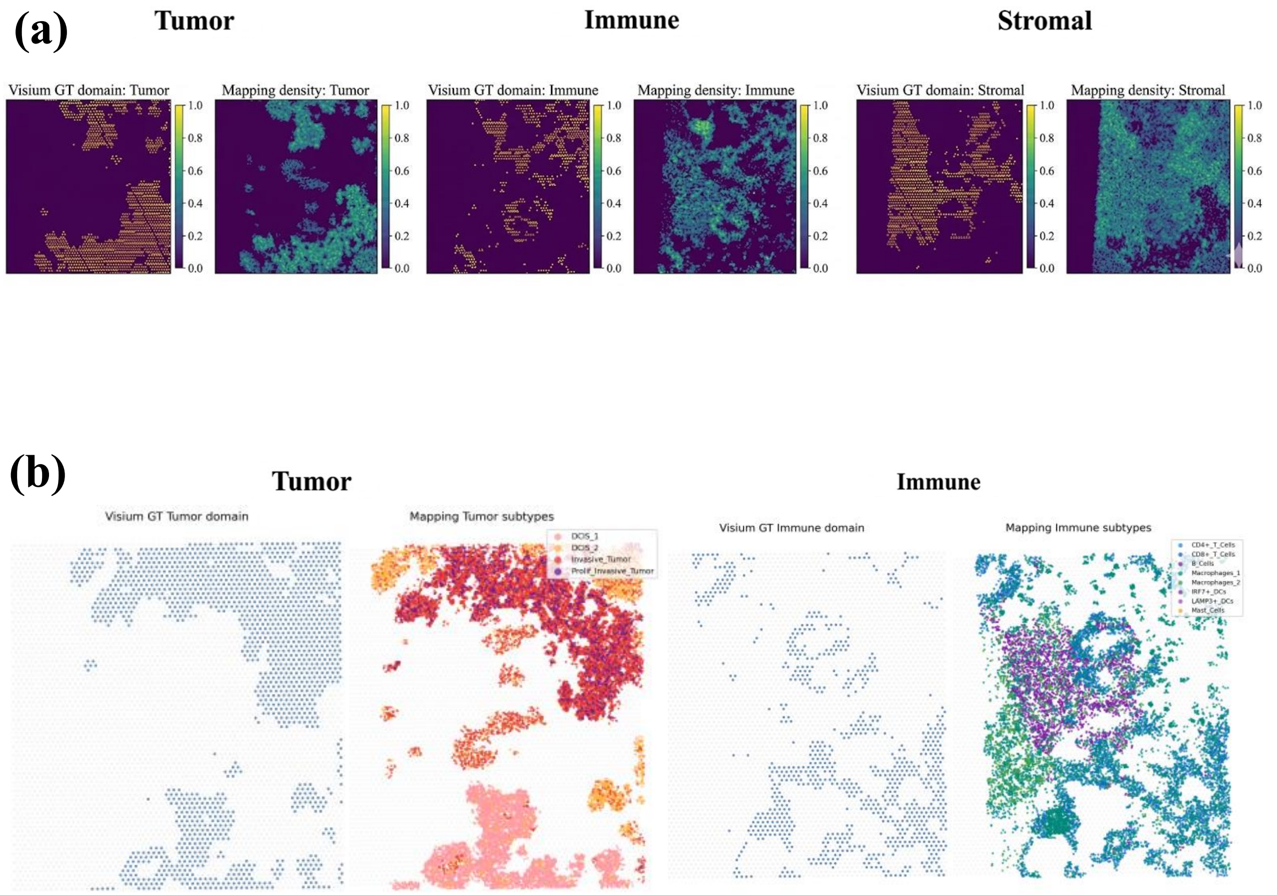

Figure S39: **Mapped single-cell densities recover tumour, immune and stromal tissue compartments.** (a) Visium ground-truth domain masks for tumour, immune and stromal regions are compared with the corresponding mapped-cell density maps. The comparison evaluates whether reconstructed cells are concentrated in the correct coarse tissue compartments rather than dispersed across incompatible regions. (b) Ground-truth tumour and immune domains are shown alongside mapped subtype-level distributions. Tumour subtypes, including DCIS and invasive tumour states, and immune subtypes, including T cells, B cells, macrophages, and dendritic-cell populations, retain domain-specific spatial localization within the reconstructed atlas.

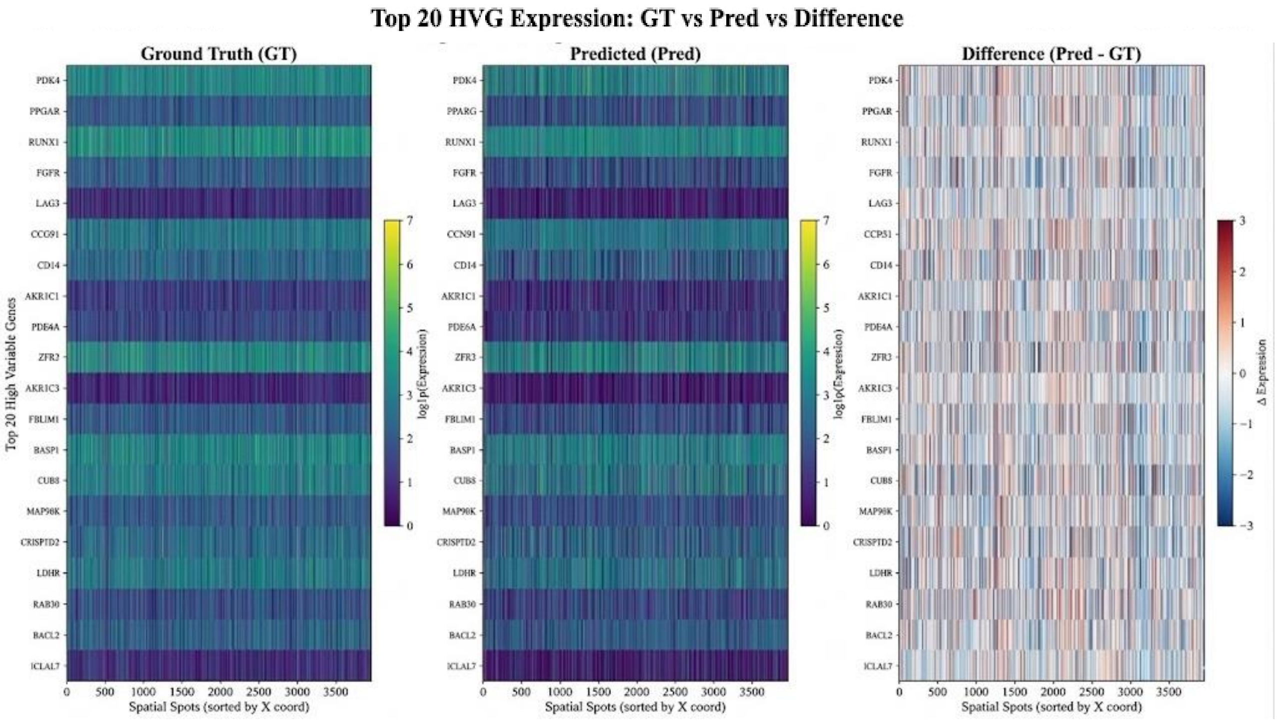

Figure S40: **Spatial expression recovery of highly variable genes in the Xenium-validated benchmark.** Heatmaps compare the Xenium-binned ground-truth expression, predicted expression and residual expression for the top 20 highly variable genes across spatial spots ordered by x coordinate. Expression values are shown on a log-transformed scale for the ground-truth and predicted matrices, while the residual heatmap shows prediction minus ground truth. Similar spatial expression patterns between the ground-truth and predicted matrices, together with residuals centered near zero, indicate recovery of spatially structured transcriptional programs in the reconstructed atlas.

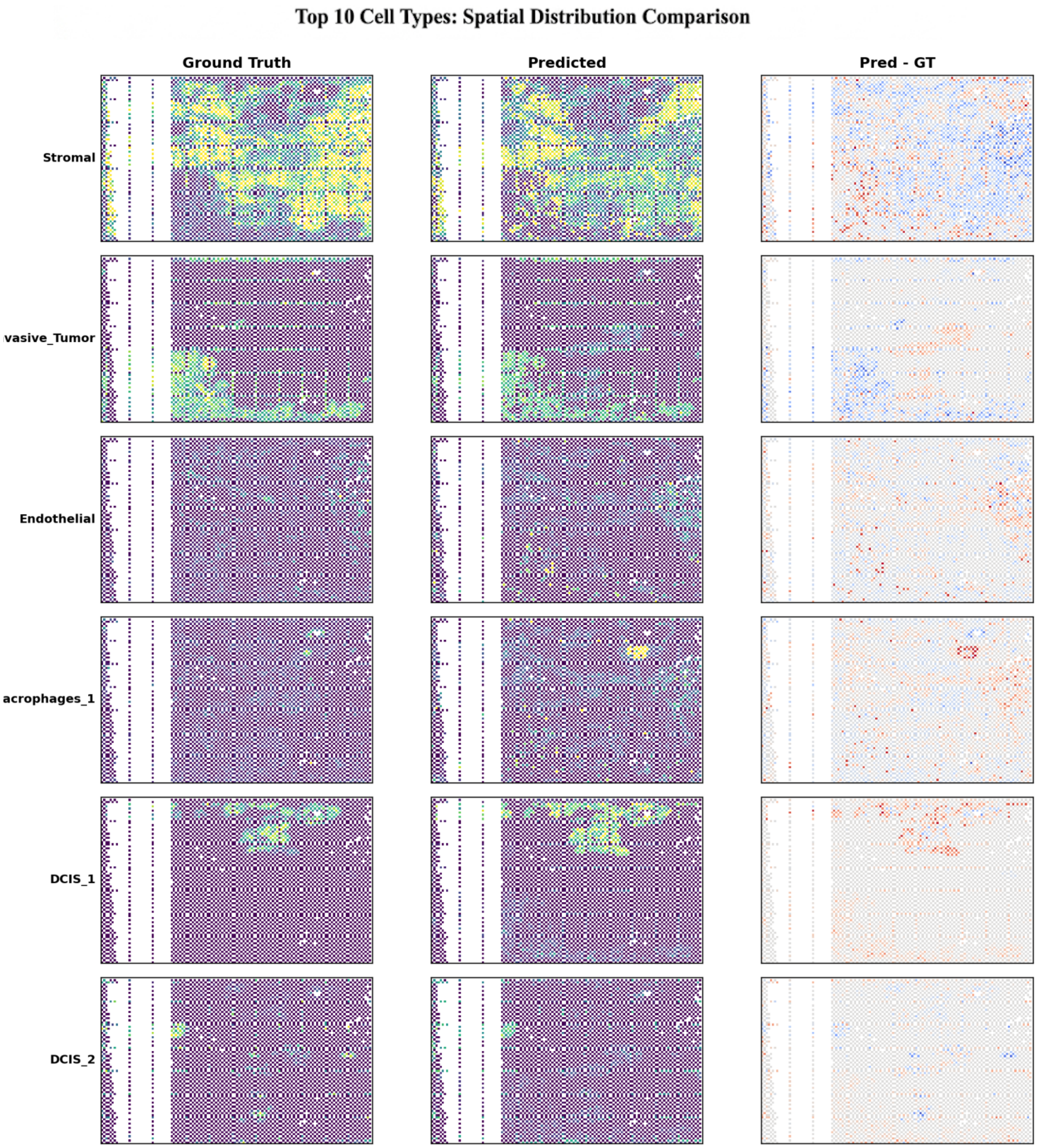

Figure S41: **Spatial distribution comparison of major cell types between Xenium-derived ground truth and reconstructed predictions.** For each of the top cell types, the left column shows the Xenium-binned ground-truth spatial proportion map, the middle column shows the predicted spatial proportion map, and the right column shows the residual map computed as prediction minus ground truth. The comparison assesses whether reconstructed cell-type proportions preserve both broad regional localization and fine-scale spatial heterogeneity. Residual maps highlight local over-prediction and under-prediction, with small and spatially diffuse residuals indicating closer agreement with the Xenium reference.

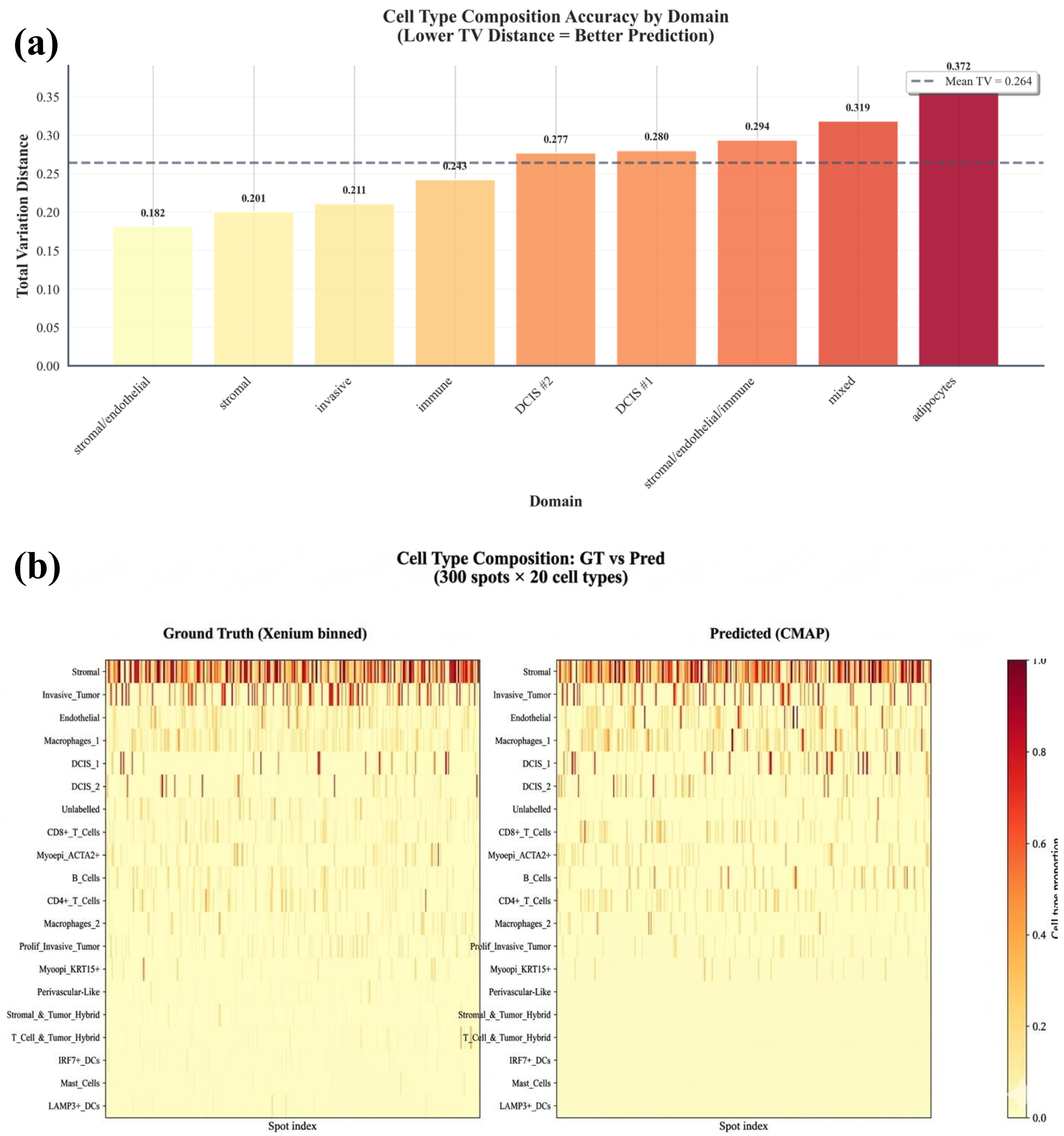

Figure S42: **Domain-resolved cell-type composition accuracy against Xenium-binned ground truth.** (a) Total variation distance was computed between predicted and ground-truth cell-type compositions within each tissue domain, with lower values indicating more accurate composition recovery. The dashed line denotes the mean total variation distance across domains. Stromal/endothelial, stromal and invasive domains showed the lowest errors, whereas mixed and adipocyte-enriched regions were more difficult to reconstruct. (b) Heatmaps compare Xenium-binned ground-truth and predicted spot-level cell-type compositions across 300 spatial spots and 20 cell types. Rows denote cell types and columns denote spots, with colour intensity representing the cell-type proportion at each spot.

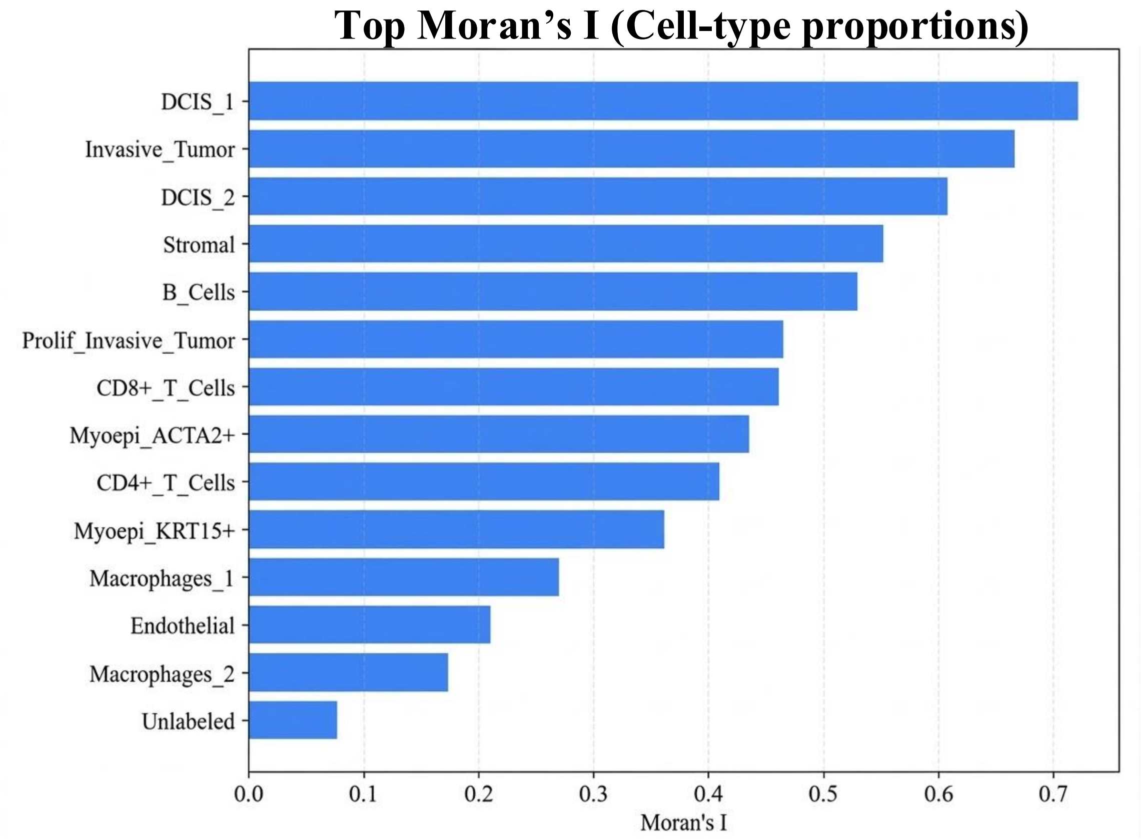

Figure S43: **Spatial autocorrelation of reconstructed cell-type proportions in the Xenium-validated breast cancer benchmark.** Moran's I was computed for the spatial proportion map of each cell type to quantify spatial coherence across the reconstructed tissue. Higher values indicate stronger spatial clustering rather than random dispersion. Tumour epithelial populations, including DCIS_1, Invasive_Tumor and DCIS_2, showed the strongest spatial autocorrelation, followed by stromal, B-cell and T-cell populations, indicating that the reconstructed atlas preserves both tumour compartmentalization and immune-stromal spatial organization.

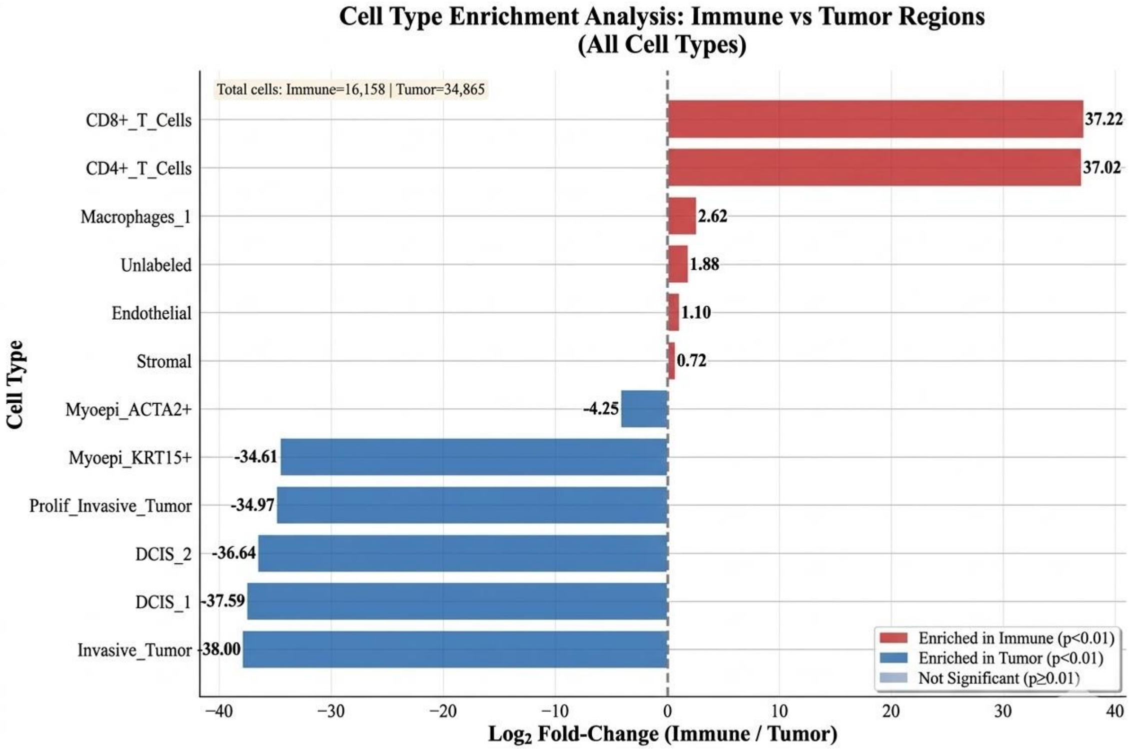

Figure S44: **Cell-type enrichment distinguishes immune and tumour regions in the reconstructed spatial atlas.** Bars show the $\log_2$ fold change of cell-type abundance in immune regions relative to tumour regions. Positive values indicate enrichment in immune regions, whereas negative values indicate enrichment in tumour regions. $CD8^+$ T cells and $CD4^+$ T cells were strongly enriched in immune regions, while epithelial tumour states, including Invasive_Tumor, DCIS_1, DCIS_2 and Prolif_Invasive_Tumor, were preferentially enriched in tumour regions. Colours denote enrichment direction and statistical significance.

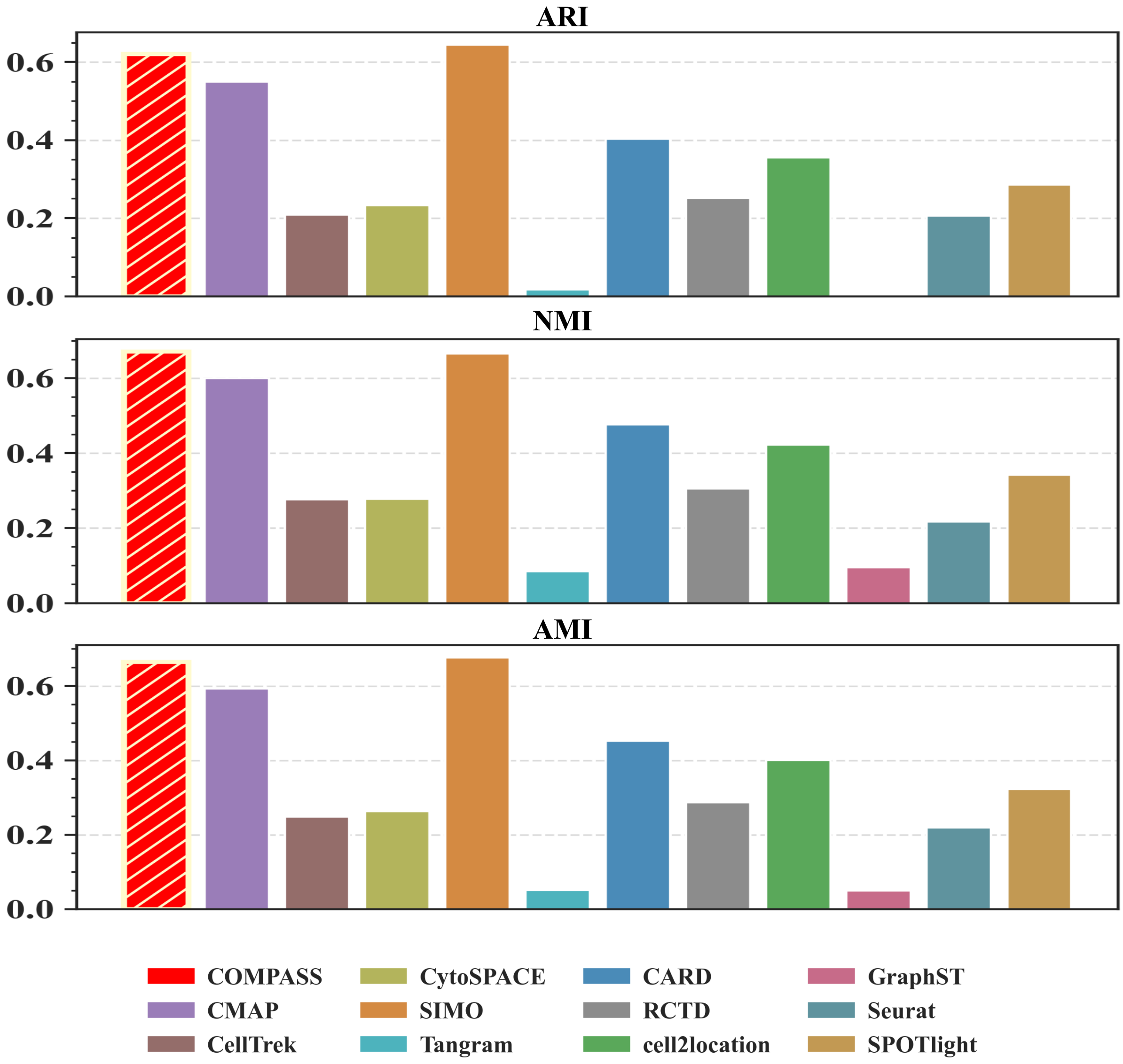

Figure S45: **External clustering agreement on the mouse olfactory bulb dataset.** Bar plots compare COMPASS and benchmark methods using ARI, NMI, and AMI between inferred spatial partitions and the laminar reference annotations of the mouse olfactory bulb. Higher values indicate stronger agreement with the known tissue architecture.

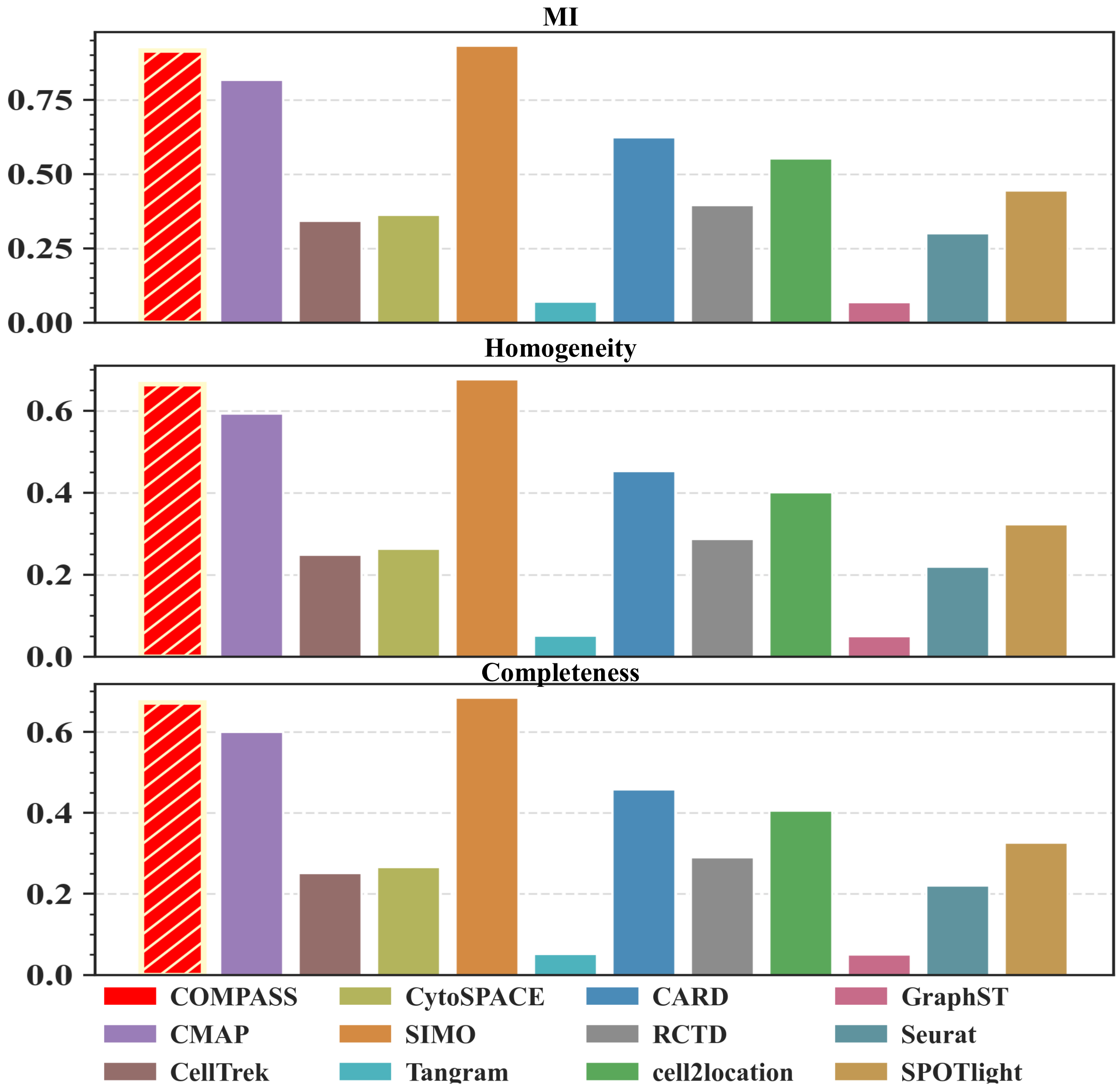

Figure S46: **Cluster information and purity on the mouse olfactory bulb dataset.** Bar plots compare COMPASS and benchmark methods using MI, homogeneity, and completeness between inferred spatial partitions and ground-truth laminar annotations. Higher values indicate better recovery of layer-specific spatial organization.

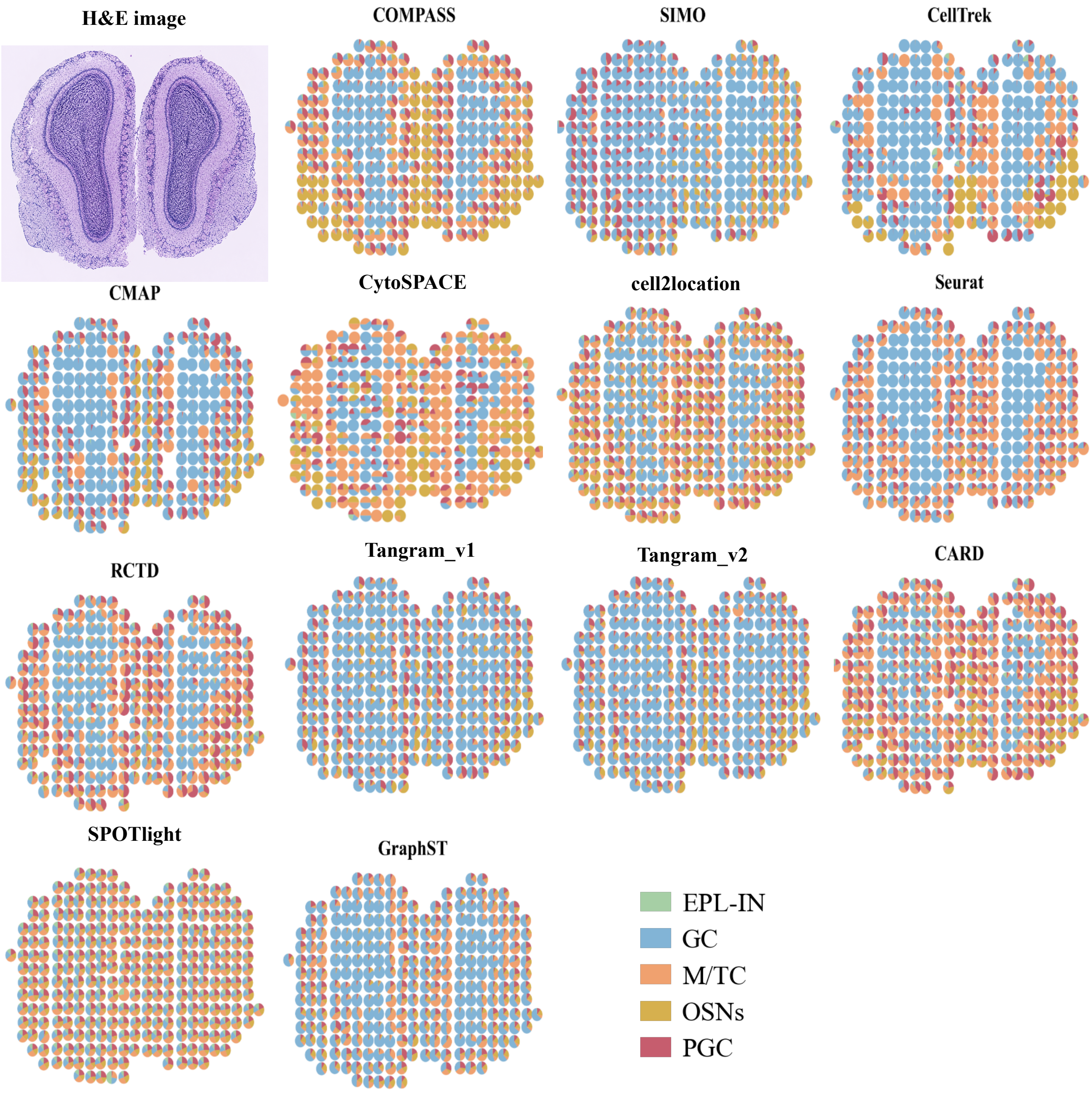

Figure S47: **Spatial cell-type composition in the mouse olfactory bulb.** The H&E image provides the anatomical reference for the mouse olfactory bulb (MOB) section profiled by Spatial Transcriptomics v1.0. Spot-level pie charts compare COMPASS with baseline mapping and deconvolution methods, showing the inferred proportions of five major MOB cell populations: granule cells (GC), olfactory sensory neurons (OSNs), periglomerular cells (PGC), mitral and tufted cells (M-TC), and external plexiform layer interneurons (EPL-IN). Each pie represents one spatial spot, with sector area proportional to the estimated cell-type fraction. COMPASS recovers an anatomically coherent laminar organization, with a GC-dominant central compartment and peripheral enrichment of olfactory bulb cell populations, whereas competing methods show stronger cross-layer mixing or incomplete spatial structure.

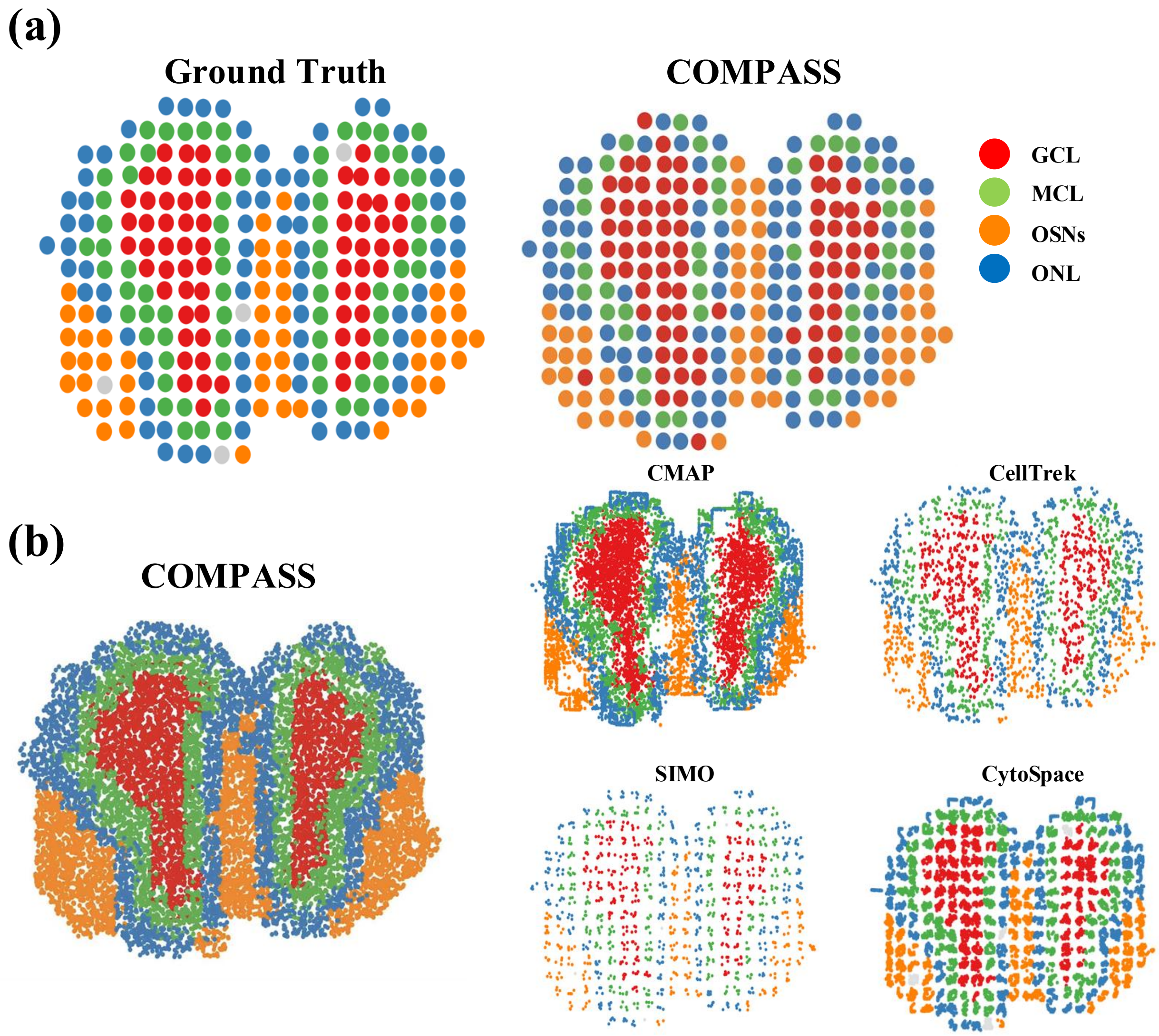

Figure S48: **Spatial domain reconstruction in the mouse olfactory bulb.** (a) Spatial maps colored by the four laminar layers of the MOB, namely the granule cell layer (GCL), mitral cell layer (MCL), olfactory sensory neurons (OSNs), and olfactory nerve layer (ONL), comparing the ground-truth laminar annotation with the reconstructions of COMPASS and four mapping baselines (CMAP, CellTrek, SIMO, and CytoSpace). (b) Single-cell map reconstructed by COMPASS, with cells colored by the four inferred spatial domains (clusters 1 to 4), which correspond to the GCL, MCL, OSNs, and ONL layers in (a). The spot-level cluster view in (a) and the cell-level mapped-cell view in (b) produced by COMPASS remain spatially continuous and highly concordant, recovering a concentric laminar organization with a GCL-dominant core and an ONL-enriched periphery, whereas competing methods show fragmented boundaries, incomplete spatial coverage, or grid-like artifacts.

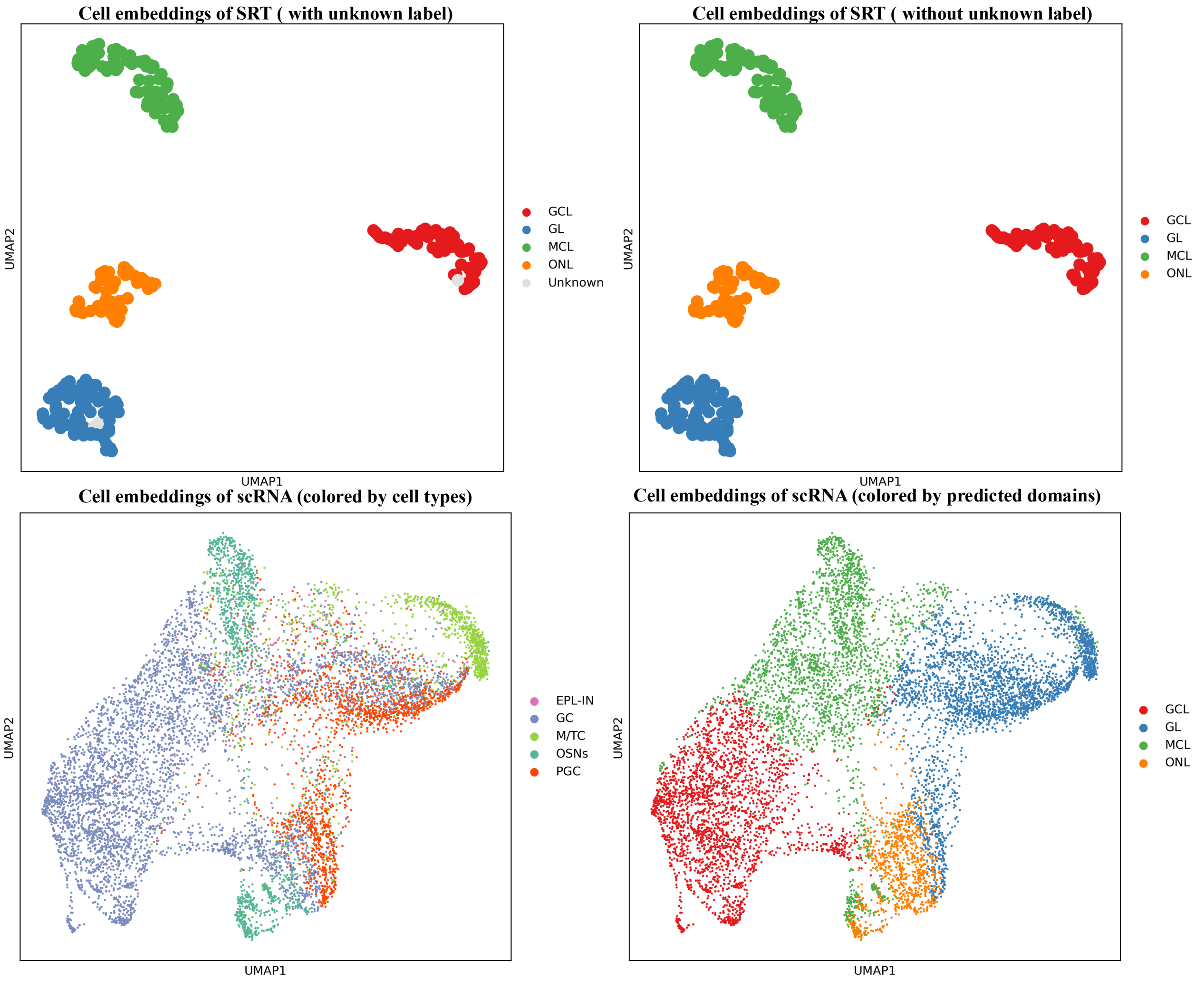

Figure S49: **Embedding structure of spatial transcriptomics and scRNA-seq data in the mouse olfactory bulb.** UMAP visualizations show spatial transcriptomics (SRT) embeddings with and without the unknown label, and scRNA-seq embeddings colored by cell types or COMPASS-predicted spatial domains. The plots illustrate the separation of laminar SRT domains and the transfer of domain structure to dissociated single cells.

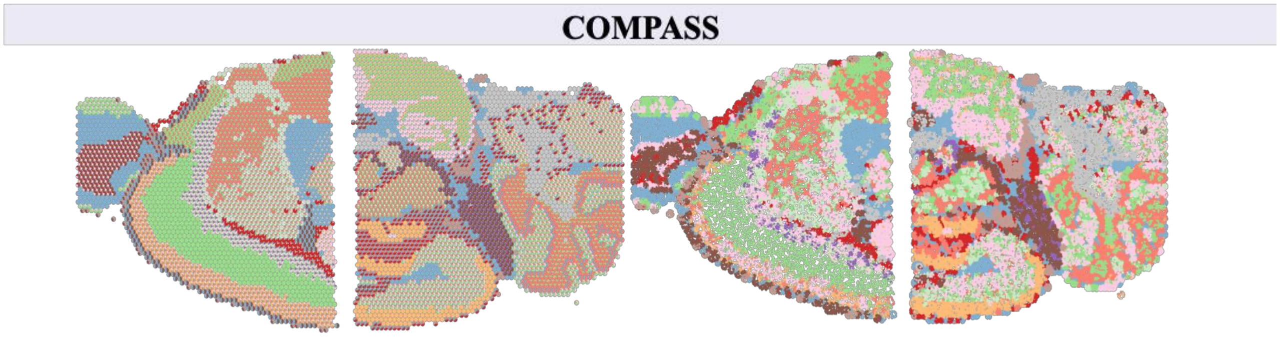

Figure S50: **Spatial reconstruction of mouse brain sections with section-wise domain inference.** COMPASS results are shown for the two sagittal mouse brain sections after separate domain clustering. The left panel shows spot-level cell-type composition pie charts, and the right panel shows the corresponding spatial spot map, demonstrating preservation of laminar and regional organization across sections.

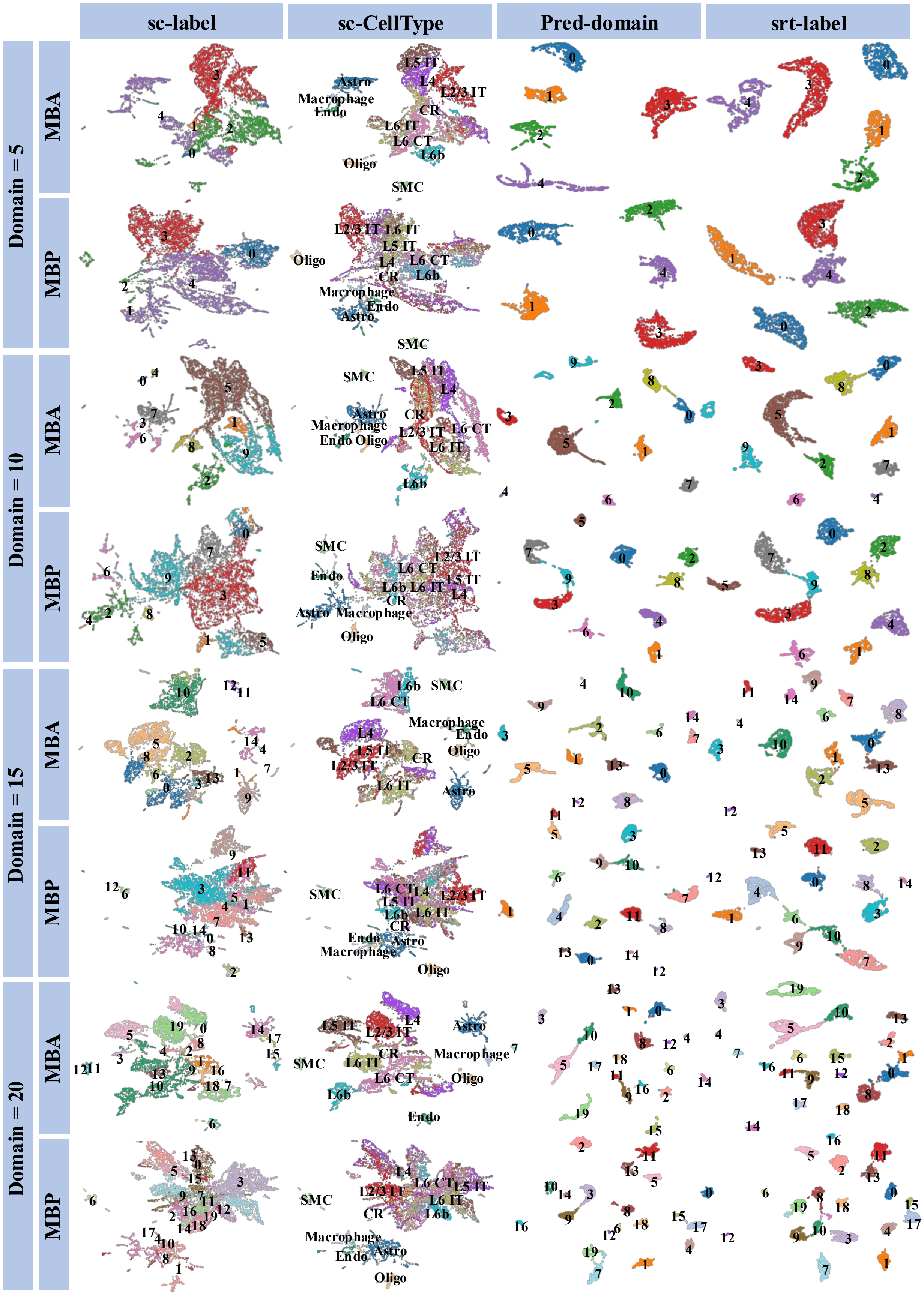
sc-label
sc-CellType
Pred-domain
srt-label
Domain = 5
Domain = 10
Domain = 15
Domain = 20
MBA
MBP

Figure S51: **Joint UMAP analysis of two sagittal mouse brain sections under different domain numbers.** UMAP visualizations show the shared RNA latent space when the two mouse brain sections, denoted MBA and MBP, were clustered jointly with domain numbers of 5, 10, 15, and 20. Columns show scRNA-seq labels, scRNA-seq cell-type annotations, predicted domains for dissociated cells, and spatial transcriptomics (SRT) domain labels, illustrating cross-section consistency of COMPASS domain inference.

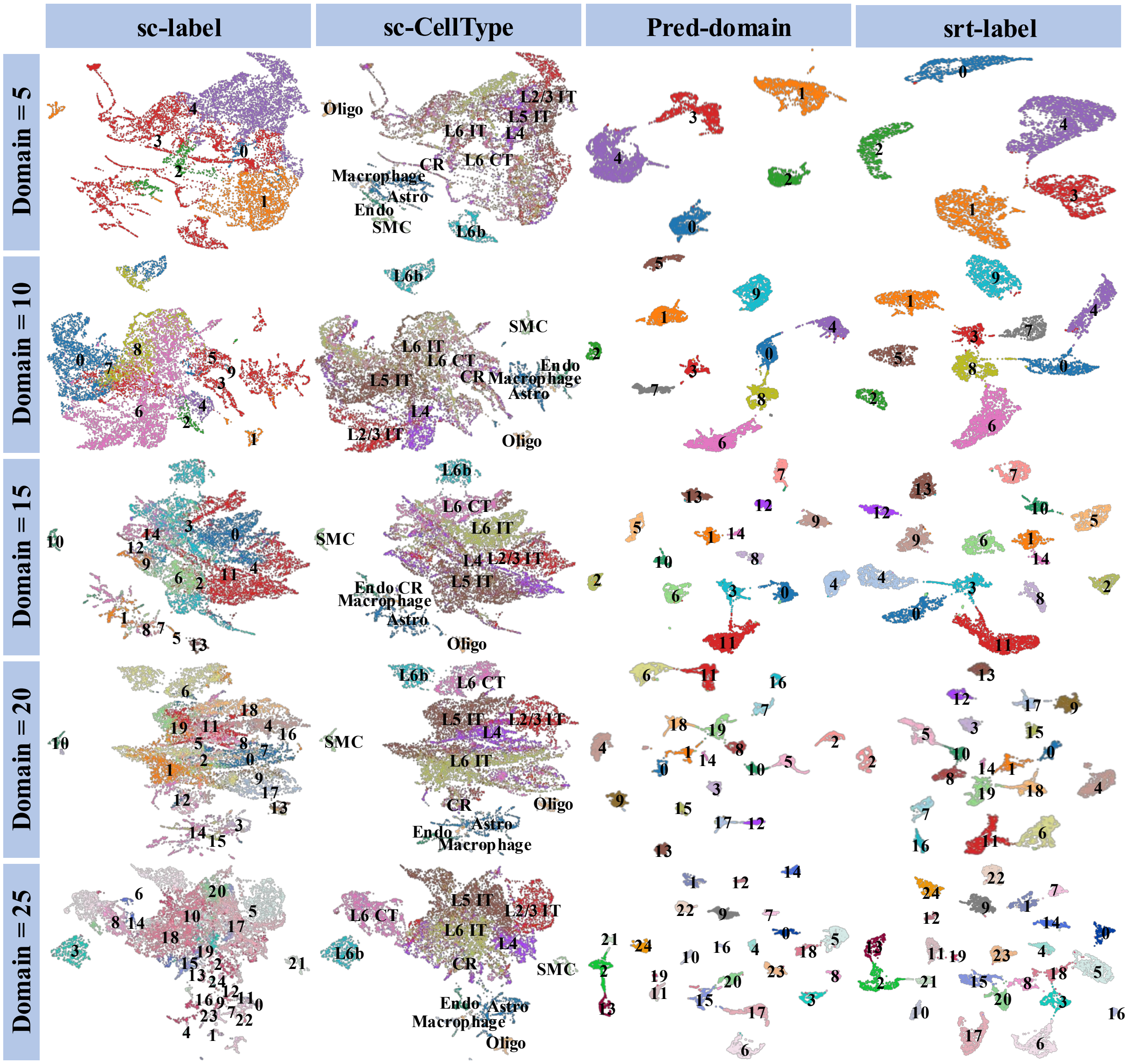

Figure S52: **Section-wise UMAP analysis of mouse brain reconstruction under different domain numbers.** UMAP visualizations show COMPASS embeddings when spatial domains were inferred separately across domain numbers of 5, 10, 15, 20, and 25. Columns show scRNA-seq labels, scRNA-seq cell-type annotations, predicted domains, and SRT domain labels, assessing the stability of domain structure and cell-type organization across clustering resolutions.

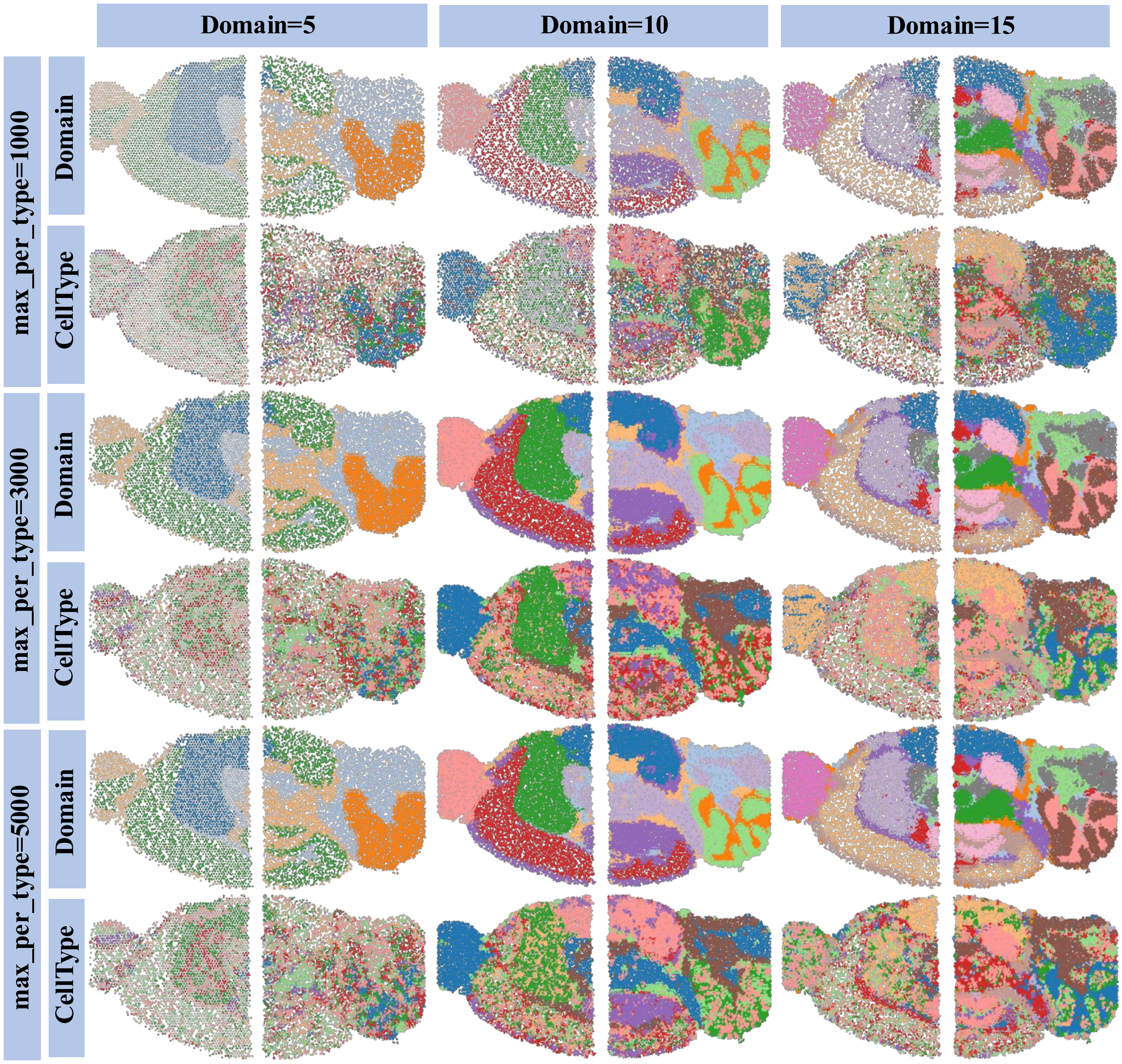

Figure S53: **Sensitivity of mouse brain spatial mapping to domain number and per-type sampling capacity.** Spatial maps compare COMPASS reconstructions across domain numbers of 5, 10, and 15 and maximum cells per type (`max_per_type`) of 1000, 3000, and 5000. For each setting, maps are colored by inferred spatial domain and mapped cell type, showing that COMPASS preserves major anatomical structures across domain-resolution and sampling-capacity choices.

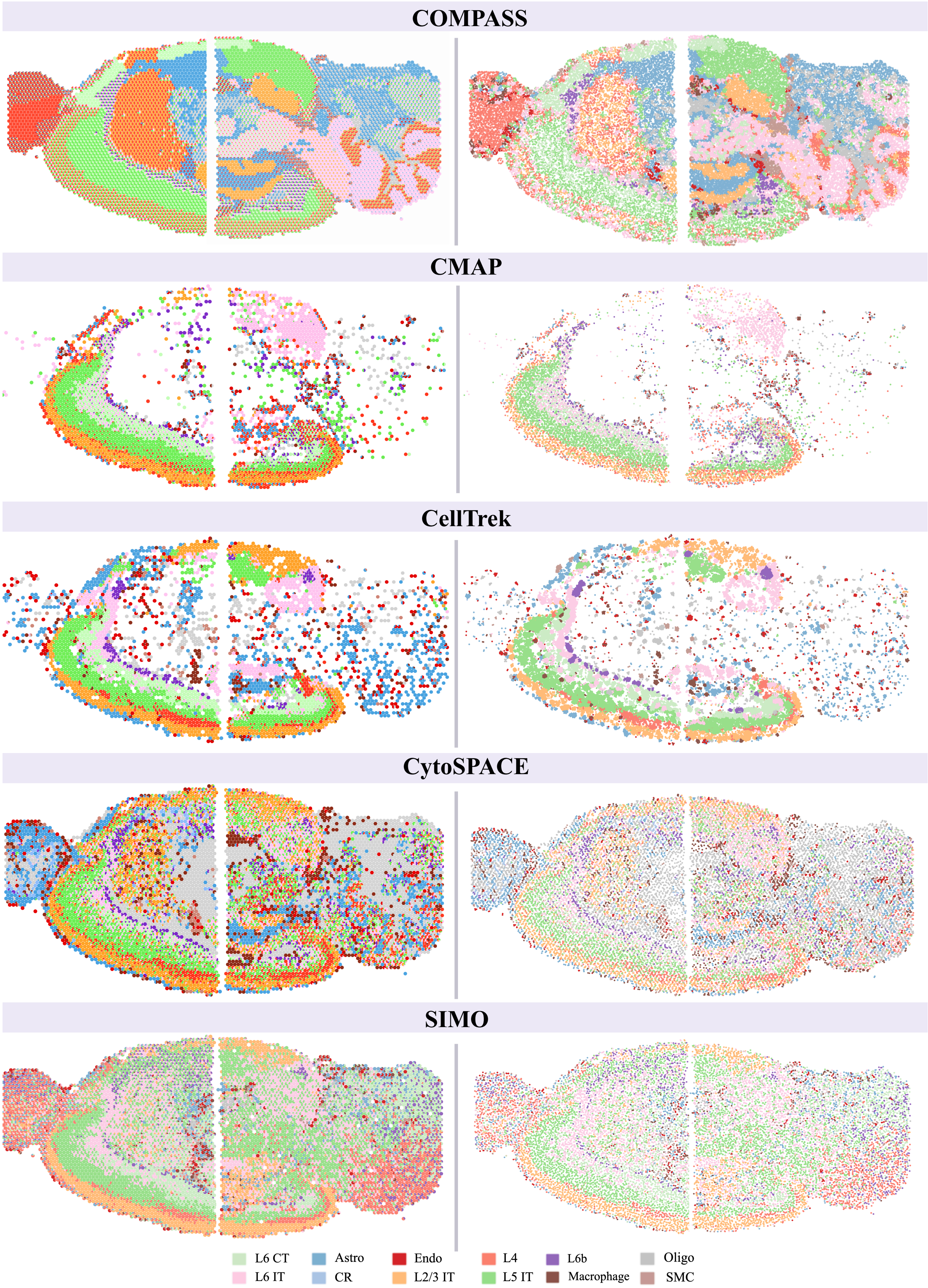
COMPASS
CMAP
CellTrek
CytoSPACE
SIMO
L6 CT
L6 IT
Astro
CR
Endo
L2/3 IT
L4
L5 IT
L6b
Macrophage
Oligo
SMC

Figure S54: **COMPASS preserves coherent cell-type organization across two sagittal mouse brain sections.** Representative single-cell spatial maps reconstructed from two 10x Genomics Visium sagittal mouse brain sections are shown for COMPASS and four mapping baselines, including CMAP, CellTrek, CytoSPACE, and SIMO. Each point represents one mapped cell and is colored by the annotated cell type shown in the legend. COMPASS jointly reconstructs the two sections with dense tissue coverage, clear cortical laminar organization, and consistent localization of neuronal and glial populations, including L2/3 IT, L5 IT, L6 CT, astrocytes, and oligodendrocytes. In contrast, baseline methods show distinct failure modes, including cell dropout and fragmented layouts in CMAP and CellTrek, disrupted or mixed laminar patterns in CytoSPACE, and cross-region mixing with weakened anatomical boundaries in SIMO.

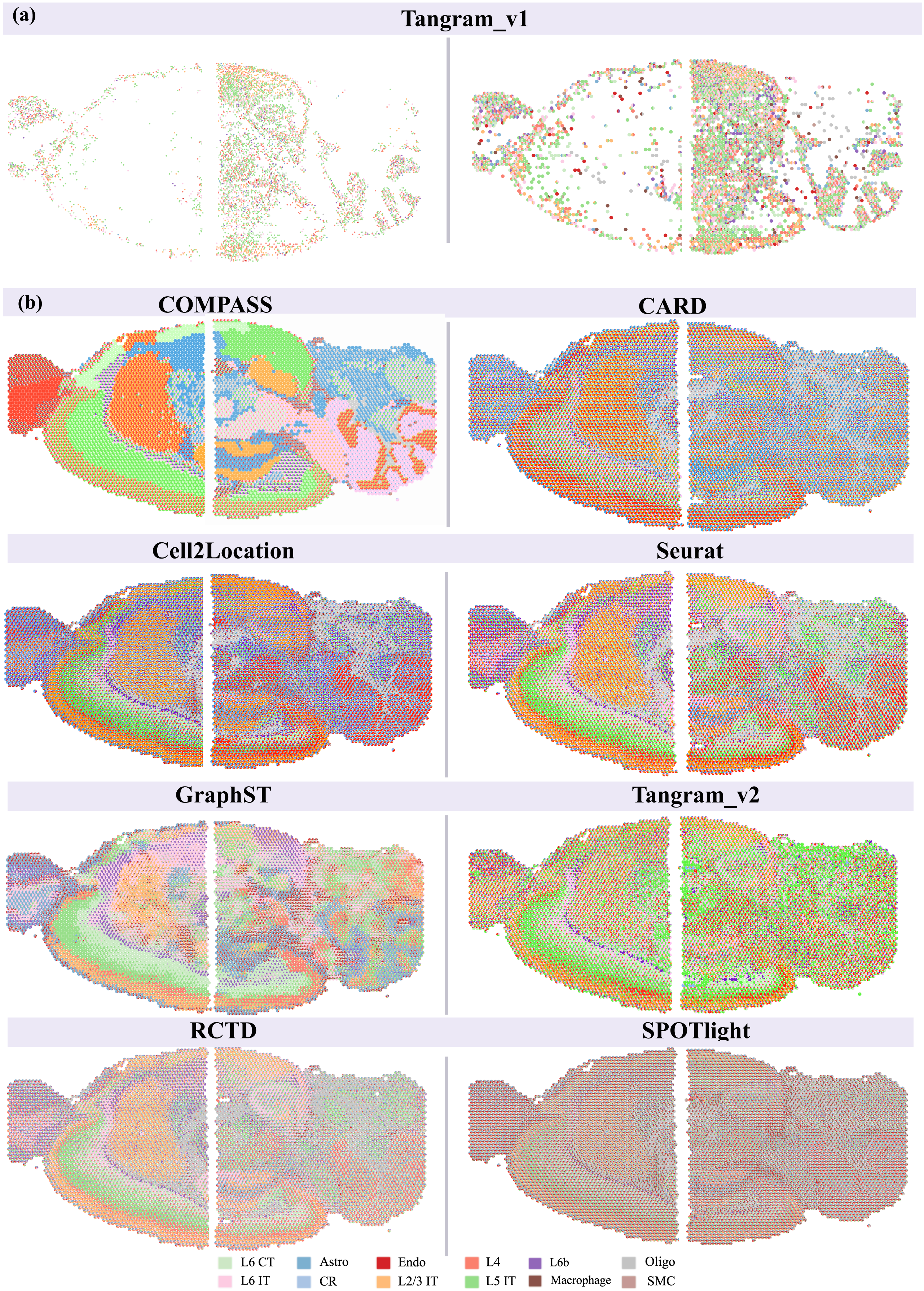
(a)
Tangram_v1
(b)
COMPASS
CARD
Cell2Location
Seurat
GraphST
Tangram_v2
RCTD
SPOTlight
L6 CT
Astro
Endo
L4
L6b
Oligo
L6 IT
CR
L2/3 IT
L5 IT
Macrophage
SMC

Figure S55: **COMPASS preserves spatially coherent cell-type compositions across two sagittal mouse brain sections.** (a) Cell-level mapping results produced by Tangram_v1 across the two Visium sagittal mouse brain sections, showing sparse recovery and substantial cross-region mixing. (b) Spatial cell-type composition maps inferred by COMPASS and benchmark methods, including CARD, Cell2Location, Seurat, GraphST, Tangram_v2, RCTD, and SPOTlight. Each pie chart represents the inferred cell-type composition at one Visium spot, with colors denoting cortical neuronal populations and non-neuronal cell types. Compared with the baseline methods, COMPASS recovers sharper laminar organization across the two sections, preserving localized distributions of major cortical populations, including L2/3 IT, L4, L5 IT, L6 IT, L6 CT, and L6b cells, as well as glial and vascular-associated populations. Baseline methods show varying degrees of over-smoothing, regional mixing, or loss of laminar contrast, consistent with the reduced spatial coherence observed in the sagittal mouse brain benchmark. Cell-type abbreviations are as indicated in the legend.

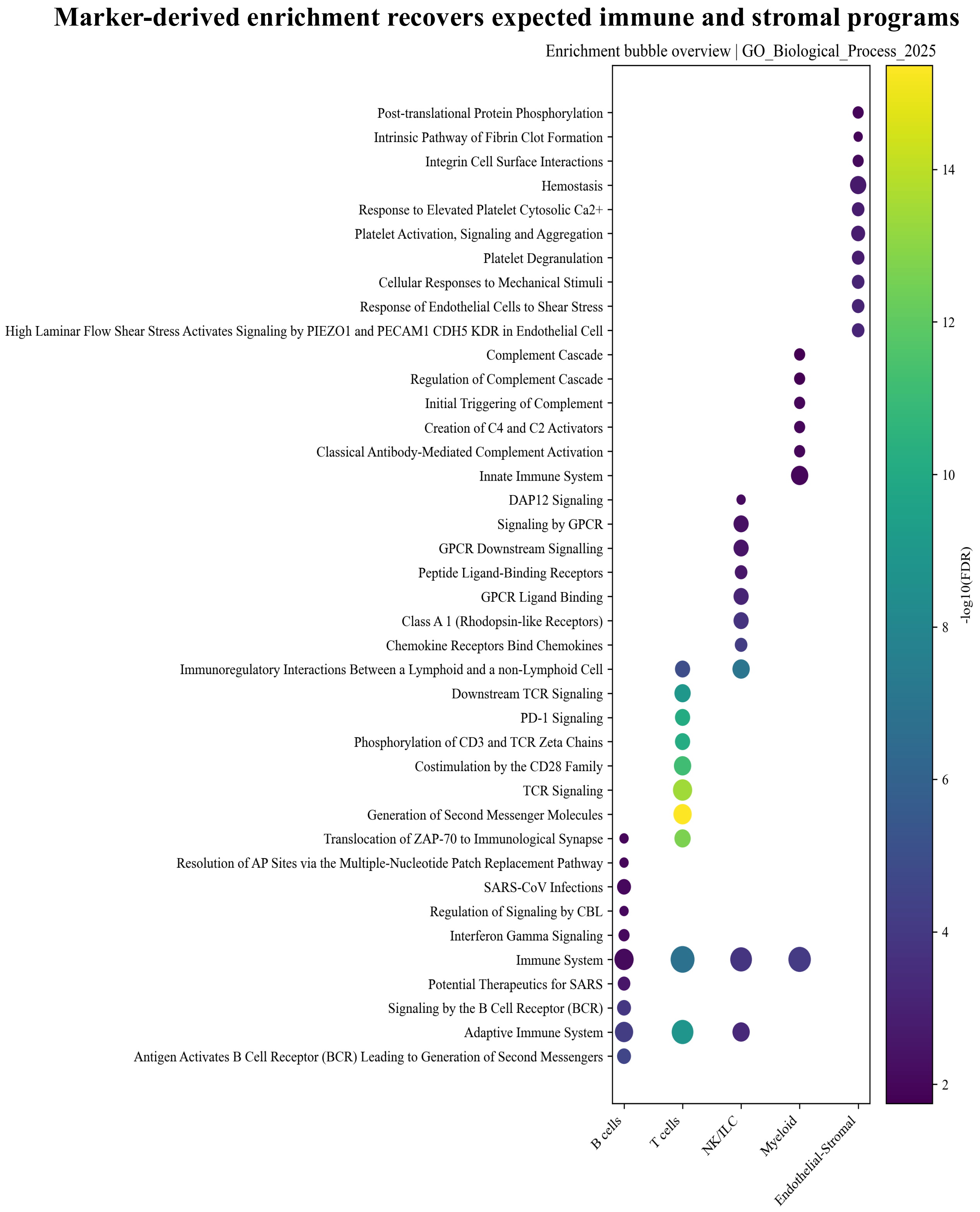

Figure S56: **Marker-derived functional enrichment of coarse lymph-node cell classes.** Bubble plot showing gene ontology biological process enrichment for marker genes associated with B cells, T cells, NK/ILC cells, myeloid cells, and endothelial–stromal cells. Dot size represents enrichment strength or gene-set overlap, and color denotes statistical significance measured by $-\log_{10}(\text{FDR})$. Enriched terms recover expected immune and stromal programs, including B-cell receptor signaling in B cells, T-cell receptor and CD3-mediated signaling in T cells, innate immune and chemokine-related programs in NK/ILC and myeloid populations, and platelet, complement, and endothelial stress-response programs in endothelial-stromal cells.

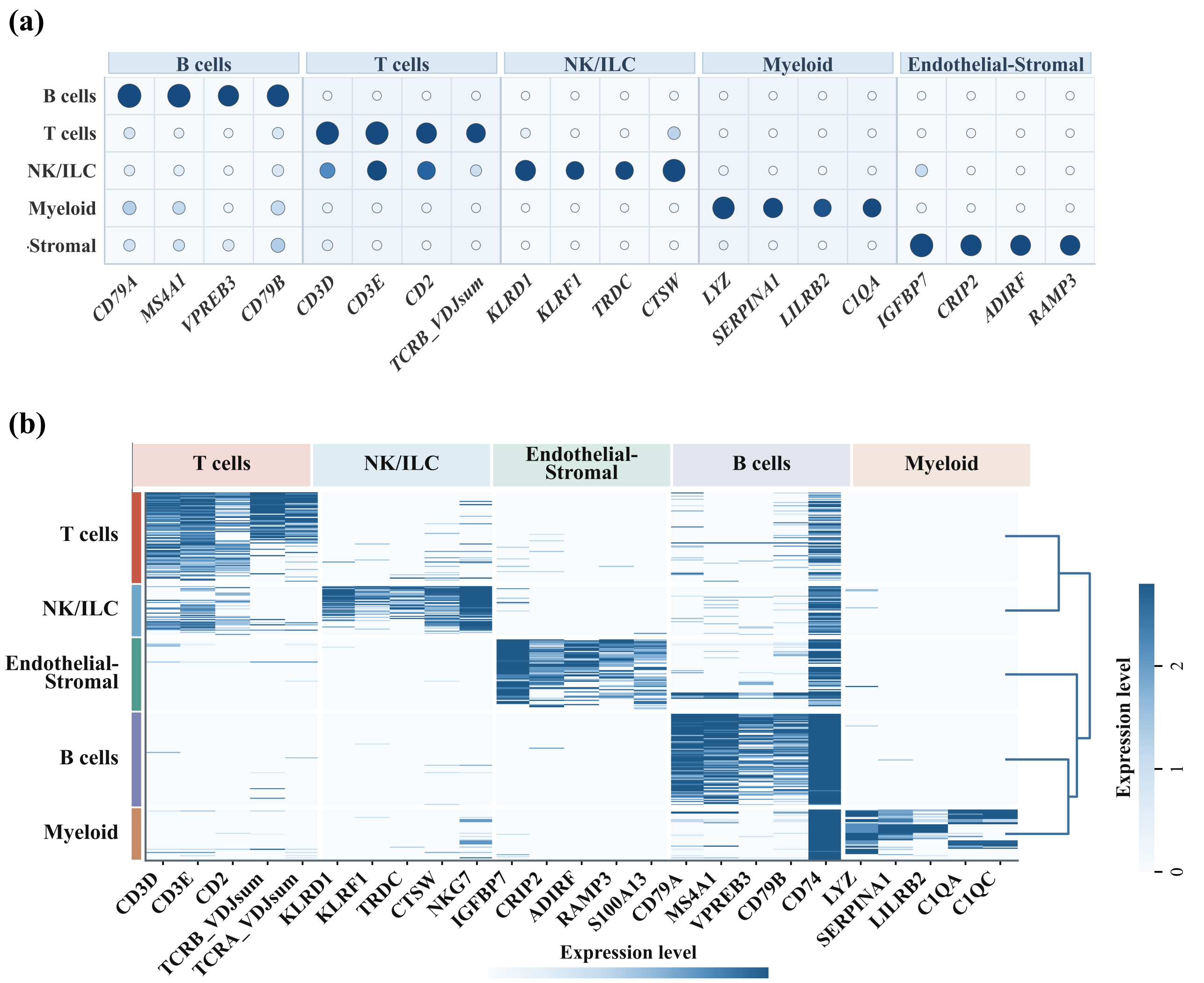

Figure S57: **Canonical marker genes distinguish major cell classes in the human lymph node dataset.** (a) Dot plot showing the expression of canonical marker genes across five coarse cell classes. B-cell markers include CD79A, MS4A1, VPREB3, and CD79B; T-cell markers include CD3D, CD3E, CD2, and TCR-related genes; NK/ILC markers include KLRD1, KLRF1, TRDC, CTSW, and NKG7; myeloid markers include LYZ, SERPINA1, LILRB2, and C1QA; endothelial–stromal markers include IGFBP7, CRIP2, ADIRF, and RAMP3. (b) Heatmap of marker-gene expression across reconstructed cells, showing that cells grouped by COMPASS exhibit lineage-specific marker programs consistent with their annotated coarse cell classes.

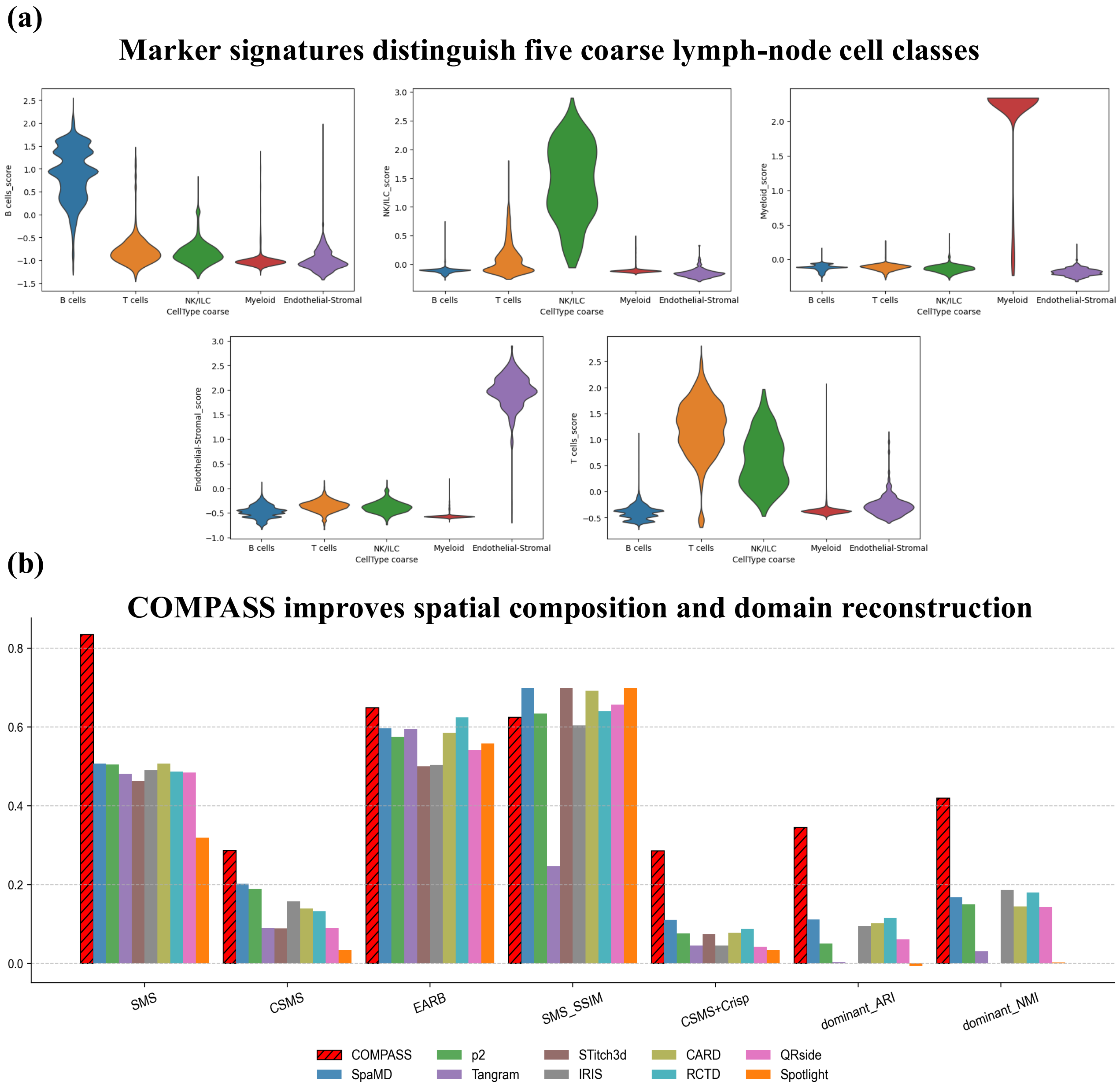

Figure S58: **Marker-based cell-class validation and spatial reconstruction metrics in the human lymph node dataset.** (a) Violin plots show representative marker-gene signatures for five coarse lymph-node cell classes, including B cells, T cells, NK/ILC cells, myeloid cells, and endothelial–stromal cells. Marker scores are selectively enriched in their expected cell classes, supporting the biological validity of the reconstructed cell identities. (b) Bar plots compare COMPASS with benchmark methods using spatial composition and domain-reconstruction metrics, including spatial map similarity (SMS), cosine map similarity (CosMS), edge- or boundary-aware agreement metrics, structural similarity of spatial maps, composite spatial-map scores, and dominant-cell-type ARI and NMI against anatomical annotations. Higher values indicate better recovery of spatial composition and tissue-domain organization.

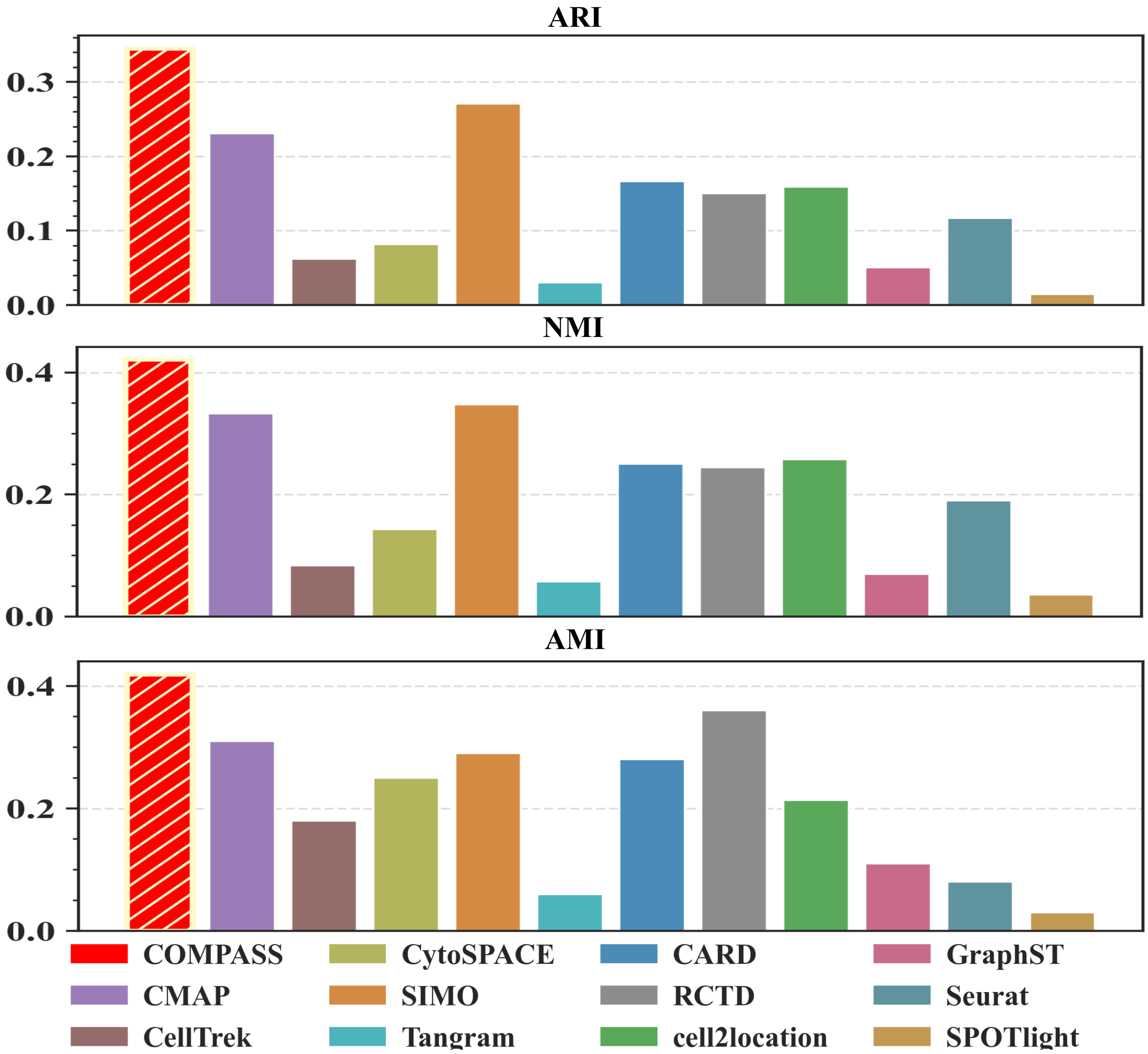

Figure S59: **External clustering agreement with anatomical regions in the human lymph node dataset.** Bar plots compare COMPASS and benchmark methods using ARI, NMI, and AMI between inferred spatial partitions and H&E-based anatomical annotations. These metrics evaluate whether the reconstructed spatial domains recover the major lymph-node structures, including cortical, follicular, medullary, hilar, and stromal regions. Higher values indicate better agreement with the annotated tissue architecture.

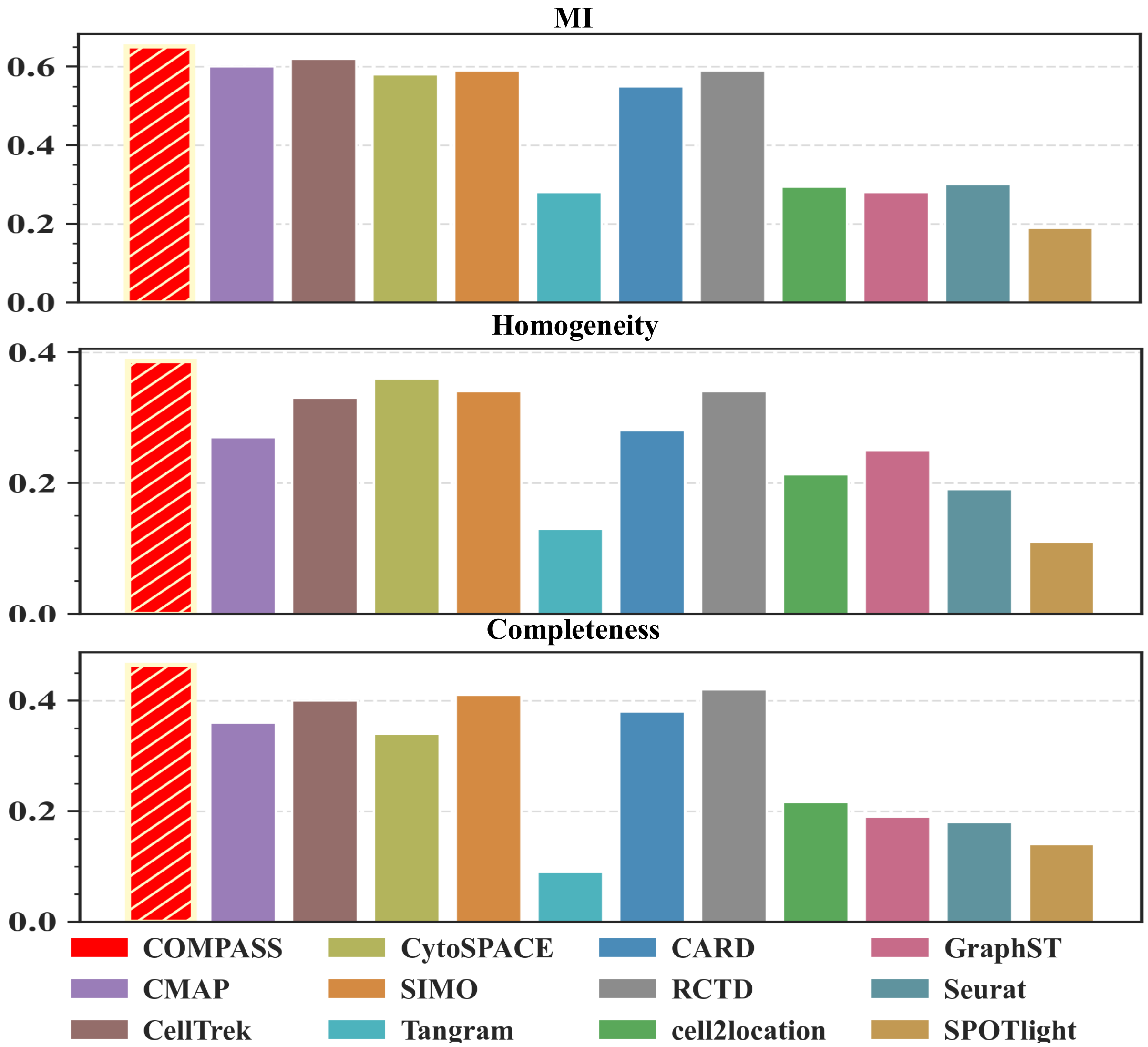

Figure S60: **Cluster information and purity metrics in the human lymph node dataset.** Bar plots compare COMPASS and benchmark methods using MI, homogeneity, and completeness between inferred spatial partitions and H&E-annotated anatomical regions. MI measures the shared information between inferred and reference partitions, homogeneity evaluates whether each inferred cluster is dominated by a single anatomical region, and completeness evaluates whether each anatomical region is assigned consistently to one inferred partition. Higher values indicate stronger concordance with lymph-node tissue organization.

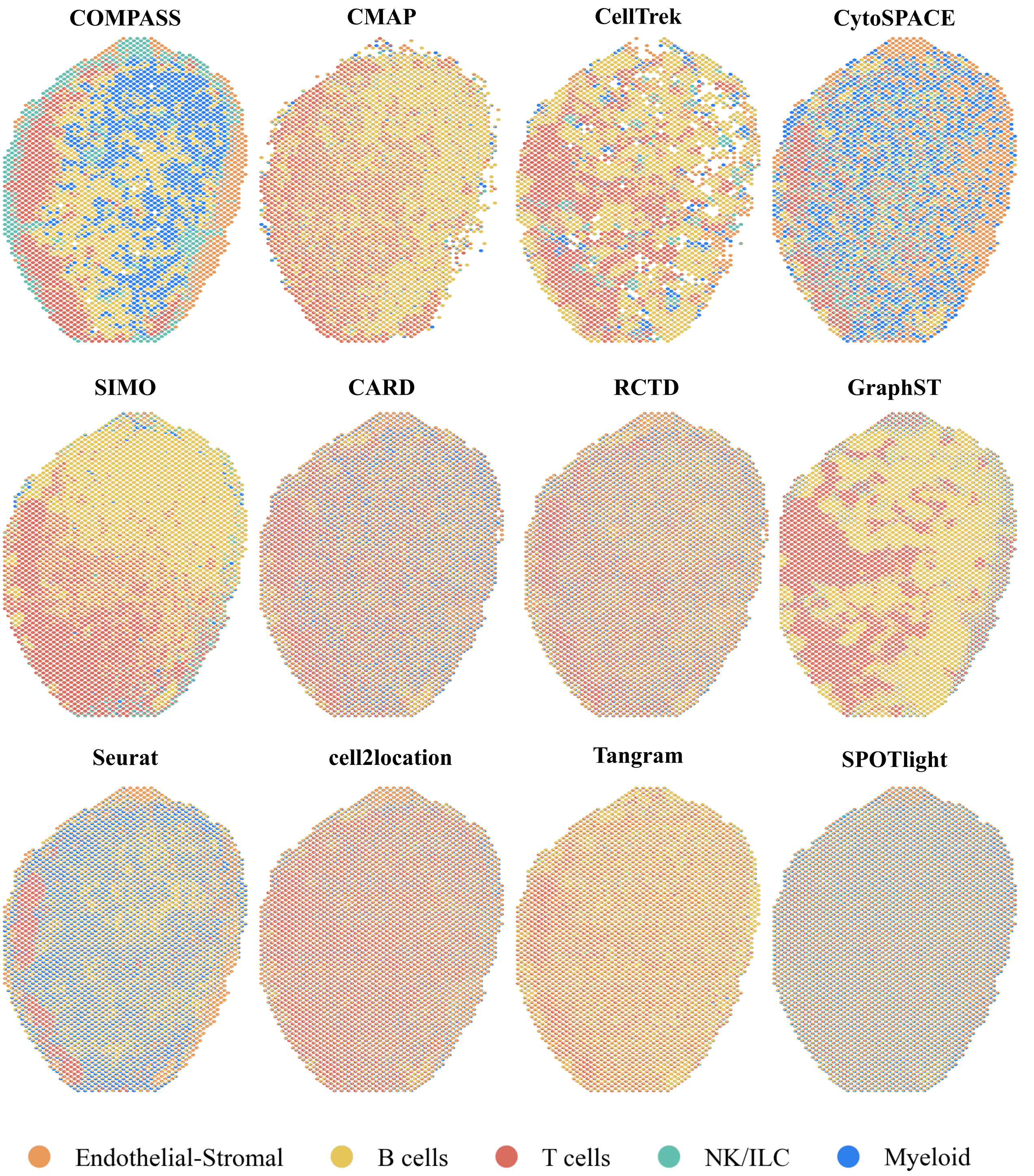

Figure S61: **Spatial cell-type composition in the human lymph node.** Spot-level pie charts show the inferred proportions of five major cell populations in the 10x Genomics Visium CytAssist human lymph node dataset: endothelial-stromal cells, B cells, T cells, NK/ILC cells, and myeloid cells. Each pie represents one spatial spot, with sector area proportional to the estimated cell-type fraction. COMPASS integrates paired spatial RNA and ADT protein measurements, whereas baseline methods rely on transcriptomic information. COMPASS recovers the layered lymph-node architecture, including B-cell enrichment in cortical and follicular regions, T-cell enrichment in the paracortical zone, and endothelial-stromal enrichment along the peripheral capsule and hilar regions.

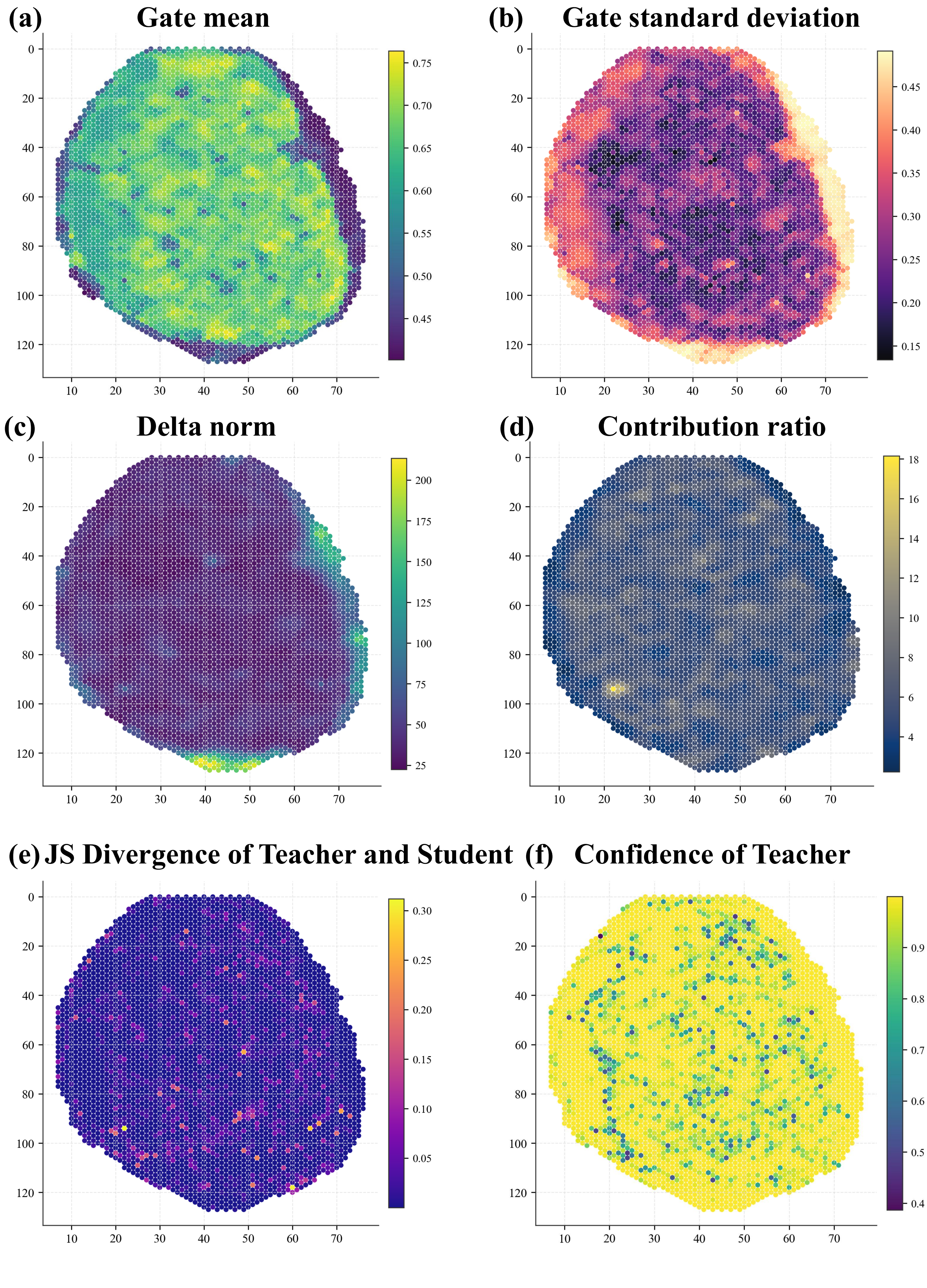
(a) Gate mean
(b) Gate standard deviation
(c) Delta norm
(d) Contribution ratio
(e) JS Divergence of Teacher and Student
(f) Confidence of Teacher

Figure S62: **Spatial distribution of RG-COD diagnostic scores in the human lymph node dataset.** Spatial maps show spot-wise RG-COD diagnostics used to assess how ADT information is incorporated into the RNA-centered representation. (a) Mean gate value across latent dimensions. (b) Standard deviation of gate values across latent dimensions. (c) Norm of the gated residual update. (d) Residual contribution ratio relative to the RNA representation. (e) JS divergence between teacher and student predictions. (f) Teacher confidence, measured as the maximum predicted domain probability of the fused teacher branch. The spatially heterogeneous gate and residual patterns indicate selective auxiliary incorporation, while the low JS divergence suggests that ADT mainly refines the RNA representation rather than inducing large teacher-student decision shifts.

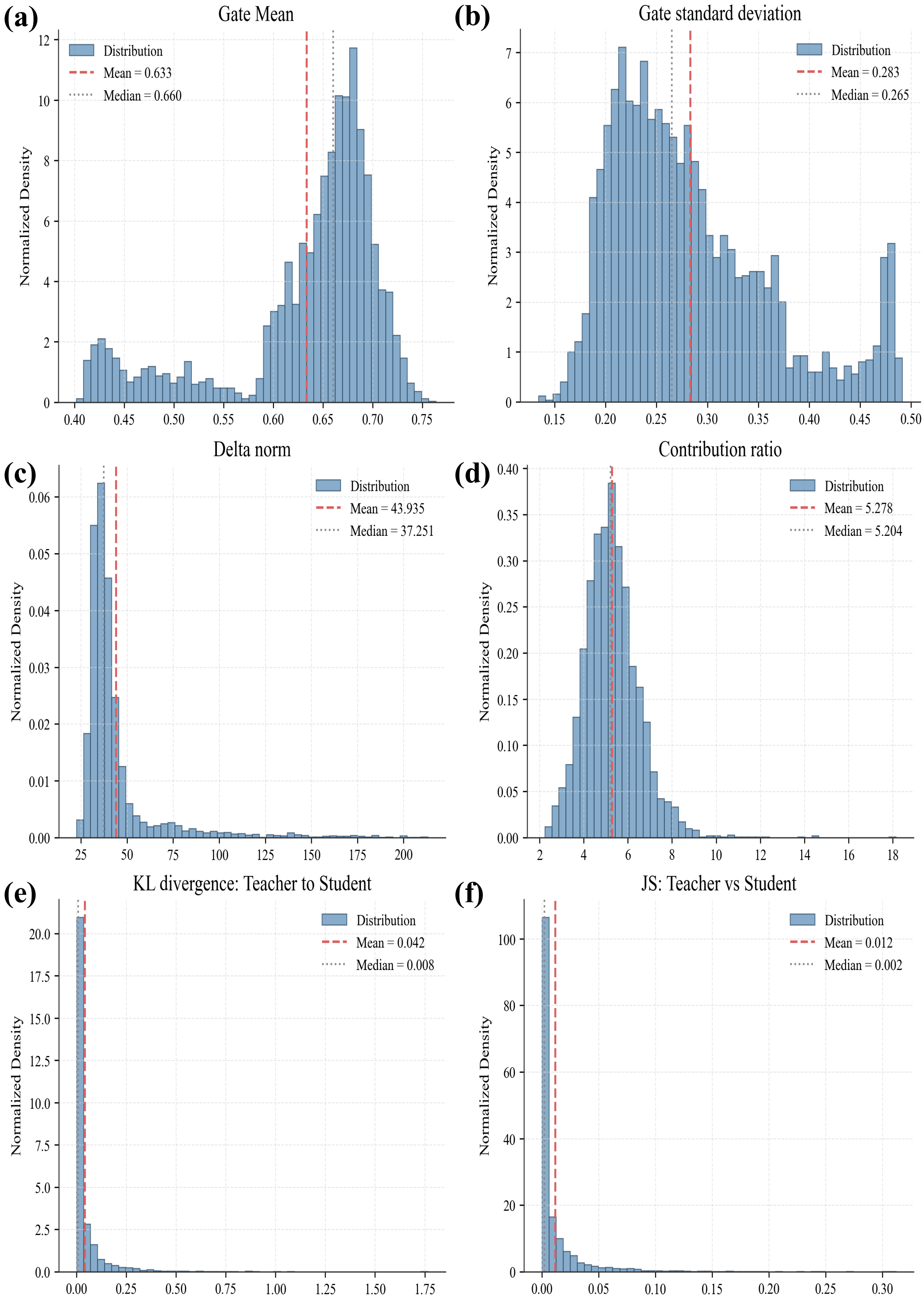
(a)
Gate Mean
Distribution
Mean = 0.633
Median = 0.660
Normalized Density
(b)
Gate standard deviation
Distribution
Mean = 0.283
Median = 0.265
Normalized Density
(c)
Delta norm
Distribution
Mean = 43.935
Median = 37.251
Normalized Density
(d)
Contribution ratio
Distribution
Mean = 5.278
Median = 5.204
Normalized Density
(e)
KL divergence: Teacher to Student
Distribution
Mean = 0.042
Median = 0.008
Normalized Density
(f)
JS: Teacher vs Student
Distribution
Mean = 0.012
Median = 0.002
Normalized Density

Figure S63: **Global distributions of RG-COD diagnostic scores in the human lymph node dataset.** Histograms summarize the spot-wise distributions of RG-COD diagnostics across the tissue section. (a) Gate mean. (b) Gate standard deviation. (c) Delta norm. (d) Residual contribution ratio. (e) KL divergence from teacher to student. (f) JS divergence between teacher and student. Red dashed lines indicate the mean and black dotted lines indicate the median. The elevated gate mean and residual contribution ratio show broad ADT usage, whereas the near-zero KL and JS divergences indicate stable transfer into the RNA-only student.

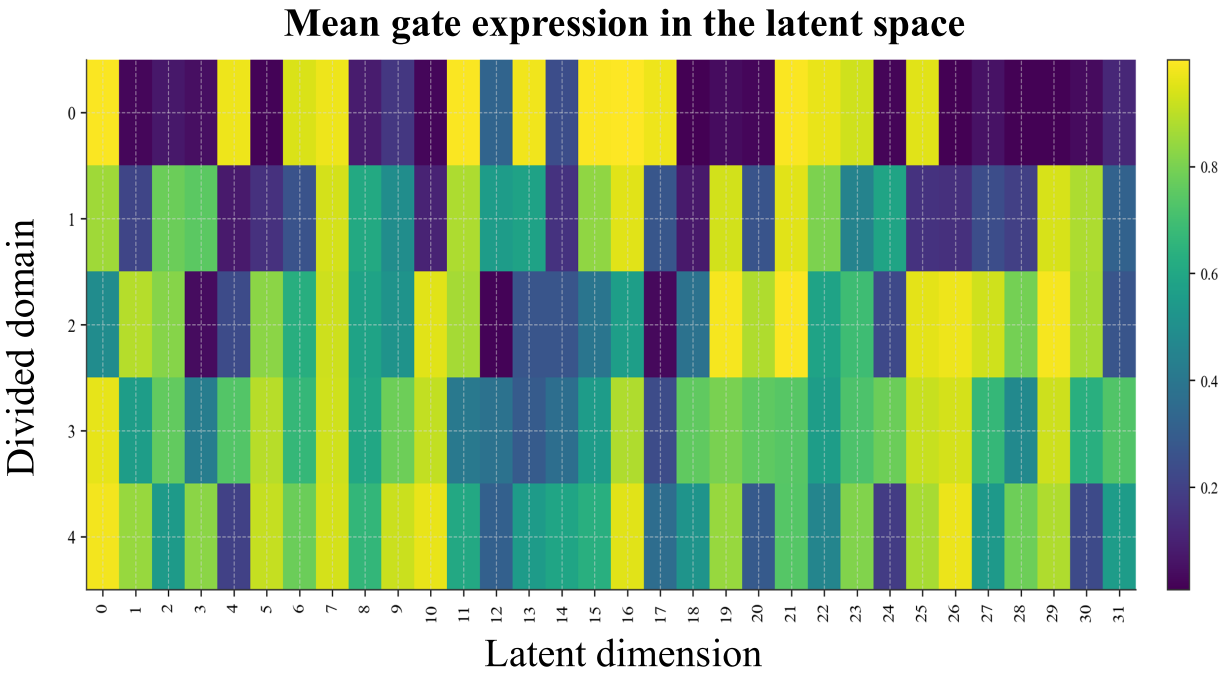

Figure S64: **Domain- and dimension-specific gate activity in the human lymph node dataset.** Heatmap showing the mean RG-COD gate value across inferred spatial domains and latent dimensions. Rows correspond to divided domains, and columns correspond to latent dimensions. The heterogeneous pattern indicates that ADT information is modulated in both a domain-specific and dimension-specific manner, rather than being uniformly fused into the RNA representation.

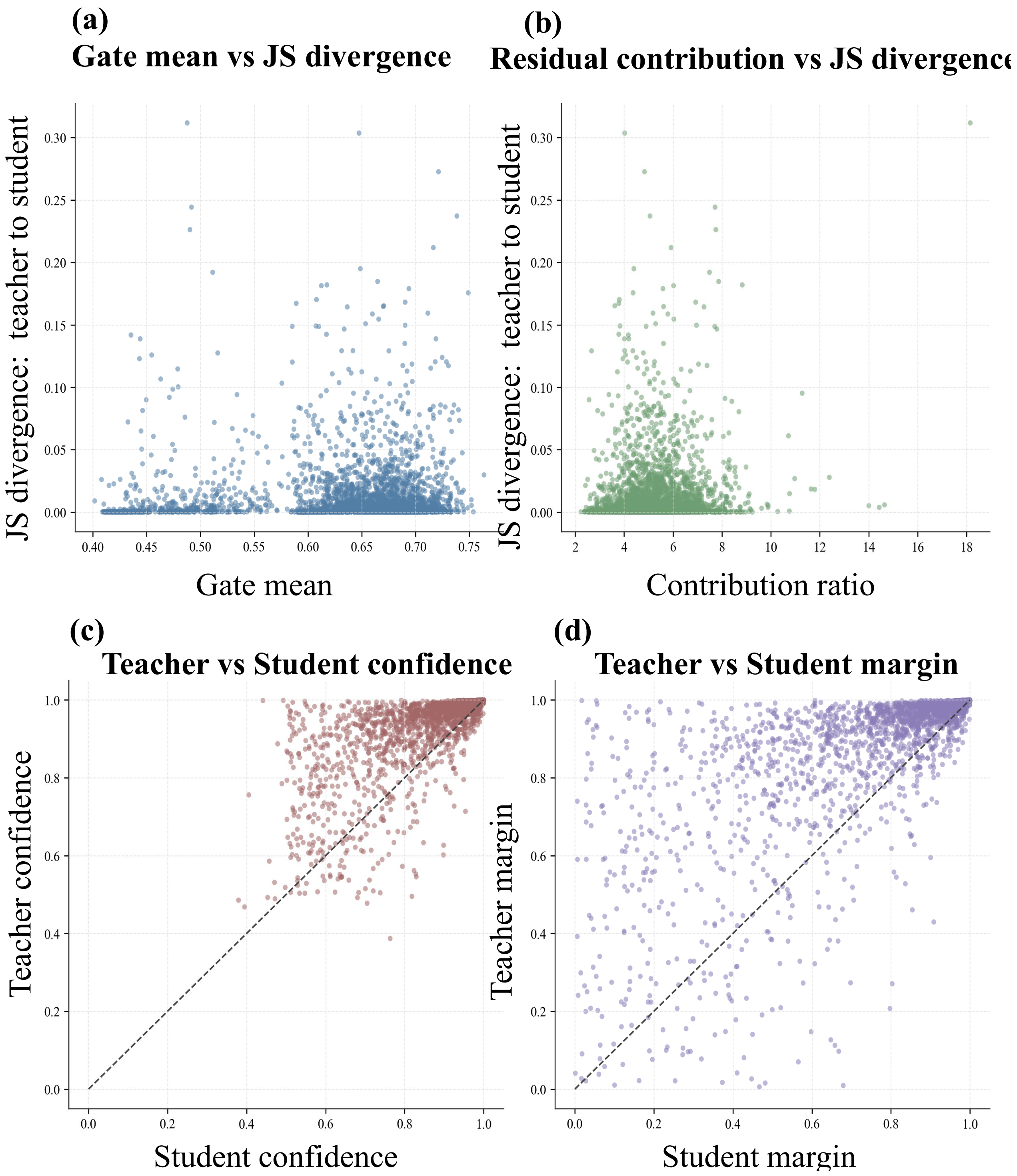

Figure S65: **Relationship between gated ADT transfer and teacher–student agreement in the human lymph node dataset.** Scatter plots examine whether stronger auxiliary incorporation is associated with teacher–student disagreement. (a) Gate mean versus teacher-student JS divergence. (b) Residual contribution ratio versus teacher-student JS divergence. (c) Teacher confidence versus student confidence. (d) Teacher margin versus student margin. Diagonal lines in (c,d) indicate equal confidence or equal margin between teacher and student. These analyses show that RG-COD can introduce substantial residual updates while keeping the student predictions aligned with the fused teacher.

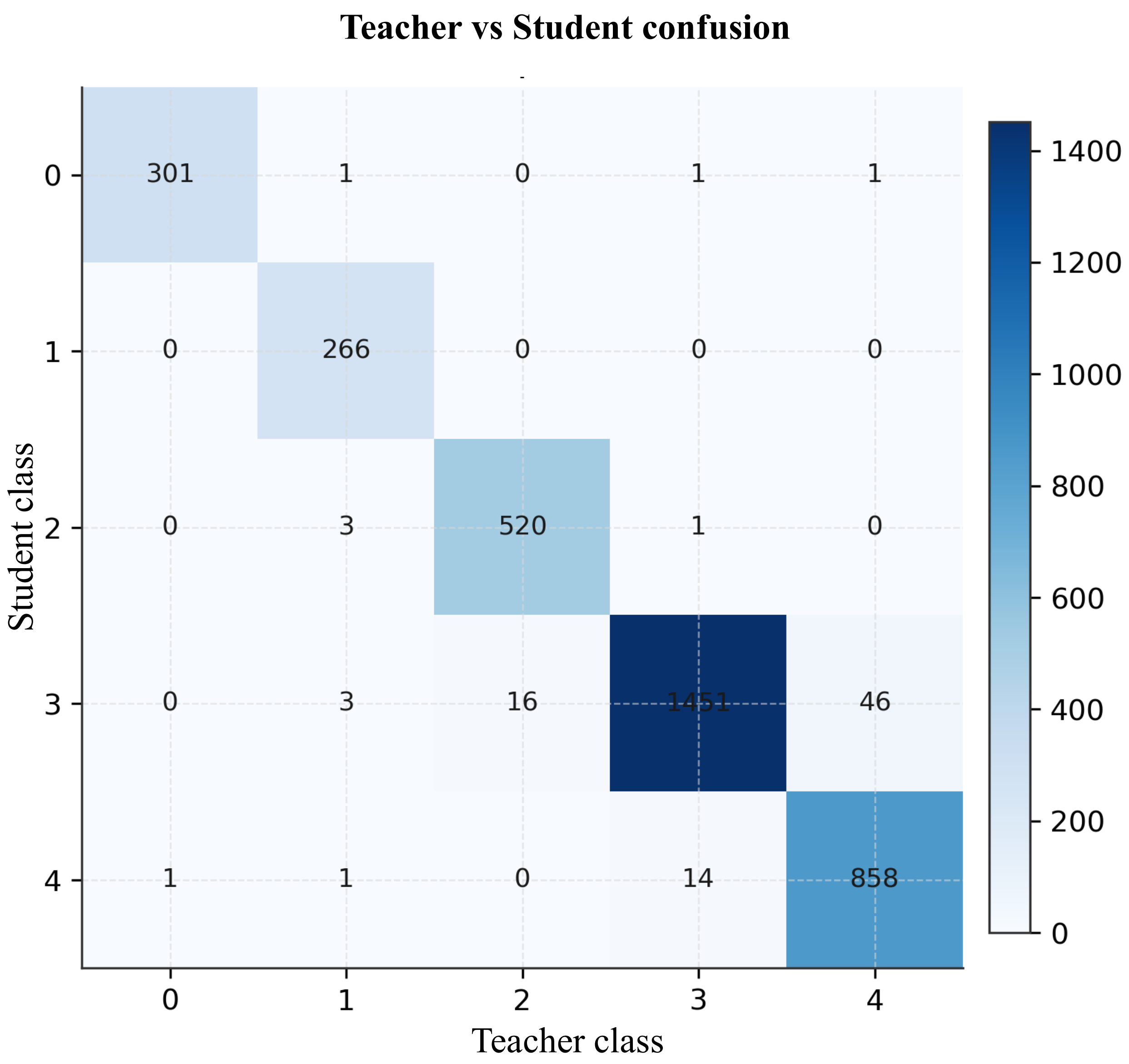

Figure S66: **Teacher-student prediction consistency in the human lymph node dataset.** Confusion matrix comparing domain classes predicted by the fused teacher branch and the RNA-only student after RG-COD training. Columns denote teacher-predicted classes and rows denote student-predicted classes, with color intensity and cell labels indicating the number of spots in each teacher–student class pair. The near-diagonal structure indicates that auxiliary-derived domain information is effectively distilled into the deployable RNA-only student while preserving consistent spatial-domain decisions.

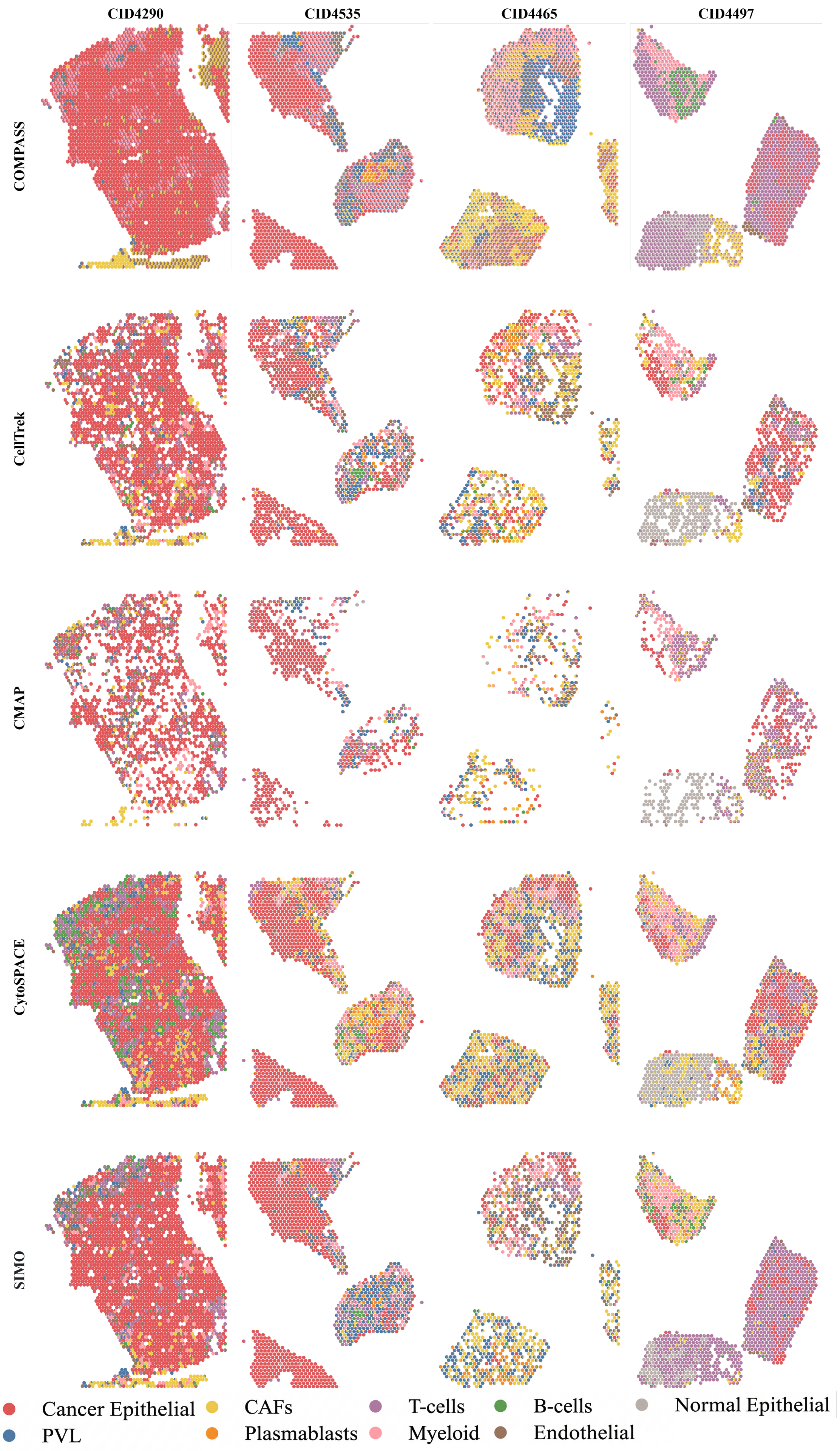
CID4290
CID4535
CID4465
CID4497
COMPASS
CellTrek
CMAP
CytoSPACE
SIMO
Cancer Epithelial
PVL
CAFs
Plasmablasts
T-cells
Myeloid
B-cells
Endothelial
Normal Epithelial

Figure S67: **Benchmark comparison of single-cell spatial reconstruction across clinical breast cancer samples.** Spatial distributions of major cell types reconstructed by COMPASS and four competing mapping methods across four primary breast cancer samples (CID4290, CID4535, CID4465, and CID4497). Columns denote individual samples, and rows denote mapping methods. Each point represents a mapped single cell colored by cell type, including cancer epithelial cells, PVL, CAFs, plasmablasts, T cells, myeloid cells, B cells, endothelial cells, and normal epithelial cells. COMPASS produced spatially complete and biologically coherent cell maps that preserved dense tumor epithelial regions and sample-specific immune and stromal organization, whereas CellTrek, CMAP, CytoSPACE, and SIMO showed varying degrees of fragmented coverage, local mixing, sparse recovery, or over-homogenized regional composition.

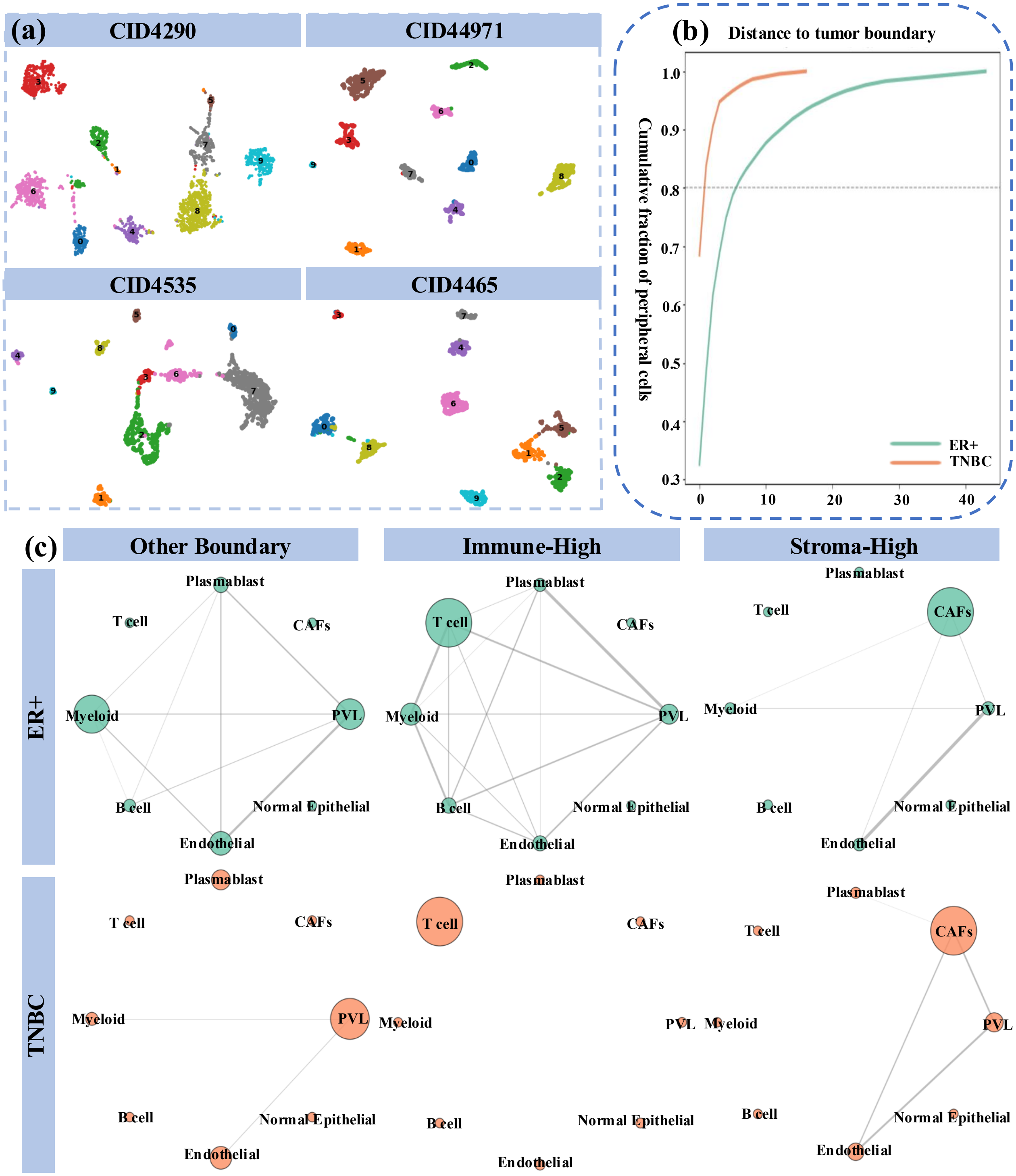

Figure S68: **Subtype-specific spatial organization and ecological interactions in breast cancer.** (a) UMAP visualizations of reconstructed single-cell atlases from four breast cancer samples, including ER+ samples CID4290 and CID4465 and TNBC samples CID44971 and CID4535, colored by major cell type. (b) Cumulative distribution curves of the distance from peripheral cells to the nearest tumor boundary in ER+ and TNBC samples, showing subtype-specific differences in tumor-boundary proximity. (c) Cell–cell interaction networks within three ecological regions, including other boundary, immune-high, and stroma-high regions, stratified by ER+ and TNBC subtypes. Node size represents the relative frequency of each cell type within the corresponding ecological region, and edges indicate significant cell-type associations, with thicker edges denoting stronger associations.

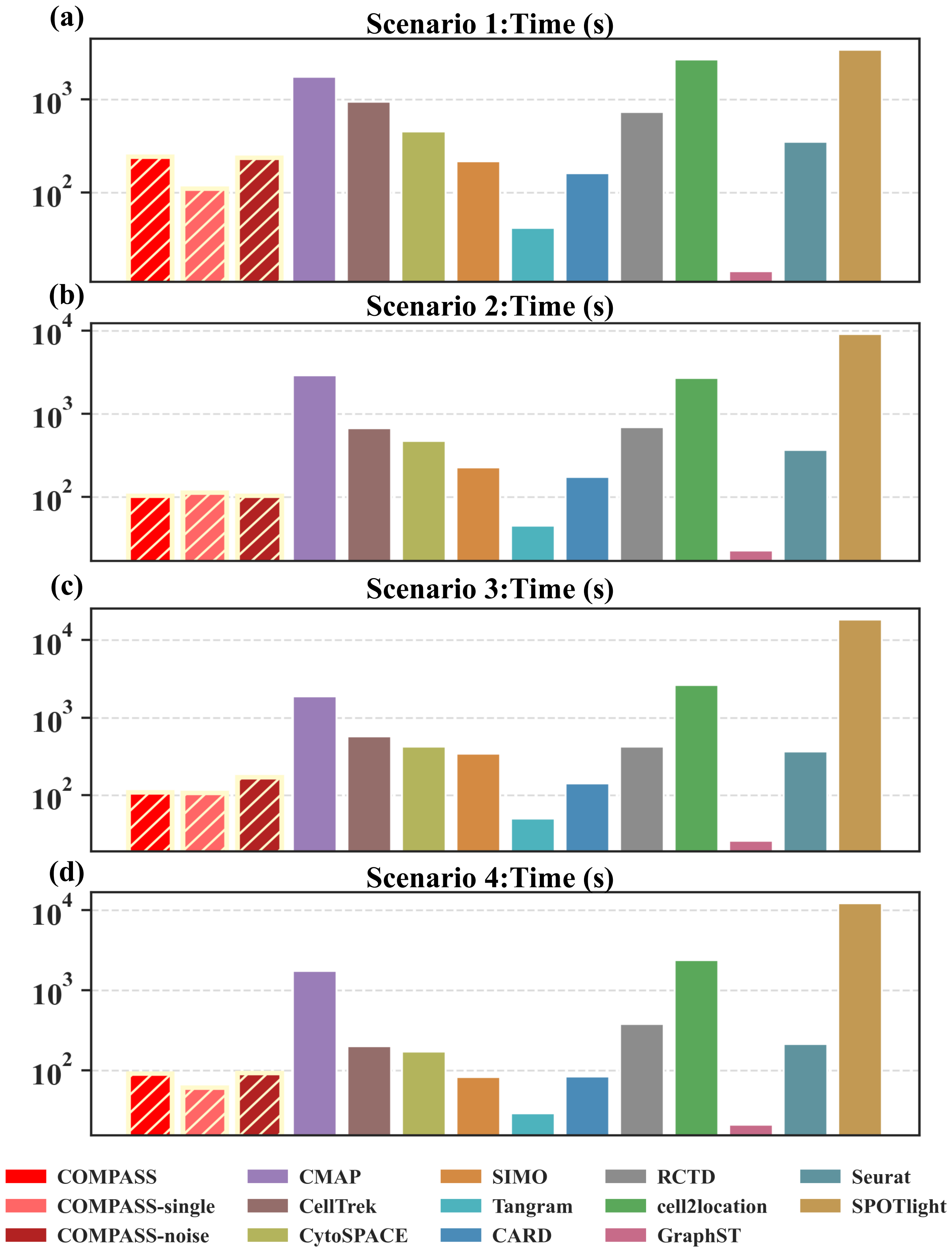

(a)
Scenario 1:Time (s)
$10^3$
$10^2$
(b)
Scenario 2:Time (s)
$10^4$
$10^3$
$10^2$
(c)
Scenario 3:Time (s)
$10^4$
$10^3$
$10^2$
(d)
Scenario 4:Time (s)
$10^4$
$10^3$
$10^2$
COMPASS
COMPASS-single
COMPASS-noise
CMAP
CellTrek
CytoSPACE
SIMO
Tangram
CARD
RCTD
cell2location
GraphST
Seurat
SPOTlight

Figure S69: **Runtime comparison across four simulation scenarios.** Bar plots compare the computational time of COMPASS, COMPASS-single, COMPASS-noise, and benchmark methods across four progressively challenging simulation settings, with the three COMPASS variants shown as hatched bars. (a) Scenario 1 uses unperturbed spatial RNA and auxiliary ATAC inputs. (b) Scenario 2 introduces measurement noise to the spatial RNA modality while leaving the auxiliary ATAC profiles intact. (c) Scenario 3 additionally masks 50% of the spot-level RNA entries to zero while retaining the paired ATAC modality. (d) Scenario 4 further removes cell states from the dissociated scRNA-seq reference on top of the Scenario 3 perturbations. Runtime is reported in seconds on a logarithmic scale. Runtime is reported in seconds on a logarithmic scale.

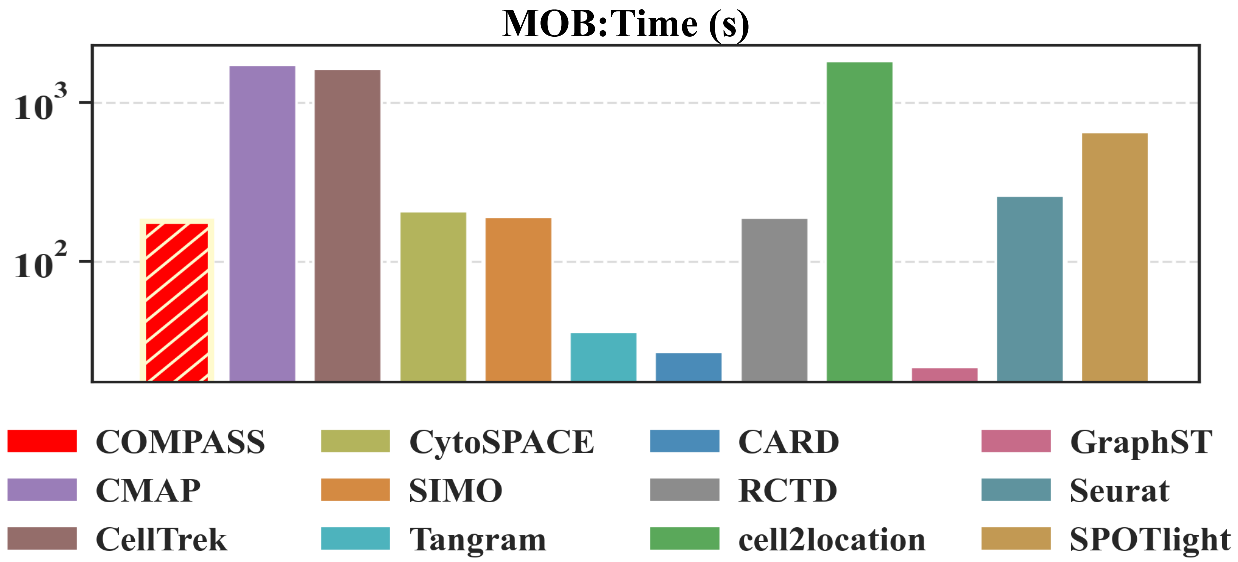

Figure S70: **Runtime comparison on the mouse olfactory bulb dataset.** Bar plots compare the computational time of COMPASS and benchmark methods on the MOB dataset, which evaluates reconstruction of the laminar olfactory bulb architecture. Runtime is reported in seconds on a logarithmic scale, summarizing the computational cost of each method under the same dataset setting.

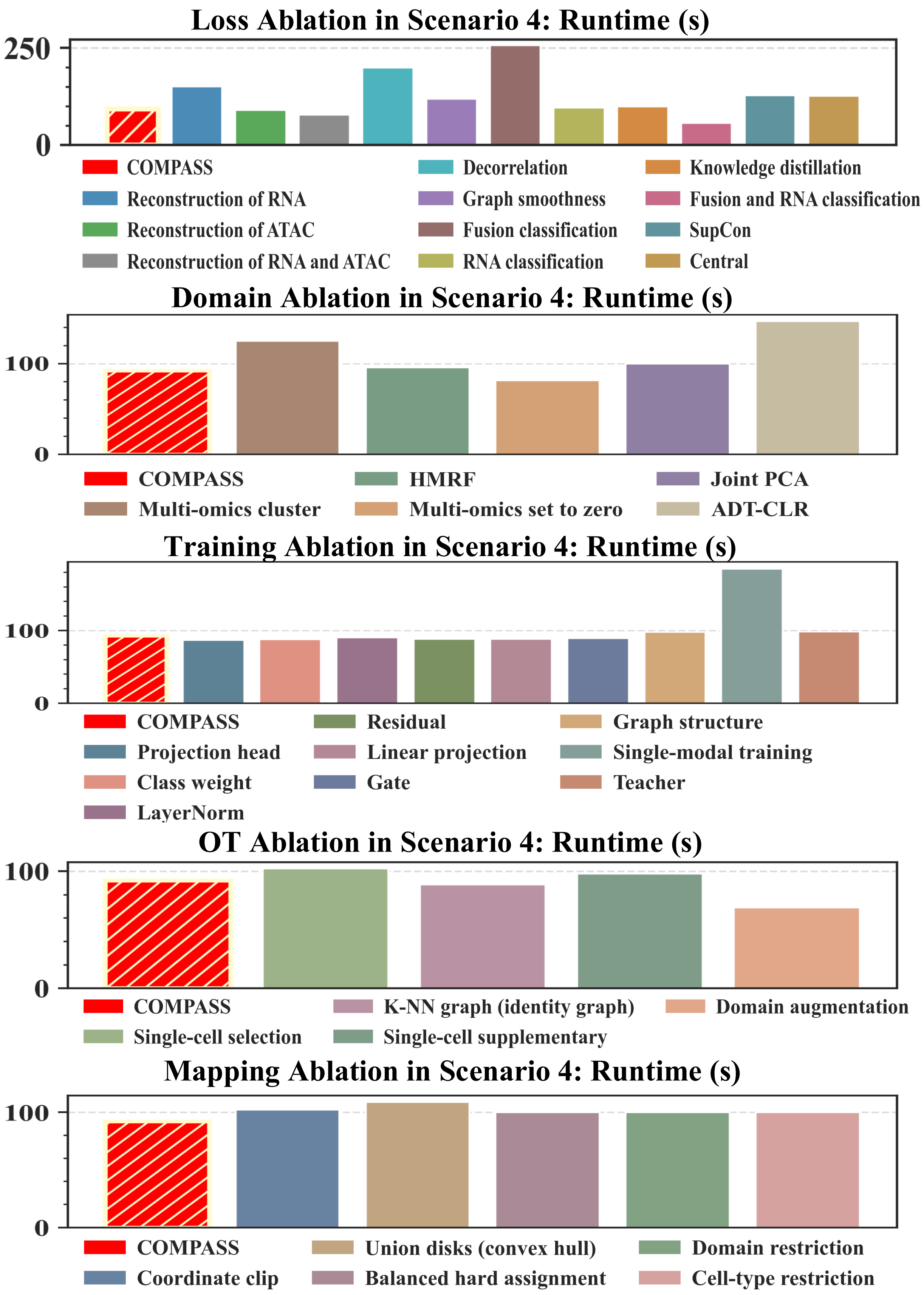

Figure S71: **Runtime of component ablation experiments in Scenario 4.** Bar plots show the computational time, in seconds, for the full COMPASS model and its ablated variants under the most challenging simulation setting. From top to bottom, panels summarize the runtime of loss-function ablations, domain-prior ablations, RG-COD training ablations, OT-based assignment ablations, and PDGR mapping ablations, with the hatched red bar denoting the full COMPASS model.

### Supplementary Notes

### 1 COMPASS method

This section provides the detailed methodological formulation of COMPASS under both single-omics and multi-omics settings. In the single-omics setting, COMPASS reconstructs a cell-resolved spatial atlas from spatial transcriptomics and dissociated scRNA-seq data by first delineating spatially coherent tissue domains from spatial RNA, using these domains as a structural prior for RNA-centered graph representation learning, and then assigning dissociated cells to spatial spots through domain-constrained and cell-type-stratified optimal transport. In the multi-omics setting, COMPASS jointly considers spatial RNA and co-measured auxiliary spatial omics during domain delineation and representation learning. Auxiliary modality-derived domain structure is transferred into the deployable RNA backbone through residual-gated cross-omics distillation, so that dissociated RNA-only cells and spatial spots can still be compared in a shared RNA-centered latent space. In both settings, the resulting spot-level assignments are further refined by Poisson-disk geometric refinement, which converts discrete cell-to-spot assignments into continuous sub-spot coordinates under locality, capacity-consistency, and minimum-separation constraints. Together, these components integrate domain-aware representation learning, structure-constrained cell-to-spot assignment, and geometry-aware coordinate reconstruction into a unified framework for precise spatial single-cell atlas assembly.

#### 1.1 Domain delineation for tissue-structure prior construction

COMPASS first infers coarse spatial domains to provide a tissue-structure prior for domain-aware representation learning and spatially constrained cell-to-spot mapping. Let $X^{\mathrm{st}} \in \mathbb{R}^{n \times g}$ denote the spatial transcriptomic matrix for $n$ spots and $g$ genes, with spatial coordinates $S = \{s_i\}_{i=1}^{n}$, $s_i \in \mathbb{R}^2$. When an auxiliary spatial omics modality is available on the same tissue section, we denote it by $X^{\mathrm{aux}} \in \mathbb{R}^{n \times p}$. COMPASS supports both RNA-only and multi-omics domain initialization, and the two settings differ only in the construction of the clustering embedding $u_i$.

In the RNA-only setting, COMPASS operates on the normalized and log-transformed spatial RNA matrix. In the implementation, preprocessing is applied adaptively: if the observed values are identified as raw count-like measurements, total-count normalization followed by a $\log(1 + x)$ transformation is performed; otherwise, the input matrix is used directly. Principal component analysis is then applied to the spot-by-gene RNA matrix, and the first $d_{\mathrm{rna}}$ principal components are retained, with $d_{\mathrm{rna}} = 30$ by default unless limited by the input dimensionality. Denoting the preprocessed spatial RNA matrix by $\widetilde{X}^{\mathrm{st}}$, the RNA-derived clustering features are

$$U^{\mathrm{rna}} = \mathrm{PCA}_{d_{\mathrm{rna}}}\left(\widetilde{X}^{\mathrm{st}}\right) \in \mathbb{R}^{n \times d_{\mathrm{rna}}}.$$

For RNA-only analysis, the spatial-domain clustering embedding is defined as

$$U = U^{\mathrm{rna}}, \qquad u_i = U_{i\cdot}.$$

In the multi-omics setting, COMPASS constructs a joint clustering embedding from spatial RNA and the co-measured auxiliary modality. The RNA branch is summarized by PCA as above. The auxiliary branch is preprocessed according to the data type. For protein or ADT measurements, centered log-ratio normalization followed by scaling and PCA is used. For chromatin accessibility measurements, TF–IDF transformation, logarithmic scaling, latent semantic indexing, and scaling are used. Let

$$U^{\text{rna}} \in \mathbb{R}^{n' \times d_{\text{rna}}}, \qquad U^{\text{aux}} \in \mathbb{R}^{n' \times d_{\text{aux}}}$$

denote the RNA and auxiliary low-dimensional features on the $n'$ spots shared by the two modalities. The two feature matrices are combined by weighted concatenation,

$$H = \left[\omega_{\text{rna}} U^{\text{rna}},\ \omega_{\text{aux}} U^{\text{aux}}\right],$$

where $\omega_{\text{rna}}$ and $\omega_{\text{aux}}$ control the relative contribution of RNA and auxiliary information. To reduce redundancy and stabilize mixture modeling, an optional joint PCA is further applied:

$$U = \begin{cases} \text{PCA}_{d_{\text{joint}}}(H), & d_{\text{joint}} > 0, \\ H, & d_{\text{joint}} = 0. \end{cases}$$

The row vector $u_i = U_{i\cdot}$ is then used as the clustering embedding of spot $i$. Thus, in both RNA-only and multi-omics modes, COMPASS performs spatial domain regularization on a unified spot-level embedding $u_i$, while allowing auxiliary omics to contribute to domain initialization when available.

A Gaussian mixture model with $K$ components is fitted to $\{u_i\}_{i=1}^{n'}$ to capture expression- or multi-omics-driven latent tissue states. The model provides posterior probabilities

$$\gamma_{ik} = p_{\text{GMM}}(y_i = k \mid u_i), \qquad k = 1, \dots, K,$$

where $y_i \in \{1, \dots, K\}$ denotes the latent domain label of spot $i$. The initial domain assignment is obtained by maximum posterior probability,

$$y_i^{(0)} = \arg\max_k \gamma_{ik}.$$

In practice, COMPASS uses a regularized GMM, with full covariance matrices attempted when numerically stable and more constrained covariance parameterizations are used as fallbacks when necessary.

To incorporate physical tissue topology, COMPASS constructs an undirected spatial neighborhood graph from the two-dimensional spot coordinates. By default, a binary $k$-nearest-neighbor graph with $k = 6$ is used, although radius-based neighborhoods can also be specified. Let $E$ denote the edge set of this spatial graph, and let $N_i$ denote the neighbors of spot $i$. Spatial domain inference is formulated as a Potts-type hidden Markov random field that balances molecular evidence and spatial smoothness:

$$\hat{\mathbf{y}} = \arg\max_{\mathbf{y}} \left\{ \sum_{i=1}^{n'} \log p_{\text{GMM}}(y_i \mid u_i) - \beta \sum_{(i,j) \in E} \mathbf{1}(y_i \neq y_j) \right\},$$

where $\beta > 0$ controls the strength of spatial regularization. Equivalently, this can be written as the minimization of the domain energy

$$\mathcal{L}_{\text{dom}}(\mathbf{y}) = \sum_{i=1}^{n'} -\log \gamma_{i,y_i} + \beta \sum_{(i,j) \in E} \mathbf{1}(y_i \neq y_j).$$

The unary term penalizes assignments inconsistent with the GMM-derived molecular state, whereas the pairwise Potts term penalizes label discontinuities between neighboring spots and therefore favors spatially contiguous domains.

Optimization is performed using iterated conditional modes. At iteration $t$, each spot is updated by minimizing its local conditional energy:

$$y_i^{(t+1)} = \arg \min_{c\in\{1,\dots,K\}} \left\{ -\log\gamma_{ic} + \beta \sum_{j\in N_i} \mathbf{1}(c \neq y_j^{(t)}) \right\}.$$

Thus, the pairwise penalty for assigning spot $i$ to candidate domain $c$ is proportional to the number of neighboring spots whose current labels differ from $c$. This iterative refinement suppresses isolated noisy assignments and improves spatial continuity while preserving molecularly supported domain boundaries. To avoid pathological over-smoothing, COMPASS checks whether the HMRF refinement[1] collapses to a degenerate single-domain solution; if so, the algorithm falls back to the original GMM initialization.

The resulting spatial domain labels $\hat{\mathbf{y}}$ are used in two downstream steps. First, they supervise the domain-aware graph representation module, including the RNA backbone and, when auxiliary omics are available, the multi-omics teacher branch. Second, they constrain cell-to-spot optimal transport within domain- and cell-type-specific buckets, thereby reducing cross-domain misassignment of transcriptionally similar cells. After domain delineation, COMPASS performs one-vs-rest differential expression analysis across inferred domains using the Wilcoxon rank-sum test on the spatial RNA matrix. The top-ranked genes from each domain are merged into a non-redundant RNA domain-marker set, with the top 50 genes per domain retained by default. These markers provide interpretable molecular signatures of the inferred spatial domains and serve as biological anchors for downstream cross-modal interpretation.

#### 1.2 Cross-modal graph alignment for RNA-centered representation learning

COMPASS learns a domain-aware RNA-centered latent space in which spatial spots and dissociated single cells can be directly compared for subsequent single-cell domain inference and domain-constrained cell-to-spot mapping. We use the same notation as in the main text. Let $X^{\mathrm{st}} \in \mathbb{R}^{n\times g}$ denote the spatial transcriptomic matrix for $n$ spatial spots and $g$ genes, and let $S = \{s_i\}_{i=1}^{n}$, $s_i \in \mathbb{R}^2$, denote the corresponding spot coordinates. Let $X^{\mathrm{aux}} \in \mathbb{R}^{n\times p}$ denote an optional auxiliary spatial omics matrix measured on the same tissue section, where $p$ is the number of auxiliary features. Let $X^{\mathrm{sc}} \in \mathbb{R}^{N\times g_{\mathrm{sc}}}$ denote the dissociated single-cell RNA-seq matrix for $N$ cells and $g_{\mathrm{sc}}$ measured genes. The spatial domain labels inferred by the domain delineation module are denoted by $\widehat{\mathbf{y}} = (\widehat{y}_1, \dots, \widehat{y}_n)$, where $\widehat{y}_i \in \{1, \dots, K\}$ and $K$ is the number of spatial domains. These domain labels provide the spot-level supervision for the graph representation module.

Because RNA is the only modality shared by spatial spots and dissociated single cells, COMPASS first aligns the two RNA datasets to a common spatial RNA template. The template is defined by the gene order of $X^{\mathrm{st}}$ together with the spatial-RNA-derived per-gene mean and standard deviation. Specifically, for gene $j \in \{1, \ldots, g\}$,

$$\mu_j^{\mathrm{st}} = \frac{1}{n} \sum_{i=1}^{n} X_{ij}^{\mathrm{st}}, \qquad \sigma_j^{\mathrm{st}} = \left[ \frac{1}{n} \sum_{i=1}^{n} \left( X_{ij}^{\mathrm{st}} - \mu_j^{\mathrm{st}} \right)^2 \right]^{1/2} + \varepsilon_\sigma,$$

where $\varepsilon_\sigma > 0$ is a small constant that avoids division by zero. The standardized spatial RNA matrix is

$$\overline{X}_{ij}^{\mathrm{st}} = \frac{X_{ij}^{\mathrm{st}} - \mu_j^{\mathrm{st}}}{\sigma_j^{\mathrm{st}}}, \qquad i = 1, \ldots, n, \quad j = 1, \ldots, g.$$

The single-cell RNA matrix is reordered to match the spatial RNA gene template. If gene $j$ is present in $X^{\mathrm{st}}$ but absent from $X^{\mathrm{sc}}$, the corresponding single-cell entries are imputed by $\mu_j^{\mathrm{st}}$, so that their standardized values are zero. Let $\overline{X}^{\mathrm{sc}} \in \mathbb{R}^{N \times g}$ denote the resulting reordered, completed, and standardized single-cell RNA matrix. PCA is then fitted on $\overline{X}^{\mathrm{st}}$, and the same transformation is applied to $\overline{X}^{\mathrm{sc}}$:

$$\widetilde{X}_{\mathrm{st}} = R_{\mathrm{rna}} \left( \overline{X}^{\mathrm{st}} \right) \in \mathbb{R}^{n \times d_{\mathrm{rna}}}, \qquad \widetilde{X}_{\mathrm{sc}} = R_{\mathrm{rna}} \left( \overline{X}^{\mathrm{sc}} \right) \in \mathbb{R}^{N \times d_{\mathrm{rna}}},$$

where $R_{\mathrm{rna}}$ denotes the PCA transformation fitted on spatial RNA and $d_{\mathrm{rna}}$ is the retained RNA feature dimension. This construction ensures that spatial RNA and single-cell RNA are represented in the same low-dimensional RNA coordinate system before graph encoding.

When an auxiliary spatial omics modality is available, multi-omics representation learning is performed on the spots shared by $X^{\mathrm{st}}$ and $X^{\mathrm{aux}}$. Let $\mathcal{I}_\cap \subseteq \{1, \ldots, n\}$ denote the set of common spots and $n_\cap = |\mathcal{I}_\cap|$. In the multi-omics setting, the spatial matrices, spatial graph, and domain labels are restricted to $\mathcal{I}_\cap$, but we keep the same notation for readability. For ADT or protein measurements, COMPASS applies centered log-ratio normalization. For spot $i$ and auxiliary feature $\ell$,

$$X_{i\ell}^{\mathrm{aux,clr}} = \log \frac{X_{i\ell}^{\mathrm{aux}} + \varepsilon_{\mathrm{clr}}}{\left[ \prod_{m=1}^{p} \left( X_{im}^{\mathrm{aux}} + \varepsilon_{\mathrm{clr}} \right) \right]^{1/p}},$$

where $\varepsilon_{\mathrm{clr}} > 0$ is a small pseudo-count. The CLR-normalized matrix is then feature-wise standardized and reduced by PCA. For chromatin accessibility data, let $B \in \mathbb{R}^{n_\cap \times p}$ denote the spot-by-peak count matrix. COMPASS uses a TF–IDF transformation followed by logarithmic scaling, latent semantic indexing, and standardization. Specifically,

$$\mathrm{tf}_{i\ell} = \frac{B_{i\ell}}{\sum_{m=1}^{p} B_{im} + \varepsilon}, \qquad \mathrm{idf}_\ell = \log \left( 1 + \frac{n_\cap}{\mathrm{df}_\ell + 1} \right),$$

where $\varepsilon > 0$ is a numerical stabilizer and $\mathrm{df}_\ell = |\{i : B_{i\ell} > 0\}|$ is the number of spots with nonzero accessibility at feature $\ell$. The transformed accessibility feature is

$$B_{i\ell}^{\mathrm{tfidf}} = \log \left( 1 + 10^4 \, \mathrm{tf}_{i\ell} \mathrm{idf}_\ell \right).$$

After modality-specific preprocessing and dimensionality reduction, the auxiliary low-dimensional feature matrix is denoted by

$$\widetilde{X}_{\mathrm{aux}} \in \mathbb{R}^{n_\cap \times d_{\mathrm{aux}}},$$

where $d_{\mathrm{aux}}$ is the retained auxiliary feature dimension. Thus, $\widetilde{X}_{\mathrm{st}}$, $\widetilde{X}_{\mathrm{sc}}$, and $\widetilde{X}_{\mathrm{aux}}$ denote low-dimensional inputs to the graph representation module rather than original count matrices.

COMPASS next constructs graph structures for spatial spots and dissociated cells. For spatial spots, an undirected $k$-nearest-neighbor graph is constructed from the Euclidean distances between spot coordinates. Let $A_{\mathrm{st}} \in \mathbb{R}^{n_{\mathrm{st}} \times n_{\mathrm{st}}}$ denote the binary spatial adjacency matrix after symmetrization, where $n_{\mathrm{st}} = n$ in RNA-only analysis and $n_{\mathrm{st}} = n_{\cap}$ in multi-omics analysis. After adding self-loops, the normalized spatial propagation matrix is

$$\widetilde{A}_{\mathrm{st}} = D_{\mathrm{st}}^{-1/2} \left(A_{\mathrm{st}} + I\right) D_{\mathrm{st}}^{-1/2},$$

where $I$ is the identity matrix and $D_{\mathrm{st}}$ is the diagonal degree matrix of $A_{\mathrm{st}} + I$. In the multi-omics setting, the same $\widetilde{A}_{\mathrm{st}}$ is used for spatial RNA and auxiliary omics because both modalities are measured on the same spot graph. For dissociated single cells, a $k$-nearest-neighbor graph is constructed in the aligned RNA feature space $\widetilde{X}_{\mathrm{sc}}$. Let $A_{\mathrm{sc}} \in \mathbb{R}^{N \times N}$ denote the corresponding binary adjacency matrix, and let

$$\widetilde{A}_{\mathrm{sc}} = D_{\mathrm{sc}}^{-1/2} \left(A_{\mathrm{sc}} + I\right) D_{\mathrm{sc}}^{-1/2}$$

denote the normalized single-cell propagation matrix, where $D_{\mathrm{sc}}$ is the degree matrix of $A_{\mathrm{sc}} + I$. In mini-batch training, $A_{\mathrm{sc}}$ and $\widetilde{A}_{\mathrm{sc}}$ are replaced by the batch-induced subgraph built from precomputed nearest neighbors.

The RNA and auxiliary modalities are encoded by two-layer graph encoders. For spatial RNA, the encoder shares the same formulation as Eq. (2) in the main text:

$$Z_{\mathrm{st}} = \widetilde{A}_{\mathrm{st}} \cdot \sigma\left(\widetilde{A}_{\mathrm{st}} \widetilde{X}_{\mathrm{st}} W_{\mathrm{rna}}^{(1)}\right) \cdot W_{\mathrm{rna}}^{(2)},$$

where $W_{\mathrm{rna}}^{(1)}$ and $W_{\mathrm{rna}}^{(2)}$ are trainable RNA encoder weights, $\sigma(\cdot)$ denotes the ReLU activation, and $Z_{\mathrm{st}} \in \mathbb{R}^{n_{\mathrm{st}} \times d_z}$ is the spatial RNA latent embedding with latent dimension $d_z$. For auxiliary spatial omics, the modality-specific encoder is

$$Z_{\mathrm{aux}} = \widetilde{A}_{\mathrm{st}} \cdot \sigma\left(\widetilde{A}_{\mathrm{st}} \widetilde{X}_{\mathrm{aux}} W_{\mathrm{aux}}^{(1)}\right) \cdot W_{\mathrm{aux}}^{(2)},$$

where $W_{\mathrm{aux}}^{(1)}$ and $W_{\mathrm{aux}}^{(2)}$ are auxiliary encoder weights, and $Z_{\mathrm{aux}} \in \mathbb{R}^{n_{\cap} \times d_z}$. Dissociated single cells are encoded by the same RNA backbone used for spatial RNA:

$$Z_{\mathrm{sc}} = \widetilde{A}_{\mathrm{sc}} \cdot \sigma\left(\widetilde{A}_{\mathrm{sc}} \widetilde{X}_{\mathrm{sc}} W_{\mathrm{rna}}^{(1)}\right) \cdot W_{\mathrm{rna}}^{(2)},$$

where $Z_{\mathrm{sc}} \in \mathbb{R}^{N \times d_z}$. The sharing of $W_{\mathrm{rna}}^{(1)}$ and $W_{\mathrm{rna}}^{(2)}$ ensures that spatial RNA spots and dissociated cells are embedded by the same deployable RNA encoder.

Each encoder is equipped with three auxiliary heads used during training. The RNA reconstruction head is denoted by $D_{\mathrm{rna}} : \mathbb{R}^{d_z} \to \mathbb{R}^{d_{\mathrm{rna}}}$, and the auxiliary reconstruction head is denoted by $D_{\mathrm{aux}} : \mathbb{R}^{d_z} \to \mathbb{R}^{d_{\mathrm{aux}}}$. These heads reconstruct low-dimensional input features, not the original count matrices. The student domain-classification head is $C_{\mathrm{student}} : \mathbb{R}^{d_z} \to \mathbb{R}^{K}$, and the teacher domain-classification head is $C_{\mathrm{teacher}} : \mathbb{R}^{d_z} \to \mathbb{R}^{K}$. Projection heads used for contrastive learning and distribution alignment are denoted by $Q_{\mathrm{rna}} : \mathbb{R}^{d_z} \to \mathbb{R}^{d_q}$ and $Q_{\mathrm{teacher}} : \mathbb{R}^{d_z} \to \mathbb{R}^{d_q}$, where $d_q$ is the projection dimension.

The following loss functions are used by the graph representation module. Let $\mathcal{I}_{\mathrm{lab}} \subseteq \{1, \ldots, n_{\mathrm{st}}\}$ be the set of spots with valid inferred domain labels, and let $n_{\mathrm{lab}} = |\mathcal{I}_{\mathrm{lab}}|$. For a logit matrix $l \in \mathbb{R}^{n_{\mathrm{st}} \times K}$, define

$$p_{ik}(l) = \frac{\exp(l_{ik})}{\sum_{k'=1}^{K} \exp(l_{ik'})}, \qquad i = 1, \ldots, n_{\mathrm{st}}, \quad k = 1, \ldots, K.$$

Let $r_k$ denote the empirical frequency of domain $k$ among labeled spots:

$$r_k = \frac{1}{n_{\mathrm{lab}}} \sum_{i \in \mathcal{I}_{\mathrm{lab}}} \mathbf{1}\left(\widehat{y}_i = k\right), \qquad k = 1, \ldots, K.$$

The class weight for domain $k$ is

$$\omega_k = \frac{(r_k + \varepsilon_w)^{-1}}{K^{-1} \sum_{k'=1}^{K} (r_{k'} + \varepsilon_w)^{-1}},$$

where $\varepsilon_w > 0$ is a numerical stabilizer. The class-weighted domain cross-entropy loss is

$$\mathcal{L}_{\mathrm{CE}}\left(l, \widehat{\mathbf{y}}\right) = -\frac{1}{n_{\mathrm{lab}}} \sum_{i \in \mathcal{I}_{\mathrm{lab}}} \omega_{\widehat{y}_i} \log p_{i,\widehat{y}_i}(l).$$

For a latent embedding $Z \in \mathbb{R}^{m \times d_z}$, an input feature matrix $X \in \mathbb{R}^{m \times d_x}$, and a reconstruction head $D : \mathbb{R}^{d_z} \to \mathbb{R}^{d_x}$, the feature reconstruction loss is

$$\mathcal{L}_{\mathrm{rec}}(Z, X; D) = \frac{1}{m d_x} \left\| D(Z) - X \right\|_F^2 .$$

In particular,

$$\mathcal{L}_{\mathrm{rec}}^{\mathrm{rna}} = \mathcal{L}_{\mathrm{rec}}\left(Z_{\mathrm{st}}, \widetilde{X}_{\mathrm{st}}; D_{\mathrm{rna}}\right), \qquad \mathcal{L}_{\mathrm{rec}}^{\mathrm{aux}} = \mathcal{L}_{\mathrm{rec}}\left(Z_{\mathrm{aux}}, \widetilde{X}_{\mathrm{aux}}; D_{\mathrm{aux}}\right).$$

The spatial smoothness loss is defined for a spatial embedding $Z \in \mathbb{R}^{n_{\mathrm{st}} \times d_z}$ as

$$\mathcal{L}_{\mathrm{smooth}}(Z) = \frac{1}{n_{\mathrm{st}}} \left\| \widetilde{A}_{\mathrm{st}} Z - Z \right\|_F^2 .$$

This loss penalizes rapid embedding changes across neighboring spots in the spatial graph.

The supervised contrastive loss is computed on projected spatial embeddings. Let $Q$ denote the projection head used for the embedding $Z$, and define the normalized projected vector

$$h_i(Z, Q) = \frac{Q(Z_i)}{\left\| Q(Z_i) \right\|_2 + \varepsilon_h}, \qquad i = 1, \ldots, n_{\mathrm{st}},$$

where $Z_i$ is the $i$-th row of $Z$ and $\varepsilon_h > 0$ avoids division by zero. For each labeled spot $i$, define its positive set as

$$\mathcal{P}(i) = \left\{ j \in \mathcal{I}_{\mathrm{lab}} : j \neq i,\, \widehat{y}_j = \widehat{y}_i \right\},$$

and define

$$\mathcal{I}_+ = \left\{ i \in \mathcal{I}_{\mathrm{lab}} : |\mathcal{P}(i)| > 0 \right\}.$$

With temperature $\tau > 0$, the supervised contrastive loss is

$$\mathcal{L}_{\mathrm{con}}\left(Z, \widehat{\mathbf{y}}; Q\right) = -\frac{1}{|\mathcal{I}_+|} \sum_{i \in \mathcal{I}_+} \log \frac{\sum_{j \in \mathcal{P}(i)} \exp\left(h_i(Z, Q)^\top h_j(Z, Q)/\tau\right)}{\sum_{j \in \mathcal{I}_{\mathrm{lab}},\, j \neq i} \exp\left(h_i(Z, Q)^\top h_j(Z, Q)/\tau\right)}.$$

The center loss uses a learnable center $m_k \in \mathbb{R}^{d_z}$ for each domain $k$. Let $M = \{m_1, \ldots, m_K\}$. The center loss is

$$\mathcal{L}_{\mathrm{center}}\left(Z, \widehat{\mathbf{y}}; M\right) = \frac{1}{n_{\mathrm{lab}}} \sum_{i \in \mathcal{I}_{\mathrm{lab}}} \left\| Z_i - m_{\widehat{y}_i} \right\|_2^2 .$$

To reduce redundant covariance among latent dimensions, COMPASS optionally uses a DeCov regularizer. For $Z \in \mathbb{R}^{m\times d_z}$, define the empirical mean

$$\bar{Z} = \frac{1}{m}\sum_{i=1}^{m} Z_i$$

and covariance matrix

$$C_Z = \frac{(Z - \mathbf{1}_m\bar{Z})^\top (Z - \mathbf{1}_m\bar{Z})}{m - 1 + \varepsilon_c},$$

where $\mathbf{1}_m \in \mathbb{R}^{m\times 1}$ is an all-one column vector and $\varepsilon_c > 0$ is a stabilizer. The DeCov loss is

$$\mathcal{L}_{\text{decov}}(Z) = \frac{1}{d_z^2}\sum_{\substack{a,b=1\\ a\neq b}}^{d_z} (C_Z)_{ab}^2.$$

In RNA-only training, spatial spots and dissociated single cells are additionally aligned at the distribution level in the projected RNA latent space. Let $\mathcal{B}_{\text{sc}} \subseteq \{1,\ldots,N\}$ denote a mini-batch of single cells, and let $B = |\mathcal{B}_{\text{sc}}|$. Let $Z_{\text{sc}}^{\mathcal{B}} \in \mathbb{R}^{B\times d_z}$ denote the single-cell embedding restricted to this mini-batch. The projected spatial and single-cell embeddings are

$$U_{\text{st}} = Q_{\text{rna}}(Z_{\text{st}}) \in \mathbb{R}^{n_{\text{st}}\times d_q}, \qquad U_{\text{sc}} = Q_{\text{rna}}(Z_{\text{sc}}^{\mathcal{B}}) \in \mathbb{R}^{B\times d_q}.$$

For a set of kernel bandwidth parameters $\Gamma$, define the multi-kernel radial basis function

$$k_\Gamma(x,y) = \sum_{\gamma\in\Gamma} \exp\left(-\gamma \|x-y\|_2^2\right).$$

The maximum mean discrepancy loss is

$$\begin{aligned}\mathcal{L}_{\text{MMD}}\left(U_{\text{st}}, U_{\text{sc}}\right) =& \frac{1}{n_{\text{st}}(n_{\text{st}}-1)}\sum_{\substack{i,i'=1\\ i\neq i'}}^{n_{\text{st}}} k_\Gamma\left(U_{\text{st},i}, U_{\text{st},i'}\right)\\ &+ \frac{1}{B(B-1)}\sum_{\substack{a,a'=1\\ a\neq a'}}^{B} k_\Gamma\left(U_{\text{sc},a}, U_{\text{sc},a'}\right)\\ &- \frac{2}{n_{\text{st}}B}\sum_{i=1}^{n_{\text{st}}}\sum_{a=1}^{B} k_\Gamma\left(U_{\text{st},i}, U_{\text{sc},a}\right).\end{aligned}$$

The CORAL loss aligns second-order statistics. For a matrix $U \in \mathbb{R}^{m\times d_q}$, define

$$\text{Cov}(U) = \frac{(U - \mathbf{1}_m\bar{U})^\top (U - \mathbf{1}_m\bar{U})}{m - 1 + \varepsilon_c}, \qquad \bar{U} = \frac{1}{m}\sum_{i=1}^{m} U_i.$$

Then

$$\mathcal{L}_{\text{CORAL}}\left(U_{\text{st}}, U_{\text{sc}}\right) = \frac{1}{4d_q^2}\left\|\text{Cov}(U_{\text{st}}) - \text{Cov}(U_{\text{sc}})\right\|_F^2.$$

COMPASS also includes distribution-balancing terms to avoid degenerate domain predictions. For spatial logits $l_{\text{st}}$, define the mean predicted domain distribution over labeled spots as

$$\bar{p}_k = \frac{1}{n_{\text{lab}}}\sum_{i\in\mathcal{I}_{\text{lab}}} p_{ik}(l_{\text{st}}), \qquad k = 1,\ldots,K.$$

The quadratic balancing loss and KL balancing loss are

$$\mathcal{L}_{\text{bal}} = \sum_{k=1}^{K} (\bar{p}_k - r_k)^2 ,$$

$$\mathcal{L}_{\text{balKL}} = \text{KL}\,(r \| \bar{p}) = \sum_{k=1}^{K} r_k \text{log} \frac{r_k + \varepsilon}{\bar{p}_k + \varepsilon},$$

where $r = (r_1, \ldots, r_K)$, $\bar{p} = (\bar{p}_1, \ldots, \bar{p}_K)$, and $\varepsilon > 0$ is a stabilizer.

When single-cell cell-type annotations are available, COMPASS uses an additional cellType–domain regularizer on the single-cell domain predictions. Let $c_a \in \mathcal{C}$ denote the cell-type label of single cell $a$, where $\mathcal{C}$ is the set of cell types. For a single-cell mini-batch $\mathcal{B}_{\text{sc}}$, let

$$q_{ak} = p_{ak}\,(l_{\text{sc}})\,, \qquad a \in \mathcal{B}_{\text{sc}}, \quad k = 1, \ldots, K,$$

where $l_{\text{sc}} = C_{\text{student}}(Z_{\text{sc}}^{\mathcal{B}})$. Let $\mathcal{B}_t = \{a \in \mathcal{B}_{\text{sc}} : c_a = t\}$ be the set of mini-batch cells of type $t$, and let $\mathcal{T}_B = \{t : |\mathcal{B}_t| \geq b_{\min}\}$ be the cell types represented by at least $b_{\min}$ cells in the mini-batch. Define the global single-cell domain distribution and the type-specific domain distribution as

$$\bar{q}_k = \frac{1}{B} \sum_{a \in \mathcal{B}_{\text{sc}}} q_{ak}, \qquad \bar{q}_{t,k} = \frac{1}{|\mathcal{B}_t|} \sum_{a \in \mathcal{B}_t} q_{ak}.$$

Let

$$u_K = \left( \frac{1}{K}, \ldots, \frac{1}{K} \right)$$

denote the uniform distribution over domains. The entropy of the domain distribution for cell type $t$ is

$$H(\bar{q}_t) = -\sum_{k=1}^{K} \bar{q}_{t,k} \text{log}\,(\bar{q}_{t,k} + \varepsilon)\,.$$

The cellType–domain regularizer is

$$\mathcal{L}_{\text{ctdom}} = \frac{1}{|\mathcal{T}_B|} \sum_{t \in \mathcal{T}_B} H(\bar{q}_t) + \text{KL}\,(u_K \| \bar{q}) - \frac{2}{|\mathcal{T}_B|(|\mathcal{T}_B| - 1)} \sum_{\substack{t,t' \in \mathcal{T}_B \\ t < t'}} \frac{1}{K} \|\bar{q}_t - \bar{q}_{t'}\|_2^2 \,.$$

The first term encourages each cell type to concentrate on a limited set of predicted domains, the second term prevents global single-cell domain collapse, and the third term encourages different cell types to have distinguishable domain-usage profiles.

In the RNA-only setting, no auxiliary spatial omics matrix is used. Spatial spots and dissociated single cells are both encoded by the same RNA graph encoder:

$$Z_{\text{st}} = \widetilde{A}_{\text{st}} \cdot \sigma \left( \widetilde{A}_{\text{st}} \widetilde{X}_{\text{st}} W_{\text{rna}}^{(1)} \right) \cdot W_{\text{rna}}^{(2)},$$

$$Z_{\text{sc}} = \widetilde{A}_{\text{sc}} \cdot \sigma \left( \widetilde{A}_{\text{sc}} \widetilde{X}_{\text{sc}} W_{\text{rna}}^{(1)} \right) \cdot W_{\text{rna}}^{(2)}.$$

The spatial and single-cell domain logits are

$$l_{\text{st}} = C_{\text{student}}\,(Z_{\text{st}})\,, \qquad l_{\text{sc}} = C_{\text{student}}\,\left(Z_{\text{sc}}^{\mathcal{B}}\right).$$

The RNA-only training objective is

$$\begin{aligned}\mathcal{L}_{\text{single}} = & \lambda_{\text{rec}}^{\text{rna}}\mathcal{L}_{\text{rec}}^{\text{rna}} + \lambda_{\text{CE}}\mathcal{L}_{\text{CE}}\left(l_{\text{st}}, \widehat{\mathbf{y}}\right) + \lambda_{\text{smooth}}\mathcal{L}_{\text{smooth}}\left(Z_{\text{st}}\right) \\ & + \lambda_{\text{con}}\mathcal{L}_{\text{con}}\left(Z_{\text{st}}, \widehat{\mathbf{y}}; Q_{\text{rna}}\right) + \lambda_{\text{center}}\mathcal{L}_{\text{center}}\left(Z_{\text{st}}, \widehat{\mathbf{y}}; M\right) + \lambda_{\text{decov}}\mathcal{L}_{\text{decov}}\left(Z_{\text{st}}\right) \\ & + \lambda_{\text{MMD}}\mathcal{L}_{\text{MMD}}\left(Q_{\text{rna}}(Z_{\text{st}}), Q_{\text{rna}}(Z_{\text{sc}}^{\mathcal{B}})\right) + \lambda_{\text{CORAL}}\mathcal{L}_{\text{CORAL}}\left(Q_{\text{rna}}(Z_{\text{st}}), Q_{\text{rna}}(Z_{\text{sc}}^{\mathcal{B}})\right) \\ & + \lambda_{\text{bal}}\mathcal{L}_{\text{bal}} + \lambda_{\text{balKL}}\mathcal{L}_{\text{balKL}} + \lambda_{\text{ctdom}}\mathcal{L}_{\text{ctdom}}.\end{aligned}$$

All $\lambda$'s are nonnegative scalar hyperparameters controlling the relative contribution of the corresponding loss terms. The symbol $\beta$ is not used for these training weights, because $\beta$ is reserved for the Potts smoothness parameter in the HMRF domain delineation objective in the main text.

In the multi-omics setting, the spatial side contains both RNA and auxiliary omics, whereas the dissociated single-cell reference contains only RNA. Therefore, COMPASS does not use a fully fused multi-omics representation as the final deployable mapping space. Instead, it uses Residual-Gated Cross-Omics Distillation to transfer auxiliary-derived domain structure into the RNA-centered backbone. The RNA and auxiliary spatial embeddings are first computed as

$$Z_{\text{st}} = \widetilde{A}_{\text{st}} \cdot \sigma\left(\widetilde{A}_{\text{st}}\widetilde{X}_{\text{st}}W_{\text{rna}}^{(1)}\right) \cdot W_{\text{rna}}^{(2)},$$

$$Z_{\text{aux}} = \widetilde{A}_{\text{st}} \cdot \sigma\left(\widetilde{A}_{\text{st}}\widetilde{X}_{\text{aux}}W_{\text{aux}}^{(1)}\right) \cdot W_{\text{aux}}^{(2)}.$$

The auxiliary embedding is projected into the RNA latent space by a learnable projection $P : \mathbb{R}^{d_z} \to \mathbb{R}^{d_z}$:

$$Z_{\text{aux}\to\text{rna}} = P\left(Z_{\text{aux}}\right).$$

A spot-specific residual gate is computed as

$$G = g\left(\left[Z_{\text{st}}, Z_{\text{aux}\to\text{rna}}\right]\right), \qquad G \in (0,1)^{n_\cap \times d_z},$$

where $g : \mathbb{R}^{2d_z} \to (0,1)^{d_z}$ is a sigmoid gating network applied row-wise and $[\cdot,\cdot]$ denotes concatenation along the feature dimension. The multi-omics teacher representation is

$$\widehat{Z}_{\text{st}} = \text{LN}\left(Z_{\text{st}} + G \odot Z_{\text{aux}\to\text{rna}}\right),$$

where $\odot$ denotes element-wise multiplication and $\text{LN}(\cdot)$ denotes layer normalization. The residual form makes auxiliary information a selective correction to the RNA embedding rather than a replacement of the RNA representation. The RNA-only spatial embedding $Z_{\text{st}}$ is the student representation.

The teacher and student domain logits are

$$l_{\text{teacher}} = C_{\text{teacher}}\left(\widehat{Z}_{\text{st}}\right), \qquad l_{\text{student}} = C_{\text{student}}\left(Z_{\text{st}}\right).$$

For temperature $T > 0$, define the softened teacher and student distributions as

$$s_{ik}^{\text{teacher}} = \frac{\exp\left(l_{\text{teacher},ik}/T\right)}{\sum_{k'=1}^{K}\exp\left(l_{\text{teacher},ik'}/T\right)}, \qquad s_{ik}^{\text{student}} = \frac{\exp\left(l_{\text{student},ik}/T\right)}{\sum_{k'=1}^{K}\exp\left(l_{\text{student},ik'}/T\right)}.$$

The knowledge-distillation loss is

$$\mathcal{L}_{\mathrm{KD}} = \frac{T^2}{n_{\mathrm{lab}}} \sum_{i \in \mathcal{I}_{\mathrm{lab}}} \sum_{k=1}^{K} \mathrm{sg}\left[s_{ik}^{\mathrm{teacher}}\right] \log \frac{\mathrm{sg}\left[s_{ik}^{\mathrm{teacher}}\right] + \varepsilon}{s_{ik}^{\mathrm{student}} + \varepsilon},$$

where $\mathrm{sg}[\cdot]$ denotes stop-gradient on the teacher distribution. The unweighted RG-COD loss corresponding to the main text is

$$\mathcal{L}_{\text{RG-COD}} = \mathcal{L}_{\mathrm{KD}} + \mathcal{L}_{\mathrm{CE}}\left(l_{\mathrm{teacher}}, \widehat{\mathbf{y}}\right) + \mathcal{L}_{\mathrm{CE}}\left(l_{\mathrm{student}}, \widehat{\mathbf{y}}\right).$$

In the weighted implementation, the same objective is written as

$$\mathcal{L}_{\text{RG-COD}}^{\mathrm{w}} = \lambda_{\mathrm{teacher}} \mathcal{L}_{\mathrm{CE}}\left(l_{\mathrm{teacher}}, \widehat{\mathbf{y}}\right) + \lambda_{\mathrm{student}} \mathcal{L}_{\mathrm{CE}}\left(l_{\mathrm{student}}, \widehat{\mathbf{y}}\right) + \lambda_{\mathrm{KD}} \mathcal{L}_{\mathrm{KD}},$$

where $\lambda_{\mathrm{teacher}}$, $\lambda_{\mathrm{student}}$, and $\lambda_{\mathrm{KD}}$ are nonnegative weights for teacher supervision, student supervision, and distillation, respectively.

The multi-omics reconstruction loss is modality-specific:

$$\mathcal{L}_{\mathrm{rec}}^{\mathrm{multi}} = \lambda_{\mathrm{rec}}^{\mathrm{rna}} \mathcal{L}_{\mathrm{rec}}^{\mathrm{rna}} + \lambda_{\mathrm{rec}}^{\mathrm{aux}} \mathcal{L}_{\mathrm{rec}}^{\mathrm{aux}},$$

where $\lambda_{\mathrm{rec}}^{\mathrm{rna}}$ and $\lambda_{\mathrm{rec}}^{\mathrm{aux}}$ weight the RNA and auxiliary reconstruction terms. The spatial regularization terms in the RG-COD branch are applied to the teacher embedding $\widehat{Z}_{\mathrm{st}}$, because this is the spatial representation that incorporates both RNA and auxiliary domain information:

$$\mathcal{L}_{\mathrm{smooth}}^{\mathrm{multi}} = \mathcal{L}_{\mathrm{smooth}}\left(\widehat{Z}_{\mathrm{st}}\right),$$

$$\mathcal{L}_{\mathrm{con}}^{\mathrm{multi}} = \mathcal{L}_{\mathrm{con}}\left(\widehat{Z}_{\mathrm{st}}, \widehat{\mathbf{y}}; Q_{\mathrm{teacher}}\right),$$

$$\mathcal{L}_{\mathrm{center}}^{\mathrm{multi}} = \mathcal{L}_{\mathrm{center}}\left(\widehat{Z}_{\mathrm{st}}, \widehat{\mathbf{y}}; M\right),$$

$$\mathcal{L}_{\mathrm{decov}}^{\mathrm{multi}} = \mathcal{L}_{\mathrm{decov}}\left(\widehat{Z}_{\mathrm{st}}\right).$$

The default multi-omics objective is

$$\begin{aligned}\mathcal{L}_{\mathrm{multi}} = & \mathcal{L}_{\mathrm{rec}}^{\mathrm{multi}} + \mathcal{L}_{\text{RG-COD}}^{\mathrm{w}} + \lambda_{\mathrm{smooth}} \mathcal{L}_{\mathrm{smooth}}^{\mathrm{multi}} \\ & + \lambda_{\mathrm{con}} \mathcal{L}_{\mathrm{con}}^{\mathrm{multi}} + \lambda_{\mathrm{center}} \mathcal{L}_{\mathrm{center}}^{\mathrm{multi}} + \lambda_{\mathrm{decov}} \mathcal{L}_{\mathrm{decov}}^{\mathrm{multi}}.\end{aligned}$$

This objective separates the roles of the two modalities. The auxiliary modality provides a domain-informative teacher signal through $\widehat{Z}_{\mathrm{st}}$, whereas the RNA student branch $Z_{\mathrm{st}}$ is trained to absorb this auxiliary-derived information through direct domain supervision and knowledge distillation. After training, the dissociated single cells are encoded by the same distilled RNA backbone:

$$Z_{\mathrm{sc}} = \widetilde{A}_{\mathrm{sc}} \cdot \sigma\left(\widetilde{A}_{\mathrm{sc}} \widetilde{X}_{\mathrm{sc}} W_{\mathrm{rna}}^{(1)}\right) \cdot W_{\mathrm{rna}}^{(2)}.$$

Thus, both the spatial RNA embedding $Z_{\mathrm{st}}$ and the dissociated-cell embedding $Z_{\mathrm{sc}}$ lie in the same deployable RNA-centered latent space. The student classifier can also be applied to $Z_{\mathrm{sc}}$ to obtain single-cell domain probabilities,

$$l_{\mathrm{sc}} = C_{\mathrm{student}}\left(Z_{\mathrm{sc}}\right), \qquad p_{ak}^{\mathrm{sc}} = \frac{\exp(l_{\mathrm{sc},ak})}{\sum_{k'=1}^{K} \exp(l_{\mathrm{sc},ak'})},$$

where $p_{ak}^{\mathrm{sc}}$ is the predicted probability that single cell $a$ belongs to spatial domain $k$. The embeddings $Z_{\mathrm{st}}$ and $Z_{\mathrm{sc}}$ are therefore directly comparable and are used by the downstream two-pass single-cell domain inference and domain-constrained optimal transport modules.

#### 1.3 Domain-constrained optimal transport for cell-to-spot assignment

After learning the shared RNA-centered latent space, COMPASS assigns dissociated single cells to spatial spots through two coupled steps. First, spatial-domain labels are inferred for dissociated single cells using a two-pass strategy that reduces the effect of class imbalance and preserves rare cell type–domain combinations. Second, the retained cells are assigned to candidate spatial spots by solving domain-constrained and cell-type-stratified optimal transport problems in the shared latent space. This section gives the detailed formulation of these two steps while following the notation used in the main text.

Let $Z_{\text{st}} \in \mathbb{R}^{n\times d_z}$ denote the spatial RNA embedding obtained from the distilled RNA backbone, where $n$ is the number of spatial spots and $d_z$ is the latent dimension. The spatial-domain labels inferred for spots are denoted by

$$\widehat{\mathbf{y}} = (\widehat{y}_1, \ldots, \widehat{y}_n), \qquad \widehat{y}_j \in \{1, \ldots, K\},$$

where $K$ is the number of spatial domains and $j$ indexes spatial spots. Let $X^{\text{sc}} \in \mathbb{R}^{N\times g_{\text{sc}}}$ denote the dissociated single-cell RNA-seq matrix for $N$ cells, and let $c_a \in \mathcal{T}$ denote the biological cell-type annotation of single cell $a$, where $\mathcal{T}$ is the set of cell types and $a = 1, \ldots, N$. The single-cell RNA matrix is first aligned to the same spatial RNA template used for representation learning. Specifically, let $G_{\text{st}}$ be the ordered gene set of the spatial RNA template, and let $\mu_g^{\text{st}}$ and $\sigma_g^{\text{st}}$ be the spatial-RNA-derived mean and standard deviation for gene $g \in G_{\text{st}}$. For each single cell $a$ and template gene $g$, the standardized input is

$$\widetilde{x}_{ag}^{\text{sc}} = \frac{\bar{x}_{ag}^{\text{sc}} - \mu_g^{\text{st}}}{\sigma_g^{\text{st}} + \varepsilon_\sigma},$$

where $\varepsilon_\sigma > 0$ is a numerical stabilizer. Here $\bar{x}_{ag}^{\text{sc}} = x_{ag}^{\text{sc}}$ if gene $g$ is present in the single-cell dataset, and $\bar{x}_{ag}^{\text{sc}} = \mu_g^{\text{st}}$ if gene $g$ is absent, so that missing template genes have standardized value zero. Let

$$\widetilde{X}_{\text{sc}} \in \mathbb{R}^{N\times d_{\text{rna}}}$$

denote the resulting low-dimensional single-cell RNA feature matrix after applying the same spatial-RNA-fitted dimension-reduction transformation used to construct $\widetilde{X}_{\text{st}}$.

The first-pass inference provides provisional spatial-domain probabilities for all single cells. To make this step scalable, COMPASS uses an identity propagation matrix and therefore performs no explicit cell–cell message passing in the first pass. Let $I_N \in \mathbb{R}^{N\times N}$ denote the identity matrix. Using the same RNA encoder weights $W_{\text{rna}}^{(1)}$ and $W_{\text{rna}}^{(2)}$ as in the main graph encoder, the first-pass single-cell embedding is

$$Z_{\text{sc}}^{(1)} = I_N \cdot \sigma\left(I_N \widetilde{X}_{\text{sc}} W_{\text{rna}}^{(1)}\right) \cdot W_{\text{rna}}^{(2)} = \sigma\left(\widetilde{X}_{\text{sc}} W_{\text{rna}}^{(1)}\right) W_{\text{rna}}^{(2)},$$

where $\sigma(\cdot)$ denotes the ReLU activation. Let $C_{\text{student}} : \mathbb{R}^{d_z} \to \mathbb{R}^K$ denote the deployable RNA-domain classifier learned by the student branch. The first-pass logits and probabilities are

$$l_{\text{sc}}^{(1)} = C_{\text{student}}\left(Z_{\text{sc}}^{(1)}\right),$$

$$p_{ak}^{(1)} = \frac{\exp\left(l_{\mathrm{sc},ak}^{(1)}\right)}{\sum_{k'=1}^{K}\exp\left(l_{\mathrm{sc},ak'}^{(1)}\right)}, \qquad a = 1,\dots,N, \quad k = 1,\dots,K.$$

The provisional domain label of cell $a$ is

$$\widetilde{y}_a^{(1)} = \arg\max_{k\in\{1,\dots,K\}} p_{ak}^{(1)}.$$

These first-pass predictions are used only to estimate the coarse domain composition of the single-cell population and to guide balanced subset construction; they are not treated as the final cell-domain labels.

To reduce domination by abundant cell populations, COMPASS constructs a representative subset by stratifying cells according to the Cartesian product of biological cell type and first-pass predicted domain. For cell type $t \in \mathcal{T}$ and provisional domain $k \in \{1,\dots,K\}$, define the joint bucket

$$\mathcal{B}_{t,k} = \left\{a : c_a = t,\ \widetilde{y}_a^{(1)} = k\right\}, \qquad A_{t,k} = |\mathcal{B}_{t,k}|.$$

Let

$$A_{t,\cdot} = \sum_{k=1}^{K} A_{t,k}, \qquad A_{\cdot,k} = \sum_{t\in\mathcal{T}} A_{t,k}$$

denote the observed cell-type and provisional-domain marginal counts. COMPASS specifies a cell-type row target

$$R_t = \min\left(M_t,\ A_{t,\cdot}\right),$$

where $M_t$ is the maximum number of retained cells allowed for cell type $t$. In the default implementation $M_t = M$ is shared across cell types. The total target subset size is

$$R_{\mathrm{tot}} = \sum_{t\in\mathcal{T}} R_t.$$

To avoid losing minority provisional domains, a domain column target $D_k$ is also defined. A default balanced choice is

$$D_k = \min\left(A_{\cdot,k},\ \left\lceil \frac{R_{\mathrm{tot}}}{K} \right\rceil\right),$$

followed by feasibility adjustment so that the final quotas do not exceed bucket availability. The initial quota for bucket $(t,k)$ preserves the within-cell-type domain composition:

$$q_{t,k}^{(0)} = R_t \frac{A_{t,k}}{A_{t,\cdot} + \varepsilon_q},$$

where $\varepsilon_q > 0$ prevents division by zero. The implemented quota matrix

$$Q = \{Q_{t,k}\}_{t\in\mathcal{T},\,k=1}^{K}$$

is an integer approximation of $q^{(0)}$ that satisfies

$$0 \le Q_{t,k} \le A_{t,k}, \qquad Q_{t,k} \in \mathbb{Z},$$

and approximately matches both the row targets and column targets:

$$\sum_{k=1}^{K} Q_{t,k} \approx R_t, \qquad \sum_{t\in\mathcal{T}} Q_{t,k} \approx D_k.$$

Equivalently, the quota adjustment can be viewed as a constrained integer projection,

$$Q \approx \arg\min_{\{q_{t,k}\}} \left\{ \sum_{t\in\mathcal{T}} \left( \sum_{k=1}^{K} q_{t,k} - R_t \right)^2 + \eta \sum_{k=1}^{K} \left( \sum_{t\in\mathcal{T}} q_{t,k} - D_k \right)^2 + \xi \sum_{t\in\mathcal{T}} \sum_{k=1}^{K} \left( q_{t,k} - q_{t,k}^{(0)} \right)^2 \right\},$$

subject to $0 \le q_{t,k} \le A_{t,k}$ and $q_{t,k} \in \mathbb{Z}$. Here $\eta \ge 0$ and $\xi \ge 0$ control the relative importance of matching provisional-domain balance and preserving the observed within-cell-type composition. In practice, this projection is implemented by rounding and feasibility adjustment. Within each bucket $\mathcal{B}_{t,k}$, cells are ranked by their first-pass confidence for domain $k$,

$$s_a^{(1)} = p_{ak}^{(1)}, \qquad a \in \mathcal{B}_{t,k},$$

and the $Q_{t,k}$ cells with the largest $s_a^{(1)}$ are retained. The selected representative cell set is denoted by

$$\mathcal{B}_{\text{map}} = \bigcup_{t\in\mathcal{T}} \bigcup_{k=1}^{K} \mathcal{B}_{t,k}^{\text{sel}}, \qquad N_{\text{map}} = |\mathcal{B}_{\text{map}}|,$$

where $\mathcal{B}_{t,k}^{\text{sel}} \subseteq \mathcal{B}_{t,k}$ is the selected subset from bucket $(t,k)$.

The second-pass inference refines domain predictions on the balanced representative subset. Let

$$\widetilde{X}_{\text{sc}}^{\text{map}} \in \mathbb{R}^{N_{\text{map}} \times d_{\text{rna}}}$$

denote the aligned low-dimensional RNA features of cells in $\mathcal{B}_{\text{map}}$. A $k$-nearest-neighbor graph is constructed among these selected cells in the aligned RNA feature space. Let

$$A_{\text{sc}}^{\text{map}} \in \mathbb{R}^{N_{\text{map}} \times N_{\text{map}}}$$

denote the symmetrized binary adjacency matrix of this graph. Its normalized propagation matrix is

$$\widetilde{A}_{\text{sc}}^{\text{map}} = (D_{\text{sc}}^{\text{map}})^{-1/2} \, (A_{\text{sc}}^{\text{map}} + I) \, (D_{\text{sc}}^{\text{map}})^{-1/2},$$

where $I$ is the identity matrix and $D_{\text{sc}}^{\text{map}}$ is the diagonal degree matrix of $A_{\text{sc}}^{\text{map}} + I$. The refined single-cell embedding is

$$Z_{\text{sc}}^{(2)} = \widetilde{A}_{\text{sc}}^{\text{map}} \cdot \sigma\left( \widetilde{A}_{\text{sc}}^{\text{map}} \widetilde{X}_{\text{sc}}^{\text{map}} W_{\text{rna}}^{(1)} \right) \cdot W_{\text{rna}}^{(2)}.$$

The second-pass logits and probabilities are

$$l_{\text{sc}}^{(2)} = C_{\text{student}}\left( Z_{\text{sc}}^{(2)} \right),$$

$$p_{ak}^{(2)} = \frac{\exp\left( l_{\text{sc},ak}^{(2)} \right)}{\sum_{k'=1}^{K} \exp\left( l_{\text{sc},ak'}^{(2)} \right)}, \qquad a \in \mathcal{B}_{\text{map}}, \quad k = 1, \ldots, K.$$

The refined domain label is

$$\widetilde{y}_a^{(2)} = \arg\max_{k \in \{1,\ldots,K\}} p_{ak}^{(2)}.$$

To prevent rare domains from disappearing after graph-based refinement, COMPASS applies a rare-domain rescue step. Let

$$N_k^{(2)} = \sum_{a \in \mathcal{B}_{\text{map}}} \mathbf{1}\left(\widetilde{y}_a^{(2)} = k\right)$$

be the number of selected cells assigned to domain $k$ after the second pass, and let $m_k^{\min}$ be the minimum desired number of cells for domain $k$. If $N_k^{(2)} < m_k^{\min}$, candidate cells that were provisionally assigned to domain $k$ in the first pass are revisited:

$$\mathcal{R}_k^{\text{cand}} = \left\{a \in \mathcal{B}_{\text{map}} : \widetilde{y}_a^{(1)} = k,\ \widetilde{y}_a^{(2)} \neq k\right\}.$$

These candidates are ranked by their first-pass confidence $p_{ak}^{(1)}$, and the top

$$r_k = \max\left(0,\ m_k^{\min} - N_k^{(2)}\right)$$

cells are relabeled to domain $k$, subject to availability. Let $\mathcal{R}_k \subseteq \mathcal{R}_k^{\text{cand}}$ denote the selected rescue set. The final single-cell domain label is

$$\widehat{y}_a^{\text{sc}} = \begin{cases} k, & a \in \mathcal{R}_k \text{ for some } k, \\ \widetilde{y}_a^{(2)}, & \text{otherwise.} \end{cases}$$

The refined embedding used for downstream transport is denoted by

$$Z_{\text{sc}} = Z_{\text{sc}}^{(2)} \in \mathbb{R}^{N_{\text{map}} \times d_z}.$$

Thus, $Z_{\text{st}}$ and $Z_{\text{sc}}$ are directly comparable in the shared RNA-centered latent space, while $\widehat{y}_j$ and $\widehat{y}_a^{\text{sc}}$ provide spot-domain and cell-domain labels for domain-restricted mapping.

Cell-to-spot mapping is performed separately within domain and cell-type buckets. For domain $k$ and cell type $t$, define the selected single-cell set

$$I_{k,t} = \{a \in \mathcal{B}_{\text{map}} : \widehat{y}_a^{\text{sc}} = k,\ c_a = t\}, \qquad n_{k,t} = |I_{k,t}|,$$

and the candidate spatial spot set

$$J_k = \{j \in \{1,\ldots,n\} : \widehat{y}_j = k\}, \qquad m_k = |J_k|.$$

Only buckets with $n_{k,t} > 0$ and $m_k > 0$ are used for optimal transport. Let

$$Z_{\text{sc}}^{k,t} = \{Z_{\text{sc},a} : a \in I_{k,t}\} \in \mathbb{R}^{n_{k,t} \times d_z}, \qquad Z_{\text{st}}^{k} = \{Z_{\text{st},j} : j \in J_k\} \in \mathbb{R}^{m_k \times d_z}$$

denote the corresponding cell and spot embeddings. For $a \in I_{k,t}$ and $j \in J_k$, the transport cost is defined by cosine distance in the shared RNA latent space:

$$C_{aj}^{(k,t)} = 1 - \frac{Z_{\text{sc},a}^{\top} Z_{\text{st},j}}{\|Z_{\text{sc},a}\|_2 \, \|Z_{\text{st},j}\|_2 + \varepsilon_C},$$

where $\varepsilon_C > 0$ is a numerical stabilizer. This cost favors assigning cells to spots with similar latent molecular states while restricting candidate matches to spatially compatible domains.

For each $(k, t)$ bucket, COMPASS solves an entropy-regularized balanced optimal transport problem[23]. Let

$$P^{(k,t)} \in \mathbb{R}_+^{n_{k,t} \times m_k}$$

be the transport plan, where $P_{aj}^{(k,t)}$ represents the amount of mass transported from cell $a$ to spot $j$ within the bucket. Let

$$\mu^{(k,t)} \in \mathbb{R}_+^{n_{k,t}}, \qquad \nu^{(k,t)} \in \mathbb{R}_+^{m_k}$$

be the prescribed cell and spot marginals satisfying

$$\sum_{a \in I_{k,t}} \mu_a^{(k,t)} = \sum_{j \in J_k} \nu_j^{(k,t)}.$$

Following the formulation in the main text, the optimal transport plan is

$$P^{(k,t)*} = \arg\min_{P \geq 0} \left\{ \left\langle P, C^{(k,t)} \right\rangle + \epsilon \sum_{a=1}^{n_{k,t}} \sum_{j=1}^{m_k} P_{aj} \left( \log P_{aj} - 1 \right) \right\}$$

subject to

$$P \mathbf{1}_{m_k} = \mu^{(k,t)}, \qquad P^\top \mathbf{1}_{n_{k,t}} = \nu^{(k,t)}.$$

Here $\epsilon > 0$ is the entropic regularization parameter, $\mathbf{1}_q$ denotes an all-one vector of length $q$, and

$$\left\langle P, C^{(k,t)} \right\rangle = \sum_{a=1}^{n_{k,t}} \sum_{j=1}^{m_k} P_{aj} C_{aj}^{(k,t)}.$$

In the default balanced setting, each cell carries one unit of mass and the spot marginal is uniform within the candidate domain:

$$\mu^{(k,t)} = \mathbf{1}_{n_{k,t}}, \qquad \nu^{(k,t)} = \frac{n_{k,t}}{m_k} \mathbf{1}_{m_k}.$$

Other spot marginals can be used when external spot-capacity priors are available.

The entropy-regularized problem is solved by Sinkhorn scaling. Define the Gibbs kernel

$$K_{aj}^{(k,t)} = \exp\left( -\frac{C_{aj}^{(k,t)}}{\epsilon} \right).$$

Starting from a positive vector $v^{(0)} \in \mathbb{R}_+^{m_k}$, the scaling vectors are updated as

$$u^{(r+1)} = \mu^{(k,t)} \oslash \left( K^{(k,t)} v^{(r)} \right),$$

$$v^{(r+1)} = \nu^{(k,t)} \oslash \left( \left( K^{(k,t)} \right)^\top u^{(r+1)} \right),$$

where $r$ is the Sinkhorn iteration index and $\oslash$ denotes element-wise division. The transport plan is reconstructed as

$$P^{(k,t)*} = \mathrm{diag}(u) K^{(k,t)} \mathrm{diag}(v).$$

The resulting soft plan satisfies the prescribed cell and spot marginals up to numerical tolerance and provides a probabilistic cell-to-spot assignment within each domain and cell-type bucket.

The soft transport plan is first used to estimate spot-level cell-type composition. For spot $j \in J_k$, the expected transported mass contributed by cell type $t$ is

$$s_{j,t}^{(k)} = \sum_{a \in I_{k,t}} P_{aj}^{(k,t)\,*}.$$

Aggregating over cell types gives the total expected load on spot $j$,

$$s_{j,\cdot}^{(k)} = \sum_{t \in \mathcal{T}} s_{j,t}^{(k)}.$$

A soft spot-level cell-type proportion can then be obtained as

$$\rho_{j,t} = \frac{s_{j,t}^{(k)}}{\sum_{t' \in \mathcal{T}} s_{j,t'}^{(k)} + \varepsilon_\rho}, \qquad j \in J_k,$$

where $\varepsilon_\rho > 0$ prevents division by zero. This soft representation preserves uncertainty in the transport plan and is useful for evaluating cell-type composition before discrete assignment.

For coordinate reconstruction, COMPASS converts the soft transport plan into a discrete one-cell-one-spot assignment. Within each $(k, t)$ bucket, define the expected spot load

$$\ell_j^{(k,t)} = \sum_{a \in I_{k,t}} P_{aj}^{(k,t)\,*}, \qquad j \in J_k.$$

Since $\sum_{j \in J_k} \ell_j^{(k,t)} = n_{k,t}$ under the default unit cell marginal, these real-valued loads are converted into integer spot capacities by largest-remainder rounding[4]. Let

$$b_j^{(k,t)} = \left\lfloor \ell_j^{(k,t)} \right\rfloor, \qquad r_j^{(k,t)} = \ell_j^{(k,t)} - b_j^{(k,t)}.$$

The remaining number of cells to allocate is

$$R^{(k,t)} = n_{k,t} - \sum_{j \in J_k} b_j^{(k,t)}.$$

The final integer capacity is

$$h_j^{(k,t)} = b_j^{(k,t)} + \mathbf{1}\left(j \in \mathcal{H}_{k,t}\right),$$

where $\mathcal{H}_{k,t}$ is the set of $R^{(k,t)}$ spots with the largest residuals $r_j^{(k,t)}$. Thus

$$h_j^{(k,t)} \in \mathbb{Z}_{\geq 0}, \qquad \sum_{j \in J_k} h_j^{(k,t)} = n_{k,t}.$$

Each spot $j \in J_k$ is expanded into $h_j^{(k,t)}$ identical assignment slots. Let $\mathcal{S}_{k,t}$ denote the resulting set of slots, and let

$$\psi : \mathcal{S}_{k,t} \to J_k$$

map each slot back to its original spot. A capacity-constrained Hungarian assignment is solved on the expanded bipartite graph. The hard-assignment cost of assigning cell $a \in I_{k,t}$ to slot $q \in \mathcal{S}_{k,t}$ is

$$H_{aq}^{(k,t)} = -\log\left(P_{a,\psi(q)}^{(k,t)\,*} + \delta\right),$$

where $\delta > 0$ is a small constant. The binary assignment matrix $A^{(k,t)} = \{A_{aq}^{(k,t)}\}$ is obtained by

$$A^{(k,t)*} = \arg \min_{A \in \{0,1\}^{n_{k,t} \times |\mathcal{S}_{k,t}|}} \sum_{a \in I_{k,t}} \sum_{q \in \mathcal{S}_{k,t}} A_{aq} H_{aq}^{(k,t)}$$

subject to

$$\sum_{q \in \mathcal{S}_{k,t}} A_{aq} = 1, \qquad \forall a \in I_{k,t},$$

$$\sum_{a \in I_{k,t}} A_{aq} = 1, \qquad \forall q \in \mathcal{S}_{k,t}.$$

The discrete spot assignment for cell $a$ is then

$$\pi(a) = \psi(q) \quad \text{where} \quad A_{aq}^{(k,t)*} = 1.$$

Repeating this procedure over all valid $(k,t)$ buckets yields a one-cell-one-spot assignment map

$$\pi : \mathcal{B}_{\text{map}} \to \{1, \ldots, n\}.$$

In memory-efficient execution, COMPASS does not need to keep all full transport matrices simultaneously. For soft composition estimation, it is sufficient to store the bucket-wise column sums $s_{j,t}^{(k)}$. For hard assignment, the relevant bucket-level transport matrix can be computed or reconstructed locally before capacity rounding and Hungarian matching. This preserves the same soft-then-hard transport formulation while reducing memory usage for large single-cell references.

Overall, the two-pass inference stage first estimates single-cell domain structure over the full scRNA-seq reference and then refines the mapping set on a balanced, confidence-enriched representative manifold. The optimal transport stage then performs cell-to-spot assignment only within compatible spatial domains and cell-type strata. By combining quota-based balancing, graph-refined single-cell domain inference, entropy-regularized balanced transport, and capacity-constrained hardening, COMPASS reduces cross-domain mismatches, prevents severe spot crowding, and produces the discrete assignment map required for subsequent sub-spot coordinate reconstruction.

#### 1.4 Poisson-disk geometric refinement for sub-spot coordinate reconstruction

After the domain-constrained optimal transport step and the subsequent capacity-constrained hard assignment, each mapped single cell is assigned to one spatial transcriptomics (ST) spot. Let

$$\mathcal{A} = \{1, \ldots, N\}$$

denote the set of mapped cells, where $N$ is the total number of cells retained after assignment. The hard cell-to-spot assignment is represented by

$$\pi : \mathcal{A} \to \{1, \ldots, n\},$$

where $n$ is the number of ST spots and $\pi(a) = j$ indicates that cell $a$ is assigned to spot $j$. This discrete assignment determines the spot membership of each cell, but it does not determine the continuous sub-spot coordinate of the cell. Therefore, Poisson-Disk Geometric Refinement (PDGR) is applied to convert the spot-level assignment into a cell-resolved spatial atlas while keeping the assignment $\pi$ fixed.

Let

$$S = \{s_j\}_{j=1}^{n}, \qquad s_j \in \mathbb{R}^2,$$

be the observed ST spot coordinates. For each spot $j$, the number of cells assigned to that spot is

$$n_j = \sum_{a=1}^{N} \mathbb{I}\{\pi(a) = j\},$$

where $\mathbb{I}\{\cdot\}$ denotes the indicator function. The set of occupied spots is defined as

$$\mathcal{J} = \{j : n_j > 0\}.$$

Thus,

$$N = \sum_{j \in \mathcal{J}} n_j.$$

The goal of PDGR is to estimate a continuous coordinate

$$\hat{s}_a \in \mathbb{R}^2, \qquad a \in \mathcal{A},$$

for every mapped cell, subject to the requirement that each cell remains associated with its assigned spot.

The geometric scale of the refinement is determined from the spatial organization of the occupied ST spots. For each occupied spot $j \in \mathcal{J}$, define the local nearest-neighbor spacing as

$$L_j^{(1)} = \min_{\substack{j' \in \mathcal{J} \\ j' \neq j}} \|s_j - s_{j'}\|_2.$$

The global characteristic spacing is then defined as

$$L = \mathrm{median}_{j \in \mathcal{J}}\, L_j^{(1)}.$$

Based on these spatial scales, PDGR introduces three scale-dependent parameters,

$$r_j = \rho_r L_j^{(1)}, \qquad d_{\min} = \rho_d L, \qquad R = \rho_R L,$$

where $\rho_r > 0$, $\rho_d > 0$, and $\rho_R > 0$ are scalar hyperparameters. The radius $r_j$ controls the maximum allowed displacement of cells assigned to spot $j$ from the corresponding spot center, $d_{\min}$ specifies the minimum separation between reconstructed single-cell coordinates, and $R$ determines the size of the spot-supported tissue region used for candidate generation.

For any point $c \in \mathbb{R}^2$ and radius $R > 0$, let

$$\mathcal{D}(c, R) = \{p \in \mathbb{R}^2 : \|p - c\|_2 \leq R\}$$

denote the closed Euclidean disk centered at $c$. The global support region for coordinate reconstruction is defined as the union of disks centered at occupied ST spots:

$$\Omega = \bigcup_{j \in \mathcal{J}} \mathcal{D}(s_j, R).$$

This construction restricts reconstructed cell coordinates to regions supported by the observed spatial measurements, rather than allowing cells to be placed arbitrarily on the tissue plane.

Within $\Omega$, PDGR first generates an overcomplete pool of candidate coordinates,

$$Q = \{q_\ell\}_{\ell=1}^{M} \subset \Omega,$$

with $M$ chosen to be larger than the total number of mapped cells. The candidate pool is generated using Bridson's Poisson-disk sampling procedure[5] with exclusion radius $d_{\min}$. Starting from an initial seed point $q_1 \in \Omega$, the algorithm iteratively proposes new candidate points around active points. For an active point $q \in Q$, a proposed point is sampled from the annular region

$$q_{\text{new}} \sim \text{Unif}\left(\left\{p \in \mathbb{R}^2 : d_{\min} \le \|p - q\|_2 < 2d_{\min}\right\} \cap \Omega\right).$$

The proposed point is accepted only if it lies inside the support region and is sufficiently separated from all previously accepted candidates:

$$q_{\text{new}} \in \Omega, \qquad \min_{q' \in Q} \|q_{\text{new}} - q'\|_2 \ge d_{\min}.$$

The procedure terminates when no additional valid candidates can be generated. The resulting candidate pool satisfies

$$\|q_\ell - q_{\ell'}\|_2 \ge d_{\min}, \qquad \ell \ne \ell',$$

and therefore provides a blue-noise-distributed set of possible single-cell locations inside the spot-supported tissue region.

Because $Q$ is overcomplete, PDGR next selects $N$ candidate coordinates and allocates them to occupied spots according to the capacities induced by the hard assignment. Let

$$x_{\ell j} \in \{0, 1\}$$

be a binary variable indicating whether candidate coordinate $q_\ell$ is selected and allocated to spot $j$. The allocation is formulated as

$$X^* = \arg\min_X \sum_{\ell=1}^{M} \sum_{j \in \mathcal{J}} x_{\ell j} \|q_\ell - s_j\|_2^2,$$

subject to

$$\sum_{\ell=1}^{M} x_{\ell j} = n_j, \qquad \forall j \in \mathcal{J},$$

$$\sum_{j \in \mathcal{J}} x_{\ell j} \le 1, \qquad \forall \ell = 1, \dots, M,$$

and

$$x_{\ell j} = 0 \quad \text{if} \quad \|q_\ell - s_j\|_2 > r_j.$$

The first constraint enforces capacity consistency by assigning exactly $n_j$ coordinates to spot $j$. The second constraint ensures that each candidate coordinate is used at most once. The third constraint imposes locality by preventing a candidate coordinate from being assigned to a spot if it lies outside the spot-specific radius $r_j$. Let

$$Q_j = \{q_\ell : x^*_{\ell j} = 1\}$$

be the set of coordinates allocated to spot $j$. Then

$$|Q_j| = n_j, \qquad \forall j \in \mathcal{J}.$$

Since the allocated coordinates are selected from the Poisson-disk candidate pool, the minimum-separation property among selected coordinates is retained.

After candidate coordinates have been allocated to spots, PDGR assigns these coordinates to individual cells within each spot. For each occupied spot $j$, define

$$\mathcal{A}_j = \{a \in \mathcal{A} : \pi(a) = j\}$$

as the set of cells assigned to spot $j$. By construction,

$$|\mathcal{A}_j| = |Q_j| = n_j.$$

Let $\mathcal{C}$ denote the set of cell types, and let

$$c_a \in \mathcal{C}$$

be the cell-type annotation of cell $a$. For each spot $j$ and cell type $t \in \mathcal{C}$, define

$$n_{j,t} = \sum_{a=1}^{N} \mathbb{I}\{\pi(a) = j,\ c_a = t\}$$

as the number of type-$t$ cells assigned to spot $j$. Thus,

$$n_j = \sum_{t \in \mathcal{C}} n_{j,t}.$$

To encourage local cell-type continuity, PDGR evaluates the expected cell-type composition around each allocated candidate coordinate using the compositions of neighboring spots. Let $\mathcal{N}(j) \subseteq \mathcal{J}$ denote the spatial neighborhood of spot $j$, defined from the spatial adjacency graph or nearest-neighbor graph of occupied spots. For a candidate coordinate $q_\ell \in Q_j$, the distance-based weight assigned to a neighboring spot $j' \in \mathcal{N}(j)$ is

$$w(q_\ell, j') = \frac{\exp\left(-\frac{\|q_\ell - s_{j'}\|_2^2}{2(L_j^{(1)})^2}\right)}{\sum_{j'' \in \mathcal{N}(j)} \exp\left(-\frac{\|q_\ell - s_{j''}\|_2^2}{2(L_j^{(1)})^2}\right)}.$$

The local affinity of coordinate $q_\ell$ to cell type $t$ is then

$$f_t(q_\ell) = \sum_{j' \in \mathcal{N}(j)} w(q_\ell, j') \frac{n_{j',t}}{n_{j'}}.$$

The vector

$$f(q_\ell) = \{f_t(q_\ell)\}_{t \in \mathcal{C}}$$

summarizes the expected cell-type composition in the spatial neighborhood of $q_\ell$.

For each spot $j$, the final cell-to-coordinate matching is obtained by a balanced linear assignment between the cells in $\mathcal{A}_j$ and the coordinates in $Q_j$. Let

$$\Sigma_j = \{\sigma_j : \mathcal{A}_j \to Q_j \mid \sigma_j \text{ is a bijection}\}$$

be the set of all one-to-one matchings between these cells and coordinates. Assigning cell $a$ to coordinate $q_\ell$ incurs the cost

$$C^{\text{loc}}_{a\ell} = 1 - f_{c_a}(q_\ell),$$

which is small when the neighborhood around $q_\ell$ is enriched for the cell type of cell $a$. The within-spot assignment is

$$\sigma_j^* = \arg\min_{\sigma_j \in \Sigma_j} \sum_{a \in \mathcal{A}_j} C^{\text{loc}}_{a,\sigma_j(a)}.$$

The reconstructed coordinate of each cell $a \in \mathcal{A}_j$ is then defined as

$$\hat{s}_a = \sigma_j^*(a), \qquad j = \pi(a).$$

The final coordinates satisfy the geometric constraints imposed by PDGR. Capacity consistency holds because each occupied spot $j$ receives exactly $n_j$ reconstructed cells:

$$|\{\hat{s}_a : \pi(a) = j\}| = n_j, \qquad \forall j \in \mathcal{J}.$$

Locality holds because every cell is placed within the spot-specific neighborhood of its assigned spot:

$$\|\hat{s}_a - s_{\pi(a)}\|_2 \le r_{\pi(a)}, \qquad \forall a \in \mathcal{A}.$$

Minimum separation holds because the final coordinates are selected from the Poisson-disk candidate pool:

$$\|\hat{s}_a - \hat{s}_b\|_2 \ge d_{\min}, \qquad a \ne b.$$

The type-coherent assignment further promotes continuity across local tissue neighborhoods by preferentially placing each cell near candidate coordinates whose surrounding spots are enriched for the same cell type.

The output of this refinement step is a reconstructed single-cell spatial atlas

$$\mathcal{T} = \{(\hat{s}_a, c_a, e_a, y_a)\}_{a=1}^{N},$$

where $\hat{s}_a \in \mathbb{R}^2$ is the reconstructed continuous coordinate of cell $a$, $c_a$ is its cell-type annotation, $e_a \in \mathbb{R}^g$ is its original single-cell gene expression profile over $g$ genes, and

$$y_a = \hat{y}_{\pi(a)}$$

is the spatial domain label inherited from the assigned ST spot.

### 2 Simulation design

#### 2.1 Robustness experiments

**Reference data and shared spatial generative framework** We constructed simulated spatial RNA and chromatin-accessibility (ATAC) datasets from a subsampled human melanoma single-cell multi-omics reference with jointly measured transcriptomic and epigenomic profiles[6]. The reference consisted of 2515 cells annotated into ten immune and malignant compartments. Spatial spots were arranged on a $24 \times 24$ square lattice with 576 spots and partitioned into eight structured sub-regions by tiling four quadrant-specific binary spatial masks. For each spot $s$, the number of contributing cells was sampled from a Conway–Maxwell–Poisson (CMP) distribution, $n_s \sim \mathrm{CMP}(\lambda_{r(s)}, \nu)$, where $r(s)$ denotes the region index, $\lambda_r$ is a region-specific rate parameter, and $\nu$ controls dispersion. Region-specific candidate cell-type sets were predefined, and cells were drawn from the reference in proportion to empirical type abundances using unbalanced sampling. Spot-level profiles were obtained by summing single-cell counts across modalities, and ground-truth cell-type proportions were recorded as $p_{s,c} = n_{s,c}/n_s$. Whenever a CMP draw yielded $n_s < 1$, we set $n_s \leftarrow 1$.

To emulate focal spatial heterogeneity, five axis-aligned rectangular areas on the lattice were selected, and all spots within them were resampled in a second round. For each such spot, the total cell count was drawn as an integer uniformly from $\{10, 11, \ldots, 20\}$. A dominant cell type was drawn uniformly from a permuted ordering of all atlas labels, and the number of dominant-type cells was set to $\lfloor \rho n_s \rfloor$ using Python-style truncation of $\rho n_s$ to an integer; remaining cells were assigned to other types by independent random draws with replacement. The expression for second-round spots was recomputed by aggregating the corresponding reference cells. After both rounds, the dominant label used for downstream attenuation was defined as the cell type with maximal $p_{s,c}$ at each spot.

For RNA, we introduced center-to-edge spatial technical variation using a spot-specific Poisson perturbation with baseline rate $\lambda_0$ and gradient coefficient $\gamma$,

$$\lambda_s = \lambda_0 \Big( 1 + \tfrac{d_s}{d_{\max}} (\gamma - 1) \Big) ,$$

where $d_s$ is the Euclidean distance of spot $s$ to the tissue centroid and $d_{\max}$ is the maximum spot–centroid distance. Poisson noise was added gene-wise; when zero protection was enabled, noise was applied only at entries that were non-zero before perturbation. Highly variable genes (HVGs) were computed separately for each reference cell type; at each spot, HVGs linked to the dominant type after both sampling rounds were multiplicatively downscaled for a randomly selected 80% subset of the top-ranked genes. Cell-resolution coordinates for evaluation were generated by sampling 7–13 cells per spot uniformly within a disk of fixed radius centred on each spot coordinate.

**Baseline matched simulation** The baseline setting applied the shared framework with modest compositional and technical perturbations. In the first round, each region was assigned a pool of cell types by randomly partitioning the global label ordering into contiguous blocks of size two to four depending on the region index, and all regions used $\lambda_r = 10$ with $\nu = 25$. The second-round compositionally skewed resampling used $\rho = 0.8$. RNA perturbation used $\lambda_0 = 15$ and $\gamma = 3$; ATAC received no additive Poisson background in this setting. HVG attenuation used the top 200 genes per dominant type and retained 20% of the original expression for the selected subset. No additional RNA-specific ambiguity mechanisms were applied beyond the shared steps above.

**Multi-omics complementarity under RNA-specific degradation** The complementarity setting used the same spatial generator but attenuated transcriptomic discriminability while leaving ATAC comparatively intact. Region–cell-type pools were specified explicitly so that predefined confusable pairs—$CD4^+$ T versus regulatory T, tumour subtype 1 versus tumour subtype 2, and mDC versus pDC—co-occurred across multiple regions. In the first round, $\lambda_r = 12$ for all regions and each region contained four to five permissible types. The second-round skew used $\rho = 0.5$. RNA perturbation was strengthened to $\lambda_0 = 35$ with $\gamma = 4$, and HVG attenuation used the top 300 genes with retained amplitude $0.08\times$ for the selected 80% subset per dominant type.

Three ordered RNA distortions were then applied to the spot-level RNA matrix only. First, for each confusable pair, genes were ranked by mean-expression separation between the two types in the reference atlas; additive Gaussian noise scaled by the non-zero spot-wise standard deviation per gene was applied, together with shrinkage toward the pairwise reference mean midpoint. Second, ambient-RNA-like background was added by scaling a global mean RNA profile to each spot in proportion to its total RNA signal. Third, for spots in which both members of a confusable pair exceeded 1% proportion, cross-type mixing was applied using pairwise reference mean profiles with additional stochastic residuals. ATAC spot matrices were not modified by these three steps. This design synthetically reduces RNA-based separability for predefined confusable pairs rather than implying loss of RNA information in real tissues generally.

**Complementary modality ablation and reduced reference size** Starting from the complementarity simulation outputs, we applied two further perturbations relevant to incomplete measurement and reference assembly. For spatial modality ablation, spots were ordered by increasing $x$ and then $y$; the lower half of spots by count along this ordering retained RNA signal only, and the upper half retained ATAC signal only, implemented by setting all RNA counts to zero in the upper half and all ATAC counts to zero in the lower half while leaving coordinates and stored proportion annotations unchanged. For reduced reference size, the single-cell RNA atlas was subsampled to 30% of cells within each type by stratified random sampling, retaining at least one cell per type. Analyses of this setting used the ablated spatial matrices together with the subsampled reference. Stochastic components of spatial simulation used fixed seeds for layout generation, second-round resampling, and noise injection as implemented in the released scripts; reference subsampling used a fixed seed for the 30% replicate.

#### 2.2 Ablation experiments

To further dissect which components drive the robustness of COMPASS, we performed a targeted ablation study under Scenario 4, the most challenging simulation setting in this study. This scenario combines three sources of perturbation: noisy spatial RNA measurements, partial loss of spot-level RNA signal, and dropout-induced incompleteness in the dissociated scRNA-seq reference. It therefore stresses all stages of the reconstruction pipeline simultaneously, including domain inference from corrupted spatial measurements, cross-omics transfer from auxiliary ATAC to the RNA backbone, cell-to-spot assignment under reference incompleteness, and sub-spot coordinate refinement. We ablated five functional blocks of COMPASS, including domain-prior construction, RG-COD training strategy, multi-objective loss design, OT-based cell-to-spot assignment, and PDGR-based geometric refinement. This design allows us to determine whether the performance gain comes from a single dominant module or from non-redundant contributions across representation learning, domain-constrained mapping, and geometric coordinate reconstruction.

**Auxiliary-aware domain priors anchor tissue structure under corrupted spatial measurements** We first examined the domain-prior module, because the inferred domains provide the upstream tissue-level structure used both to supervise representation learning and to constrain downstream cell-to-spot mapping. The domain ablation showed that auxiliary-aware domain construction was one of the most influential components. Removing the multi-omics signal by setting the auxiliary input to zero increased RMSE from 0.1439 to 0.2291, corresponding to a 59.2% increase, while PCC decreased from 0.6828 to 0.3025 and UQI dropped from 0.6553 to 0.1181. Replacing the full domain strategy with a naive multi-omics clustering variant caused a similarly large degradation, increasing RMSE by 50.5% and reducing PCC by 42.1%. These changes were much larger than the degradation caused by removing HMRF smoothing alone, which increased RMSE by only 4.8%. This pattern indicates that the main source of domain-level robustness is not merely spatial smoothing, but the construction of an informative multi-omics tissue partition. HMRF still contributed to spatial localization, as its ablation increased PCD from 101.70 to 115.14, suggesting that spatially coherent domains are particularly important for preserving coordinate-level consistency. Overall, the domain ablation supports the role of the domain prior as a structural anchor that prevents transcriptionally similar cells from being matched across spatially incompatible tissue regions.

**RG-COD teacher-student transfer preserves an RNA-deployable cross-omics representation** Given that the domain prior defines the spatial structure to be transferred, we next asked how RG-COD training transmits this structure into the RNA-deployable representation. Among the training ablations, removing the teacher branch caused the most pronounced failure. SSIM decreased from 0.4281 to 0.2472, a 42.3% reduction, PCC decreased by 19.8%, and PCD increased from 101.70 to 116.32. This confirms that the fused teacher is not a dispensable auxiliary head, but the main route through which ATAC-derived domain information is distilled into the RNA-only student. Ablating the gate, class weighting, linear projection, or graph structure caused milder but consistent losses, indicating that these components mainly stabilize how auxiliary information is projected, weighted, and injected into the RNA backbone. Notably, some variants such as the LayerNorm-related ablation improved several spot-level composition metrics, but still worsened PSD and PCD by more than 10%. This discrepancy shows that better spot-level proportions do not necessarily imply better single-cell localization. The full RG-COD design therefore provides a more balanced solution across composition recovery, domain consistency, and coordinate-level mapping accuracy.

**Reconstruction, domain supervision, and distillation define the core training signal** After establishing the importance of the RG-COD architecture, we further decomposed the loss function to determine which training signals are essential for maintaining this representation. The loss ablation showed a clear hierarchy. Individual regularizers, including decorrelation, graph smoothness, supervised contrastive learning, and center loss, mainly acted as stabilizing terms and produced moderate metric changes when removed individually. In contrast, disrupting the core reconstruction and supervision signals caused severe collapse. Removing the coupled RNA and ATAC reconstruction terms increased RMSE from 0.1439 to 0.2416, a 67.9% increase, reduced SSIM from 0.4281 to 0.1394, and nearly eliminated PCC, which dropped to 0.0205. A similarly strong failure occurred when fused, and RNA classification supervision was simultaneously removed. These results indicate that reconstruction maintains modality-specific structure, whereas domain classification aligns the learned space with biologically and spatially meaningful partitions. Knowledge distillation occupied an intermediate but important role: removing it increased RMSE to 0.1700, reduced SSIM to 0.3607, and increased JSD from 0.4335 to 0.4858. Thus, the objective function is not simply a collection of auxiliary penalties. Its core reconstruction, supervision, and distillation terms jointly enforce the cross-omics-to-RNA transfer required for mapping RNA-only dissociated cells.

**Balanced domain-cell-type OT prevents assignment collapse under reference incompleteness** We then evaluated the OT assignment block, which operates downstream of representation learning and determines whether the learned domain-aware embedding can be converted into reliable cell-to-spot correspondences. The strongest OT-stage failure occurred when balanced single-cell selection was ablated. RMSE increased from 0.1439 to 0.1652, PCC decreased from 0.6828 to 0.5698, and JSD increased from 0.4335 to 0.6521, corresponding to a 50.4% increase in distributional inconsistency. PCD also increased from 101.70 to 123.62. These changes indicate that balanced CellType-by-domain sampling is essential for preventing dominant cell types or high-confidence domains from overwhelming minority compartments under reference dropout. In contrast, replacing the second-pass kNN graph with an identity graph or removing domain augmentation caused smaller but consistent degradation, suggesting that graph-based refinement and domain replenishment mainly improve the stability of scRNA-side domain prediction before transport. Interestingly, the single-cell supplementary variant improved several composition-level metrics, including RMSE and PCC, but still worsened PCD. This again shows that composition recovery alone is insufficient for evaluating cell-resolved reconstruction. The OT ablation therefore suggests that the assignment module primarily controls distributional balance, mapping coverage, and the translation of representation quality into spatially valid cell placement.

**PDGR enforces locality and type-coherent geometry during sub-spot reconstruction** We finally examined the mapping and PDGR refinement block, which converts discrete spot memberships into continuous sub-spot coordinates. This stage addresses a different source of ambiguity from the representation and OT modules: even after a cell is assigned to the correct spot, its within-spot coordinate remains underdetermined. Ablating cell-type-aware placement caused the largest mapping-stage degradation, increasing RMSE from 0.1439 to 0.1802, a 25.2% increase, while reducing PCC from 0.6828 to 0.4710 and increasing PCD to 116.32. This indicates that type-aware placement is important not only for visual refinement but also for preserving local cell-type neighborhoods that determine spot-level composition. Removing coordinate clipping caused a different failure mode: JSD increased from 0.4335 to 0.5938 and PCD rose to 118.03, suggesting that locality constraints are needed to prevent cells from drifting away from their assigned spot-supported region. Replacing the union-of-disks support with a convex-hull support improved SSIM, but worsened UQI, JSD, and PCD, indicating that a more permissive global support can create smoother maps without preserving biologically plausible local geometry. Balanced hard assignment and domain restriction produced more moderate changes, consistent with their role in maintaining capacity consistency and spatial compatibility during the transition from soft transport to discrete cell placement. These results support PDGR as a necessary geometric refinement step rather than a cosmetic visualization procedure.

Across the five ablation groups, the most severe failures arose from disrupting auxiliary-aware domain construction, removing the RG-COD teacher pathway, or eliminating the coupled reconstruction and classification signals. These components form the representation-level core of COMPASS and explain why the gap between COMPASS and COMPASS-single widens when the spatial RNA signal deteriorates. OT and PDGR ablations produced more selective but biologically meaningful failures, especially in JSD, PSD, PCD, and local composition consistency, indicating that assignment and geometric refinement suppress downstream mapping artifacts that cannot be corrected by representation learning alone. The full model therefore achieves its strongest overall performance not because every component independently improves every metric, but because different modules constrain different degrees of freedom in an otherwise underdetermined reconstruction problem.

The runtime analysis further showed that these improvements were not obtained through computationally prohibitive variants. In Scenario 4, the full COMPASS pipeline completed in approximately 92 s, and most ablation variants remained within the same order of magnitude. Some simplified variants were faster because they removed training or mapping components, whereas a few variants incurred additional overhead due to less efficient optimization or assignment behavior. Overall, the ablation results support a non-redundant architecture in which domain-aware representation learning, RG-COD-based auxiliary transfer, balanced OT assignment, and PDGR coordinate refinement each suppress distinct failure modes while maintaining practical computational cost.

### 3 Description of compared methods

We compared COMPASS with five established spatial mapping methods and six spatial deconvolution methods. For the spatial mapping methods, we selected CMAP[2], CellTrek[7], CytoSPACE[8], SIMO[9], and Tangram[10]. CMAP efficiently maps large-scale individual cells to exact spatial locations by integrating single-cell and spatial data through a divide-and-conquer strategy, which involves spatial domain division, optimal spot alignment, and precise coordinate prediction. CellTrek achieves single-cell spatial mapping through co-embedding and metric learning approaches, utilizing a multivariate random forest model to construct a cell-spot distance matrix for spatial charting. CytoSPACE formulates the single-cell-to-spot assignment as a convex optimization problem, matching single-cell transcriptomes to spatial expression profiles via the shortest augmenting path algorithm to ensure a globally optimal alignment. SIMO is designed for the spatial integration of multi-omics datasets through probabilistic alignment, employing a fused Gromov-Wasserstein optimal transport framework to calculate the mapping relationship between cells and spots. Tangram uses a deep learning framework to map single-cell data onto spatial locations by optimizing the alignment between cell-type expression and spatial transcriptomics profiles. Beyond spatial mapping, we also assessed the spatial deconvolution capabilities of COMPASS by comparing it with six state-of-the-art deconvolution methods: CARD[11], RCTD[12], GraphST[13], Seurat[14], cell2location[15], and SPOTlight[16]. CARD adopts a conditional autoregressive (CAR) model to incorporate spatial smoothness, combining a regression-based deconvolution framework with spatial priors to borrow cell-type composition information across locations and improve robustness. RCTD models spatial transcriptomics counts using a Poisson mixture framework, where spot-level expression is represented as a weighted combination of reference cell-type profiles, with platform effects explicitly accounted for. GraphST utilizes a graph self-supervised contrastive learning method to learn informative spot representations by minimizing the embedding distance between spatially adjacent spots, facilitating accurate spatially informed cell-type deconvolution. Seurat leverages an anchor-based integration strategy, employing dimensionality reduction techniques to compute prediction scores that transfer cell-type annotations from reference single-cell data to spatial spots. Cell2location is based on a hierarchical Bayesian model that leverages negative binomial likelihoods and variational inference to jointly model cell-type abundance and technical effects. SPOTlight employs a seeded non-negative matrix factorization (NMF) regression approach to learn cell-type-specific topic profiles from single-cell RNA-seq data and subsequently deconvolute spatial transcriptomics spots into cell-type proportions.

All methods were implemented using their official software packages and executed following recommended guidelines whenever available. For the spatial mapping methods, CMAP was implemented using the CMAP R package (v1.0.0) with default parameters, except that `cluster_k` (the number of clusters) was matched to COMPASS, and the cell-filtering threshold was relaxed to 0.5 to retain more cells. CellTrek was implemented using the CellTrek R package (v0.0.94) following the authors' recommended settings, with the exception of setting `repel_r = NULL` for automatic repelling radius calculation (one-fourth of the spot distance). CytoSPACE (v1.1.0) was executed in Python using its official package with default parameters, with the solver method set to 'lapjv' and the sampling method set to 'duplicates'. SIMO (v1.0.0) was run in Python using the simo package with default settings, except that the Leiden clustering resolution was adjusted to align the cluster count with COMPASS for each dataset. Tangram (v1.0.4) was implemented in Python using `mode='constrained'` to map cells while respecting the provided cell density priors, without manual tuning of other default parameters. Regarding the spatial deconvolution methods, CARD was implemented using the CARD R package (v1.1) following the authors' recommended settings. RCTD was implemented using the spacexr R package (v2.2.1) with default parameters. GraphST (v1.0.0) was executed in Python, with the training epochs set to 1,200 to ensure optimal model convergence. Seurat was implemented using the Seurat R package (v4.3.0), employing its standard label transfer workflow with 30 principal components. Cell2location (v0.1.4) was executed in Python; its negative binomial regression model was trained for 250 epochs, and the spatial mapping model was trained for 30,000 epochs, with the expected number of cells per location adjusted based on the ratio of the number of cells to the number of spots. SPOTlight was implemented using the SPOTlight R package (v1.2.0), retaining marker genes with an AUC score greater than 0.5 and downsampling to 100 cells per cell type. To ensure a fair and rigorous comparison, the preprocessing steps for all aforementioned methods were kept strictly consistent with that of COMPASS.

### 4 Details on evaluations in simulations and real data sets

#### 4.1 External and internal clustering validation

To quantitatively assess the concordance between inferred spatial domains and ground-truth annotations, we adopted several external clustering validation metrics based on label agreement. Let $n$ denote the total number of spatial spots, and let $n_{ij}$ be the number of spots assigned to ground-truth class $i$ and inferred cluster $j$. The corresponding marginal sums are defined as $a_i = \sum_j n_{ij}$ and $b_j = \sum_i n_{ij}$. Each metric is calculated as follows:

**Adjusted Rand Index (ARI)**

$$\mathrm{ARI} = \frac{\sum_{ij}\binom{n_{ij}}{2} - \frac{\sum_i \binom{a_i}{2}\sum_j \binom{b_j}{2}}{\binom{n}{2}}}{\frac{1}{2}\left[\sum_i \binom{a_i}{2} + \sum_j \binom{b_j}{2}\right] - \frac{\sum_i \binom{a_i}{2}\sum_j \binom{b_j}{2}}{\binom{n}{2}}}$$

ARI measures the pairwise agreement between two partitions while correcting for random assignment. It ranges from $-1$ to 1, where higher values indicate stronger agreement between inferred spatial domains and ground-truth annotations.

**Mutual Information (MI)**

$$\mathrm{MI}(Y,\hat{Y}) = \sum_i \sum_j \frac{n_{ij}}{n} \log\left(\frac{n\, n_{ij}}{a_i b_j}\right)$$

MI quantifies the amount of shared information between inferred and reference labels. A larger MI indicates stronger dependency between the two partitions, although its magnitude is affected by the number of clusters and label distributions.

**Normalized Mutual Information (NMI)**

$$\mathrm{NMI}(Y,\hat{Y}) = \frac{\mathrm{MI}(Y,\hat{Y})}{\sqrt{H(Y)\,H(\hat{Y})}}$$

where

$$H(Y) = -\sum_i \frac{a_i}{n}\log\frac{a_i}{n}, \quad H(\hat{Y}) = -\sum_j \frac{b_j}{n}\log\frac{b_j}{n}$$

NMI rescales MI to the range $[0,1]$, enabling comparison across different clustering structures. Higher values indicate better alignment between inferred and ground-truth partitions.

**Adjusted Mutual Information (AMI)**

$$\mathrm{AMI}(Y,\hat{Y}) = \frac{\mathrm{MI}(Y,\hat{Y}) - \mathbb{E}[\mathrm{MI}(Y,\hat{Y})]}{\frac{1}{2}\big(H(Y)+H(\hat{Y})\big) - \mathbb{E}[\mathrm{MI}(Y,\hat{Y})]}$$

AMI corrects MI for chance agreement by subtracting its expected value under random labeling. It provides a more robust comparison across different clustering settings.

**Homogeneity**

$$\text{Homogeneity} = 1 - \frac{H(Y \mid \hat{Y})}{H(Y)}$$

Homogeneity evaluates whether each inferred cluster contains only samples from a single ground-truth class. Higher values indicate purer clusters.

**Completeness**

$$\text{Completeness} = 1 - \frac{H(\hat{Y} \mid Y)}{H(\hat{Y})}$$

Completeness evaluates whether all samples belonging to the same ground-truth class are assigned to the same inferred cluster. Higher values indicate better preservation of true class structure.

where the conditional entropies are defined as

$$H(Y \mid \hat{Y}) = -\sum_j \sum_i \frac{n_{ij}}{n} \log \frac{n_{ij}}{b_j}, \quad H(\hat{Y} \mid Y) = -\sum_i \sum_j \frac{n_{ij}}{n} \log \frac{n_{ij}}{a_i}.$$

To assess the intra-domain homogeneity of cell type compositions and inter-domain separation for inferred spatial domains without manual annotations, we adopted three complementary internal clustering validation metrics, using per-spot cell type proportions as input features and inferred spatial domains as cluster labels. Each metric is calculated as follows:

**Silhouette Coefficient**

$$\text{Silhouette Coefficient} = \frac{1}{N} \sum_{i=1}^{N} \frac{D_2(i) - D_1(i)}{\max\left(D_2(i), D_1(i)\right)}$$

where $N$ is the total number of spatial spots, $D_1(i)$ denotes the mean distance between spot $i$ and all other spots within its assigned domain, and $D_2(i)$ is the minimum mean distance between spot $i$ and spots in any other domain. This metric ranges from $-1$ to 1, with higher values indicating superior intra-domain compactness and inter-domain distinction.

**Calinski-Harabasz**

$$\text{CH} = \frac{\text{SS}_B/(k-1)}{\text{SS}_W/(N-k)}$$

where $\text{SS}_B$ is the sum of squared distances between each domain centroid and the global centroid, $\text{SS}_W$ is the sum of squared distances between each spot and its domain centroid, $k$ is the number of inferred spatial domains, and $N$ is the total number of spatial spots. A higher CH index corresponds to more well-separated and compact spatial domains.

**Davies-Bouldin**

$$\text{DB} = \frac{1}{k} \sum_{i=1}^{k} \max_{j \neq i} \left( \frac{\bar{s}_i + \bar{s}_j}{d(c_i, c_j)} \right)$$

where $k$ is the number of inferred spatial domains, $\bar{s}_i$ is the mean distance between spots in domain $i$ and the domain centroid, $c_i$ and $c_j$ are the centroids of domains $i$ and $j$, and

$d(c_i, c_j)$ is the distance between the two centroids. A lower DB index indicates better clustering performance with minimal overlap between distinct spatial domains.

#### 4.2 Spatial smoothness

To systematically assess the degree of spatial autocorrelation in the inferred embeddings, we computed Moran's $I$ and Geary's $C$. These indices quantify global and localized spatial dependencies, respectively, and are formulated as:

$$I = \frac{n}{\sum_{i=1}^{n}\sum_{j=1}^{n} w_{ij}} \frac{\sum_{i=1}^{n}\sum_{j=1}^{n} w_{ij}(x_i - \bar{x})(x_j - \bar{x})}{\sum_{i=1}^{n}(x_i - \bar{x})^2},$$

$$C = \frac{n-1}{2\sum_{i=1}^{n}\sum_{j=1}^{n} w_{ij}} \frac{\sum_{i=1}^{n}\sum_{j=1}^{n} w_{ij}(x_i - x_j)^2}{\sum_{i=1}^{n}(x_i - \bar{x})^2},$$

where $n$ represents the total number of spatial units, $x_i$ and $x_j$ denote the variable of interest at spots $i$ and $j$, and $\bar{x}$ is the mean of $x$. The spatial connectivity is encoded in a shared binary weight matrix, where the element $w_{ij} = 1$ if spot $i$ is contiguous to spot $j$, and $w_{ij} = 0$ otherwise (with $w_{ii} = 0$).

#### 4.3 Cell composition accuracy

To evaluate the performance of cell-type proportion reconstruction, we adopted three metrics to quantify the similarities and disparities between predicted and ground-truth cell-type proportion profiles, including RMSE, SSIM, and ERGAS. Each metric is calculated as follows:

**RMSE**

$$RMSE = \sqrt{\frac{1}{N}\sum_{i=1}^{N}(x_i - y_i)^2}$$

where $N$ is the total number of spatial spots, $x_i$ and $y_i$ are the ground-truth and predicted cell-type proportion at the $i$-th spot, respectively. This non-negative metric decreases with higher prediction accuracy.

**SSIM**

$$SSIM(x, y) = \frac{(2\mu_x\mu_y + c_1)(2\sigma_{xy} + c_2)}{(\mu_x^2 + \mu_y^2 + c_1)(\sigma_x^2 + \sigma_y^2 + c_2)}$$

where $\mu_x$, $\mu_y$, $\sigma_x^2$, $\sigma_y^2$ denote the mean and variance of the ground-truth $x$ and predicted $y$ profiles, respectively, and $\sigma_{xy}$ is the covariance of $x$ and $y$. Stabilization terms $c_1 = (k_1L)^2$ and $c_2 = (k_2L)^2$ adopt default parameters $k_1 = 0.01$ and $k_2 = 0.03$, where $L$ is the dynamic range of input values. SSIM ranges from $-1$ to 1, with 1 representing identical profiles and higher values indicating better structural pattern preservation.

**ERGAS**

$$ERGAS = 100 \cdot \frac{h}{l}\sqrt{\frac{1}{K}\sum_{k=1}^{K}\frac{RMSE(B_k)^2}{\bar{y}_k^2}}$$

where $K$ is the number of cell-type groups, $RMSE(B_k)$ is the RMSE of the $k$-th group, and $\bar{y}_k$ is the mean ground-truth proportion of the $k$-th group. The pixel size ratio $\frac{h}{l}$ between fused and reference images is set to 1.0 in this study. Lower ERGAS values correspond to superior reconstruction performance.

#### 4.4 Cell mapping coherency

To evaluate single-cell spatial mapping consistency and discriminative power of inferred spatial domains, we used three metrics based on mapped cell count distributions per spot: JSD, CV, and Objective score. Each metric is calculated as follows:

**JSD**

$$JSD(P_{d_1} \parallel P_{d_2}) = \frac{1}{2} D_{KL}(P_{d_1} \parallel M) + \frac{1}{2} D_{KL}(P_{d_2} \parallel M)$$

$$\text{Mean JSD} = \frac{2}{K(K-1)} \sum_{1 \le d_1 < d_2 \le K} JSD(P_{d_1} \parallel P_{d_2})$$

where $P_{d_1}, P_{d_2}$ are cell count frequency distributions of domains $d_1, d_2$, $M = \frac{1}{2}(P_{d_1} + P_{d_2})$, $D_{KL}(\cdot \parallel \cdot)$ is Kullback-Leibler divergence, and $K$ is the total number of inferred spatial domains. Higher Mean JSD indicates more distinct inter-domain cell mapping patterns.

**CV**

$$CV_d = \frac{\sigma_d}{\mu_d} \quad \text{Mean CV} = \frac{1}{K} \sum_{d=1}^{K} CV_d$$

where $\sigma_d$ and $\mu_d$ are the standard deviation and mean of mapped cell counts for spots in domain $d$, and $K$ is the total number of inferred spatial domains. Lower Mean CV indicates more homogeneous intra-domain cell distribution.

**Objective**

$$\text{Objective} = \text{Mean JSD} - \text{Mean CV}$$

This composite score jointly evaluates between-domain discriminability and within-domain uniformity. Higher values indicate superior performance, with distinct inter-domain patterns and consistent intra-domain cell allocation.

#### 4.5 Intra-domain count homogeneity

To assess the uniformity of mapped cell counts across spatial spots within inferred domains, we adopted the Mean Entropy metric based on Shannon entropy, calculated as:

$$H(d) = -\sum_{v \in V_d} p_{d,v} \log_2(p_{d,v}) \quad \text{Mean Entropy} = \frac{1}{K} \sum_{d=1}^{K} H(d)$$

where $H(d)$ is the Shannon entropy of domain $d$, $V_d$ is the set of unique cell count values in domain $d$, $p_{d,v}$ is the occurrence frequency of value $v$, and $K$ is the total number of inferred spatial domains. Higher values indicate a more uniform cell count distribution across spots, while lower values reflect a more skewed distribution.

### 5 Data description

**Mouse Olfactory Bulb Data** This study utilized the mouse olfactory bulb spatial transcriptomic data generated by the Spatial Transcriptomics v1.0 technology. Following previous studies, we focused on the mouse OB Section 12 for downstream analyses, which covers a total of 16034 genes across 282 spatial spots. The matched mouse OB single-cell RNA sequencing (scRNA-seq) reference dataset used in this study contains 18560 genes across 12801 cells, covering five major cell types: granule cells (GC, $n = 8614$), olfactory sensory neurons (OSNs, $n = 1200$), periglomerular cells (PGC, $n = 1693$), mitral and tufted cells (M-TC, $n = 1133$), and external plexiform layer interneurons (EPL-IN, $n = 161$).

**Mouse Sagittal Brain Data** This dataset comprises publicly available adult C57BL/6 mouse brain data from 10x Genomics, including sagittal and coronal sections. The anterior brain section consists of gene expression measurements in the form of read counts for 18449 genes and 2696 spatial locations, and the posterior section contains gene expression measurements in the form of read counts for 18496 genes and 3339 spatial locations. To perform the single-cell spatial mapping tasks, scRNA-seq data of the mouse brain were used as a reference, including 45768 gene measurements from 10530 cells. Specifically, 12 cell types with meaningful biological characterizations were selected, including astrocyte (Astro, $n = 556$), Cajal-Retzius cells (CR, $n = 17$), endothelial cell (Endo, $n = 166$), Layer 2/3 Intratelencephalic (L2/3 IT, $n = 1178$), Layer 4 (L4, $n = 1352$), Layer 5 Intratelencephalic (L5 IT, $n = 2964$), Layer 6 Corticothalamic Neurons (L6 CT, $n = 1225$), Layer 6 Intratelencephalic (L6 IT, $n = 2173$), Layer 6b Neurons (L6b, $n = 473$), Macrophage ($n = 131$), oligodendrocyte (Oligo, $n = 186$), and smooth muscle cell (SMC, $n = 109$).

**High-resolution Xenium data** This study used paired spatial omics data from adjacent FFPE breast cancer sections reported by Janesick et al., integrating Xenium and Visium data from GEO SuperSeries GSE243280, including GSE243168 and GSE243275. Xenium outputs included cell-level spatial coordinates, transcript detection results, morphology images, and metadata, while Visium outputs included gene expression profiles, spot coordinates, scale factors, and histology images. These data were used for spatial registration, coordinate harmonization, and integrative analysis. Following the 10x Genomics registration workflow, Xenium coordinates were transformed into the morphology image coordinate system, mapped to the Visium full-resolution space, and then converted to the H&E hires image coordinate system. Only the overlapping region between the two platforms was retained for downstream analyses. Specifically, the Visium spatial transcriptomic data consists of gene expression measurements for 18085 genes across 4992 spatial spots. The matched Xenium single-cell resolution dataset used as a reference contains 313 genes across 167780 cells, covering 20 cell groups: B Cells ($n = 4987$), CD4+ T Cells ($n = 8453$), CD8+ T Cells ($n = 6940$), DCIS 1 ($n = 12923$), DCIS 2 ($n = 11683$), Endothelial ($n = 8931$), IRF7+ DCs ($n = 494$), Invasive Tumor ($n = 34374$), LAMP3+ DCs ($n = 298$), Macrophages 1 ($n = 11174$), Macrophages 2 ($n = 1624$), Mast Cells ($n = 167$), Myoepi ACTA2+ ($n = 7078$), Myoepi KRT15+ ($n = 2860$), Perivascular-Like ($n = 847$), Prolif Invasive Tumor ($n = 3775$), Stromal ($n = 41422$), Stromal & T Cell Hybrid ($n = 607$), T Cell & Tumor Hybrid ($n = 589$), and Unlabeled ($n = 8554$). Furthermore, due to the limited number of genes captured in the single-cell data, we directly utilized the common genes shared between the two datasets for the analyses.

**Human lymph node data** This study utilized the human lymph node spatial multi-omics (RNA and protein) data generated by the 10x Genomics Visium CytAssist platform. Based on manual annotations, we focused on the spatially matched RNA and ADT data for downstream analyses, which covers a total of 18085 genes and 31 proteins across 3484 spatial spots. These spots encompass ten distinct anatomical regions: medulla cords ($n = 1188$), medulla sinuses ($n = 797$), cortex ($n = 753$), pericapsular adipose tissue ($n = 320$), capsule ($n = 204$), subcapsular sinus ($n = 74$), follicle ($n = 60$), medulla vessels ($n = 57$), hilum ($n = 23$), and trabeculae ($n = 8$). The matched human scRNA-seq reference dataset used in this study contains 10237 genes across 73260 cells, covering five major lymphatic tissue sources (Tonsil, mLN, Spleen, iLN, and tLN). For the purpose of this study, the cell types were merged into five major categories: B cells ($n = 38147$), T cells ($n = 30393$), Endothelial-Stromal cells ($n = 738$), Myeloid cells ($n = 1079$), and NK/ILC ($n = 2903$).

**Clinical breast cancer data** The dataset was obtained from the NCBI GEO database (GSE176078). We selected four representative primary breast cancer samples, encompassing two major clinical subtypes: the ER+ subtype and the TNBC subtype. The ER+ subtype includes two samples (sample IDs: CID4290, CID4465), representing oestrogen receptor-positive tumours with typical luminal characteristics. Specifically, the Visium spatial transcriptomic data for CID4290 consists of 19237 genes across 2426 spatial spots. Its matched scRNA-seq reference contains 29733 genes across 5789 cells, covering 8 cell types: B-cells ($n = 117$), CAFs ($n = 280$), Cancer Epithelial ($n = 4053$), Endothelial ($n = 298$), Myeloid ($n = 341$), Normal Epithelial ($n = 18$), PVL ($n = 140$), and T-cells ($n = 542$). For CID4465, the spatial data includes 18820 genes across 1211 spots, while the paired scRNA-seq data contains 29733 genes across 1564 cells, covering 9 cell types: B-cells ($n = 33$), CAFs ($n = 379$), Cancer Epithelial ($n = 124$), Endothelial ($n = 294$), Myeloid ($n = 181$), Normal Epithelial ($n = 10$), Plasmablasts ($n = 110$), PVL ($n = 317$), and T-cells ($n = 116$). The TNBC subtype comprises two samples (sample IDs: CID44971, CID4535), representing triple-negative breast cancer with high heterogeneity and a complex immune microenvironment. For CID44971, the spatial data consists of 18967 genes across 1162 spots, and the matched scRNA-seq reference encompasses 29733 genes across 7986 cells, including 9 cell types: B-cells ($n = 369$), CAFs ($n = 582$), Cancer Epithelial ($n = 894$), Endothelial ($n = 217$), Myeloid ($n = 684$), Normal Epithelial ($n = 735$), Plasmablasts ($n = 48$), PVL ($n = 91$), and T-cells ($n = 4366$). For CID4535, the spatial transcriptomic data captures 19237 genes across 1125 spots, and the paired scRNA-seq data comprises 29733 genes across 3961 cells, covering 9 cell types: B-cells ($n = 56$), CAFs ($n = 102$), Cancer Epithelial ($n = 2223$), Endothelial ($n = 219$), Myeloid ($n = 255$), Normal Epithelial ($n = 22$), Plasmablasts ($n = 96$), PVL ($n = 592$), and T-cells ($n = 396$).

### References

[1] Zhu Q, Shah S, Dries R, Cai L, Yuan GC. Identification of spatially associated subpopulations by combining scRNAseq and sequential fluorescence in situ hybridization data. Nature biotechnology. 2018;36(12):1183-90.

[2] Ke J, Xu J, Liu J, Yang Y, Guo C, Xie B, et al. High-resolution mapping of single cells in spatial context. Nature Communications. 2025;16(1):6533.

[3] Dai B, Yi L, Wang P, Li H, Hu P, Song Y, et al. 3d-OT: a deep geometry-aware framework for heterogeneous slices alignment of spatial multi-omics. Nature Methods. 2026:1-12.

[4] Pukelsheim F. Proportional representation. Springer; 2017.

[5] Bridson R. Fast Poisson disk sampling in arbitrary dimensions. SIGGRAPH sketches. 2007;10(1):1.

[6] Russell AJ, Weir JA, Nadaf NM, Shabet M, Kumar V, Kambhampati S, et al. Slide-tags enables single-nucleus barcoding for multimodal spatial genomics. Nature. 2024;625(7993):101-9.

[7] Vahid MR, Brown EL, Steen CB, Zhang W, Jeon HS, Kang M, et al. High-resolution alignment of single-cell and spatial transcriptomes with CytoSPACE. Nature biotechnology. 2023;41(11):1543-8.

[8] Wei R, He S, Bai S, Sei E, Hu M, Thompson A, et al. Spatial charting of single-cell transcriptomes in tissues. Nature biotechnology. 2022;40(8):1190-9.

[9] Yang P, Jin K, Yao Y, Jin L, Shao X, Li C, et al. Spatial integration of multi-omics single-cell data with SIMO. Nature communications. 2025;16(1):1265.

[10] Biancalani T, Scalia G, Buffoni L, Avasthi R, Lu Z, Sanger A, et al. Deep learning and alignment of spatially resolved single-cell transcriptomes with Tangram. Nature methods. 2021;18(11):1352-62.

[11] Ma Y, Zhou X. Spatially informed cell-type deconvolution for spatial transcriptomics. Nature biotechnology. 2022;40(9):1349-59.

[12] Cable DM, Murray E, Zou LS, Goeva A, Macosko EZ, Chen F, et al. Robust decomposition of cell type mixtures in spatial transcriptomics. Nature biotechnology. 2022;40(4):517-26.

[13] Long Y, Ang KS, Li M, Chong KLK, Sethi R, Zhong C, et al. Spatially informed clustering, integration, and deconvolution of spatial transcriptomics with GraphST. Nature communications. 2023;14(1):1155.

[14] Stuart T, Butler A, Hoffman P, Hafemeister C, Papalexi E, Mauck WM, et al. Comprehensive integration of single-cell data. cell. 2019;177(7):1888-902.

[15] Kleshchevnikov V, Shmatko A, Dann E, Aivazidis A, King HW, Li T, et al. Cell2location maps fine-grained cell types in spatial transcriptomics. Nature biotechnology. 2022;40(5):661-71.

[16] Elosua-Bayes M, Nieto P, Mereu E, Gut I, Heyn H. SPOTlight: seeded NMF regression to deconvolute spatial transcriptomics spots with single-cell transcriptomes. Nucleic acids research. 2021;49(9):e50-0.
